# Social interactions between people of same and different generations shape longitudinal changes in interpersonal neural synchrony, loneliness, and social connection

**DOI:** 10.1101/2025.10.14.682029

**Authors:** Ryssa Moffat, Guillaume Dumas, Emily S. Cross

## Abstract

Loneliness is globally acknowledged as a severe and burgeoning health risk, fuelling interest in helping people of all ages form meaningful social connections. One promising approach consists of intergenerational social programs. While behavioural and qualitative evidence derived from such programs promise health and wellbeing benefits, the physiological consequences of repeated intergenerational encounters remain unknown. Insight into physiological changes will shed light on the mechanisms of social connection. We charted longitudinal changes in interpersonal neural synchrony (INS) in 31 intergenerational (older/younger adult) and 30 same generation (younger adult) dyads across a six-session creative drawing program. At each session, dyads completed self-report measures, drew together and alone, and had their cortical activation recorded with fNIRS. In both groups, INS was greater while dyads drew together than alone. Across sessions, intergenerational dyads’ INS decreased and same generation dyads’ INS increased. INS in RIFG∼RTPJ and RIFG∼RIFG were predictive of loneliness levels and feelings of social closeness, respectively. This exploratory longitudinal research reinforces the multi-faceted nature of INS dynamics as social connections are forged.

## 1 Introduction

Experiences of loneliness can have serious negative consequences for physical and mental health. As a result, widespread efforts are being made to combat loneliness (1). According to meta-analyses of behavioural studies, community programs providing opportunities for repeated contact between generations can improve older adults’ (65+ years) physical, social, and cognitive health (2), while also reducing younger adults’ stereotypes about aging (3,4). Moreover, physicians are increasingly prescribing attendance at such programs–a practice called social prescribing (5)–further highlighting the value the medical field attributes to such programs. To date the existing evidence base for intergenerational community programs consists of behavioural and qualitative studies. While offering promising results in terms of self-reported benefits and social engagement, such studies offer little insight into how or why intergenerational relationships develop on a physiological level. In this study, we sought to provide a physiological perspective on the trajectory of intergenerational relationship development using a longitudinal hyperscanning approach, comparing intergenerational and same generation dyads who completed a six-week creative drawing program. Specifically, we tracked the trajectories of dyads’ interpersonal neural synchrony (INS) and changes in social connectedness.

The following sections briefly describe the benefits of intergenerational social interactions, what is known about INS in intergenerational interactions, and the benefits of longitudinal perspectives on INS. Following this, we detail the aims and hypotheses of the present study.

### 1.1 Intergenerational social interactions combat loneliness by offering social connections

Loneliness can be defined as “a negative, subjective emotional state resulting from a discrepancy between one’s desired and actual experiences of connection” (6). Put more succinctly, loneliness can be thought of as *perceived* social isolation. Policy makers and researchers have long emphasised the severe risk that loneliness poses to physical and mental health (1,7,8). Public figures and media outlets go so far as to compare the adverse health impacts (e.g., mortality) of loneliness to those of smoking cigarettes (9). Considering everyday functioning, longitudinal studies show that loneliness and social isolation can precipitate declines in cognitive functioning including verbal ability, numeracy, and short-term memory (10,11). Loneliness is also associated with reduced perceptual speed (12) and biases toward negative evaluations of social information (13–15). At the brain level, chronic loneliness and episodes of social isolation have been associated with altered structure (typically reduction in tissue volume) and altered function of prefrontal and posterior superior temporal cortices, as well as the insula, hippocampus, and amygdala (16–18). Attentional, visual, and default mode networks are also reported to show stronger functional connectivity in lonely individuals, which is believed to be associated with the bias toward processing social information negatively and a tendency toward increased rumination (18–20).

To combat loneliness, policy makers, researchers (1,21,22), and health practitioners (5) promote the creation of opportunities for meaningful social interactions for all members of society, and especially for those from populations most susceptible to loneliness. This typically involves initiating community programs that provide a platform for regular socialising opportunities (6,23–25). Some community programs are intentionally designed to bring together members of different age groups, i.e., to be intergenerational. One reason for this is that people of all ages experience loneliness (26–28). Another is that intergenerational social interactions can improve social and physical wellbeing for older adults, as well as increase empathy and feelings of social closeness toward older adults from younger participants (2). The observed improvements seem to stem from repeated intergenerational contact (29–31). In this context, repeated intergenerational contact requires regular engagement in intergenerational community programs, which is most likely to be achieved by ensuring that programs offer meaningful social interactions. One approach that has been shown to be particularly successful in this domain is arts-based and creative activities (32). Indeed, a growing body of evidence documents the social and physical health benefits of intergenerational arts programs for participants of all ages (30,33–35).

Despite the mounting behavioural and qualitative evidence that repeated intergenerational interactions yield social benefits, including reduced loneliness, little is known about the physiological aspects that contribute to *how* and *why* these intergenerational programs offer such benefits. This is important because the neurophysiological signatures of budding intergenerational relationships have the potential to be harnessed as tools to promote the formation of new social connections, yet they remain unknown (36).

### 1.2 Interpersonal neural synchrony (INS) during collaborative intergenerational interactions

One source of physiological information is brain activity. Recording brain activity from two or more people while they interact is commonly referred to as ‘hyperscanning’ and offers a measure of interpersonal neural synchrony (INS; the extent to which patterns of brain activity are temporally aligned). Mobile functional near-infrared spectroscopy (fNIRS) and electroencephalography (EEG) are particularly well suited to recording brain activity during social interactions in real-world settings (36,37). Hyperscanning with non-mobile techniques, such as functional magnetic resonance imaging (fMRI) and magneto-encephalography (MEG), is possible, though it offers less realistic social interactions and is less logistically feasible (38,39).

The role of INS in social behaviour remains debated (40–43), with two main theoretical conceptualisations: common cognitive processing and mutual prediction. The common cognitive processing framework posits that interacting people perceive similar stimuli, which results in both people’s brains responding in comparable ways at comparable time delays (44). This framework can explain why individuals viewing the same movie or simultaneously imitating an instructor may show heightened INS. Capturing the interactive and turn-based aspects of real social interactions, the mutual prediction framework posits that interacting people predict potential changes in the environment that they share, such as social information about each other, and that the interdependence of these continuous predictions leads to heightened INS (45–47). We embed our interpretation of INS during collaborative drawing in the mutual prediction framework, in that we would argue that collaborative drawing inherently involves different (even if complementary) stimuli for each participant created by the other.

#### 1.2.1 INS in social brain networks differs between solo and dyadic tasks

In studies comparing INS while dyads perform tasks collaboratively and alone, INS is consistently greater for collaborative performance than solo performance. This pattern has been observed for finger tapping (48), cognitive tasks such as n-back and object detection tasks (49,50), singing (51), tracing shapes on a screen using computer buttons or a stylus (52,53), drawing specific verbs as in Pictionary (54), among other activities. A meta-analysis of fNIRS hyperscanning studies by Czeszumski et al. (55) reports that during collaboration, INS is commonly observed between both interactants’ prefrontal cortex (PFC), inferior frontal cortex (IFG) or temporoparietal junction (TPJ). INS in the prefrontal and inferior frontal cortices is proposed to reflect initiation and reception of joint attention (56–58), a social competence that depends on cognitive control (59,60). INS in the TPJ, a key node of the Theory of Mind (ToM) network (61,62), is believed to reflect perspective taking and integration of social information (55). Complementary findings from EEG hyperscanning studies suggest that the strength and nature of collaboration shape the levels of INS recorded over frontal and prefrontal brain regions for alpha, beta, and theta frequency bands (63–67). Specifically, the alpha band may index joint attention and visual processing, while beta and theta bands may reflect joint encoding of sensorimotor information (37,63,64). In combination, the findings from fNIRS and EEG hyperscanning studies indicate that INS levels during collaboration are shaped by processes such as joint attention and perspective taking, which underpin the mutual prediction framework (40).

Many hyperscanning studies on collaboration can be considered intergenerational, in that they examine collaboration between parent-child/infant or student-teacher dyads (for reviews, see: ,68,69). Though these studies offer the field of social neuroscience a valuable starting point for understanding intergenerational collaboration, two gaps in the literature have yet to be filled: First, the hyperscanning literature on collaboration distinctly lacks any representation of older adults. Second, the hyperscanning literature more generally offers very few longitudinal perspectives into INS exceeding two time points. We now briefly explore the importance of addressing these gaps.

#### 1.2.2 Older adults are underrepresented in hyperscanning research

To date, older adults’ representation in the hyperscanning literature is limited to two hyperscanning studies (70,71), one study simulating INS (72), and one position piece arguing for the inclusion of older adults in such studies (36). It is critical to address this knowledge gap as the real-world settings in which older adults interact with younger adults and children (with their children and grandchildren, in assisted living, when seeking medical care, in community programs or through volunteer work) are numerous. Moreover, the proportion of older adults in the world’s population is projected to continue growing until the end of this century (73), meaning that filling this knowledge gap will benefit a steadily increasing number of people.

In the one study to date exploring intergenerational INS involving older adults, Dikker et al. (72) employed a neurocognitive model to simulate EEG-based INS during intergenerational social interactions between grandparents, parents, and grandchildren. In this seminal study, the authors found greater simulated INS between grandparent-grandchild dyads and grandparent-parent-child triads, relative to grandparent-parent and parent-child dyads. The authors report that the increase in simulated grandparent-child INS may be driven by age-related changes in the spectrum of brain activity. The authors also encourage the collection of real-world data to validate these simulations (72). These findings underscore how including people of all ages, particularly underrepresented older adults, in intergenerational hyperscanning will enhance our understanding of INS and a wider array of types of social connections.

#### 1.2.3 INS trajectories across days, weeks, and years remain uncharted

Most hyperscanning studies offer snapshots of INS in a single moment in time, and compare INS between contexts, tasks, or mental states. The *trajectory* in INS over days or weeks remains unknown, despite the value of granularity insights for understanding changes in behaviour related to the investigated contexts, task, and mental states. Longitudinal hyperscanning studies typically include two time points, as opposed to a denser sampling of the trajectory of INS. Numerous reviews from different domains of social neuroscience (e.g., developmental, clinical, communication, etc.) recognise the lack of longitudinal insights into INS and call for longitudinal hyperscanning studies to be conducted (e.g., 43,74–79). The discrepancy between the number of calls for studies tracking INS trajectories and the paucity of studies reporting them substantiates the critical need for additional multi-session hyperscanning studies. Concretely, multi-session longitudinal designs can fill in the blanks in our understanding of the dynamics of forming and maintaining social connections–a field recently coined ‘relational neuroscience’ (43).

To the best of our knowledge, five hyperscanning studies have recorded INS at multiple sessions and subsequently analysed the aggregated time points, as opposed to reporting changes in INS across sessions (63,80–83). These studies are reviewed in greater detail in a position piece by Moffat et al. (36). Only one fNIRS hyperscanning study offers insights into the trajectory of INS during relationship formation (84). Sened et al.’s (84) proof-of-concept study tracked the trajectory of INS over bilateral IFG in 8 clinician-patient dyads at the first, third and fifth sessions of a 6-session intervention, where dyads completed imagery and cognitive behavioural training procedures. Sened et al. report increasing INS across sessions, as well as an association between increasing INS and reduced anxiety symptoms, improved wellbeing and perceived quality of session, but not the therapeutic alliance (i.e., the patient’s rating of quality of the collaborative relationship between clinician and patient). One further study takes a unique approach to understanding INS in the context of relationship formation, though only encompassing two time points (85). INS was recorded from 11 dyads at two sessions, the first within 5 days of the dyad becoming flatmates, and the second after 7 months of co-living. Wang et al. report that social closeness can be predicted from INS with approximately 69% accuracy, using a deep neural network model and EEG hyperscanning recordings from frontotemporal brain regions. These two studies (84,85) provide preliminary evidence that INS between dyads increases as dyads grow more familiar with each other, however a larger sample size and number of time points would provide a richer picture of INS trajectories in developing relationships.

#### 1.2.4 Cross-sectional research suggests that social closeness modulates INS

Taking a cross-sectional approach to understand social closeness and relationship development, some studies compare romantic couples, friends and strangers at single moments in time. Such studies suggest that the nature of a dyad’s relationship influences INS during forms of interaction including joint motor action, conversation, and classroom learning (43,53,70,80,86–88). Djalovski et al. (86) recruited romantic couples, friends and strangers to complete a finger tapping task as dyads, and a conversation-based empathy-giving task while brain activity was measured using EEG. During the finger tapping task, INS was greatest between romantic couples, followed by friends, and lowest between strangers. The authors suggest that cumulative joint experience with a specific partner enhances prediction of the partners movements, resulting in greater INS. The empathy-giving task revealed the opposite gradient with romantic couples showing the least INS, friends showing more, and strangers showing the most INS. Djalovski et al. propose that strangers monitor each other for potential negative responses more closely than couples to avoid the consequences of a potential misunderstanding. In another study examining verbal communication, Speer et al. (88) recruited friends and strangers to engage in prompted conversations with demarcated turns. The authors report that across the span of conversation, friends showed decreasing INS and strangers showed increasing INS, with the reductions in INS corresponding to more exploratory conversations and increases in INS seeking to converge on common ground. Though phrased differently, Speer et al.’s interpretation is closely aligned to Djalovski et al.’s interpretation: The less well a dyad knows each other, the greater the INS during verbal communication. To explore this further, we highlight that a central difference between the motor and verbal communication tasks is that the motor task (i.e., drawing specific objects with an ‘Etch A Sketch’) is highly specified, setting both dyad members’ expectations clearly and symmetrically, whereas the verbal communication tasks (i.e., describing a troubling or distressing event and conversing with demarcated turns) are temporally asymmetrical and inherently unpredictable. Within the framework of mutual prediction, we propose that social closeness and the degree of predictability of behaviour induced by the task contribute to the opposite INS gradients.

Other cross-sectional hyperscanning studies provide evidence that social closeness can shape the arousal that people experience while interacting, as well as INS. In a conversation study, Long et al. (87) recorded brain activity with fNIRS while participants engaged in supportive and conflictual conversations. The authors observed that romantic couples showed greater INS than control dyads when engaging in debate. INS was associated with dyads’ arousal ratings, suggesting the dynamic range of arousal in conflict-related conversations with a romantic partner moderates INS. Further, Zhang et al. (70) compared INS during a competitive button-pressing task performed by older romantic couples and older strangers and found that romantic couples were more competitive and showed greater INS than strangers. These studies suggest that dyads who are socially closer explore more of the ‘dynamic range’ of arousal.

Considered together, these studies suggest that when dyads made up of strangers meet for the first time and engage in a creative, unconstrained activity, such as drawing, INS is likely to be low. Across repeated encounters, INS levels are likely to increase. Drawing on Dikker’s model simulating intergenerational INS, one would expect that generational constellations may impact INS levels and may interact with the trajectory of INS as dyads become socially closer.

### 1.3 Present study: Tracking intergenerational INS longitudinally, across repeated collaborative interactions

In this preregistered study (https://osf.io/hz6tm), we undertook exploratory analyses to achieve three aims:

1. To characterise INS in intergenerational dyads made up of one young and one older adult.
2. To chart the trajectory of INS within same generation and intergenerational dyads, as relationships form across repeated social interactions.
3. To characterise the relationship between INS and differing levels of loneliness and social closeness, and varying attitudes toward other generations.

To achieve these aims, we recruited 31 intergenerational and 30 same generation dyads to complete a six-session creative drawing program. At each session, we collected measures of participants’ loneliness and attitudes toward other generations, then recorded dyads’ brain activity using fNIRS while they drew independently and collaboratively with oil pastels. Participants then reported how socially close they felt to their drawing partner. We calculated INS using wavelet transform coherence (WTC) for all pairs of four regions of interest (ROIs), which were the bilateral IFG and TPJ. According to fNIRS best practices, we report both fNIRS signals (oxygenated haemoglobin [HbO] and deoxygenated haemoglobin [HbR]), with HbO results presented in the main manuscript and HbR results presented in the S1 Appendix. We employed a *Bayesian estimation* (as opposed to hypothesis testing) approach (89–92). Specifically, we built hierarchical models to estimate the magnitude and uncertainty of estimates using posterior predictive distributions. We summarise the distributions of model parameters and contrasts using point estimates and 95% credible intervals called 95% HPDs (highest posterior density regions). We use this estimation approach to assess the sign, magnitude and uncertainty of parameter estimates from our models and to assess the extent to which values of interest (e.g., 0, reflecting the absence of differences between groups or drawing conditions) are among the most credible values in a distribution (89). For example, if we compute a contrast between drawing alone and together, we obtain a distribution of the predicted difference between conditions. If the 95% HPD for this distribution excludes 0, we infer a *difference* between conditions. Aligned with the estimation approach (89–92), we do not compute Bayes factors. Further methodological details are presented in Methods, as well as Appendices S2 and S4. Detailed results tables with point estimates and 95% HPD are presented in Appendices S3, S5, and S6, separately from the results text for ease of reading.

Our findings pertaining to INS in intergenerational and same generation dyads, tracked across 6 sessions in a real-world context resembling a community art program, deliver first insights into INS involving older adults, multi-session trajectories of INS, as well as the relationship between INS and measures of social connection.

## 2 Results

### 2.1 Self-reported loneliness, social closeness, and attitudes toward generations

We first investigated how self-reported loneliness, social closeness within dyads, as well as attitudes towards generations differ between intergenerational and same generation dyads (Fig A in S3 Appendix).

#### Loneliness

At the beginning of the first session, both groups reported mid-low levels of loneliness (intergenerational mean = 1.50, SD = 1.30; same generation mean = 2.33, SD = 1.65 on a scale from 0 ‘least lonely’ to 6 ‘most lonely’). Contrasts indicate that at the first session and for all sessions combined, intergenerational dyads reported feeling less lonely than same generation dyads (session 1 = 0.68 and combined sessions = 0.61 points higher). Across sessions, feelings of loneliness showed small reductions for both groups (∼ 1 % per session), with no observed difference in rate of reduction between groups (Table 1).

**Table 1.** Unstandardised point estimates from models and group contrasts per self-reported measure with 95% HPD in square brackets. Point estimates for 1) the first session, 2) all six sessions combined, and 3) the slope of change across sessions. Self-report measures are loneliness, social closeness, and attitudes toward generations (allophilia). Visualisations of distributions for each measure and trajectory across sessions by age (i.e., older or younger adult) and group in Fig A and B in S3 Appendix.

|  | Loneliness | Social Closeness | Allophilia |
| --- | --- | --- | --- |
| **First session** |  |  |  |
| Intergen | 1.54 [1.16, 1.90] | 3.32 [2.97, 3.69] | -0.56 [-3.90, 2.74] |
| Same gen | 2.22 [1.85, 2.59] | 2.87 [2.51, 3.24] | -5.20 [-8.60, -1.90] |
| Intergen-Same gen | -0.68 [-1.19, -0.15] | 0.45 [-0.06, 0.96] | 4.65 [0.05, 8.97] |
| **Sessions combined** |  |  |  |
| Intergen | 1.28 [1.02, 1.54] | 4.08 [3.76, 4.39] | -1.19 [-3.45, 1.05] |
| Same gen | 1.89 [1.62, 2.16] | 3.52 [3.19, 3.38] | -5.66 [-7.93, 3.28] |
| Intergen-Same gen | -0.61 [-0.99, -0.25] | 0.56 [0.10, 1.00] | 4.48 [1.29, 7.56] |
| **Change across six sessions** |  |  |  |
| Intergen | -0.06 [-0.10, -0.02] | 0.24 [0.19, 0.27] | -0.21 [-0.55, 0.12] |
| Same gen | -0.07 [-.11, - 0.03] | 0.25 [0.20, 0.30] | -0.02 [-0.37, 0.30] |
| Intergen-Same gen | 0.01 [-0.05, 0.07] | -0.01 [-0.08, 0.07] | -0.18 [-0.66, 0.28] |

#### Social closeness

At the end of the first session, both groups reported medium levels of loneliness (intergenerational mean = 3.32, SD = 1.63; same generation mean = 2.87, SD = 1.17 on a scale from 1 ‘no overlap’ to 7 ‘complete overlap’, where more overlap indicates greater social closeness). At the first session, contrasts indicate a trend toward intergenerational dyads showing greater feelings of social closeness than same generation dyads (0.56 higher; 95% HPD includes 0, 6% of HPD < 0). For all sessions combined, intergenerational dyads reported greater feelings of social closeness, compared to same generation dyads (0.56 higher). Across sessions, feelings of social closeness increased for both groups (∼ 4 % per session), with no observed difference in rate between groups (Table 1).

#### Attitudes toward generations (allophilia)

At the beginning of the first session, participants in both groups reported feeling similarly close to their own and other generations (intergenerational mean = 0.01, SD = 14.48; same generation mean = -5.87, SD = 13.39 on a scale from -85 ‘feel closer to own generation’ to 85 ‘feel closer to other generation’). At the first session and for all sessions combined, same generation dyads reported feeling closer to their own generation than the intergenerational dyads (session 1 = 4.65 and combined sessions = 4.48 points higher), who reported feeling equally close to their own and the other generation. Across sessions, allophilia scores stayed stable (i.e., no observed change across sessions for either group; Table 1).

#### Relationships between measures of social connection

As presented in Table 2, greater social closeness was associated with reduced feelings of loneliness for same generation dyads. A matching trend was observed for intergenerational dyads (95% HPD include 0, 4% of HPD > 0). Greater social closeness was associated with more positive attitudes toward other generations for intergeneration dyads but not same generation dyads. Greater loneliness was associated with more positive attitudes toward other generations for same generation dyads but not intergenerational dyads. None of these associations differed between groups.

**Table 2.** Unstandardised point estimates from models of relationships between self-reported measures per group with 95% HPD in square brackets. Self-report measures are loneliness, social closeness, and attitudes toward generations (allophilia).

|  | Loneliness | Social Closeness |
| --- | --- | --- |
| **Social Closeness (Intergen)** | -0.13 [-0.27, 0.01] |  |
| **Social Closeness (Samegen)** | -0.16 [-0.28, -0.04] |  |
| **Allophilia (Intergen)** | 0.01 [-0.00, 0.02] | 0.02 [0.01, 0.04] |
| **Allophilia (Samegen)** | 0.02 [0.01, 0.03] | 0.01 [-0.01, 0.20] |

### 2.2 INS

#### 2.2.1 Real dyads vs. pseudo dyads

The first step in our analysis of INS was to perform contrasts between the posterior predictive distributions computed for real and pseudo dyads’ INS levels. Pseudo dyads consist of pairs of participants who never interacted, but match a group type (e.g., both younger adults, or one older and one younger adult). We first did this for all sessions combined and then for changes in INS levels across sessions per combination of ROI pair, task, and group. These contrasts serve to assess whether the level of observed INS (or the slope of change in INS) differs meaningfully from a null distribution of INS levels (or slopes) made up of dyads who never interacted with one another. Further statistical analyses can then be performed on the ROI pairs where the posterior predictive distributions for real and pseudo dyads differ (i.e., the 95% HPD of the distribution of the difference excludes zero). To summarise these distributions for model parameters (e.g., INS levels for real dyads) and for contrasts (e.g., the difference between dyad types), we report point estimates from models and contrasts with 95% HPD in Tables A-D in S5 Appendix.

##### INS levels for all sessions combined

All differences between real and pseudo dyads corresponded to greater INS for real than pseudo dyads (black squares in Fig 1A; solid lines in Fig 1B; Table B in S5 Appendix). The ROI pairs showing differences were RIFG∼RIFG for intergenerational dyads drawing alone, LIFG∼LIFG, LTPJ∼LTPJ, RIFG∼RIFG, LIFG∼RIFG, LIFG∼LTPJ, RIFG∼LTPJ for intergenerational dyads drawing together, no ROI pairs for same-generation dyads drawing alone, and LIFG∼RIFG and RIFG∼RTPJ for same-generation dyads drawing together. HbR analyses revealed similar results, in that LTPJ∼LTPJ, LIFG∼RIFG, and LIFG∼LTPJ showed differences between real and pseudo dyads for intergenerational dyads drawing together (reported in full in S1 Appendix).

**Fig 1.**
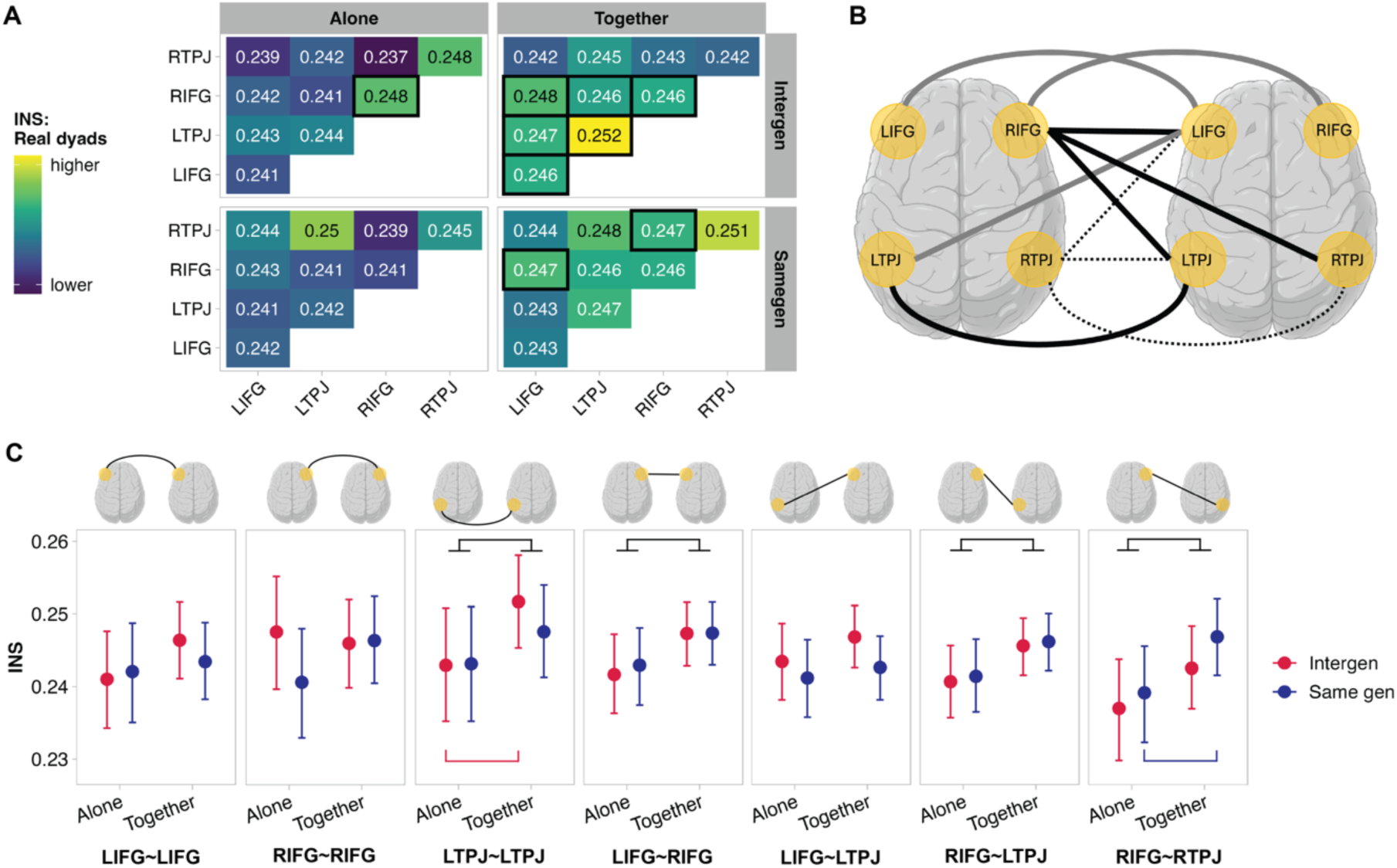
A) INS for all ROI pairs per task and group; black squares indicate that INS levels differ between real and pseudo dyads (i.e., 95% HPD of contrast distribution excludes 0). B) Visualisation of all possible ROI pairs (n=10). Dashed lines indicate ROI pairs where INS did not differ between real and pseudo dyads. Solid lines indicate ROI pairs where INS differed between real and pseudo dyads. Black solid lines indicate ROI pairs where task, group, and task*group contrasts showed differences. Grey solid lines indicate ROI pairs where contrasts showed no differences for task, group, or task*group contrasts. C) INS levels as point estimates with 95% HPD for all ROI pairs that differed between real and pseudo dyads, for which task, group, and task*group contrasts were computed. LIFG∼LIFG: No differences. RIFG∼RIFG: No differences. LTPJ∼LTPJ: Task difference, drawing together > alone. Interaction, drawing together > alone for intergenerational dyads. LIFG∼RIFG: Task difference, drawing together > alone. LIFG∼LTPJ: No differences. RIFG∼LTPJ: Task difference, drawing together > alone. RIFG∼RTPJ: Task difference, drawing together > alone. Interaction, drawing together > alone for same generation dyads. Full list of point estimates from models and contrasts with 95% HPDs available in Tables A-B in S5 Appendix for HbO. HbR analyses visualised in Fig C in S3 Appendix, showed the same pattern of results for LIFG∼RIFG and RIFG∼LTPJ. Brains created in BioRender. Moffat, R. (2026) https://BioRender.com/6p18lgg. A and C were created using code shared in the OSF repository “Longitudinal perspectives on intergenerational inter-brain synchrony”: https://osf.io/xcgp6. Code for A: https://osf.io/xcgp6/files/v5kyc. Code to generate individual values, compute summary values, and generate each panel in C: https://osf.io/xcgp6/files/8rgu5. Table of summary values shown in C: https://osf.io/xcgp6/files/ya5hn.

##### Change in INS levels across sessions

We observed a more positive slope of change for all ROI pairs that differed between real and pseudo dyads, with one exception (black squares Fig 2A; solid lines in Fig 2B; Table D in S5 Appendix). The ROI pairs showing differences were LIFG∼RTPJ for intergenerational dyads drawing alone, LTPJ∼LPTJ and RIFG∼RTPJ for intergenerational dyads drawing together, no ROI pairs for same-generation dyads drawing alone, and RTPJ∼RTPJ and LIFG∼LTPJ for same-generation dyads drawing together. HbR analyses showed little similarity (reported in full in S1 Appendix).

**Fig 2.**
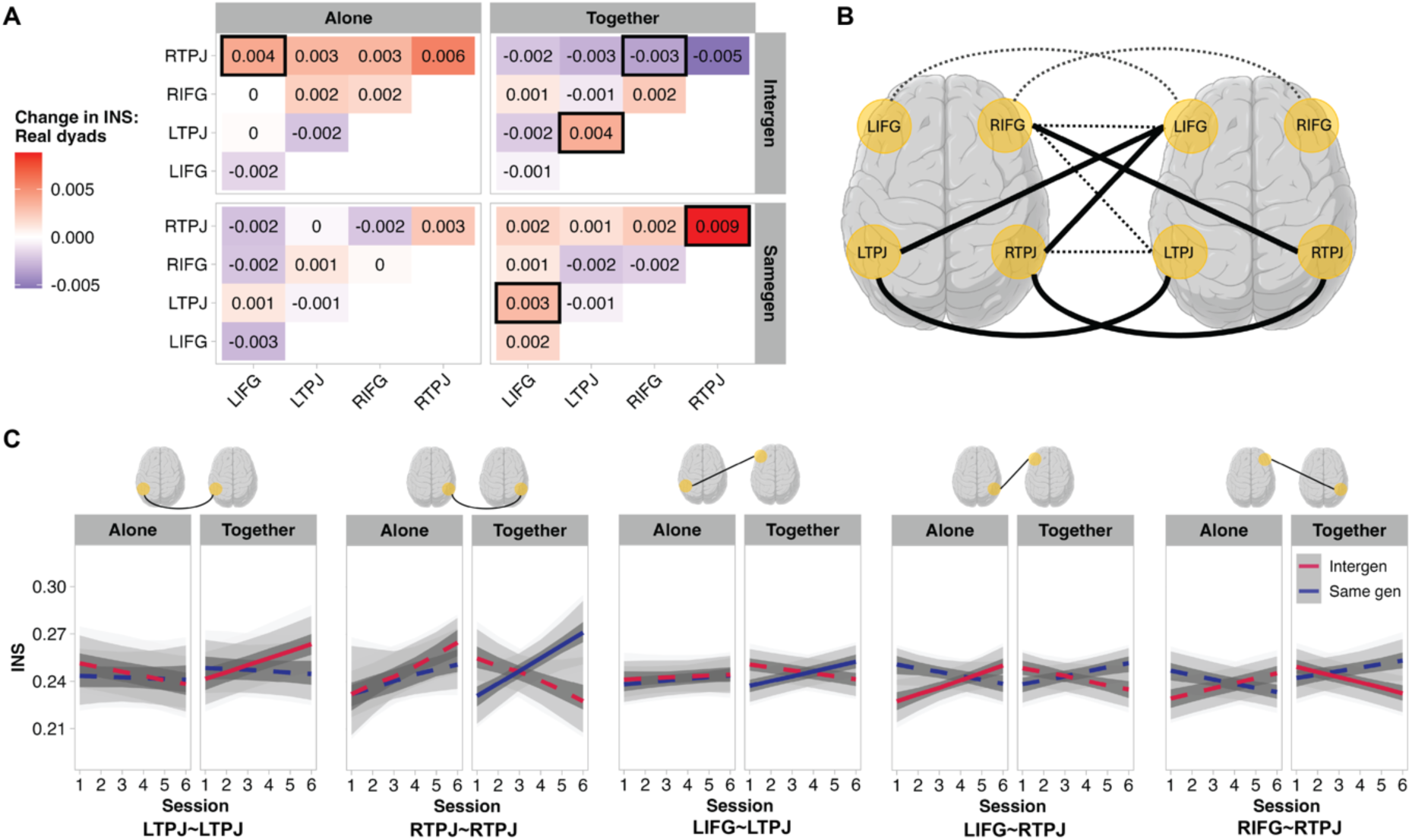
A) Slope of change in INS across six sessions; black squares indicate that INS levels differ between real and pseudo dyads (i.e., 95% HPD of contrast distribution excludes 0). B) Visualisation of all possible ROI pairs (n=10). Dashed lines indicate ROI pairs where the slope of change in INS did not differ between real and pseudo dyads. Solid lines indicate ROI pairs where the slope of change in INS differed between real and pseudo dyads. Black solid lines indicate ROI pairs where task, group, and task*group contrasts showed differences. C) Slope of change in INS across sessions for all ROI pairs that differed between real and pseudo dyads, for which task, group, and task*group contrasts were computed. Solid lines indicate that the 95% HPD of an INS∼Session slope excludes 0, indicating an association. Dashed lines indicate that the 95% HPD of an INS∼Session slope includes 0, indicating no association. Grey shading shows 95%, 89%, and 50% intervals of the posterior predictive distributions. LTPJ∼LTPJ: Interaction effect, drawing together > alone for intergenerational dyads. RTPJ∼RTPJ: Interaction effects, drawing together < alone for intergenerational dyads and intergenerational < same generation dyads for drawing together. LIFG∼LTPJ: Group effect, intergenerational < same generation dyads. LIFG∼RTPJ: Interaction effects, drawing together < alone for intergenerational dyads, intergenerational > same generation dyads for drawing alone, and intergenerational < same generation dyads for drawing together. RIFG∼RTPJ: Interaction effects, drawing together < alone for intergenerational dyads, intergenerational > same generation dyads for drawing alone, and intergenerational < same generation dyads for drawing together. Full list of point estimates from models and contrasts with 95% HPD available in Tables C-D in S5 Appendixfor HbO. HbR analyses, visualised in Fig D in S3 Appendix, showed the same pattern of results for RTPJ∼RTPJ and the same interaction effect of drawing together < alone for intergenerational dyads. Brains created in BioRender. Moffat, R. (2026) https://BioRender.com/6p18lgg. A and C were created using code shared in the OSF repository “Longitudinal perspectives on intergenerational inter-brain synchrony”: https://osf.io/xcgp6. Code for A: https://osf.io/xcgp6/files/v5kyc. Code for C: https://osf.io/xcgp6/files/8rgu5.

#### 2.2.2 Composition of non-homologous ROI pairs in intergenerational dyads

For the intergenerational dyads specifically, we examined whether INS differed as a function of the generational composition of the ROI pairs. In other words, whether an ROI belonging to an older or younger member of a dyad influenced INS levels for all sessions or slopes across sessions. Full table of estimates and contrasts with HPD available in Table E in S5 Appendix.

##### INS levels for all sessions combined

When intergenerational dyads drew together, there was a trend of greater INS when the LIFG belonged to the older adult and the LTPJ belonged to the younger adult than vice versa (95% HPD include 0, 3% of HPD < 0). We observed no differences in INS for any composition of any ROI pair for drawing alone. **Change in INS levels across sessions.** We observed no differences in the INS slopes across sessions for any composition of any ROI pair, for drawing alone or together.

#### 2.2.3 Contrasting drawing conditions and groups

##### INS levels for all sessions combined

We observed no differences between groups but did observe differences between tasks. We observed greater INS for drawing together than alone in LTPJ∼LTPJ, LIFG∼RIFG, RIFG∼LTPJ and RIFG∼RTPJ. With respect to the interaction between group and task, we found that in LTPJ∼LTPJ, INS was greater for drawing together than alone for intergenerational dyads. In RIFG∼RTPJ, INS was greater for drawing together than alone for same generation dyads. These differences correspond to ∼0.01 difference in coupling (equivalent to 39 % of the range between the minimum and maximum 95% HPD value observed in real and pseudo dyads for all sessions combined; Table A in S5 Appendix). Point estimates and 95% HPD per ROI pair, task and group from models are visualised in Fig 1C. A full list of point estimates and 95% HPDs from contrasts is presented in Table F in S5 Appendix. We also contrasted the first and second instances of drawing together. We found no differences and ruled out order-related changes in INS level while drawing together (Table G in S5 Appendix).

##### Change in INS levels across sessions

We observed no difference between drawing alone and together but did observe the following differences between groups: We observed a more positive INS slope for the same generation than the intergenerational dyads in LIFG∼LTPJ. We also found interactions between group and task. Intergenerational dyads showed a more positive INS slope for drawing together than alone in LTPJ∼LPTJ, and more positive INS slopes for drawing alone than together in RTPJ∼RTPJ, LIFG∼RTPJ, and RIFG∼RTPJ. For drawing together, the INS slope was more positive for same-generation dyads than intergenerational dyads. Same generation dyads further showed more positive INS slopes than intergenerational dyads for drawing together in RTPJ∼RTPJ, LIFG∼RTPJ and RIFG∼RTPJ. Intergenerational dyads showed more positive INS slopes than same generation dyads when drawing alone in LIFG∼RTPJ and RIFG∼RTPJ. These differences correspond to changes in the magnitude of ∼0.006 to 0.011 change in coupling from one session to the next (equivalent to 32 % of the range between the minimum and maximum 95% HPD values observed in real and pseudo dyads for all sessions combined; Table A in S5 Appendix). Point estimates and 95% HPD per ROI pair, task and group from models are visualised in Fig 2C. A full list of point estimates and 95% HPDs from contrasts is presented in Tables H-J in S5 Appendix.

#### 2.2.4 Relationship between INS and self-report measures

##### Loneliness

*Summed.* We did not observe an association between INS and summed loneliness scores. Neither group nor task influenced this relationship. We observed an interaction in RIFG∼RTPJ, for the intergenerational dyads, whereby the INS∼summed loneliness relationship was more negative for drawing alone than together (Fig 3; Tables K-L in S5 Appendix). *Difference.* We did not observe an association between INS and loneliness difference scores, and group did not influence this relationship. Task influenced the INS∼loneliness difference relationship for RIFG∼RIFG, whereby the relationship was more negative for drawing together than alone. We observed interactions for RIFG∼RIFG, RIFG∼RTPJ, and LTPJ∼LTPJ. For RIFG∼RIFG, we observed a task difference for intergenerational dyads only. For LTPJ∼LPTJ, we observed a task difference for intergenerational, but not same generation dyads and a difference between groups for drawing alone but not together. For RIFG∼RTPJ, we observed a task difference for same generation dyads only (Fig 3; Tables M-N in S5 Appendix).

**Fig 3.**
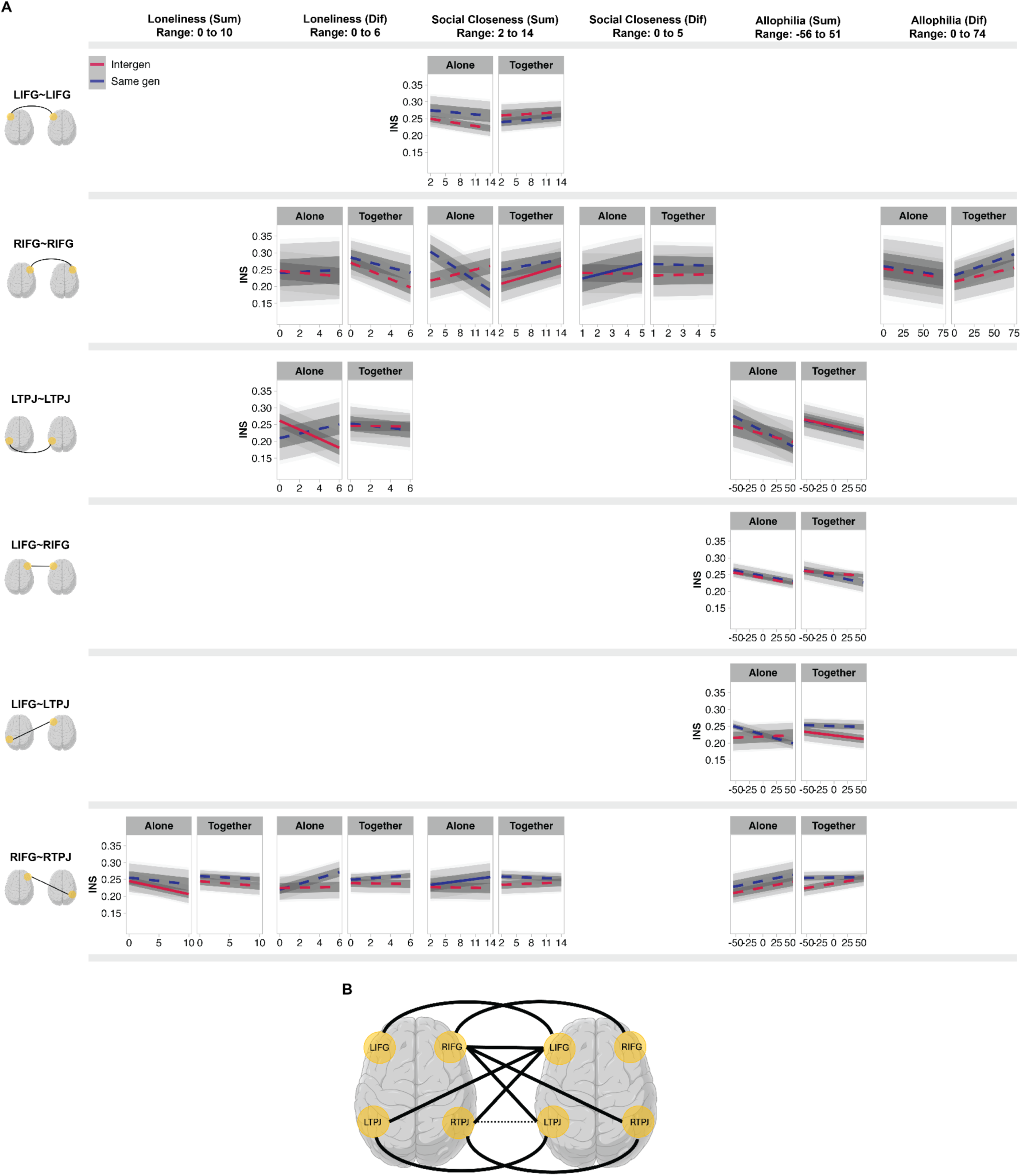
A) Slope of change in INS across levels of self-report measures for ROI pairs with at least one real vs. pseudo difference revealed by contrasts (point estimates from models and contrasts with 95 % HPD in Tables K-V in S5 Appendix). Lines indicate whether the 95% HPD for each INS∼Self-report measure slope excludes 0 (solid), indicating an association, or includes 0 (dashed), indicating no association. Shading shows 95%, 89%, and 50% intervals for the posterior predictive distributions. Results of task, group, and task*group contrasts for the panels above are as follows. *LIFG∼LIFG Social closeness (Sum):* Task difference, negative relationship for drawing alone and drawing alone < together. Interaction, together > alone for same generation. *RIFG∼RIFG Loneliness (Dif):* Task difference, drawing together < alone. *RIFG∼RIFG Social Closeness (Sum):* Group difference, intergenerational > same generation dyads. Interaction, intergenerational > same generation for drawing alone. *RIFG∼RIFG Social Closeness (Dif):* Interactions, drawing together > alone for intergenerational dyads, drawing together < alone for same generation dyads, and intergenerational > same generation for drawing alone. *RIFG∼RIFG Allophilia (Dif):* Task difference, positive relationship for drawing together and drawing together > alone. Interaction difference, drawing together > alone for same generation dyads. *LTPJ∼LTPJ Loneliness (Dif):* Interaction difference, intergenerational < same generation dyads for drawing alone. *LTPJ∼LTPJ Allophilia (Sum):* Negative relationship. Group difference, negative relationship for intergenerational dyads only. Task difference, negative relationship for drawing alone and together. *LIFG∼RIFG Allophilia (Sum):* Negative relationship. Group difference, negative relationship for same generation dyads only. *LIFG∼LTPJ Allophilia (Sum):* Interaction, drawing together < alone for intergenerational dyads. *RIFG∼RTPJ Loneliness (Sum):* No differences. *RIFG∼RTPJ Loneliness (Dif):* Interaction, drawing together < alone for same generation dyads only. *RIFG∼RTPJ Social Closeness (Sum):* Positive relationship. Group difference, positive relationship for same generation dyads and intergenerational < same generation. Task difference, positive relationship for drawing alone. Interactions, drawing together < alone for same generation dyads, and intergenerational < same generation for drawing alone. *RIFG∼RTPJ Allophilia (Sum):* Group difference, intergenerational > same generation dyads. B) Visualisation of all possible ROI pairs (n=10). Dashed lines indicate ROI pairs where i) neither INS for all sessions combined nor ii) the slope of change in INS differed between real and pseudo dyads. Solid lines indicate ROI pairs where INS (for sessions combined or slope of change across sessions) differed between real and pseudo dyads. Black solid lines indicate ROI pairs where task, group, and task*group contrasts showed differences. Analyses of HbR, visualised in Fig E in S3 Appendix, showed similar pattern of results for Social Closeness (Sum) and Allophilia (Dif) in RIFG∼RIFG, as well as Loneliness (Dif) in LTPJ∼LTPJ. Brains created in BioRender. Moffat, R. (2026) https://BioRender.com/6p18lgg. Fig 3 was created using code shared in the OSF repository “Longitudinal perspectives on intergenerational inter-brain synchrony”: https://osf.io/xcgp6. Code for individual panels in this figure: https://osf.io/xcgp6/files/6xty4.

##### Social closeness

*Summed*. We observed a positive relationship between summed social closeness scores and INS in RIFG∼RTPJ. In LIFG∼LIFG, we observed a task difference, whereby there was a more positive relationship between INS and summed social closeness for drawing together than alone. We observed a group difference for RIFG∼RIFG, whereby intergenerational dyads showed a more positive relationship than same generation dyads, and RIFG∼RTPJ where intergenerational dyads showed a more negative relationship than same generation dyads. We observed interactions for LIFG∼LIFG and RIFG∼RIFG, RIFG∼RTPJ. In LIFG∼LIFG there was a task difference for same generation dyads only. In RIFG∼RIFG there was a group difference for drawing alone only. In RIFG∼RTPJ, we observed a task difference for the same generation dyads only and a group difference only when dyads drew alone (Fig 3; Tables O-P in S5 Appendix). *Difference.* We did not observe an association between INS and social closeness difference scores. Neither group nor task influenced this relationship. We found an interaction for RIFG∼RIFG: There was a task difference for both groups and a group difference when drawing alone only (Fig 3; Tables Q-R in S5 Appendix).

##### Attitudes towards generations (allophilia)

*Summed*. We observed a negative relationship between summed allophilia scores and INS in LTPJ∼LPTJ and LIFG∼RIFG. We also found a group difference for RIFG∼RTPJ, where the relationship between INS and the summed allophilia scores was more positive for the intergenerational dyads than same generation dyads. An interaction was observed in LIFG∼LTPJ, with a task difference for intergenerational dyads only (Fig 3; Tables S-T in S5 Appendix). *Difference*. We did not observe an association between INS and allophilia difference scores, and group did not influence this relationship. We observed a task difference in RIFG∼RIFG, where the relationship between INS and the difference in allophilia scores was more positive for drawing together than alone. We found an interaction for RIFG∼RIFG, where we observed a task difference for same generation dyads only (Fig 3; Tables U-V in S5 Appendix).

## 3 Discussion

We invited intergenerational and same generation dyads to complete six session of creative drawing (resembling a community art program), and charted dyads’ levels of loneliness, social closeness, attitudes toward generations and INS as they formed social connections across 6 sessions. Our first aim was to characterise INS within intergenerational dyads comprising one young and one older adult. In a large sample (sessions combined), we show that INS among intergenerational dyads is largely comparable to same generation dyads. Specifically, both dyad types show greater INS in LTPJ∼LTPJ, LIFG∼RIFG, RIFG∼LTPJ, and RIFG∼RTPJ for drawing together than alone, with the greatest differences in LIFG∼RIFG for intergeneration dyads and RIFG∼RTPJ for same generation dyads. Our second aim was to chart the trajectory of INS in same generation and intergenerational dyads, across repeated social interactions. While drawing together, ROI pairs that include the right TPJ (RTPJ∼RTPJ, LIFG∼RTPJ, and RIFG∼RTPJ) showed decreasing INS for intergenerational dyads and increasing INS for same generation dyads. Our third aim was to characterise the relationship between INS and social connectedness, via loneliness, social closeness, and attitudes toward generations. We found that the relationship between loneliness and INS in RIFG∼RIFG differed when dyads drew alone and together: Greater similarity in dyadic levels of loneliness predicted greater INS when dyads drew together and lower INS when they drew alone. Greater social closeness predicted increasing INS in RIFG∼RTPJ for all dyads and greater INS in RIFG∼RIFG for intergenerational than same generation dyads. With respect to dyadic attitudes towards generations, we observed that INS in LIFG∼RIFG and LTPJ∼LTPJ was elevated among dyads who felt further from other generations (i.e., closer to their own). Below, we explore the implications of these findings.

Individuals’ levels of loneliness and social closeness decreased and increased, respectively, across sessions, for both generational constellations. These findings match the target outcomes of the community programs proposed by policy makers, researchers, and health practitioners (1,5,21,22) and indicate that social connections were indeed formed. Practically, our results reveal a 1% reduction in loneliness scores per session and a 4% increase in social closeness per session. These incremental changes highlight the potential of repeated encounters. Further research with more than 6 sessions would be useful for ascertaining if and at which point these changes in loneliness and social closeness may plateau.

Considering all sessions together, we observed that the generational constellation (i.e., intergenerational vs. same generation) predicted differences in individual’s levels of loneliness, social closeness and attitudes toward other generations. For example, simply being in the presence of a similar or quite differently aged person may alter one’s perceptions of one’s loneliness. If the other person is of a similar age, one might feel more lonely than if the person is older, in which case one would self-report feeling less lonely – perhaps by virtue of social comparison (93). This finding should be validated in future work by comparing intergenerational dyads with same generation dyads composed of two older adults, as well as two younger adults. Greater feelings of social closeness among intergenerational, as compared to same generation, dyads may relate to overcoming perceived generational differences or generational stereotypes (94). Notably, the distributions of difference in social closeness did not differ between groups, suggesting that the increased social closeness observed for intergenerational dyads was not driven by older or younger adults’ responses alone, but rather by both age groups. Intergenerational dyads held more charitable attitudes toward other generations than same generation dyads, aligning with findings that contact with older adults reduces younger adults’ negative stereotypes of and anxiety regarding older adults (95). Among intergenerational dyads, individuals’ attitudes toward generations were linked to their social closeness ratings, highlighting possible bidirectional interplay between attitudes and social closeness. Moreover, among same generation dyads, greater loneliness was associated with a weaker preference towards one’s ‘own’ (younger) generation, highlighting the window of opportunity for intergenerational social activities in combatting loneliness.

This study offers novel insights into INS during collaboration in dyads composed of older and younger adults, adding rich empirical data to the two existing studies that involve older adults (70,72). The fNIRS hyperscanning literature on collaboration focuses on the prefrontal and inferior frontal cortices and the TPJ (55). The bilateral IFG play a role in the initiation and reception of joint attention (56–58) and the bilateral TPJ are central to the integration of social information (61,62). Moreover, with some exceptions (e.g., 70), hyperscanning studies typically only report homologous pairs of brain areas (e.g., LIFG∼LIFG), for which the interpretation of heightened INS may be simpler. We preregistered and now report exploratory analyses of INS that include non-homologous ROI pairs (e.g., LIFG∼RIFG) and offer possible reasoning for observations of heightened INS.

In our large sample (all sessions combined), we show that INS in intergenerational dyads is largely comparable to same generation dyads. That is, we observed greater INS in LTPJ∼LTPJ, LIFG∼RIFG, RIFG∼LTPJ, and RIFG∼RTPJ for drawing together than alone. This aligns with the existing findings that collaboration co-occurs with greater INS in brain regions that process social information. Considered in the context of the mutual prediction framework (45–47), these findings suggest that collaborative drawing results in heightened integration of social information (LTPJ∼LTPJ) via joint attention (LIFG∼RIFG), and perhaps, integration of the other person’s maintenance of joint attention as social information (RIFG∼LTPJ and RIFG∼RTPJ). We also observed robust differences in INS while drawing together and alone in separate ROI pairs for each group: LIFG∼RIFG for the intergenerational dyads and RIFG∼RTPJ for the same generation dyads. We cautiously speculate that intergenerational dyads’ level of cognitive control while drawing together is particularly high and aligned within dyads, and that same generation dyads may evaluate and integrate information regarding the strength or continuity of their partner’s joint attention. This interpretation requires further investigation using paradigms that can quantify joint attention and cognitive control, both of which would deepen our understanding of mutual prediction as a mechanism of INS.

INS in IFG is proposed by some researchers to be unique to verbal communication, though a few studies show that INS in IFG may also play a role in non-verbal communication (55). With our very large sample, we can provide strong evidence that INS involving the IFG occurs during non-verbal communication, such as silent collaborative drawing.

Cross-sectional studies suggest that INS levels during collaboration decrease with higher levels of familiarity (such as friendship or stable romantic relationships) that require repeated social interactions to establish (53,70,80,86–88). We document INS trajectories during collaboration and independent drawing by two (initial) strangers with the highest temporal granularity to date – 6 consecutive time points. Our findings demonstrate that INS changes from a dyad’s first encounter to the final recorded session, entailing ∼6 hours of time spent together and ∼90 minutes of collaborative activity spread across 6 weeks.

In our preregistration, we hypothesised that INS would increase across sessions for both groups, with a steeper rate of change for intergenerational dyads (see Section 5.1 for preregistration details). We included this (weak) hypothesis to document our initial thoughts collected from the limited intergenerational and/or longitudinal hypercanning research. Specifically, the prediction of increasing INS with a steeper slope for intergenerational dyads was based on Dikker and colleagues’ theoretical and simulated predictions about age-related differences in INS during verbal communication (72,96). In contrast, our real-world data were collected while participants were specifically instructed to refrain from verbal communication during collaborative drawing, potentially leading to the difference between predicted and observed results. We observed that while dyads drew together, ROI pairs that include right TPJ (RTPJ∼RTPJ, LIFG∼RTPJ, and RIFG∼RTPJ) showed decreasing INS across sessions among intergenerational dyads but increasing INS among same generation dyads. This has potential implications for the mutual prediction framework, as well as the field of relational neuroscience, that merit further empirical investigation. For example, this may suggest that, in the context of collaborative drawing, intergenerational dyads’ monitoring and integration of their partner’s levels of joint attention may decrease with increasing familiarity, whereas same generation dyads may monitor each other’s levels of joint attention more closely as familiarity increases. Across sessions, INS increased in RTPJ∼RTPJ for same generation dyads and LTPJ∼LTPJ for intergenerational dyads. This difference in lateralisation may be related to the integration of “matched” or “mismatched” predictions. Whereas the right TPJ is associated with identifying “mismatched” predictions, the left TPJ encodes both “matched” and “mismatched” predictions (97). Accordingly, it is possible that same generation dyads become increasingly sensitive to “mismatched” predictions with increasing familiarity while intergenerational dyads may encode more “matched” predictions with increasing familiarity. Alternatively for intergenerational dyads, the frequency or absolute amount of mismatched predictions may decrease with increasing familiarity.

We found that the relationship between loneliness and INS in RIFG∼RIFG differed when dyads drew alone and together: Greater similarity in dyadic levels of loneliness predicted greater INS when dyads drew together and lower INS when they drew alone. This may point to how differences in loneliness can shape processes involved in mutual prediction, such as joint attention and perspective taking (40). Experiences of loneliness predict changes in cognition, such as perceptual speed, memory, and negatively biased perceptions of social information (10,11,13–15). It follows that these changes in cognition may co-occur with changes in joint attention or may alter the facility with which individuals engage in joint attention (98). It is possible that lonelier individuals may require more numerous neural resources to maintain joint attention or alternatively present hypervigilant tendencies, both of which could result in elevated INS in RIFG∼RIFG (19,99,100).

To understand INS beyond the changes in social connectedness incurred by repeated social interactions, we examined the extent to which dyads’ self-reported feelings of social closeness were reflected in INS. We observed that greater reported social closeness predicted greater INS in RIFG∼RTPJ for all dyads and in RIFG∼RIFG for the intergenerational dyads. We interpret the increasing INS in RIFG∼RTPJ as meaning that dyads who feel close to one another monitor and integrate each other’s joint attention more than dyads who feel less socially close. Drawing on hyperscanning studies that find links between arousal and INS (70,87,101,102), we tentatively propose that the increasing INS for intergenerational dyads in RIFG∼RIFG may reflect joint attention as well as heightened arousal or the facilitation of joint attention via heightened arousal.

With respect to dyadic attitudes towards generations, we observed that INS in LIFG∼RIFG and LTPJ∼LTPJ decreased as feelings of closeness to other generations increased. As this analysis is particularly exploratory in nature, with no empirical precedent, we offer a tentative suggestion that similarity in attitudes towards generations may give dyads a more internally consistent basis upon which they form predictions about each other’s behaviour (i.e., elsewhere termed “common ground”; ,88). These findings can serve as a useful point of departure for research examining the impact of different sources of ‘common ground’ on INS. For example, studies could explore whether cumulative life experience (e.g., number of years lived) gives older adults an advantage for finding common ground and/or whether similarities in socialisation (e.g., culture, age, socio-economic status, hobbies) play bigger role in establishing common ground that could shape the mutual prediction underpinning INS levels.

As recommended by Simons et al. (103), we make explicit the constraints on generality of this work. Our findings only generalise to the targeted brain regions (i.e., bilateral IFG and TPJ). Though our study offers typical coverage for fNIRS hyperscanning studies, we encourage the use of high-density fNIRS optode arrays in future research on this topic (104). We observed overlap between HbO and HbR results for analyses of all sessions together and INS trajectories across sessions, with some overlap in HbO and HbR results for relationships between INS and self-reported measures. Our adequate pairing of sample size and conservative Bayesian approach notwithstanding, we encourage other researchers to replicate these findings. Moreover, the INS self-report measures are not necessarily generalisable across cultures, due to differences in formation of relationships (105,106). Though our findings pertaining to changes in loneliness and social closeness across sessions are in line with findings from behavioural and qualitative studies performed as part of more elaborate and enriching community art programs, our sessions were attended by one dyad at a time, with partners assigned based on availability. Future studies could expand our paradigm to small group sessions with multiple dyads and free partner choice to reflect community programs more closely. We have no reason to believe that the results depend on other characteristics of the participants, materials, or context.

One design limitation of this study is that we did not have the resources to include a group of same generation dyads composed of two older adults, though this would have been the optimal design. Without such a group, we cannot be sure that differences between groups are purely related to the generational constellation and not age-related differences in neural processing. Future research should compare same generation dyads across different age ranges. Another design limitation pertains to gender pairings. We could not control for differences resulting from same gender and different gender pairings, because the combination of different ages and different genders resulted in eight unique gender pairings, each with a small number of dyads ( Fig A in S2 Appendix). Future research could focus on specific intergenerational gender pairings to gain a deeper understanding of the role of gender in INS trajectories.

A technical limitation of this study is that three optodes of one Photon Cap device unexpectedly became defective partway through data collection (and were replaced as rapidly as possible). This resulted in the exclusion of several participants’ right TPJ channels for the affected sessions. Thus, the number of data points for ROI pairs involving right TPJ, particularly RTPJ∼RPTJ, is lower than for other ROI pairs (Fig B in S2 Appendix). Our Bayesian approach accounted for this difference in sample size per ROI pair by partial pooling (for an excellent tutorial, see: ,107), and our selection of the ‘best channel’ per ROI helped ensure the quality of signals included in the analysis. Another approach would be to aggregate signals from all channels of adequate signal quality per ROI. In Fig C in S2 Appendix, we show that this approach yields a comparable pattern of numerically similar point estimates. Interested readers can find the strengths and weaknesses of each approach in the Methods. Finally, we used WTC (a linear measure) to quantify INS. WTC is the dominant method for computing INS in fNIRS signals (108), even though the interpersonal synchrony is widely believed to be non-linear. Approaches from information theory show promise for delivering insights into non-linear aspects of INS (109) and should be considered in future research.

## 4 Conclusion

In this study, we documented INS in intergenerational dyads including younger and older adults and charted the trajectory of INS during the formation of social connections facilitated by collaborative and creative activity. Our main findings were that, in same generation and intergenerational dyads, INS was greater while dyads drew together than alone. Across sessions, intergenerational dyads’ INS decreased and same generation dyads’ INS increased. Heightened feelings of social closeness were associated with greater INS in RIFG∼RTPJ, and greater differences in loneliness levels between dyad members predicted lower INS in RIFG∼RIFG. This longitudinal, intergenerational study reinforces that the dynamics of INS during the formation of social connections are multifaceted and merit further empirical attention.

## 5 Methods

The data presented here are part of the recently published *InterGenSynchrony Dataset* (110), which has been curated according to the Brain Imaging Data Structure (BIDS) and described in detail by our research team (111). The analysis presented here was completed before the curation of the dataset. We therefore share all of the data (non-BIDS formatted) and code for the present work on the OSF repository “Longitudinal perspectives on intergenerational inter-brain synchrony”: https://osf.io/xcgp6.

### 5.1 Ethics statement

We obtained written informed consent from all participants. Ethical approval was obtained from the Ethics Committee of the Canton Zurich (Ref: 2023-01073). The study was conducted according to the Declaration of Helsinki.

### 5.2 Preregistration

Prior to data collection, we preregistered our research plan to collect a multimodal data set (including fNIRS recordings, motion capture, self-report measures, behavioural collaboration tasks, as well as the experimenters’ subjective ratings of collaboration). We specified that we would likely disseminate our findings across multiple publications, examining specific analyses in individual papers, due to the scale and complexity of the data collected. Due to the lack of existing research on longitudinal hyperscanning and intergenerational hyperscanning, we conceptualised this research as *exploratory* (though we regrettably did not explicitly state this in our preregistration). Nonetheless, our preregistration includes broadly formulated predictions. Table A in S4 Appendix details where each component of the preregistered analyses can be found (i.e., already disseminated or in progress), and summarises the alignment of our preregistered predictions with our findings.

#### Preregistered sample size

We aimed to collect useable data from a minimum of 30 same generation and 30 intergenerational dyads between November 2023 and June 2024. This minimum reflected our best estimation of how we could maximise our financial, temporal, and human resources to address our preregistered research questions (112). Data collection began on November 15, 2023 and ended on June 18, 2024 as planned (Fig D in S2 Appendix).

#### Deviations from preregistration

We intended to include only older adults aged 70+ years. However, due to the difficulty of the recruitment process, we were concerned that we would not meet our preregistered minimum of 30 intergenerational dyads and therefore included one 69-year-old participant.

### 5.3 Participants

We recruited 122 community-dwelling participants from Zurich, Switzerland. Of these, 31 were older adults (aged 69+ years) and 91 were younger adults (aged 18–35 years). We assigned participants to intergenerational dyads (n = 31) and same generation dyads (n = 30) based on availability for sessions (e.g., matching people available at the same day/time for 6 consecutive weeks). Due to the logistical challenge posed by recruiting for and scheduling a multi-session study of this scale, specifically balanced or otherwise targeted gender pairings were not prioritised (see Fig A and D in S2 Appendix for a summary of gender pairings and session spacing). Intergenerational dyads consisted of an older adult (76 ± 4 years); 18 female, 13 male) with a younger adult (24 ± 4 years; 19 female, 11 male, 1 other). Same generation dyads consisted of two younger adults (22.5 ± 3.6 years; 36 female, 24 male). All dyads started the study as strangers. To be included, participants had no known history of neurological or psychological disorders (e.g., stroke, concussion, ADHD, autism, schizophrenia, depression) and were fluent in German. Seven additional dyads (3 same generation, 4 intergenerational) completed the initial session, then left the study due to scheduling difficulties. In accordance with our preregistration, their data are not included in the analysis.

We recruited participants via the University of Zurich’s Healthy Longevity Centre, ETH Zurich’s DESCIL student pool, clubs and organizations for senior citizens (e.g., theatre groups and choirs), and social media.

### 5.4 Measures

#### 5.4.1 Self-report measures

##### Loneliness

Participants completed the *6-Item Scale for Overall, Emotional, and Social Loneliness* (113) at the beginning of each session.

##### Social closeness

Participants indicated how socially close they felt to the other member of their dyad using the *Inclusion of Other in the Self Scale* (114) at the end of each session. This measure uses Venn diagrams with incremental increases from no overlap to complete overlap as visual analogue representation of social relationships.

##### Attitudes toward other generations

The *Allophilia Scale* (115) is typically used to characterise participants’ attitudes toward an outgroup (e.g., people of another ethnicity or political leaning). We adapted this scale in three variants: a) “younger people”, “older people”, and “people my age”. At the beginning of each session all participants completed the scale with “people my age”. Then older adults completed the scale with “younger people” and younger adults completed the scale with “older people”. To obtain baseline-corrected attitudes towards other generations, we subtracted each participant’s score for “people my age” from their score for the age group they did not belong to.

#### 5.4.2 fNIRS recordings

We recorded fNIRS signals using two Cortivision Photons Cap (Cortivision sp. z o.o., Lublin, Poland) and Cortivew software. Each device had 16 LED sources, which emitted wavelengths of 760 and 850 nm, and 10 detectors (26 optodes in total). The sample rate was 4-6 Hz. We positioned optodes according to the 10-5 EEG positions (116) over bilateral IFG and TPJ based on MNI-space coordinates derived from the Neurosynth database (https://neurosynth.org/). Specifically, we used the search terms ‘ifg’ and ‘tpj’ and recorded the coordinates of the area with the highest activation in each the left and right hemisphere, defining the ‘centre’ of our ROIs. These were the following: *left IFG:* -48, 22, 8; *right IFG:* 50, 22, 16; *left TPJ:* -56, -54, 20; and *right TPJ:* 56, -52, 18. See Fig E in S2 Appendix for visualisation of ROI centre coordinates and Fig F in S2 Appendix for optode arrangement.

##### Digitisation of optodes

After placement of the cap according to fiducial markers and before attaching optodes, we recorded a 3D image of each participant’s head using Structure Sensor Pro (model SA39) attached to an Apple iPad Pro (v. 10). The 3D head scan was processed using MATLAB (v. 2023b) and the Fieldtrip toolbox (117). Labels for all detectors and sensors and fiducial reference points were assigned to the 3D head scan and then co-registered to MNI space. Next, we obtained the interoptode distance per channel, the coordinates of the point of measurement between optodes making up each channel, as well as distance from the point of measurement to the centre of the ROI. See Fig G in S2 Appendix for visualisation of optode distances from ROI centres.

### 5.5 Procedure

Dyads attended six sessions, with approximately one week between sessions (Fig D in S2 Appendix shows schedule of sessions and number of days between sessions per dyad). Sessions were led by a female lead-researcher who provided instructions following a script (available on OSF: https://osf.io/zms2v) and answered participants’ technical and scientific queries. A second female researcher was present to aid with technical steps and did not speak unless directly addressed. Both researchers politely deflected participants’ social inquiries.

The order of events at each session is visualised in Fig 4A. At the beginning of each session, dyads completed self-report measures of loneliness and attitudes towards generations (allophilia) on an iPad, with a felt divider between participants providing privacy. Then the fNIRS caps were placed on participants’ heads and the second researcher recorded 3D models of each participant’s head with cap on (participants instructed to sit very still while researcher circled participant with 3D scanner). Next, the lead researcher reminded that participants could converse at any time except while drawing. The researchers attached optodes to the caps, ensuring that optodes came in direct contact with the scalp by clearing hair. Then fNIRS devices were turned on, paired with recording laptops (n = 2) and fNIRS signals were verified for best possible calibration. Video recording was initiated (one GoPro11 Black was positioned with a direct view of participants as shown in Fig 4B, to allow for future motion tracking analyses).

**Fig 4.**
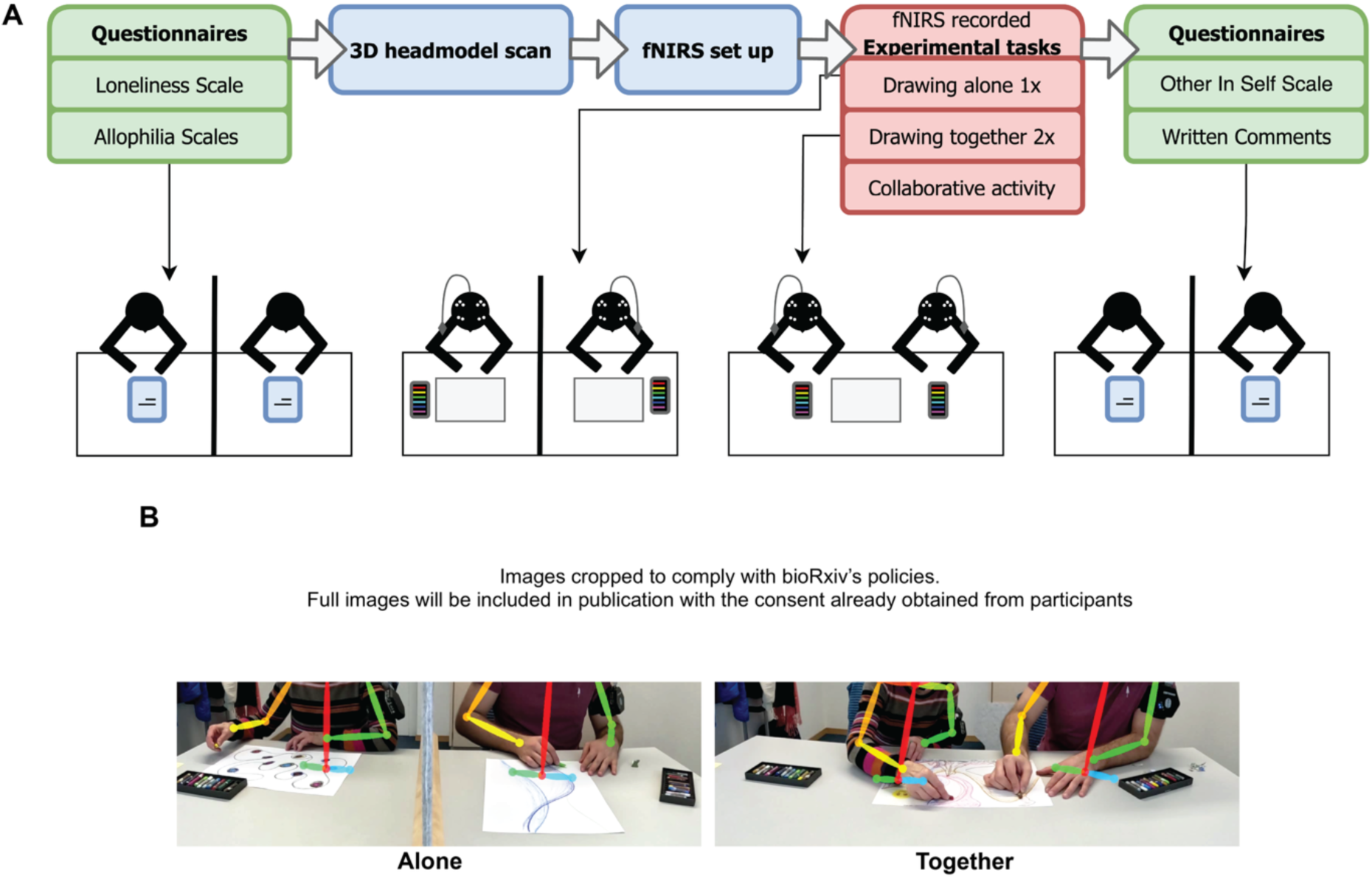
A) Flowchart visualising the protocol followed in each session. B) Left: An intergenerational dyad drawing alone on separate pages with a grey felt divider between them, blocking their view of each other. Right: The same intergenerational dyad drawing together on one page. The overlay of colourful lines shows the 2D motion tracking, which was captured, but is not reported here. Photographs taken by Ryssa Moffat. Both participants depicted in these photographs have provided written consent for their likeness to be published in open-access online research outputs.

Next, participants drew three drawings with oil pastels, without talking. Participants were allotted 5 min per drawing for each of 3 drawings, totalling 15 min of drawing per session. Participants were instructed that they would complete three drawings, the first drawing alone and the second and third together with their partner on a single piece of paper. They were instructed that they would have 5 min per drawing and could draw anything they liked. No further instructions were provided (e.g., no suggestions for motifs). While participants drew alone, they were separated by a felt divider, ensuring that they could not see each other. The divider was removed when they drew together. In cases where participants conversed to plan their drawings, the researchers waited for a natural pause to begin the 5 min timer. Each drawing was labelled and digitised after the session.

After completing the three drawings, the participants completed a short collaborative activity, which differed at each session to keep participants motivated. These include sorting pastels, a divergent thinking task, and a joint verbal fluency task, and a 54-piece jigsaw puzzle (to date only the puzzle collaboration is reported elsewhere (118)). Then, fNIRS recordings were stopped, and still wearing the fNIRS caps, participants completed a self-report measure of social closeness and provided written commentary on their experience in the session (Fig 4A). After the final session, the lead researcher held a semi-structured interview with each participant individually to gain qualitative insights into the participant’s experience.

Self-report measures were administered via RedCap. fNIRS recordings were synchronised using triggers sent from PsychoPy (v. 2022.2.5; ,119) marking the beginning of each drawing and the collaborative activity.

### 5.6 Data analysis

#### 5.6.1 Interbrain synchrony

##### 5.6.1.1 Individual level

###### Channel selection

We aimed to record from the identical position on participants’ heads at each session. However, due to the stretchy nature of neoprene caps and the inevitability of hair growth/haircuts, optode positions varied from session to session. This is also an acknowledged challenge in single-session studies, where a single optode arrangement will fall differently on individual participants’ heads (120,121). To overcome inter-session and inter-subject differences in optode positioning, we prioritise proximity to the centre of each ROI (121). That is, for each participant at each session, we selected the best channel per ROI. First, we excluded channels without visible cardiac rhythms (∼1 Hz), with a scalp-coupling index (SCI; 122) of <.7 for frequencies between 0.7-1.35 Hz, and with an interoptode distance <20 mm or >40 mm. Next, we ranked the remaining channels by distance from the centre of the ROI (closest to furthest) and selected the closest channel as the ‘best channel’ (Fig H in S2 Appendix). The ‘best channel’ approach optimises the test-retest reliability of signals across sessions and between individuals–both assumed to be stable in longitudinal group-based analyses (123,124). In other words, selecting the ‘best channel’ according to our criteria ensures that all included signals stem from comparable brain regions, both in anatomical location and in number (e.g., we avoid comparing INS values computed from a single adequate channel to INS values reflecting the average of 5 channels, most of which lie further from the centre of the ROI). Spatial/anatomical consistency aside, consistency in ROI size is particularly relevant given evidence that ROI size impacts the amplitude of measured hemodynamic responses in task-based designs (125). See Table B in S4 Appendix for a description of the number of recordings and sessions excluded, as well as reasons for exclusion.

###### Preprocessing

We preprocessed the fNIRS signals using MNE (version 1.71; ,126,127), MNE-NIRS (version 0.71; ,128). As summarised in Fig H in S2 Appendix, the signal was converted from raw intensity to optical density. Motion artefacts were corrected using the temporal derivative distribution repair algorithm (129) and short channel regression was applied using the short channel with the highest SCI (128). Recordings with no short channels with an SCI <.7 were excluded (Table B in S4 Appendix). Next, the signal was converted to concentrations of HbO and HbR using the Modified Beer–Lambert law (130,131) using an age-specific partial pathlength factor to account for age-related differences in the biological tissues probed by NIR light (132,133). This algorithm is suitable for participants aged 0-70 years. For participants aged >70 years, we specified the age parameter as 70. Last, a bandpass filter from 0.015 to 0.4 Hz was applied.

##### 5.6.1.2 Dyad level

###### Wavelet transform coherence (WTC)

We computed INS values separately for each 5-min drawing task (alone, 1^st^ drawing together, and 2^nd^ drawing together), all possible ROI pairs (N = 10; homologous ROI pairs = LIFG∼LIFG, RIFG∼RIFG, LTPJ∼LTPJ, RTPJ∼RTPJ; non homologous ROI pairs = LIFG∼RIFG, LIFG∼LTPJ, LIFG∼RTPJ, LTPJ∼RTPJ, RIFG∼LTPJ, RIFG∼RTPJ) for each real dyad and pseudo dyad (Fig H in S2 Appendix). Pseudo dyads are dyads made by pairing participants who never interacted, but align with our group types (e.g., both younger adults, or one older and one younger adult). The numbers of pseudo dyads per group, session, and ROI are listed in Table C in S4 Appendix (mean = 3467, SD = 1621). We computed INS values for pseudo dyads to obtain a null distribution against which to compare INS in real dyads. Differences in INS between real and pseudo dyads for specific ROI pairs indicate that INS levels are the result of the real interactions of real pairs and do not occur spuriously between any two people completing the same tasks in the same setting. WTC was computed using HyPyP (version 0.5.0b11; ,134) with a complex Morlet wavelet, for which the period range was defined as 5-50 seconds (0.02-0.2 Hz).

##### 5.6.1.3 Group level

For our preregistered statistical analyses, we take a *Bayesian estimation approach*, where, as we summarised in the Introduction, a central aim is to describe the sign, magnitude, and uncertainty of effect sizes in unstandardized, raw units (89–92). In contrast to (Bayesian or Frequentist) hypothesis testing approaches, the Bayesian estimation approach does not seek to test hypotheses or to quantify the strength of evidence for or against hypotheses. Kruschke and Liddell (89) provide an accessible overview of Bayesian and Frequentist estimation approaches and hypothesis testing approaches, arguing that Bayesian estimation approaches are useful for disseminating estimates with quantified uncertainty to enhance a field’s understanding of phenomena of interest. Kruschke and Liddell also describe how “we can assess the extent to which particular values of interest, such as null values, are among the most credible values” (89, p. 184) using this estimation approach. In practical terms, when we compute a contrast between the posterior predictive distributions for drawing alone and together, we obtain a distribution of the predicted difference between conditions. If the 95% HPD for this distribution excludes 0, we infer a difference between conditions. We therefore present our results as recommended by proponents of Bayesian estimation (89,90,92), we describe distributions using a point estimate and credible intervals (95% HPD) in figure and table format and provide our code (https://osf.io/xcgp6/) to generate complete distributions for closer inspection.

We fit multi-level models using the brms package (version 22.2; ,135) in the R language (version 4.5.1; ,136) within the RStudio IDE (version 2025.05.1+513; ,137). We defined weakly informed priors for our models (reported in Table D in S4 Appendix), thereby imposing a constrained distribution on our expected results (89,90).

###### Self-report measures

To assess the extent to which loneliness, social closeness, and attitudes toward generations change across the six sessions, we fit the following model for each self-report measure separately, using individual scores (i.e., not dyadic sum or difference scores): *Score ∼ 1 + Group * Session + (1|DyadID/ParticipantID)*. We subsequently computed contrasts between groups (intergenerational vs. same generation).

###### INS for sessions combined

We first sought to establish which, if any, ROI pairs showed different INS levels for real dyads relative to pseudo dyads. We fit separate models for each ROI pair (n = 10), task (n = 2; drawing alone or drawing together), and group (n = 2; intergenerational or same generation): *INS ∼ Dyad Type * Session*, where *Dyad Type* refers to real and pseudo dyads. Contrasts between real and pseudo dyads per model identified the ROI pairs for further analysis (Fig H in S2 Appendix).

###### Changes in INS across sessions

We first sought to establish which, if any, ROI pairs showed different INS trajectories across sessions for real dyads relative to pseudo dyads. We fit separate models for each ROI pair (n = 10), task (n = 2; drawing alone or drawing together), and group (n = 2; intergenerational or same generation): *INS ∼ Dyad Type * Session*, where *Dyad Type* refers to real and pseudo dyads. Contrasts between real and pseudo dyads per model identified the ROI pairs for further analysis (Fig H in S2 Appendix).

###### Further analyses with identified ROI pairs

We first fit models to assess whether intergenerational dyads show any directionality for non-homologous ROI pairs (e.g., high INS for older adult’s left IFG and younger adult’s right IFG, but not the inverse). Next, we fit one model for all sessions combined and one for changes across sessions including only real dyads and the respectively identified ROI pairs (models shown second tier of Fig H in S2 Appendix). We computed planned contrasts to examine task differences (drawing alone vs. drawing together), group (intergenerational vs. same generation), and the interaction between task and group. Finally, we fit a model (shown in second-last tier of Fig H in S2 Appendix) to all ROI pairs identified (sessions combined and across sessions) to examine the extent to which self-report measures of loneliness, social closeness and attitudes toward generations predict levels of INS. To capture dyadic aspects of each measure, we included summed and absolute difference scores per dyad (138,139). We report HbO findings in the main report and HbR findings in the S1 Appendix, with commentary on commonalities throughout the main report.

## Author contributions

RM: Conceptualization, Resources, Data curation, Formal analysis, Investigation, Visualization, Writing – original draft.

GD: Methodology, Software, Writing – review & editing.

ESC: Conceptualization, Funding acquisition, Project administration, Writing – review & editing

## Conflict of interest

The authors declare no potential conflict of interest.

## Data availability statement

These data were collected after the submission of the following preregistration: https://osf.io/hz6tm. All of the data collected and analysed for this study, as well as all of our code, can be found in our repository entitled ‘Longitudinal perspectives on intergenerational inter-brain synchrony’ on OSF: https://osf.io/xcgp6/.

## Supplementary information

Supplementary Materials, including supplementary Methods figures and tables, as well as supplementary Results figures and tables, are included in the file titled “SupplementaryMaterials.pdf”.

## Funding

Two Cortivision Photon Cap devices were provided to Ryssa Moffat through the Cortivision Pathfinder Program (grant number CPP-2023/09/01 to RM). https://www.cortivision.com

Guillaume Dumas was in part supported by the Institute for Data Valorization (Award Number: CF00137433 to GD). https://ivado.ca/en/

Guillaume Dumas was in supported by the Fonds de recherche du Québec (Award Number; 285289 to GD). https://frq.gouv.qc.ca/

Guillaume Dumas was in part supported by the Natural Sciences and Engineering Research Council of Canada (Award Number: DGECR-2023-00089 to GD). https://www.nserc-crsng.gc.ca/

Oil pastels were provided to Ryssa Moffat by Caran D’Ache (agreement number 712517 to RM). https://www.carandache.com/

Ryssa Moffat and Emily S. Cross were supported by and receive salaries from the Social Brain Sciences Lab at ETH Zurich. https://sbs.ethz.ch/

With the exception of the Social Brain Sciences Lab at ETH Zurich, the funders had no role in study design, data collection and analysis, decision to publish, or preparation of the manuscript. Prof Emily S. Cross leads the Social Brain Sciences Lab at ETH Zurich.

## Acknowledgements

From the Social Brain Science Lab at ETH Zurich, we thank Tessa Portier and Medea Häuselmann for their assistance with data collection, Linda Fanconi for scheduling participants’ sessions, Alistair Gadola for coding participant behaviour, Fanny Mougel for digitising the drawings. We thank Courtney Casale, Chantal Nagel, Tessa Portier, and Leia Jekel for proofreading the numerous iterations of this manuscript. From the CHU Sainte-Justine Azrieli Research Center of the University of Montreal, we thank Patrice Fortin for collaborating to make HyPyP suitable for fNIRS hyperscanning data and providing hands-on assistance while package functions were in development. We also thank the survey center operated by UZH Healthy Longevity Center, University of Zurich, for their assistance with recruitment of older adults from the Zurich community.

## Captions

**S1 Appendix.** Full reporting of HbR results.

**S2 Appendix.** Supplementary methods figures.

**S3 Appendix.** Supplementary results figures.

**S4 Appendix.** Supplementary methods tables.

**S5 Appendix.** Supplementary results tables for HbO.

**S5 Appendix.** Supplementary results tables for HbR.

## Notes

https://osf.io/xcgp6/overview

### Competing Interest Statement

The authors have declared no competing interest.

### Summary of Updates

Title changed, references to supplementary material changed.

## References

1. WHO. Social isolation and loneliness among older people: Advocacy brief [Internet]. World Health Organization; 2021. Available from: https://www.who.int/publications/i/item/9789240030749

2. Petersen J. A meta-analytic review of the effects of intergenerational programs for youth and older adults. Educational Gerontology. 2023 Mar 4;49(3):175–89. doi:10.1080/03601277.2022.2102340

3. Bartlett SP, Solomon P, Gellis Z. Comparative effectiveness of intergenerational service-learning programs on student outcomes of knowledge, attitude, and ageism. Educational Gerontology. 2021 Dec 2;47(12):559–73. doi:10.1080/03601277.2021.2011596

4. Gonzales E, Morrow-Howell N, Gilbert P. Changing medical students’ attitudes toward older adults. Gerontology & Geriatrics Education. 2010 Aug 26;31(3):220–34. doi:10.1080/02701960.2010.503128 PubMed PMID: 20730650.

5. Scarpetti G, Shadowen H, Williams GA, Winkelmann J, Kroneman M, Groenewegen PP, et al. A comparison of social prescribing approaches across twelve high-income countries. Health Policy. 2024 Apr;142:104992. doi:10.1016/j.healthpol.2024.104992

6. WHO. From loneliness to social connection [Internet]. World Health Organization; 2025. Available from: https://www.who.int/publications/i/item/978240112360

7. House JS, Landis KR, Umberson D. Social relationships and health. Science. 1988 Jul 29;241(4865):540–5. doi:10.1126/science.3399889

8. Wang F, Gao Y, Han Z, Yu Y, Long Z, Jiang X, et al. A systematic review and meta-analysis of 90 cohort studies of social isolation, loneliness and mortality. Nat Hum Behav. 2023 Jun 19. doi:10.1038/s41562-023-01617-6

9. The Lancet. Loneliness as a health issue. The Lancet. 2023 Jul;402(10396):79. doi:10.1016/S0140-6736(23)01411-3

10. Ayalon L, Shiovitz-Ezra S, Roziner I. A cross-lagged model of the reciprocal associations of loneliness and memory functioning. Psychology and Aging. 2016 May;31(3):255–61. doi:10.1037/pag0000075

11. Cachón-Alonso L, Hakulinen C, Jokela M, Komulainen K, Elovainio M. Loneliness and cognitive function in older adults: Longitudinal analysis in 15 countries. Psychology and Aging. 2023 Dec;38(8):778–89. doi:10.1037/pag0000777

12. Drewelies J, Windsor TD, Duezel S, Demuth I, Wagner GG, Lindenberger U, et al. Age trajectories of perceptual speed and loneliness: Separating between-person and within-person associations. Neupert S, editor. The Journals of Gerontology: Series B. 2022 Jan 11;77(1):118–29. doi:10.1093/geronb/gbab180

13. Cacioppo JT, Hawkley LC. Perceived social isolation and cognition. Trends in Cognitive Sciences. 2009 Oct;13(10):447–54. doi:10.1016/j.tics.2009.06.005

14. Okruszek Ł, Piejka A, Krawczyk M, Schudy A, Wiśniewska M, Żurek K, et al. Owner of a lonely mind? Social cognitive capacity is associated with objective, but not perceived social isolation in healthy individuals. Journal of Research in Personality. 2021 Aug;93:104103. doi:10.1016/j.jrp.2021.104103

15. Spithoven AWM, Bijttebier P, Goossens L. It is all in their mind: A review on information processing bias in lonely individuals. Clinical Psychology Review. 2017 Dec;58:97–114. doi:10.1016/j.cpr.2017.10.003

16. Cacioppo S, Capitanio JP, Cacioppo JT. Toward a neurology of loneliness. Psychological Bulletin. 2014;140(6):1464–504. doi:10.1037/a0037618

17. Düzel S, Drewelies J, Gerstorf D, Demuth I, Steinhagen-Thiessen E, Lindenberger U, et al. Structural Brain Correlates of Loneliness among Older Adults. Sci Rep. 2019 Sep 19;9(1):13569. doi:10.1038/s41598-019-49888-2

18. Lam JA, Murray ER, Yu KE, Ramsey M, Nguyen TT, Mishra J, et al. Neurobiology of loneliness: A systematic review. Neuropsychopharmacol. 2021 Oct;46(11):1873–87. doi:10.1038/s41386-021-01058-7

19. Brilliant D, Takeuchi H, Nouchi R, Yokoyama R, Kotozaki Y, Nakagawa S, et al. Loneliness inside of the brain: evidence from a large dataset of resting-state fMRI in young adult. Sci Rep. 2022 May 12;12(1):7856. doi:10.1038/s41598-022-11724-5

20. Lieberz J, Shamay-Tsoory SG, Saporta N, Esser T, Kuskova E, Stoffel-Wagner B, et al. Loneliness and the social brain: How perceived social isolation impairs human interactions. Advanced Science. 2021 Nov;8(21):2102076. doi:10.1002/advs.202102076

21. Holt-Lunstad J, Perissinotto C. Social isolation and loneliness as medical issues. N Engl J Med. 2023 Jan 19;388(3):193–5. doi:10.1056/NEJMp2208029

22. Wigfield A, Turner R, Alden S, Green M, Karania VK. Developing a new conceptual framework of meaningful interaction for understanding social isolation and loneliness. Social Policy and Society. 2022 Apr;21(2):172–93. doi:10.1017/S147474642000055X

23. Bessaha ML, Sabbath EL, Morris Z, Malik S, Scheinfeld L, Saragossi J. A systematic review of loneliness interventions among non-elderly adults. Clin Soc Work J. 2020 Mar;48(1):110–25. doi:10.1007/s10615-019-00724-0

24. Noone C, Yang K. Community-based responses to loneliness in older people: A systematic review of qualitative studies. Health & Social Care in the Community. 2022;30(4):e859–73. doi:10.1111/hsc.13682

25. O’Rourke HM, Collins L, Sidani S. Interventions to address social connectedness and loneliness for older adults: a scoping review. BMC Geriatr. 2018 Dec;18(1):214. doi:10.1186/s12877-018-0897-x

26. Cigna US. Loneliness Index [Internet]. Cigna US; 2018. Available from: https://www.multivu.com/players/English/8294451-cigna-us-loneliness-survey/#:~:text=The%20survey%20of%20more%20than,people%20who%20really%20understand%20them.

27. Hawkley LC, Buecker S, Kaiser T, Luhmann M. Loneliness from young adulthood to old age: Explaining age differences in loneliness. International Journal of Behavioral Development. 2022 Jan;46(1):39–49. doi:10.1177/0165025420971048

28. Qualter P, Vanhalst J, Harris R, Van Roekel E, Lodder G, Bangee M, et al. Loneliness across the life span. Perspect Psychol Sci. 2015 Mar;10(2):250–64. doi:10.1177/1745691615568999

29. Lee K, Jarrott SE, Juckett LA. Documented outcomes for older adults in intergenerational programming: A scoping review: research. Journal of Intergenerational Relationships. 2020 Apr 2;18(2):113–38. doi:10.1080/15350770.2019.1673276

30. Lokon E, Li Y, Kunkel S. Allophilia: Increasing college students’ “liking” of older adults with dementia through arts-based intergenerational experiences. Gerontology & Geriatrics Education. 2020 Oct 1;41(4):494–507. doi:10.1080/02701960.2018.1515740

31. Newman S, Ward C, Smith T, Wilson J, McCrea J. Intergenerational programs: Past, present and future. Rev. ed. New York: Routledge; 2014. 264 p.

32. WHO. What is the evidence on the role of the arts in improving health and well-being? A scoping review [Internet]. Copenhagen: World Health Organization; 2019. Available from: https://iris.who.int/handle/10665/329834

33. Anderson S, Fast J, Keating N, Eales J, Chivers S, Barnet D. Translating knowledge: Promoting health through intergenerational community arts programming. Health Promotion Practice. 2017 Jan;18(1):15–25. doi:10.1177/1524839915625037

34. Jenkins LK, Farrer R, Aujla IJ. Understanding the impact of an intergenerational arts and health project: A study into the psychological well-being of participants, carers and artists. Public Health. 2021 May;194:121–6. doi:10.1016/j.puhe.2021.02.029

35. Rubin SE, Gendron TL, Wren CA, Ogbonna KC, Gonzales EG, Peron EP. Challenging gerontophobia and ageism through a collaborative intergenerational art program. Journal of Intergenerational Relationships. 2015 Jul 3;13(3):241–54. doi:10.1080/15350770.2015.1058213

36. Moffat R, Casale CE, Cross ES. Mobile fNIRS for exploring inter-brain synchrony across generations and time. Frontiers in Neuroergonomics. 2024;4. doi:10.3389/fnrgo.2023.1260738

37. Liu D, Liu S, Liu X, Zhang C, Li A, Jin C, et al. Interactive Brain Activity: Review and Progress on EEG-Based Hyperscanning in Social Interactions. Frontiers in Psychology. 2018;9:1862. doi:10.3389/fpsyg.2018.01862

38. Hirata M, Ikeda T, Kikuchi M, Kimura T, Hiraishi H, Yoshimura Y, et al. Hyperscanning MEG for understanding mother-child cerebral interactions. Front Hum Neurosci. 2014 Mar 4;8. doi:10.3389/fnhum.2014.00118

39. Misaki M, Kerr KL, Ratliff EL, Cosgrove KT, Simmons WK, Morris AS, et al. Beyond synchrony: the capacity of fMRI hyperscanning for the study of human social interaction. Social Cognitive and Affective Neuroscience. 2021 Jan 18;16(1–2):84–92. doi:10.1093/scan/nsaa143

40. Hamilton AF de C. Hyperscanning: Beyond the hype. Neuron. 2021 Feb;109(3):404–7. doi:10.1016/j.neuron.2020.11.008

41. Novembre G, Iannetti GD. Hyperscanning alone cannot prove causality. Multibrain stimulation can. Trends in Cognitive Sciences. 2021 Feb;25(2):96–9. doi:10.1016/j.tics.2020.11.003

42. Moreau Q, Dumas G. Beyond correlation versus causation: Multibrain neuroscience needs explanation. Trends in Cognitive Sciences. 2021;25(7):542–3. doi:10.1016/j.tics.2021.02.011

43. De Felice S, Chand T, Croy I, Engert V, Goldstein P, Holroyd CB, et al. Relational neuroscience: Insights from hyperscanning research. Neuroscience & Biobehavioral Reviews. 2025 Feb;169:105979. doi:10.1016/j.neubiorev.2024.105979

44. Burgess AP. On the interpretation of synchronization in EEG hyperscanning studies: a cautionary note. Front Hum Neurosci. 2013;7. doi:10.3389/fnhum.2013.00881

45. Bilek E, Zeidman P, Kirsch P, Tost H, Meyer-Lindenberg A, Friston K. Directed coupling in multi-brain networks underlies generalized synchrony during social exchange. NeuroImage. 2022 May;252:119038. doi:10.1016/j.neuroimage.2022.119038

46. Wolpert DM, Doya K, Kawato M. A unifying computational framework for motor control and social interaction. Frith CD, Wolpert DM, editors. Phil Trans R Soc Lond B. 2003 Mar 29;358(1431):593–602. doi:10.1098/rstb.2002.1238

47. Kingsbury L, Huang S, Wang J, Gu K, Golshani P, Wu YE, et al. Correlated neural activity and encoding of behavior across brains of socially interacting animals. Cell. 2019 Jul;178(2):429–446.e16. doi:10.1016/j.cell.2019.05.022

48. Holper L, Scholkmann F, Wolf M. Between-brain connectivity during imitation measured by fNIRS. NeuroImage. 2012;63(1):212–22. doi:10.1016/j.neuroimage.2012.06.028

49. Cui X, Bryant DM, Reiss AL. NIRS-based hyperscanning reveals increased interpersonal coherence in superior frontal cortex during cooperation. NeuroImage. 2012;59(3):2430–7. doi:10.1016/j.neuroimage.2011.09.003

50. Dommer L, Jäger N, Scholkmann F, Wolf M, Holper L. Between-brain coherence during joint n-back task performance: A two-person functional near-infrared spectroscopy study. Behavioural Brain Research. 2012 Oct;234(2):212–22. doi:10.1016/j.bbr.2012.06.024

51. Osaka N, Minamoto T, Yaoi K, Azuma M, Shimada YM, Osaka M. How two brains make one synchronized mind in the inferior frontal cortex: fNIRS-based hyperscanning during cooperative singing. Frontiers in Psychology [Internet]. 2015 [cited 2023 Oct 25];6. Available from: https://www.frontiersin.org/articles/10.3389/fpsyg.2015.01811

52. Li L, Wang H, Luo H, Zhang X, Zhang R, Li X. Interpersonal neural synchronization during cooperative behavior of basketball players: a fNIRS-based hyperscanning study. Frontiers in Human Neuroscience [Internet]. 2020 [cited 2023 Oct 25];14. Available from: https://www.frontiersin.org/articles/10.3389/fnhum.2020.00169

53. Sun B, Xiao W, Lin S, Shao Y, Li W, Zhang W. Cooperation with partners of differing social experience: An fNIRS-based hyperscanning study. Brain and Cognition. 2021 Nov;154:105803. doi:10.1016/j.bandc.2021.105803

54. Xie H, Karipidis II, Howell A, Schreier M, Sheau KE, Manchanda MK, et al. Finding the neural correlates of collaboration using a three-person fMRI hyperscanning paradigm. Proc Natl Acad Sci USA. 2020 Sep 15;117(37):23066–72. doi:10.1073/pnas.1917407117

55. Czeszumski A, Liang SHY, Dikker S, König P, Lee CP, Koole SL, et al. Cooperative behavior evokes interbrain synchrony in the prefrontal and temporoparietal cortex: A systematic review and meta-analysis of fNIRS hyperscanning studies. eNeuro. 2022 Mar;9(2):ENEURO.0268-21.2022. doi:10.1523/ENEURO.0268-21.2022

56. Koechlin E, Ody C, Kouneiher F. The architecture of cognitive control in the human prefrontal cortex. Science. 2003 Nov 14;302(5648):1181–5. doi:10.1126/science.1088545

57. Redcay E, Kleiner M, Saxe R. Look at this: The neural correlates of initiating and responding to bids for joint attention. Front Hum Neurosci. 2012;6. doi:10.3389/fnhum.2012.00169

58. Yoshioka A, Tanabe HC, Sumiya M, Nakagawa E, Okazaki S, Koike T, et al. Neural substrates of shared visual experiences: A hyperscanning fMRI study. Social Cognitive and Affective Neuroscience. 2021 Dec 30;16(12):1264–75. doi:10.1093/scan/nsab082

59. Ding K, Wang H, Wang Q, Li H, Li C. Inhibitory control associated with the neural mechanism of joint attention in preschoolers: An fNIRS evidence. International Journal of Psychophysiology. 2023 Oct;192:53–61. doi:10.1016/j.ijpsycho.2023.08.006

60. Mundy P. A review of joint attention and social-cognitive brain systems in typical development and autism spectrum disorder. Eur J of Neuroscience. 2018 Mar;47(6):497–514. doi:10.1111/ejn.13720

61. Jacoby N, Bruneau E, Koster-Hale J, Saxe R. Localizing pain matrix and theory of mind networks with both verbal and non-verbal stimuli. NeuroImage. 2016 Feb;126:39–48. doi:10.1016/j.neuroimage.2015.11.025

62. Saxe R, Kanwisher N. People thinking about thinking people: The role of the temporo-parietal junction in “theory of mind.” NeuroImage. 2003 Aug;19(4):1835–42. doi:10.1016/S1053-8119(03)00230-1 PubMed PMID: 12948738.

63. Dikker S, Wan L, Davidesco I, Kaggen L, Oostrik M, McClintock J, et al. Brain-to-brain synchrony tracks real-world dynamic group interactions in the classroom. Current Biology. 2017 May;27(9):1375–80. doi:10.1016/j.cub.2017.04.002

64. Dumas G, Nadel J, Soussignan R, Martinerie J, Garnero. Inter-brain synchronization during social interaction. PLOS ONE. 2010;5(8):e12166.

65. Labonté-LeMoyne É, Léger PM, Resseguier B, Bastarache-Roberge MC, Fredette M, Sénécal S, et al. Are we in flow neurophysiological correlates of flow states in a collaborative game. In: Proceedings of the 2016 CHI Conference Extended Abstracts on Human Factors in Computing Systems [Internet]. San Jose California USA: ACM; 2016 [cited 2025 Aug 28]. p. 1980–8. Available from: https://dl.acm.org/doi/10.1145/2851581.2892356 doi:10.1145/2851581.2892356

66. Sinha N, Maszczyk T, Wanxuan Z, Tan J, Dauwels J. EEG hyperscanning study of inter-brain synchrony during cooperative and competitive interaction. In: 2016 IEEE International Conference on Systems, Man, and Cybernetics (SMC) [Internet]. 2016 [cited 2025 Aug 28]. p. 004813–8. Available from: https://ieeexplore.ieee.org/document/7844990/doi:10.1109/SMC.2016.7844990

67. Toppi J, Borghini G, Petti M, He EJ, De Giusti V, He B, et al. Investigating Cooperative Behavior in Ecological Settings: An EEG Hyperscanning Study. Yao D, editor. PLoS ONE. 2016 Apr 28;11(4):e0154236. doi:10.1371/journal.pone.0154236

68. De Felice S, Hamilton AFDC, Ponari M, Vigliocco G. Learning from others is good, with others is better: The role of social interaction in human acquisition of new knowledge. Phil Trans R Soc B. 2023 Feb 13;378(1870):20210357. doi:10.1098/rstb.2021.0357

69. Nguyen T, Bánki A, Markova G, Hoehl S. Studying parent-child interaction with hyperscanning. In: Progress in Brain Research [Internet]. Elsevier; 2020 [cited 2023 Jun 7]. p. 1–24. Available from: https://linkinghub.elsevier.com/retrieve/pii/S0079612320300455doi:10.1016/bs.pbr.2020.05.003

70. Zhang Q, Liu Z, Qian H, Hu Y, Gao X. Interpersonal competition in elderly couples: A functional near-infrared spectroscopy hyperscanning study. Brain Sciences. 2023 Mar 31;13(4):600. doi:10.3390/brainsci13040600

71. Liang ZW, Zhan ZJ, Liu YC, Xu HZ, Yu J. Interpersonal neural synchronization underlies interactive concept learning in older adults. npj Sci Learn. 2025 Oct 31;10(1):75. doi:10.1038/s41539-025-00368-5

72. Dikker S, Brito NH, Dumas G. It takes a village: A multi-brain approach to studying multigenerational family communication. Developmental Cognitive Neuroscience. 2024 Feb;65:101330. doi:10.1016/j.dcn.2023.101330

73. United Nations. World population prospects 2024: Summary of results. New York: United Nations; 2024.

74. Adel L, Moses L, Irvine E, Greenway KT, Dumas G, Lifshitz M. A systematic review of hyperscanning in clinical encounters. Neuroscience & Biobehavioral Reviews. 2025 Sep;176:106248. doi:10.1016/j.neubiorev.2025.106248

75. Bi X, Cui H, Ma Y. Hyperscanning Studies on Interbrain Synchrony and Child Development: A Narrative Review. Neuroscience. 2023 Oct;530:38–45. doi:10.1016/j.neuroscience.2023.08.035

76. Hoehl S, Bánki A, Brzozowska A, Carollo A, Kostorz K, Nguyen T, et al. A developmental framework of interpersonal neural synchrony. Developmental Review. 2025 Dec;78:101234. doi:10.1016/j.dr.2025.101234

77. Kelsen BA, Sumich A, Kasabov N, Liang SHY, Wang GY. What has social neuroscience learned from hyperscanning studies of spoken communication? A systematic review. Neuroscience & Biobehavioral Reviews. 2022 Jan;132:1249–62. doi:10.1016/j.neubiorev.2020.09.008

78. Noreika V, Georgieva S, Wass S, Leong V. 14 challenges and their solutions for conducting social neuroscience and longitudinal EEG research with infants. Infant Behavior and Development. 2020 Feb;58:101393. doi:10.1016/j.infbeh.2019.101393

79. Turk E, Vroomen J, Fonken Y, Levy J, Van Den Heuvel MI. In sync with your child: The potential of parent–child electroencephalography in developmental research. Developmental Psychobiology. 2022 Apr;64(3). doi:10.1002/dev.22221

80. Bevilacqua D, Davidesco I, Wan L, Chaloner K, Rowland J, Ding M, et al. Brain-to-brain synchrony and learning outcomes vary by student–teacher dynamics: Evidence from a real-world classroom electroencephalography study. Journal of Cognitive Neuroscience. 2019 Mar 1;31(3):401–11. doi:10.1162/jocn_a_01274

81. Ellingsen DM, Isenburg K, Jung C, Lee J, Gerber J, Mawla I, et al. Brain-to-brain mechanisms underlying pain empathy and social modulation of pain in the patient-clinician interaction. Proc Natl Acad Sci USA. 2023 Jun 27;120(26):e2212910120. doi:10.1073/pnas.2212910120

82. Grahl A, Anzolin A, Isenburg K, Lee J, Ellingsen DM, Jung C, et al. Brain and behavioral correlates of the patient-clinician relationship: A longitudinal fMRI hyper-scanning study of chronic pain patients. The Journal of Pain. 2021 May;22(5):602. doi:10.1016/j.jpain.2021.03.097

83. Grahl A, Anzolin A, Ellis S, Barton-Zuckerman M, Lee J, Jung C, et al. The patient-clinician relationship, expectancy and prior experience can modulate fibromyalgia treatment outcomes: a longitudinal fmrihyperscan study. The Journal of Pain. 2023 Apr;24(4):74–5. doi:10.1016/j.jpain.2023.02.217

84. Sened H, Gorst Kaduri K, Nathan Gamliel H, Rafaeli E, Zilcha-Mano S, Shamay-Tsoory S. Inter-brain plasticity as a mechanism of change in psychotherapy: A proof of concept focusing on test anxiety. Psychotherapy Research. 2025 Jan 20;1–15. doi:10.1080/10503307.2025.2451798

85. Wang D, Ren Y, Chen W. Relationship evolution shapes inter-brain synchrony in affective sharing: The role of self-expansion. Brain Struct Funct. 2024 Jul 25;229(9):2269–83. doi:10.1007/s00429-024-02841-0

86. Djalovski A, Dumas G, Kinreich S, Feldman R. Human attachments shape interbrain synchrony toward efficient performance of social goals. NeuroImage. 2021 Feb;226:117600. doi:10.1016/j.neuroimage.2020.117600

87. Long Y, Chen C, Wu K, Zhou S, Zhou F, Zheng L, et al. Interpersonal conflict increases interpersonal neural synchronization in romantic couples. Cerebral Cortex. 2022 Jul 21;32(15):3254–68. doi:10.1093/cercor/bhab413

88. Speer SPH, Mwilambwe-Tshilobo L, Tsoi L, Burns SM, Falk EB, Tamir DI. Hyperscanning shows friends explore and strangers converge in conversation. Nat Commun. 2024 Sep 5;15(1):7781. doi:10.1038/s41467-024-51990-7

89. Kruschke JK, Liddell TM. The Bayesian New Statistics: Hypothesis testing, estimation, meta-analysis, and power analysis from a Bayesian perspective. Psychon Bull Rev. 2018 Feb;25(1):178–206. doi:10.3758/s13423-016-1221-4

90. McElreath R. Statistical Rethinking; A Bayesian Course with Examples in R and Stan. 2nd Edition. New York: Chapman and Hall/CRC; 2020. 612 p.

91. Cumming G. The New Statistics: Why and How. Psychol Sci. 2014 Jan 1;25(1):7–29. doi:10.1177/0956797613504966

92. Frank MC, Braginsky M, Cachia J, Coles NA, Hardwicke TE, Hawkins RD, et al. Experimentology: an open science approach to experimental psychology methods. Cambridge, Massachusetts: The MIT Press; 2025.

93. Arnold AJ, Kappes HB, Klinenberg E, Winkielman P. The role of comparisons in judgments of loneliness. Front Psychol. 2021 Mar 24;12:498305. doi:10.3389/fpsyg.2021.498305

94. Galinsky AD, Ku G, Wang CS. Perspective-taking and self-other overlap: Fostering social bonds and facilitating social coordination. Group Processes & Intergroup Relations. 2005 Apr;8(2):109–24. doi:10.1177/1368430205051060

95. Liao T, Zhuoga C, Chen X. Contact with grandparents and young people’s explicit and implicit attitudes toward older adults. BMC Psychol. 2023 Sep 26;11(1):289. doi:10.1186/s40359-023-01344-7

96. Dikker S, Mech EN, Gwilliams L, West T, Dumas G, Federmeier KD. Exploring age-related changes in inter-brain synchrony during verbal communication. In: Psychology of Learning and Motivation [Internet]. Elsevier; 2022 [cited 2023 Jun 22]. Available from: https://linkinghub.elsevier.com/retrieve/pii/S007974212200024Xdoi:10.1016/bs.plm.2022.08.003

97. Doricchi F, Lasaponara S, Pazzaglia M, Silvetti M. Left and right temporal-parietal junctions (TPJs) as “match/mismatch” hedonic machines: A unifying account of TPJ function. Physics of Life Reviews. 2022 Sep;42:56–92. doi:10.1016/j.plrev.2022.07.001

98. Corbin IM, Dhand A. Unshared minds, decaying worlds: towards a pathology of chronic loneliness. The Journal of Medicine and Philosophy: A Forum for Bioethics and Philosophy of Medicine. 2024 Jul 11;49(4):354–66. doi:10.1093/jmp/jhae020

99. Hampshire A, Chamberlain SR, Monti MM, Duncan J, Owen AM. The role of the right inferior frontal gyrus: inhibition and attentional control. NeuroImage. 2010 Apr;50(3):1313–9. doi:10.1016/j.neuroimage.2009.12.109

100. Saporta N, Scheele D, Lieberz J, Nevat M, Kanterman A, Hurlemann R, et al. Altered activation in the action observation system during synchronization in high loneliness individuals. Cerebral Cortex. 2022 Dec 20;33(2):385–402. doi:10.1093/cercor/bhac073

101. Bast N, Polzer L, Raji N, Schnettler L, Kleber S, Lemler C, et al. Early intervention increases reactive joint attention in autistic preschoolers with arousal regulation as mediator. Eur Child Adolesc Psychiatry. 2025 May 10. doi:10.1007/s00787-025-02738-1

102. Lachat F, Hugueville L, Lemaréchal JD, Conty L, George N. Oscillatory brain correlates of live joint attention: a dual-EEG study. Front Hum Neurosci. 2012;6. doi:10.3389/fnhum.2012.00156

103. Simons DJ, Shoda Y, Lindsay DS. Constraints on Generality (COG): A Proposed Addition to All Empirical Papers. Perspect Psychol Sci. 2017 Nov;12(6):1123–8. doi:10.1177/1745691617708630

104. Guan S, Li Y, Geng Y, Li D, Xu Q, Niu P, et al. Advancing inter-brain synchrony measurement: A Comparative hyperscanning study of diffuse optical tomography and functional near-infrared spectroscopy. NeuroImage. 2026 Jan;325:121663. doi:10.1016/j.neuroimage.2025.121663

105. Goodwin R. Personal relationships across cultures [Internet]. 1st ed. Routledge; 2013 [cited 2025 Sep 12]. Available from: https://www.taylorfrancis.com/books/9780203434161 doi:10.4324/9780203434161

106. Lu P, Oh J, Leahy KE, Chopik WJ. Friendship importance around the world: Links to cultural factors, health, and well-being. Front Psychol. 2021 Jan 18;11:570839. doi:10.3389/fpsyg.2020.570839

107. Nalborczyk L, Batailler C, Vilain A, Bürkner PC. An introduction to Bayesian multilevel models using brms: A case study of gender effects on vowel variability in standard indonesian. Journal of Speech, Language, and Hearing Research. 2019;62:1225–42. doi:10.1044/2018_JSLHR-S-18-0006

108. Hakim U, De Felice S, Pinti P, Zhang X, Noah J, Ono Y, et al. Quantification of inter-brain coupling: A review of current methods used in haemodynamic and electrophysiological hyperscanning studies. NeuroImage. 2023 Oct;280:120354. doi:10.1016/j.neuroimage.2023.120354

109. Chidichimo E, Luppi AI, Mediano PAM, Leong V, Dumas G, Canales-Johnson A, et al. Towards an informational account of interpersonal coordination. Nat Rev Neurosci. 2025 Nov 17. doi:10.1038/s41583-025-00989-0

110. Moffat R, Cross ES. InterGenSynchrony Dataset [Internet]. OpenNeuro; 2026. https://openneuro.org/datasets/ds008192

111. Moffat R, Cross ES. A longitudinal fNIRS hypercanning dataset of same generation and intergenerational relationship development and collaboration. bioRxiv; 2026. doi:10.64898/2026.06.26.734715

112. Lakens D. Sample Size Justification. Collabra: Psychology. 2022 Mar 22;8(1):33267. doi:10.1525/collabra.33267

113. De Jong Gierveld J, van Tilburg T. A 6-item scale for overall, emotional, and social loneliness: Confirmatory tests on survey data. Res Aging. 2006 Sep;28(5):582–98. doi:10.1177/0164027506289723

114. Aron A, Aron EN, Smollan D. Inclusion of other in the self scale and the structure of interpersonal closeness. Journal of Personality and Social Psychology. 1992;63(4):17. doi:10.1037/0022-3514.63.4.596

115. Pittinsky TL, Rosenthal SA, Montoya RM. Measuring positive attitudes toward outgroups: Development and validation of the Allophilia Scale. In: Tropp LR, Mallett RK, editors. Moving beyond prejudice reduction: Pathways to positive intergroup relations. [Internet]. Washington: American Psychological Association; 2011 [cited 2023 Mar 23]. p. 41–60. Available from: http://content.apa.org/books/12319-002 doi:10.1037/12319-002

116. Oostenveld R, Praamstra P. The five percent electrode system for high-resolution EEG and ERP measurements. Clinical Neurophysiology. 2001;112(4):713–9. doi:10.1016/S1388-2457(00)00527-7 PubMed PMID: 11275545.

117. Oostenveld R, Fries P, Maris E, Schoffelen JM. Fieldtrip: open source software for advanced analysis of MEG, EEG, and invasive electrophysiological data. Computational Intelligence and Neuroscience. 2011;2011:1–9. doi:10.1155/2011/156869

118. Moffat R, Naudszus LA, Cross ES. Cardiac synchrony during collaborative drawing: A longitudinal comparison of same generation and intergenerational dyads. Ann NY Acad Sci. 2026 Apr;1558(1):e70272. doi:10.1111/nyas.70272

119. Peirce JW, Hirst RJ, MacAskill MR. Building Experiments in PsychoPy [Internet]. 2nd ed. London: Sage; 2022. Available from: https://uk.sagepub.com/en-gb/eur/building-experiments-in-psychopy/book273700

120. Zimeo Morais GA, Balardin JB, Sato JR. FNIRS Optodes’ Location Decider (fOLD): A toolbox for probe arrangement guided by brain regions-of-interest. Scientific Reports. 2018;8:1–11. doi:10.1038/s41598-018-21716-z

121. De Felice S, Hakim U, Gunasekara N, Pinti P, Tachtsidis I, Hamilton A. Having a chat and then watching a movie: how social interaction synchronises our brains during co-watching. Oxford Open Neuroscience. 2024 Feb 1;3:kvae006. doi:10.1093/oons/kvae006

122. Pollonini L, Olds C, Abaya H, Bortfeld H, Beauchamp MS, Oghalai JS. Auditory cortex activation to natural speech and simulated cochlear implant speech measured with functional near-infrared spectroscopy. Hearing Research. 2014;309:84–93. doi:10.1016/j.heares.2013.11.007

123. Blasi A, Lloyd-Fox S, Johnson Mark H, Elwell C. Test–retest reliability of functional near infrared spectroscopy in infants. Neurophotonics. 2014 Sep 8;1(2):025005. doi:10.1117/1.NPh.1.2.025005

124. Collins-Jones LH, Cooper RJ, Bulgarelli C, Blasi A, Katus L, McCann S, et al. Longitudinal infant fNIRS channel-space analyses are robust to variability parameters at the group-level: An image reconstruction investigation. NeuroImage. 2021 Aug;237:118068. doi:10.1016/j.neuroimage.2021.118068

125. Shader MJ, Luke R, Gouailhardou N, Mckay CM. The use of broad vs restricted regions of interest in functional near-infrared spectroscopy for measuring cortical activation to auditory-only and visual-only speech. Hearing Research. 2021;406:108256. doi:10.1016/j.heares.2021.108256 PubMed PMID: 34051607.

126. Gramfort A, Luessi M, Larson E, Engemann DA, Strohmeier D, Brodbeck C, et al. MEG and EEG data analysis with MNE-Python. Frontiers in Neuroscience. 2013;7:1–13. doi:10.3389/fnins.2013.00267

127. Gramfort A, Luessi M, Larson E, Engemann DA, Strohmeier D, Brodbeck C, et al. MNE software for processing MEG and EEG data. NeuroImage. 2014;86:446–60. doi:10.1016/j.neuroimage.2013.10.027 PubMed PMID: 24161808.

128. Luke R, Larson E, Shader MJ, Innes-Brown H, Van Yper L, Lee AKC, et al. Analysis methods for measuring passive auditory fNIRS responses generated by a block-design paradigm. Neurophotonics. 2021;8(2):025008. doi:10.1117/1.nph.8.2.025008

129. Santosa H, Huppert TJ, Fishburn F, Zhai X. Investigation of the sensitivity-specificity of canonical-and deconvolution-based linear models in evoked functional near-infrared spectroscopy. Neurophotonics. 2019;6(2):1–10. doi:10.1117/1.NPh.6.2.025009

130. Delpy DT, Cope M, Van Der Zee P, Arridge S, Wray S, Wyatt J. Estimation of optical pathlength through tissue from direct time of flight measurement. Physics in Medicine and Biology. 1988;33(12):1433–42. doi:10.1088/0031-9155/33/12/008 PubMed PMID: 3237772.

131. Kocsis L, Herman P, Eke A. The modified Beer-Lambert law revisited. Physics in Medicine and Biology. 2006;51(5):N91–8. doi:10.1088/0031-9155/51/5/N02 PubMed PMID: 16481677.

132. Garcia-Castro G. Implementing the Differential Pathlength Factor (DPF) in Python: The Scholkmann Method [Internet]. Available from: https://gongcastro.github.io/blog/dpf-scholkmann/dpf-scholkmann.html

133. Scholkmann F, Wolf M. General equation for the differential pathlength factor of the frontal human head depending on wavelength and age. J Biomed Opt. 2013 Oct 11;18(10):105004. doi:10.1117/1.JBO.18.10.105004

134. Ayrolles A, Brun F, Chen P, Djalovski A, Beauxis Y, Delorme R, et al. HyPyP: a Hyperscanning Python Pipeline for inter-brain connectivity analysis. Social Cognitive and Affective Neuroscience. 2021 Jan 18;16(1–2):72–83. doi:10.1093/scan/nsaa141

135. Bürkner PC. brms: An R Package for Bayesian Multilevel Models Using Stan. J Stat Soft. 2017;80(1):1–28. doi:10.18637/jss.v080.i01

136. R Core Team. R: A language and environment for statistical computing [Internet]. Vienna, Austria: R Foundation for Statistical Computing; 2022. Available from: http://www.r-project.org/

137. RStudio Team. RStudio: Integrated Development for R [Internet]. Boston, MA: RStudio, Inc.; 2020. Available from: http://www.rstudio.com/

138. Boukarras S, Placidi V, Rossano F, Era V, Aglioti SM, Candidi M. Interpersonal physiological synchrony during dyadic joint action is increased by task novelty and reduced by social anxiety. Psychophysiology. 2025 Mar;62(3):e70031. doi:10.1111/psyp.70031 PubMed PMID: 40097345; PubMed Central PMCID: PMC11913774.

139. Kenny DA, Kashy DA, Cook WA. Using multilevel modeling to study dyads. In: Dyadic Data Analysis, [Internet]. Guilford Publications; 2006. p. 78–99. Available from: https://ebookcentral.proquest.com/lib/ethz/detail.action?docID=362550

