## Appendices (S1-S6)/ Supplementary materials for "Social interactions between people of same and different generations shape longitudinal changes in interpersonal neural synchrony, loneliness, and social connection"


| Item | Pages |
|---|---|
| ***S1 Appendix*** (HbR Results) | 2-3 |
| ***S2 Appendix*** (Supp. methods figures) | |
| Fig A | 4 |
| Fig B | 4 |
| Fig C | 5 |
| Fig D | 5 |
| Fig E | 6 |
| Fig F | 7 |
| Fig G | 7 |
| Fig H | 8 |
| ***S3 Appendix*** (Supp. results figures) | |
| Fig A | 9 |
| Fig B | 9 |
| Fig C | 10 |
| Fig D | 11 |
| Fig E | 12-13 |
| ***S4 Appendix*** (Supp. methods tables) | |
| Table A | 14 |
| Table B | 15 |
| Table C | 16 |
| Table D | 16 |
| ***S5 Appendix*** (HbO Supp. results tables) | |
| Table A | 17-18 |
| Table B | 19 |
| Table C | 20-21 |
| Table D | 22 |
| Table E | 23 |
| Table F | 24 |
| Table G | 25 |
| Table H | 26 |
| Table I | 27-28 |
| Table J | 29-30 |
| Table K | 31-32 |
| Table L | 33-34 |
| Table M | 35-36 |
| Table N | 37-38 |
| Table O | 39-40 |
| Table P | 41-42 |
| Table Q | 43-44 |
| Table R | 45-46 |
| Table S | 47-48 |
| Table T | 49-50 |
| Table U | 51-52 |
| Table V | 53-54 |
| ***S6 Appendix*** (HbO Supp. results tables) | |
| Table A | 55-56 |
| Table B | 57 |
| Table C | 58-59 |
| Table D | 60 |
| Table E | 61 |
| Table F | 62 |
| Table G | 63 |
| Table H | 64-65 |
| Table I | 66-67 |
| Table J | 68-69 |
| Table K | 70-71 |
| Table L | 72-73 |
| Table M | 74-75 |
| Table N | 76-77 |
| Table O | 78-79 |
| Table P | 80-81 |
| Table Q | 82-83 |
| Table R | 84-85 |
| Table S | 86-87 |
| Table T | 88-89 |
| Table U | 90-91 |

**Supplementary Materials:** Moffat, Dumas and Cross (2026). *Social interactions between people of same and different generations shape longitudinal changes in interpersonal neural synchrony, loneliness, and social connection.*

### S1 Appendix – Analyses of INS in HbR

##### 1. Real dyad vs. pseudo dyads

**INS levels for all sessions combined.** We found that all differences corresponded to greater INS for real than pseudo dyads (black squares in Fig C in S3 Appendix [panel A]; solid lines in Fig C in S3 Appendix [panel B]). The ROI pairs showing differences were no ROI pairs for intergenerational dyads drawing alone, LTPJ~LTPJ, LIFG~RIFG, and LIFG~LTPJ for intergenerational dyads drawing together, no ROI pairs for same-generation dyads drawing alone, and RIFG~RIFG and RIFG~LTPJ for same-generation dyads drawing together. Estimates and contrasts with HPD in Tables A-B in S6 Appendix.

**Change in INS levels across sessions.** We observed a more positive slope of change for two ROI pairs that differed between real and pseudo pairs, and a differing negative slope of change for one ROI pair (black squares in Fig D in S3 Appendix [panel A]; solid lines in Fig D in S3 Appendix [panel B]). The ROI pairs showing differences were RTPJ~RTPJ and RIFG~RTPJ for intergenerational dyads drawing alone, LIFG~LIFG for intergenerational dyads drawing together, no ROI pairs for same-generation dyads drawing alone or together. Estimates and contrasts with HPD in Tables C-D in S6 Appendix.

##### 2. Composition of non-homologous ROI pairs in intergenerational dyads

**INS levels for all sessions combined.** For intergenerational dyads drawing together, we observed a trend of greater INS when the LIFG belonged to the older adult and the LTPJ belonged to the younger adult than vice versa (95% HPD include 0, 7% of HPD < 0). A pattern was observed where INS was greater when the RIFG belonged to the older adult and the LTPJ belonged to the younger adult than vice versa. **Change in INS levels across sessions.** No differences in the INS slopes across sessions for any composition of any ROI pair. A full list of point estimates and 95% HPDs from contrasts is presented in Table E in S6 Appendix.

##### 3. Contrasting drawing conditions and groups

**INS levels for all sessions combined.** We observed no differences between groups but did observe differences between tasks. We observed greater INS for drawing together than alone in LIFG~RIFG and RIFG~LTPJ. We also found an interaction between group and task. In LIFG~RIFG, there was a task difference for intergenerational, but not same-generation, dyads (see Supplementary Results Figure 3C). A full list of point estimates and 95% HPDs from contrasts is presented in Table F in S6 Appendix.

**Change in INS levels across sessions.** We observed no differences between groups or tasks but did observe interactions between group and task. Intergenerational dyads showed a more positive INS slope for drawing alone than together in RTPJ~RTPJ and RIFG~RTPJ while same generation dyads did not. In RTPJ~RPTJ, we also observed a more positive slope for intergenerational than same generation dyads when drawing alone (see Fig D in S3 Appendix [panel C]). A full list of point estimates and 95% HPDs from contrasts is presented in Tables G-I in S6 Appendix.

##### 4. Relationship between INS and self-report measures

**Loneliness.** *Summed.* We observed an association, whereby greater loneliness predicted lower INS in LIFG~RIFG, as well as a matching group difference for same generation dyads, and a matching task difference for drawing alone (Fig E in S3 Appendix; Tables J-K in S6 Appendix). No interactions between group and task were observed. *Difference.* We found an association, whereby greater loneliness predicted lower INS in LTPJ~LTPJ, as well as a matching group difference for intergenerational dyads, and matching task difference for drawing together. RIFG~RTPJ showed a task difference, whereby the INS~loneliness relationship was more positive for drawing alone than together (Fig E in S3 Appendix; Tables L-M in S6 Appendix). No interactions between group and task were observed.

**Social closeness (OIS).** *Summed*. We did not observe an association between INS and summed social closeness scores. Neither group nor task influenced this relationship. We observed an interaction in RIFG~RTPJ, where intergenerational dyads showed a more positive INS~social closeness relationship than same generation dyads when drawing together. In LTPJ~LPTJ, intergenerational dyads also showed a more negative INS~social closeness relationship than same generation dyads when drawing alone (Fig E in S3 Appendix; Tables N-O in S6 Appendix). *Difference.* We did not observe an association between INS and social closeness difference scores. We found a group difference, whereby intergenerational dyads showed a positive relationship in RIFG~RIFG. We also observed a task difference, whereby the relationship in LIFG~RIFG differed between drawing together and alone. We also observed interactions between group and task. In RTPJ~RTPJ, intergenerational dyads showed a more negative slope than same generation dyads while drawing together. In RIFG~LTPJ, intergenerational dyads showed a more positive slope than same generation dyads when drawing together (Fig E in S3 Appendix; Tables P-Q in S6 Appendix).

**Allophilia.** *Summed*. We did not observe an association between INS and summed allophilia scores. Group did not influence this relationship. We observed a task difference in RTPJ~RTPJ, with a more positive slope for drawing together than drawing alone. We also observed interactions between group and task. In LTPJ~LPTJ, we found a more positive relationship for drawing alone than together for intergenerational dyads. In RTPJ~RTPJ, we observed a more positive relationship for drawing together than alone for same generation dyads (Fig E in S3 Appendix; Tables R-S in S6 Appendix). *Difference*. We did not observe an association between INS and allophilia difference scores. A group difference was found in LTPJ~LTPJ, where intergenerational dyads showed a more positive relationship, that differed from same generation dyads. A task difference was observed, where the INS~allophilia (dif) relationship was more positive for drawing together than alone in LIFG~RIFG. Interactions between group and task were observed in RIFG~RIFG, LTPJ~LTPJ, and RPTJ~RTPJ. For RIFG~RIFG, same generation dyads showed a more positive relationship relative to intergenerational dyads while drawing alone. For LTPJ~LTPJ and RTPJ~RTPJ, intergenerational dyads showed a more positive relationship than same generation dyads while drawing together (Fig E in S3 Appendix; Tables T-U in S6 Appendix).

**Supplementary Materials:** Moffat, Dumas and Cross (2026). *Social interactions between people of same and different generations shape longitudinal changes in interpersonal neural synchrony, loneliness, and social connection.*

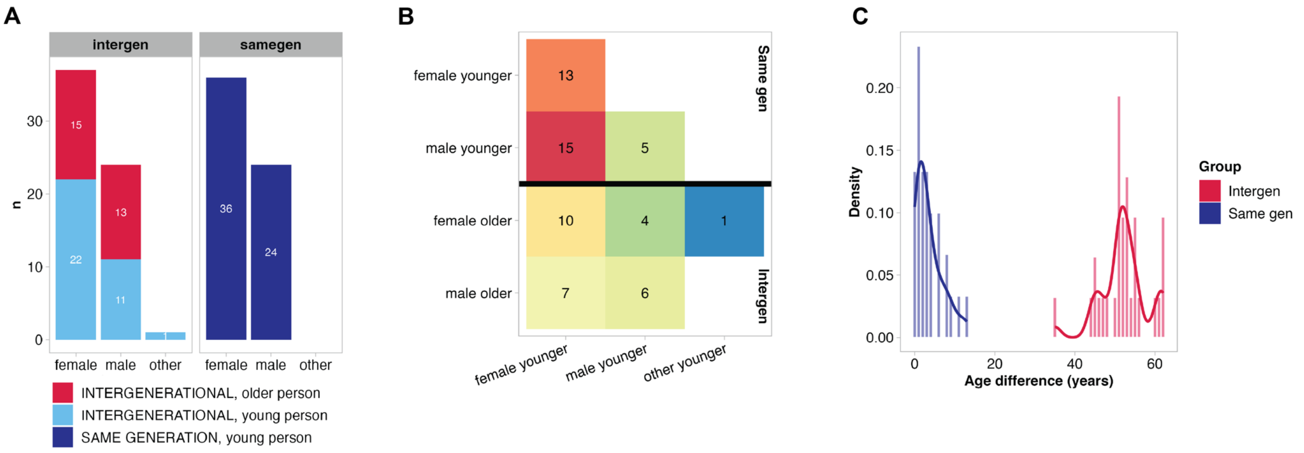


**S2 Appendix – Fig A.** A) Gender distribution per group. B) Gender/age pairings per group. C) Age difference with dyads, per group.

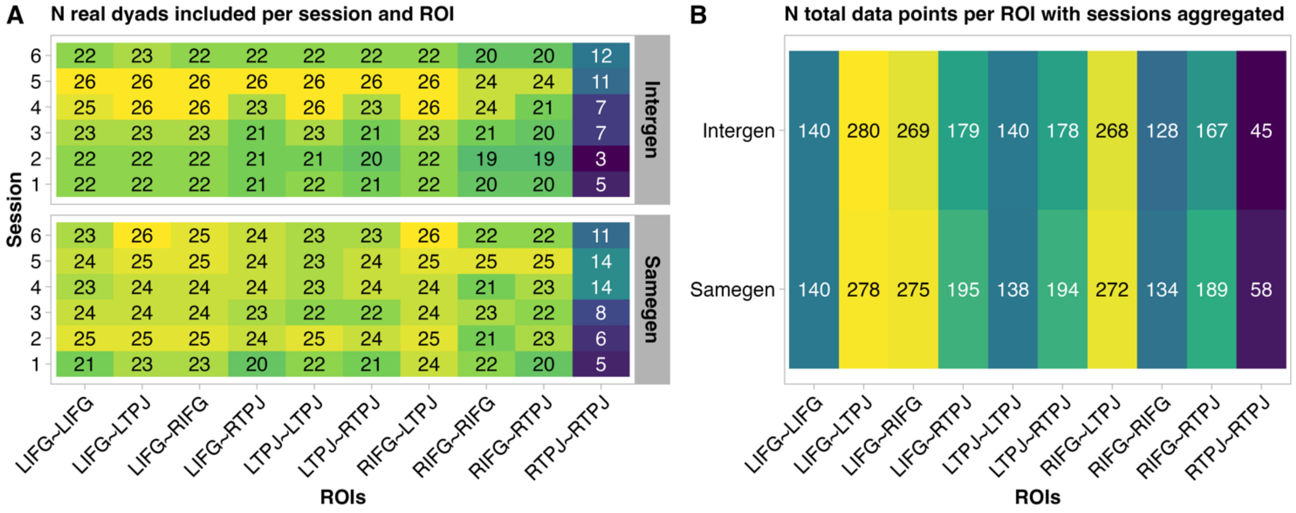


**S2 Appendix – Fig B.** A) Number of real dyads per ROI pair per session. B) Sum number of real dyads aggregated across sessions. In both A and B, the lower numbers for ROI pairs including RTPJ are the result of a period of data collection during which RTPJ optodes on one fNIRS device were broken. Data collection was not paused during this time, as not to interfere with the 6-session experimental design. New optodes were promptly ordered and installed.

**Supplementary Materials:** Moffat, Dumas and Cross (2026). *Social interactions between people of same and different generations shape longitudinal changes in interpersonal neural synchrony, loneliness, and social connection.*

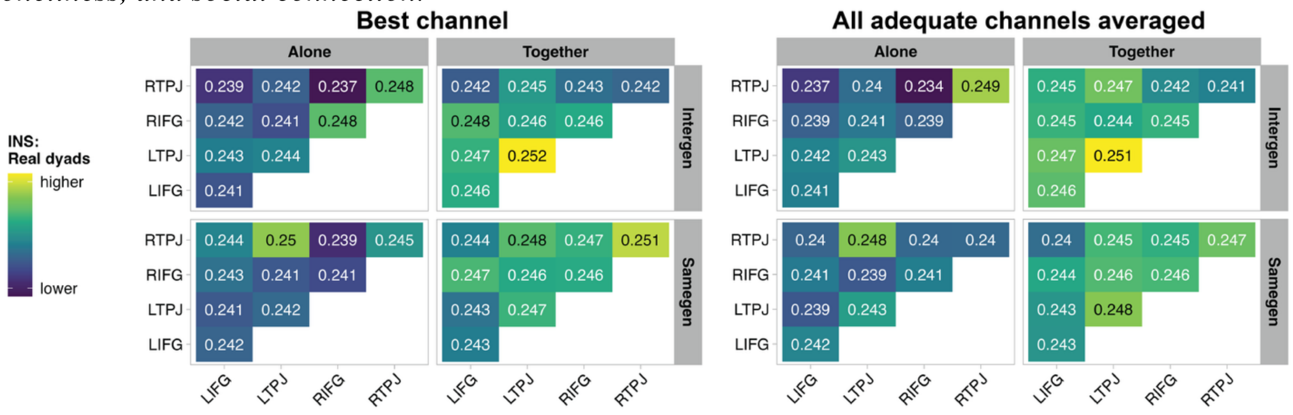


**S2 Appendix – Fig C.** Comparison of INS point estimates using the best channel (left) and the averaged INS values for all channels in each ROI (right). The best channels were selected as described in the Methods. Adequate channels had visual evidence of heart beats, channel lengths between 20 and 40 mm, and scalp-coupling indices < .7. Visual inspection reveals INS levels to be numerically similar, and the overall pattern of levels to be similar. For example, in both RIFG~RTPJ Intergen Alone is lowest value and LTPJ~LTPJ Intergen Together is highest value.

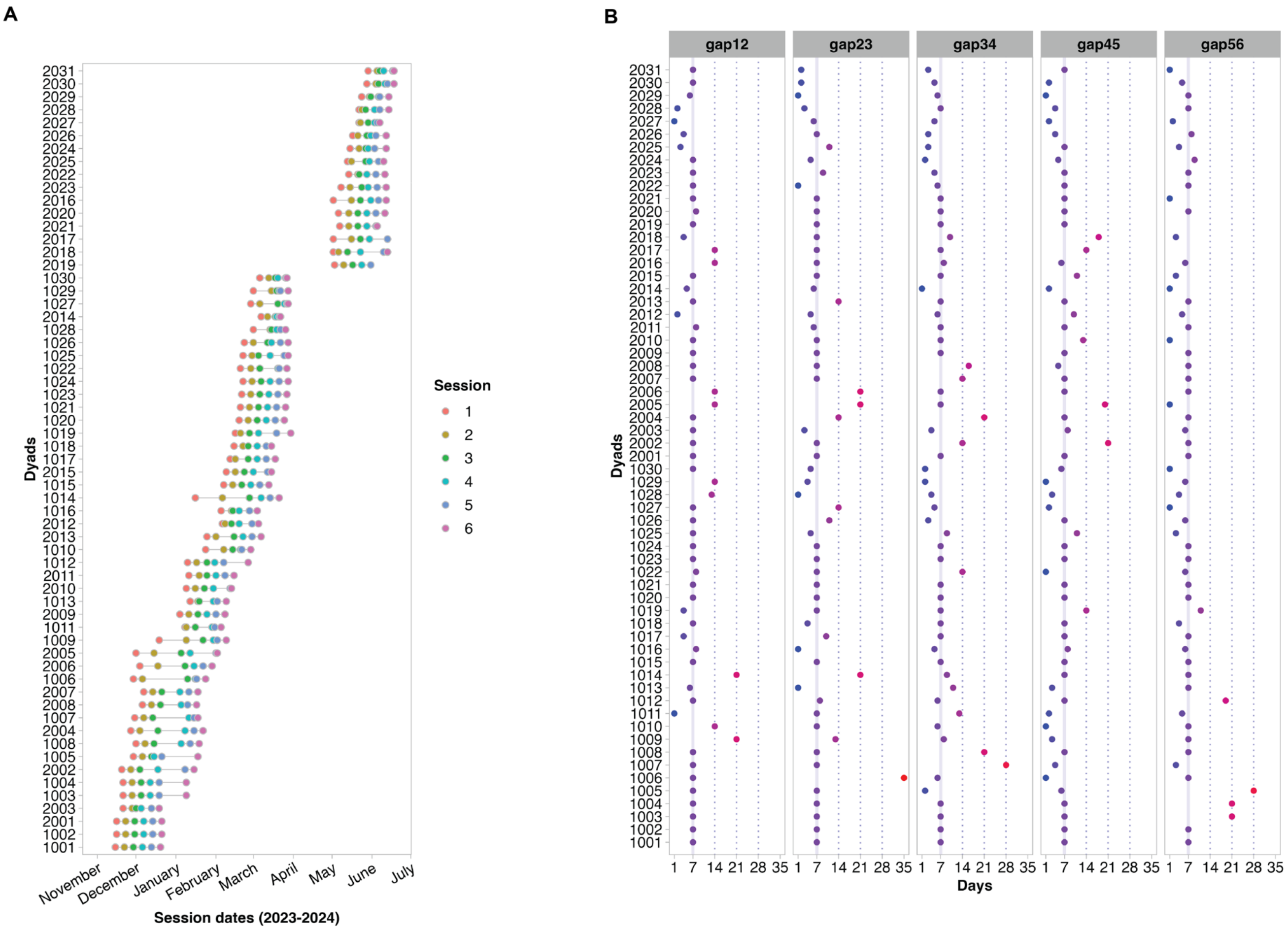


**S2 Appendix – Fig D.** A) Schedule of sessions per dyad. B) Number of days between sessions for each dyad and gap between sessions. The label *gap12* refers to the number of days between session 1 and session 2, *gap23* to the number of days between session 2 and session 3, and so on.

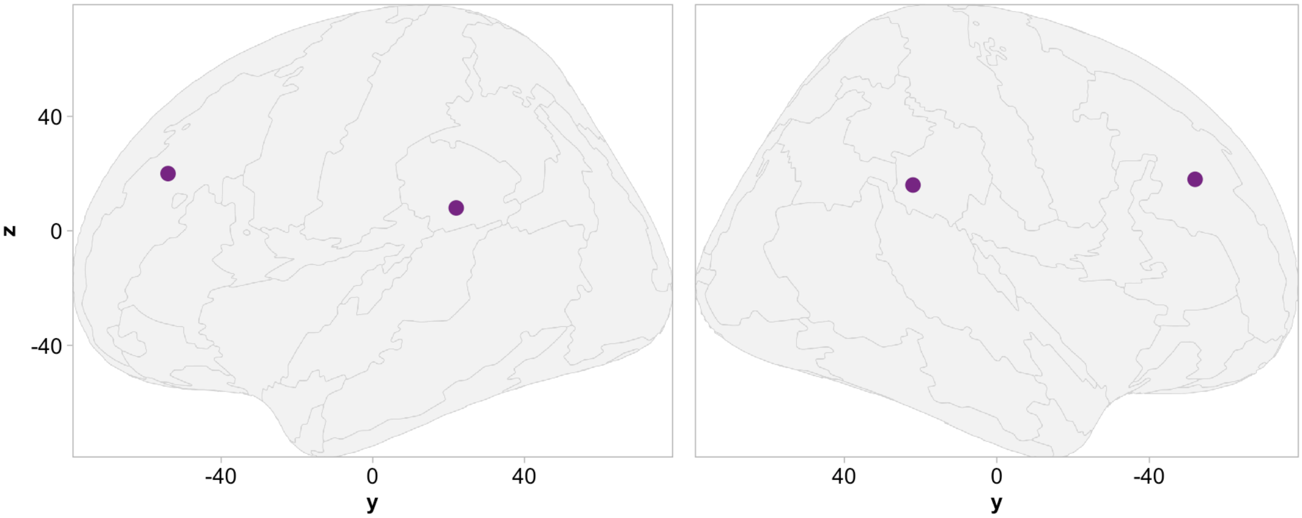


**S2 Appendix – Fig E.** Purple circles show centre of each ROI in MNI coordinates.

**Supplementary Materials:** Moffat, Dumas and Cross (2026). *Social interactions between people of same and different generations shape longitudinal changes in interpersonal neural synchrony, loneliness, and social connection.*

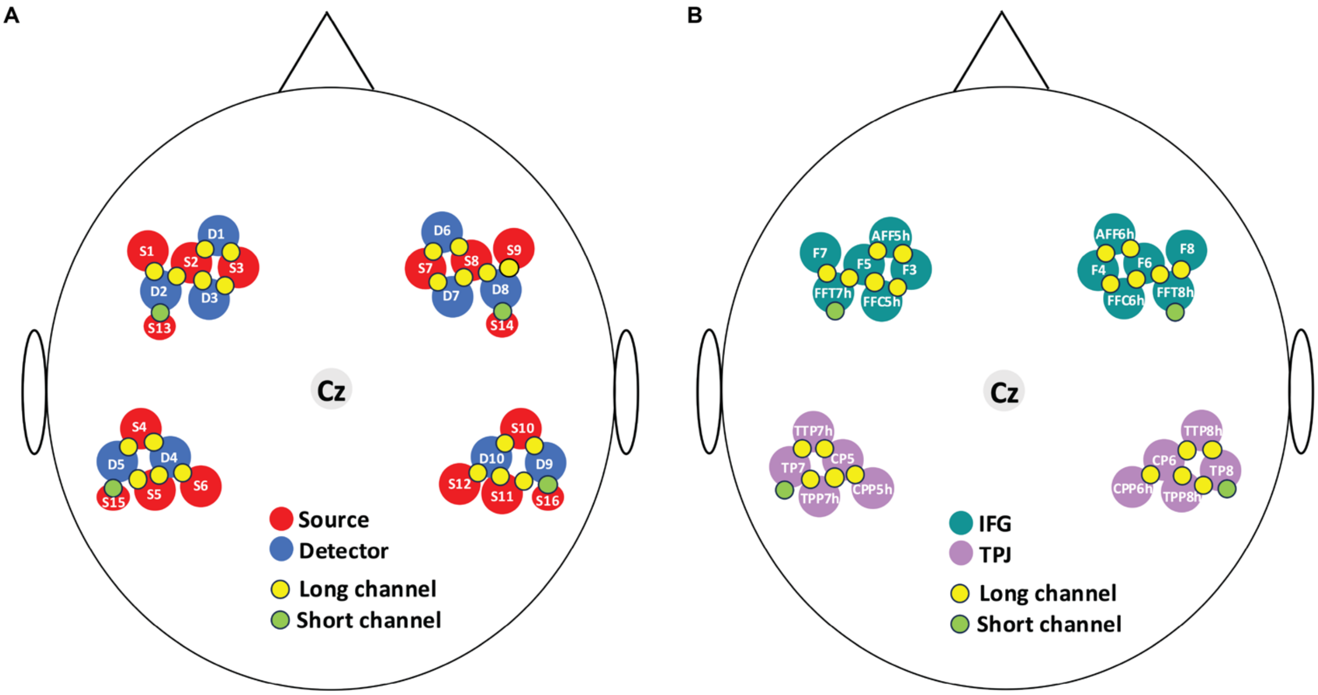


**S2 Appendix – Fig F.** A) Arrangement optodes, where numbers correspond to sensor numbers on Cortivision Photon Cap Device (S13-S16 are sources specifically designed to measure short channel signals). B) EEG 5-10 positions of each optode within the four ROIs (bilateral IFG and TPJ).

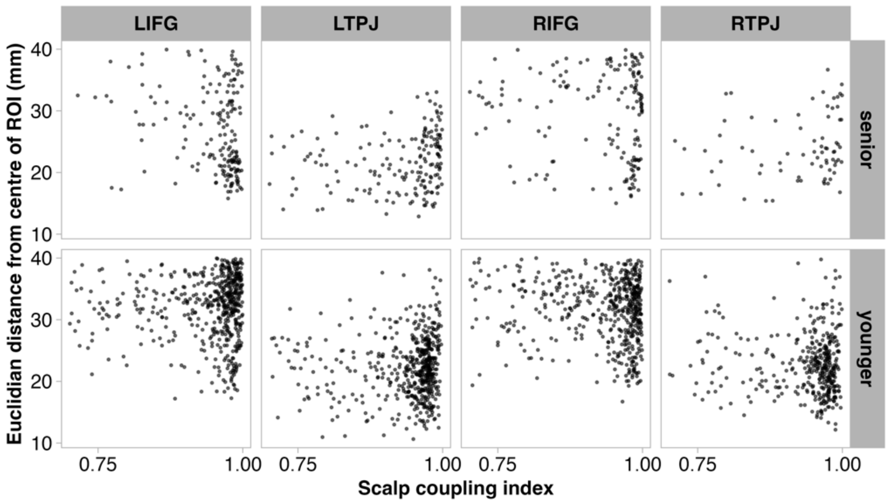


**S2 Appendix – Fig G.** Distances of selected channels from the coordinates in the centre of each ROI. One component of the distance (i.e., ~15mm) is the distance from the centre of the ROI being inside the cortex and the channel positions being on the scalp. Seniors = participants aged 69+ years. Younger = participants aged 18-35 years.

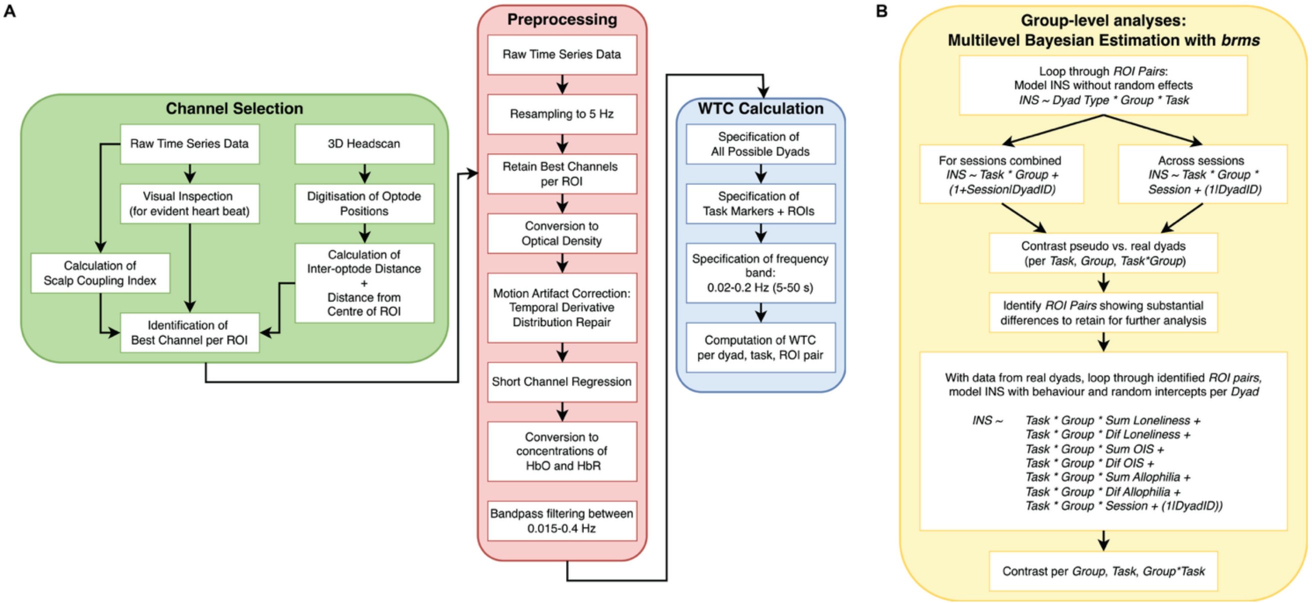


**S2 Appendix – Fig H.** A) Flowchart visualising the procedure for selecting which channels to include in analysis for individual participants, preprocessing steps for individual participants and calculation of WTC for dyads. B) Flowchart visualising the order in which group-level analyses were performed in the R package brms, and showing the models fit.

**Supplementary Materials:** Moffat, Dumas and Cross (2026). *Social interactions between people of same and different generations shape longitudinal changes in interpersonal neural synchrony, loneliness, and social connection.*

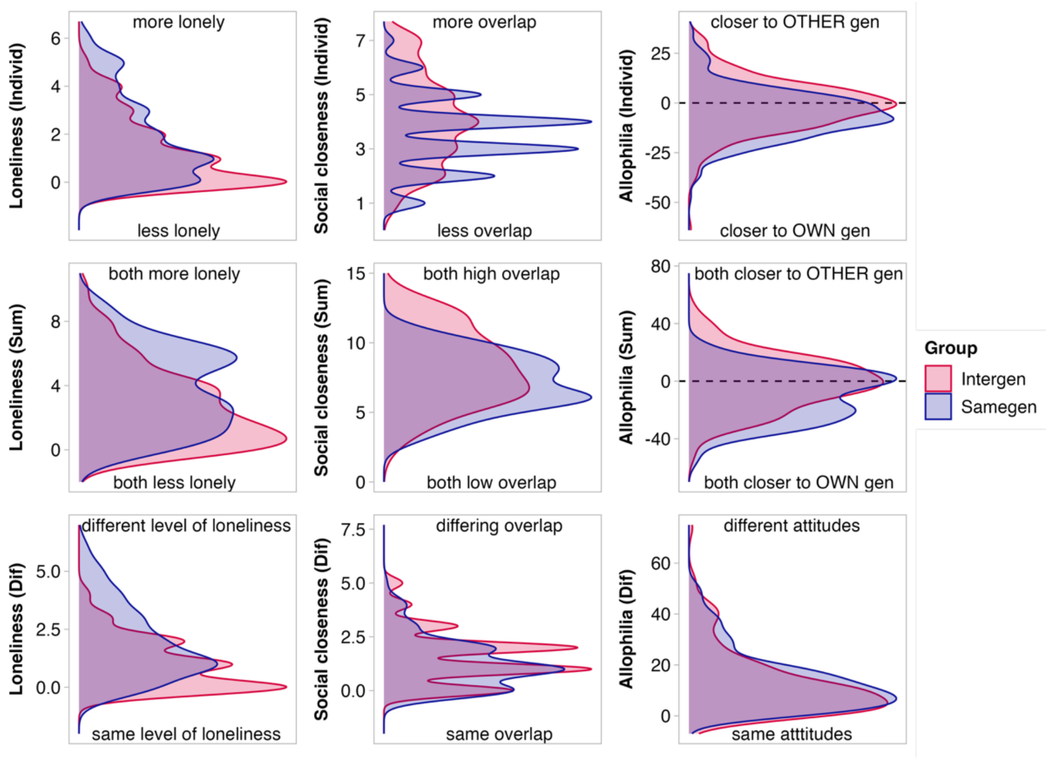


**S3 Appendix – Fig A.** Top: Distributions of self-reported loneliness, social closeness (OIS), and attitudes toward generations (allophilia) of individual participants. Middle: Distribution of summed scores per dyad. Bottom: Distribution of difference scores per dyads.

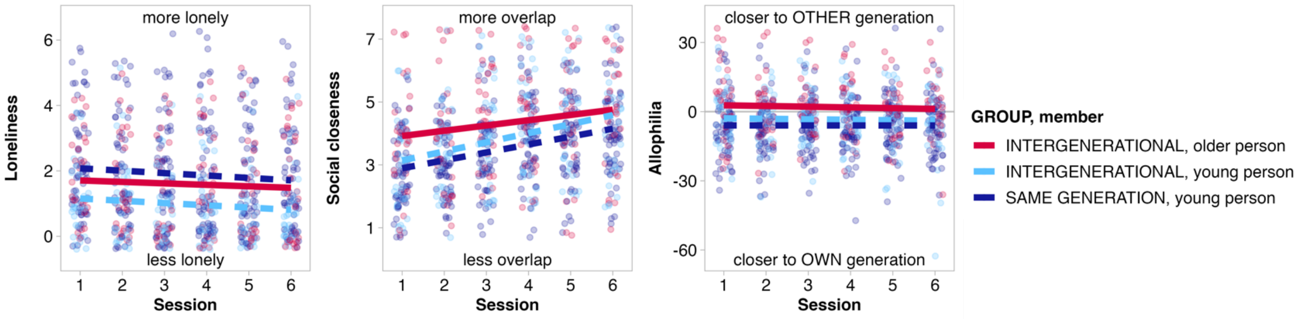


**S3 Appendix – Fig B.** Self-report scores for loneliness, social closeness, and attitudes towards generations (i.e., allophilia) plotted with younger and older members of intergenerational dyads separated.

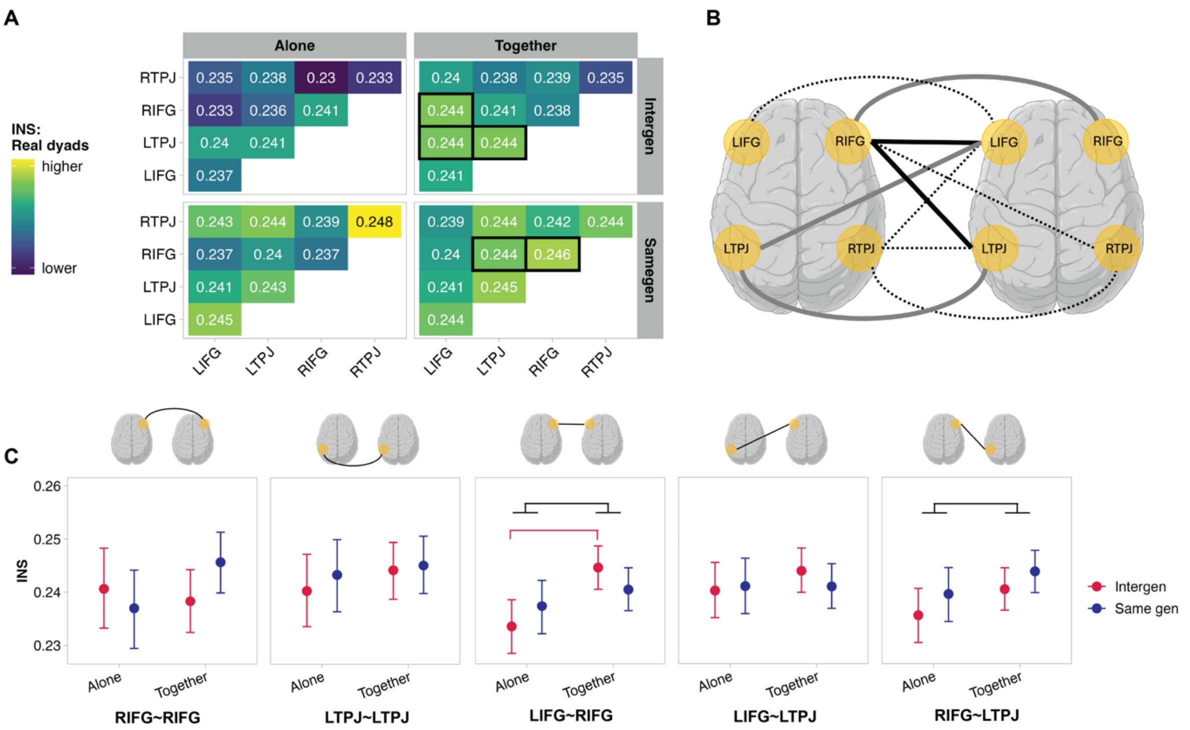


**S3 Appendix – Fig C.** HbR. INS for all ROI pairs per task and group; black squares indicate that INS levels differ between real and pseudo dyads (i.e., 95% HPD of contrast distribution excludes 0). B) Visualisation of all possible ROI pairs (n=10). Dashed lines indicate ROI pairs where INS did not differ between real and pseudo dyads. Solid lines indicate ROI pairs where INS differed between real and pseudo dyads. Black solid lines indicate ROI pairs where task, group, and task*group contrasts showed differences. Grey solid lines indicate ROI pairs where contrasts showed no differences for task, group, or task*group contrasts. C) INS levels as point estimates with 95% HPD for all ROI pairs that differed between real and pseudo dyads, for which task, group, and task*group contrasts were computed. RIFG~RIFG: No differences. LTPJ~LTPJ: No differences. LIFG~RIFG: Task difference, together > alone. Interaction, together > alone for intergenerational group. LIFG~LTPJ: No differences. RIFG~LTPJ: Task difference, together > alone. Full list of point estimates from models and contrasts with 95% HPDs available in Tables A-B in S6 Appendix. Brains created in BioRender. Moffat, R. (2026) https://BioRender.com/6p18lgg. A and C were created using code shared in the OSF repository "Longitudinal perspectives on intergenerational inter-brain synchrony": https://osf.io/xcgp6. Code for A: https://osf.io/xcgp6/files/pu7h2. Code to generate individual values, compute summary values, and generate each panel in: https://osf.io/xcgp6/files/4f8qw. Table of summary values shown in C: https://osf.io/xcgp6/files/4vynd.

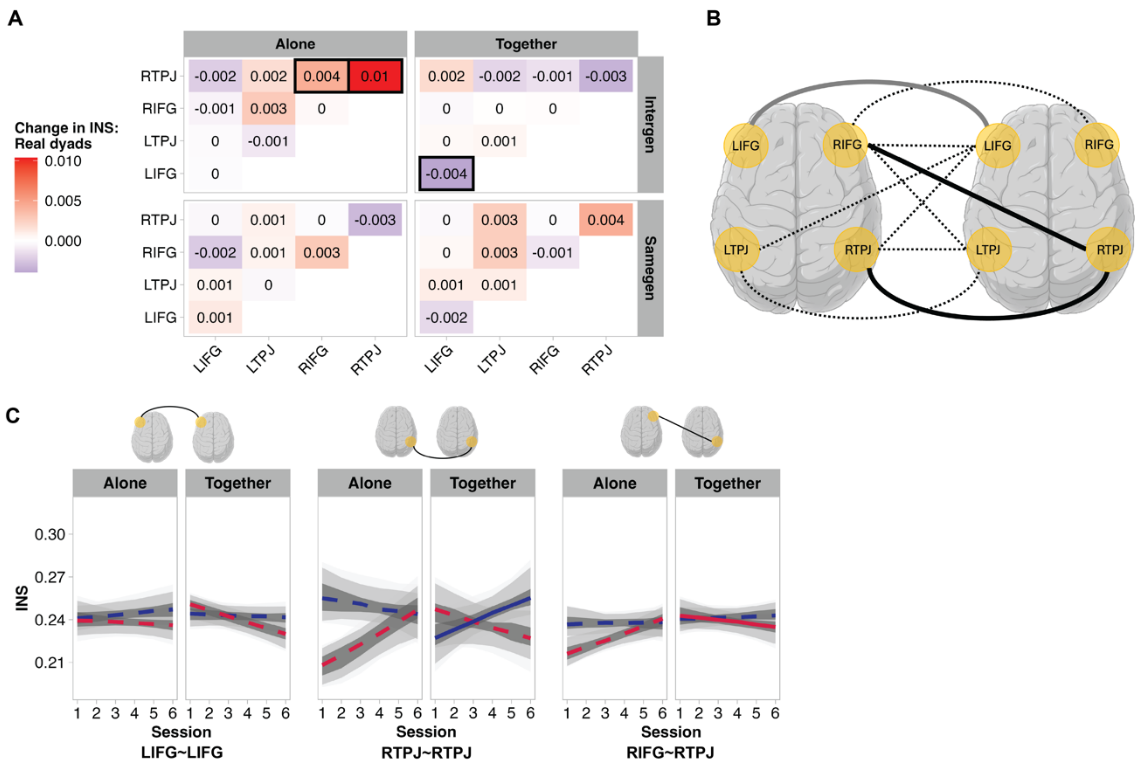


**S3 Appendix – Fig D.** HbR. A) Slope of change in INS across six sessions; black squares indicate that INS levels differ between real and pseudo dyads (i.e., 95% HPD of contrast distribution excludes 0). B) Visualisation of all possible ROI pairs (n=10). Dashed lines indicate ROI pairs where the slope of change in INS did not differ between real and pseudo dyads. Solid lines indicate ROI pairs where the slope of change in INS differed between real and pseudo dyads. Black solid lines indicate ROI pairs where task, group, and task*group contrasts showed differences. C) Slope of change in INS across sessions for all ROI pairs that differed between real and pseudo dyads, for which task, group, and task*group contrasts were computed. Solid lines indicate that the 95% HPD of an INS~Session slope excludes 0, indicating an association. Dashed lines indicate that the 95% HPD of an INS~Session slope includes 0, indicating no association. Grey shading shows 95%, 89%, and 50% intervals of the posterior predictive distributions. LIFG~LIFG: No differences. RTPJ~RTPJ: Interactions, together < alone for intergenerational group and intergenerational > same generation for alone. RIFG~RTPJ: Interaction, together < alone for intergenerational group. Full list of point estimates from models and contrasts with 95% HPDs available in Tables C-D in S6 Appendix. Brains created in BioRender. Moffat, R. (2026) https://BioRender.com/6p18lgg. A and C were created using code shared in the OSF repository "Longitudinal perspectives on intergenerational inter-brain synchrony": https://osf.io/xcgp6. Code for A: https://osf.io/xcgp6/files/pu7h2. Code for C: https://osf.io/xcgp6/files/4f8qw.

**Supplementary Materials:** Moffat, Dumas and Cross (2026). *Social interactions between people of same and different generations shape longitudinal changes in interpersonal neural synchrony, loneliness, and social connection.*

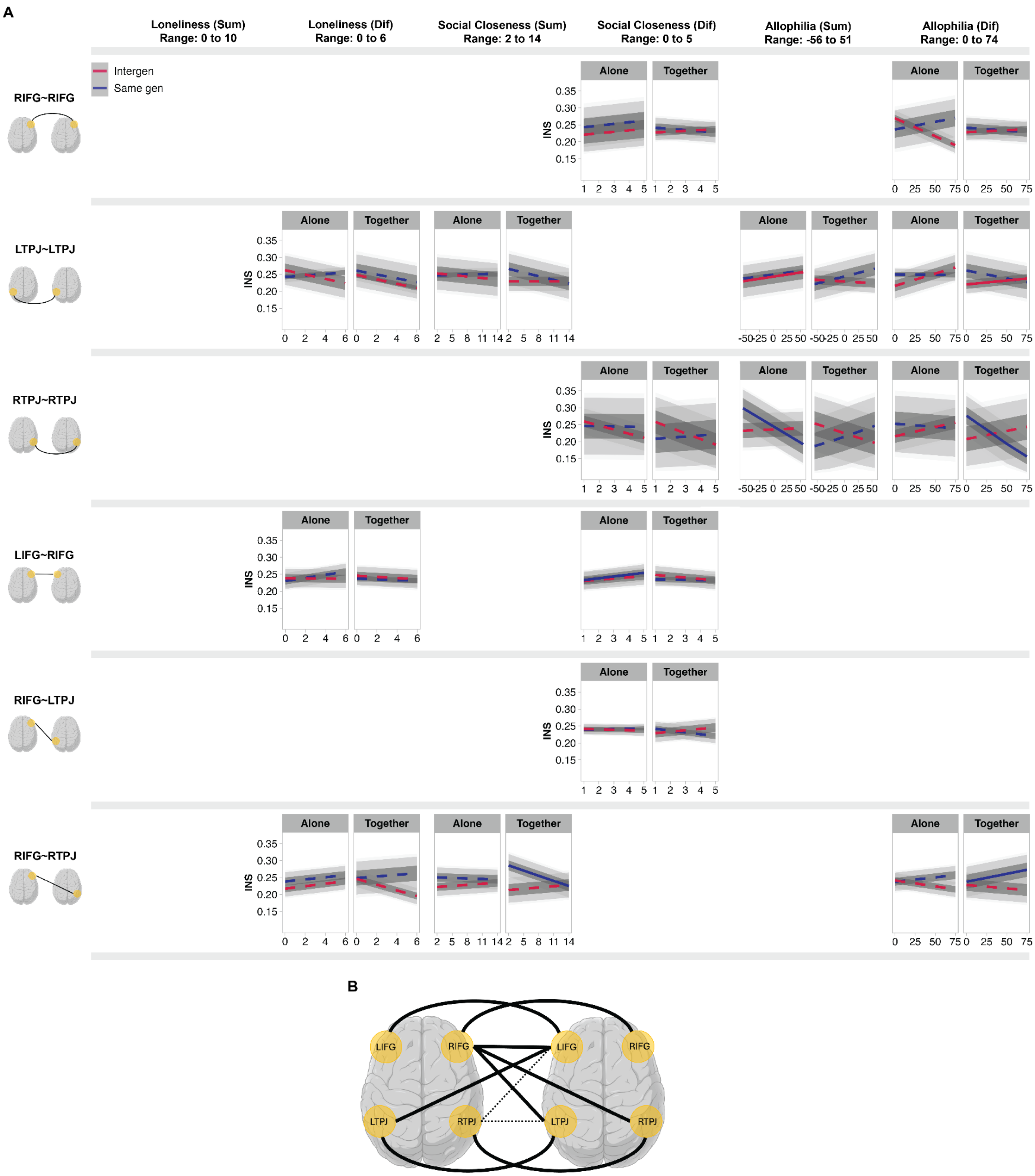

**S3 Appendix – Fig E.** HbR. A) Slope of change in INS across levels of self-report measures for ROI pairs with at least one real vs. pseudo difference revealed by contrasts. Lines indicate whether the 95% HPD for each INS~Self-report measure slope excludes 0 (solid), indicating an association, or includes 0 (dashed), indicating no association. Shading shows 95%, 89%, and 50% intervals for the posterior predictive distributions. Results of task, group, and task*group contrasts for the panels above are as follows. *RIFG~RIFG Social Closeness (Sum):* Group difference, positive relationship for intergenerational dyads only. *RIFG~RIFG Allophilia (Dif):* Interaction, intergenerational < same generation dyads for drawing alone. *LTPJ~LTPJ Loneliness (Dif):* Negative relationship. Group difference, negative relationship remains for intergenerational dyads only. Task difference, negative relationship remains for drawing together only. *LTPJ~LTPJ Social Closeness (Sum):* Interaction difference, intergenerational < same generation dyads for drawing alone. *LTPJ~LTPJ Allophilia (Sum):* Interaction, drawing alone > together for intergenerational dyads. *LTPJ~LTPJ Allophilia (Dif):* Group difference, negative relationship for intergenerational dyads and intergenerational > same generation dyads. Interaction, intergenerational > same generation dyads for drawing together. *RTPJ~RTPJ Social Closeness (Dif):* Interaction, intergenerational < same generation dyads for drawing together. *RTPJ~RTPJ Allophilia (Sum):* Task difference, drawing alone < together. Interaction, drawing alone < together for same generation dyads. *RTPJ~RTPJ Allophilia (Dif):* Interaction, intergenerational > same generation dyads for drawing together. *LIFG~RIFG Loneliness (Dif):* Negative relationship. Group difference, negative relationship for same generation dyads. Task difference, negative relationship for drawing alone. *LIFG~RIFG Social Closeness (Dif):* Task difference, positive relationship for drawing alone and drawing alone > together. *RIFG~LTPJ Social Closeness (Dif):* Interaction, intergenerational > same generation dyads for drawing together. *RIFG~RTPJ Loneliness (Dif):* Task difference, drawing alone > together. *RIFG~RTPJ Social Closeness (Sum):* Group difference, intergenerational > same generation dyads. Interaction, intergenerational > same generation dyads for drawing together. *RIFG~RTPJ Allophilia (Dif):* No differences. B) Visualisation of all possible ROI pairs (n=10). Dashed lines indicate ROI pairs where i) neither INS for all sessions combined nor ii) the slope of change in INS differed between real and pseudo dyads. Solid lines indicate ROI pairs where INS (for sessions combined or slope of change across sessions) differed between real and pseudo dyads. Black solid lines indicate ROI pairs where task, group, and task*group contrasts showed differences. Brains created in BioRender. Moffat, R. (2026) https://BioRender.com/6p18lgg. This figure was created using code shared in the OSF repository "Longitudinal perspectives on intergenerational inter-brain synchrony": https://osf.io/xcgp6. Code for individual panels in this figure: https://osf.io/xcgp6/files/ptyhe.

**Supplementary Materials:** Moffat, Dumas and Cross (2026). *Social interactions between people of same and different generations shape longitudinal changes in interpersonal neural synchrony, loneliness, and social connection.*

**S4 Appendix – Table A.** Comparison of preregistered predictions/analysis plan and the analyses undertaken to date. Preregistration is available on the Open Science Framework (https://osf.io/hz6tm). In this preregistration, we stated our plan to disseminate our findings across multiple publications.

| # | Prediction | Status of analysis |
|---|---|---|
| 1. | We predict that IOS and allophilia will increase across sessions, and that loneliness will decrease (simple terms). These effects may be more pronounced in the intergenerational group (2-way interaction). | Presented in Results of main manuscript. |
| 2. | We predict that the degree of synchrony in the dyad's drawing-hand movements and the dyad's head movements will increase across sessions for joint drawing, but not solo drawing (2-way interaction). This effect may be stronger for intergenerational pairs than same generation pairs (3-way interaction). | Movement data have not yet been analysed. Analysis not yet performed. |
| 3. | We predict that the degree of synchrony in the dyad's drawing-hand movements and the dyad's head movements [while drawing together] will be positively associated with the self-in-other scores, in a way that does not change across sessions (2-way interaction). This effect may be stronger for intergenerational pairs than same generation pairs (3-way interaction). | Movement data have not yet been analysed. Analysis not yet performed. |
| 4. | We predict congruent increases in INS and movement synchrony for drawing together, relative to alone (2-way interaction). We will examine all ROI pairs, have not formed strong predictions (3-way interaction). These effects may grow stronger across sessions and/or differ between groups (4-/5-way interaction). | Movement data have not yet been analysed. Analysis not yet performed. |
| 5. | We predict that INS during join drawing will be associated with IOS, allophilia, and loneliness (2-way interactions). These will likely become stronger across sessions and/or differ between groups (3-/4-way interactions) | Relationships between self-report measures and INS reported in Results of main manuscript (3-way interaction only). We did not pursue the 4-way interaction as we realized that the interpretability of these results would be low, particularly without directional hypotheses. |
| 6. | We predict that INS during join drawing will be associated with ratings of coherence and temporal overlap during drawing (2-way interactions). These effects will likely become stronger across sessions and/or differ between groups (3-/4-way interactions). | During data collection, the research team discussed switching from subjective experimenter ratings of drawing coherence to ratings from a less partial population with minimal knowledge of the study, as well as objective kinematic measures of temporal overlap. Ratings of the drawing coherence (called 'collaboration visible in drawings') have since been collected (Moffat et al., 2026). Hand overlap can be extracted from movement coordinates in future analyses. |

**S4 Appendix – Table B.** Breakdown of reasons for excluding dyads data. Recordings/dyads only included once in list. E.g., if a recording/dyad is counted among reasons unrelated to signal quality, they are not re-counted among reasons related to signal quality. In total 91 recordings of 732 recordings were excluded, impacting 85 of 366 dyadic sessions.

| Reason | N individual recordings excluded | N dyadic sessions impacted |
|---|---|---|
| ***Unrelated to signal quality*** | | |
| Recordings lost in transition between recording laptops | 2 | 2 |
| Triggers missing or unreliable | 7 | 4 |
| Participant unable to complete session (could not attend final session/consumed alcohol) | 5 | 3 |
| Multi-part recording (bluetooth or battery dropout) | 6 | 4 |
| ***Related to signal quality*** | | |
| No short channels of adequate quality | 71 | 69 |
| No long channels of adequate quality (pilot montage used) | 0 | 0 |
| **Total** | **91** | **85** |

**S4 Appendix – Table C.** Number of pseudo dyads per session, group and ROI.

| ROIs | Group | Session 1 | Session 2 | Session 3 | Session 4 | Session 5 | Session 6 |
|---|---|---|---|---|---|---|---|
| LIFG~RIFG | Intergen | 3386 | 3633 | 3980 | 4294 | 4323 | 3455 |
| LIFG~RIFG | Samegen | 5434 | 6268 | 6354 | 6120 | 6676 | 6198 |
| LTPJ~RTPJ | Intergen | 1679 | 1892 | 2437 | 2868 | 3193 | 2478 |
| LTPJ~RTPJ | Samegen | 3800 | 4726 | 4474 | 5003 | 5472 | 5024 |
| LIFG~LIFG | Intergen | 1754 | 1970 | 2083 | 2215 | 2188 | 1818 |
| LIFG~LIFG | Samegen | 2680 | 3378 | 3216 | 3137 | 3297 | 3137 |
| LTPJ~LTPJ | Intergen | 1726 | 1888 | 1977 | 2242 | 2188 | 1818 |
| LTPJ~LTPJ | Samegen | 2828 | 3378 | 3138 | 3217 | 3298 | 3137 |
| LIFG~LTPJ | Intergen | 3482 | 3858 | 4059 | 4457 | 4376 | 3636 |
| LIFG~LTPJ | Samegen | 5508 | 6756 | 6355 | 6355 | 6597 | 6277 |
| LIFG~RTPJ | Intergen | 1716 | 1949 | 2507 | 2854 | 3193 | 2478 |
| LIFG~RTPJ | Samegen | 3699 | 4726 | 4530 | 4941 | 5471 | 5025 |
| RIFG~RIFG | Intergen | 1630 | 1675 | 1899 | 2082 | 2134 | 1639 |
| RIFG~RIFG | Samegen | 2753 | 2905 | 3137 | 2982 | 3378 | 3059 |
| RTPJ~RTPJ | Intergen | 352 | 403 | 734 | 875 | 1145 | 820 |
| RTPJ~RTPJ | Samegen | 1270 | 1647 | 1588 | 1939 | 2264 | 2005 |
| RIFG~LTPJ | Intergen | 3355 | 3557 | 3876 | 4321 | 4323 | 3455 |
| RIFG~LTPJ | Samegen | 5583 | 6268 | 6277 | 6197 | 6677 | 6199 |
| RIFG~RTPJ | Intergen | 1622 | 1794 | 2382 | 2765 | 3144 | 2339 |
| RIFG~RTPJ | Samegen | 3750 | 4384 | 4473 | 4818 | 5537 | 4961 |

**S4 Appendix – Table D.** The weakly informed priors used to impose a constrained distribution on expected results.

| Outcome measure | Numeric range | Syntax for priors |
|---|---|---|
| INS<br>Real and pseudo dyads | 0 to 1 | weakly_informative_priors <- c(<br>set_prior('normal(0, 1)', class = 'Intercept'),<br>set_prior('normal(0, .5)', class = 'b')) |
| INS<br>Real dyads only | 0 to 1 | weakly_informative_priors <- c(<br>set_prior('normal(0, 1)', class = 'Intercept'),<br>set_prior('normal(0, .5)', class = 'b'),<br>set_prior('normal(0, .01)', class = 'sd')) |
| Loneliness | 0 to 6 | weakly_informative_priors <- c(<br>set_prior('normal(3, 1.5)', class = 'Intercept'),<br>set_prior('normal(0, 1)', class = 'b'),<br>set_prior('normal(0, .1)', class = 'sd')) |
| Social closeness | 1 to 7 | weakly_informative_priors <- c(<br>set_prior('normal(3, 5)', class = 'Intercept'),<br>set_prior('normal(0, 5)', class = 'b'),<br>set_prior('normal(0, 1)', class = 'sd')) |
| Allophilia | - 85 to 85 | weakly_informative_priors <- c(<br>set_prior('normal(0, 10)', class = 'Intercept'),<br>set_prior('normal(0, 5)', class = 'b'),<br>set_prior('normal(0, 1)', class = 'sd')) |

**S5 Appendix – Table A.** HbO. Estimates of INS levels for all sessions combined, for real and pseudo dyads per ROI pair and task in unstandardised units.

| Dyad Type | Group | ROI Pair | Task | Estimate | Lower HPD | Upper HPD |
|---|---|---|---|---|---|---|
| Pseudo | Intergen | LIFG~RIFG | Alone | 0.24 | 0.239 | 0.24 |
| Real | Intergen | LIFG~RIFG | Alone | 0.242 | 0.237 | 0.247 |
| Pseudo | Intergen | LIFG~RIFG | Together | 0.241 | 0.24 | 0.241 |
| Real | Intergen | LIFG~RIFG | Together | 0.248 | 0.244 | 0.251 |
| Pseudo | Intergen | LTPJ~RTPJ | Alone | 0.24 | 0.239 | 0.241 |
| Real | Intergen | LTPJ~RTPJ | Alone | 0.242 | 0.236 | 0.248 |
| Pseudo | Intergen | LTPJ~RTPJ | Together | 0.241 | 0.241 | 0.242 |
| Real | Intergen | LTPJ~RTPJ | Together | 0.245 | 0.24 | 0.25 |
| Pseudo | Intergen | LIFG~LIFG | Alone | 0.241 | 0.24 | 0.242 |
| Real | Intergen | LIFG~LIFG | Alone | 0.241 | 0.235 | 0.248 |
| Pseudo | Intergen | LIFG~LIFG | Together | 0.241 | 0.241 | 0.242 |
| Real | Intergen | LIFG~LIFG | Together | 0.246 | 0.241 | 0.252 |
| Pseudo | Intergen | LTPJ~LTPJ | Alone | 0.242 | 0.241 | 0.243 |
| Real | Intergen | LTPJ~LTPJ | Alone | 0.244 | 0.237 | 0.25 |
| Pseudo | Intergen | LTPJ~LTPJ | Together | 0.243 | 0.243 | 0.244 |
| Real | Intergen | LTPJ~LTPJ | Together | 0.252 | 0.247 | 0.257 |
| Pseudo | Intergen | LIFG~LTPJ | Alone | 0.24 | 0.24 | 0.241 |
| Real | Intergen | LIFG~LTPJ | Alone | 0.243 | 0.239 | 0.248 |
| Pseudo | Intergen | LIFG~LTPJ | Together | 0.241 | 0.241 | 0.241 |
| Real | Intergen | LIFG~LTPJ | Together | 0.247 | 0.243 | 0.25 |
| Pseudo | Intergen | LIFG~RTPJ | Alone | 0.238 | 0.237 | 0.239 |
| Real | Intergen | LIFG~RTPJ | Alone | 0.239 | 0.233 | 0.245 |
| Pseudo | Intergen | LIFG~RTPJ | Together | 0.24 | 0.239 | 0.24 |
| Real | Intergen | LIFG~RTPJ | Together | 0.242 | 0.237 | 0.246 |
| Pseudo | Intergen | RIFG~RIFG | Alone | 0.239 | 0.238 | 0.239 |
| Real | Intergen | RIFG~RIFG | Alone | 0.248 | 0.241 | 0.255 |
| Pseudo | Intergen | RIFG~RIFG | Together | 0.24 | 0.239 | 0.24 |
| Real | Intergen | RIFG~RIFG | Together | 0.246 | 0.241 | 0.252 |
| Pseudo | Intergen | RTPJ~RTPJ | Alone | 0.238 | 0.237 | 0.239 |
| Real | Intergen | RTPJ~RTPJ | Alone | 0.248 | 0.236 | 0.259 |
| Pseudo | Intergen | RTPJ~RTPJ | Together | 0.239 | 0.238 | 0.24 |
| Real | Intergen | RTPJ~RTPJ | Together | 0.242 | 0.232 | 0.252 |
| Pseudo | Intergen | LTPJ~RIFG | Alone | 0.239 | 0.239 | 0.24 |
| Real | Intergen | LTPJ~RIFG | Alone | 0.241 | 0.236 | 0.245 |
| Pseudo | Intergen | LTPJ~RIFG | Together | 0.241 | 0.24 | 0.241 |
| Real | Intergen | LTPJ~RIFG | Together | 0.246 | 0.242 | 0.249 |
| Pseudo | Intergen | RIFG~RTPJ | Alone | 0.238 | 0.237 | 0.238 |
| Real | Intergen | RIFG~RTPJ | Alone | 0.237 | 0.231 | 0.243 |
| Pseudo | Intergen | RIFG~RTPJ | Together | 0.238 | 0.238 | 0.239 |
| Real | Intergen | RIFG~RTPJ | Together | 0.243 | 0.238 | 0.248 |
| Pseudo | Samegen | LIFG~RIFG | Alone | 0.242 | 0.241 | 0.242 |
| Real | Samegen | LIFG~RIFG | Alone | 0.243 | 0.238 | 0.247 |
| Pseudo | Samegen | LIFG~RIFG | Together | 0.243 | 0.243 | 0.243 |

| Dyad Type | Group | ROI Pair | Task | Estimate | Lower HPD | Upper HPD |
|---|---|---|---|---|---|---|
| Real | Samegen | LIFG~RIFG | Together | 0.247 | 0.244 | 0.251 |
| Pseudo | Samegen | LTPJ~RTPJ | Alone | 0.245 | 0.244 | 0.245 |
| Real | Samegen | LTPJ~RTPJ | Alone | 0.25 | 0.244 | 0.255 |
| Pseudo | Samegen | LTPJ~RTPJ | Together | 0.244 | 0.244 | 0.244 |
| Real | Samegen | LTPJ~RTPJ | Together | 0.248 | 0.243 | 0.252 |
| Pseudo | Samegen | LIFG~LIFG | Alone | 0.242 | 0.242 | 0.243 |
| Real | Samegen | LIFG~LIFG | Alone | 0.242 | 0.235 | 0.248 |
| Pseudo | Samegen | LIFG~LIFG | Together | 0.243 | 0.242 | 0.243 |
| Real | Samegen | LIFG~LIFG | Together | 0.243 | 0.238 | 0.248 |
| Pseudo | Samegen | LTPJ~LTPJ | Alone | 0.245 | 0.244 | 0.245 |
| Real | Samegen | LTPJ~LTPJ | Alone | 0.242 | 0.235 | 0.249 |
| Pseudo | Samegen | LTPJ~LTPJ | Together | 0.246 | 0.245 | 0.246 |
| Real | Samegen | LTPJ~LTPJ | Together | 0.247 | 0.242 | 0.252 |
| Pseudo | Samegen | LIFG~LTPJ | Alone | 0.241 | 0.241 | 0.242 |
| Real | Samegen | LIFG~LTPJ | Alone | 0.241 | 0.237 | 0.246 |
| Pseudo | Samegen | LIFG~LTPJ | Together | 0.243 | 0.242 | 0.243 |
| Real | Samegen | LIFG~LTPJ | Together | 0.243 | 0.239 | 0.246 |
| Pseudo | Samegen | LIFG~RTPJ | Alone | 0.241 | 0.241 | 0.242 |
| Real | Samegen | LIFG~RTPJ | Alone | 0.244 | 0.238 | 0.249 |
| Pseudo | Samegen | LIFG~RTPJ | Together | 0.241 | 0.241 | 0.242 |
| Real | Samegen | LIFG~RTPJ | Together | 0.244 | 0.239 | 0.248 |
| Pseudo | Samegen | RIFG~RIFG | Alone | 0.241 | 0.241 | 0.242 |
| Real | Samegen | RIFG~RIFG | Alone | 0.241 | 0.234 | 0.247 |
| Pseudo | Samegen | RIFG~RIFG | Together | 0.243 | 0.242 | 0.243 |
| Real | Samegen | RIFG~RIFG | Together | 0.246 | 0.241 | 0.251 |
| Pseudo | Samegen | RTPJ~RTPJ | Alone | 0.244 | 0.243 | 0.245 |
| Real | Samegen | RTPJ~RTPJ | Alone | 0.245 | 0.235 | 0.255 |
| Pseudo | Samegen | RTPJ~RTPJ | Together | 0.242 | 0.241 | 0.243 |
| Real | Samegen | RTPJ~RTPJ | Together | 0.251 | 0.242 | 0.26 |
| Pseudo | Samegen | LTPJ~RIFG | Alone | 0.242 | 0.241 | 0.242 |
| Real | Samegen | LTPJ~RIFG | Alone | 0.241 | 0.237 | 0.246 |
| Pseudo | Samegen | LTPJ~RIFG | Together | 0.243 | 0.243 | 0.244 |
| Real | Samegen | LTPJ~RIFG | Together | 0.246 | 0.243 | 0.25 |
| Pseudo | Samegen | RIFG~RTPJ | Alone | 0.242 | 0.241 | 0.242 |
| Real | Samegen | RIFG~RTPJ | Alone | 0.239 | 0.233 | 0.244 |
| Pseudo | Samegen | RIFG~RTPJ | Together | 0.242 | 0.242 | 0.242 |
| Real | Samegen | RIFG~RTPJ | Together | 0.247 | 0.242 | 0.251 |

**S5 Appendix – Table B.** HbO. Contrasts between real and pseudo dyads' INS levels for all sessions combined per ROI pair and task in unstandardised units.

| Contrast | Group | ROI Pair | Task | Estimate | Lower HPD | Upper HPD | No overlap |
|---|---|---|---|---|---|---|---|
| Real - Pseudo | Intergen | LIFG~RIFG | Alone | -0.002 | -0.007 | 0.003 | |
| Real - Pseudo | Intergen | LIFG~RIFG | Together | -0.007 | -0.011 | -0.003 | No overlap |
| Real - Pseudo | Intergen | LTPJ~RTPJ | Alone | -0.002 | -0.008 | 0.004 | |
| Real - Pseudo | Intergen | LTPJ~RTPJ | Together | -0.004 | -0.009 | 0.001 | |
| Real - Pseudo | Intergen | LIFG~LIFG | Alone | 0 | -0.007 | 0.006 | |
| Real - Pseudo | Intergen | LIFG~LIFG | Together | -0.005 | -0.01 | 0 | No overlap |
| Real - Pseudo | Intergen | LTPJ~LTPJ | Alone | -0.002 | -0.008 | 0.005 | |
| Real - Pseudo | Intergen | LTPJ~LTPJ | Together | -0.009 | -0.014 | -0.004 | No overlap |
| Real - Pseudo | Intergen | LIFG~LTPJ | Alone | -0.003 | -0.008 | 0.001 | |
| Real - Pseudo | Intergen | LIFG~LTPJ | Together | -0.006 | -0.009 | -0.002 | No overlap |
| Real - Pseudo | Intergen | LIFG~RTPJ | Alone | -0.001 | -0.007 | 0.005 | |
| Real - Pseudo | Intergen | LIFG~RTPJ | Together | -0.002 | -0.007 | 0.003 | |
| Real - Pseudo | Intergen | RIFG~RIFG | Alone | -0.009 | -0.016 | -0.002 | No overlap |
| Real - Pseudo | Intergen | RIFG~RIFG | Together | -0.007 | -0.012 | -0.001 | No overlap |
| Real - Pseudo | Intergen | RTPJ~RTPJ | Alone | -0.01 | -0.021 | 0.002 | |
| Real - Pseudo | Intergen | RTPJ~RTPJ | Together | -0.003 | -0.013 | 0.007 | |
| Real - Pseudo | Intergen | LTPJ~RIFG | Alone | -0.002 | -0.006 | 0.003 | |
| Real - Pseudo | Intergen | LTPJ~RIFG | Together | -0.005 | -0.009 | -0.001 | No overlap |
| Real - Pseudo | Intergen | RIFG~RTPJ | Alone | 0 | -0.006 | 0.006 | |
| Real - Pseudo | Intergen | RIFG~RTPJ | Together | -0.004 | -0.009 | 0.001 | |
| Real - Pseudo | Samegen | LIFG~RIFG | Alone | -0.001 | -0.006 | 0.004 | |
| Real - Pseudo | Samegen | LIFG~RIFG | Together | -0.004 | -0.008 | -0.001 | No overlap |
| Real - Pseudo | Samegen | LTPJ~RTPJ | Alone | -0.005 | -0.011 | 0.001 | |
| Real - Pseudo | Samegen | LTPJ~RTPJ | Together | -0.004 | -0.008 | 0.001 | |
| Real - Pseudo | Samegen | LIFG~LIFG | Alone | 0 | -0.006 | 0.007 | |
| Real - Pseudo | Samegen | LIFG~LIFG | Together | -0.001 | -0.006 | 0.005 | |
| Real - Pseudo | Samegen | LTPJ~LTPJ | Alone | 0.002 | -0.004 | 0.009 | |
| Real - Pseudo | Samegen | LTPJ~LTPJ | Together | -0.001 | -0.006 | 0.004 | |
| Real - Pseudo | Samegen | LIFG~LTPJ | Alone | 0 | -0.005 | 0.005 | |
| Real - Pseudo | Samegen | LIFG~LTPJ | Together | 0 | -0.004 | 0.003 | |
| Real - Pseudo | Samegen | LIFG~RTPJ | Alone | -0.003 | -0.008 | 0.003 | |
| Real - Pseudo | Samegen | LIFG~RTPJ | Together | -0.002 | -0.007 | 0.002 | |
| Real - Pseudo | Samegen | RIFG~RIFG | Alone | 0.001 | -0.006 | 0.007 | |
| Real - Pseudo | Samegen | RIFG~RIFG | Together | -0.003 | -0.009 | 0.002 | |
| Real - Pseudo | Samegen | RTPJ~RTPJ | Alone | -0.001 | -0.011 | 0.009 | |
| Real - Pseudo | Samegen | RTPJ~RTPJ | Together | -0.009 | -0.017 | 0 | |
| Real - Pseudo | Samegen | LTPJ~RIFG | Alone | 0 | -0.004 | 0.005 | |
| Real - Pseudo | Samegen | LTPJ~RIFG | Together | -0.003 | -0.006 | 0.001 | |
| Real - Pseudo | Samegen | RIFG~RTPJ | Alone | 0.003 | -0.003 | 0.008 | |
| Real - Pseudo | Samegen | RIFG~RTPJ | Together | -0.005 | -0.009 | 0 | No overlap |

**S5 Appendix – Table C.** HbO. Estimates of slope of change in INS across sessions for real and pseudo dyads per ROI pair and task in unstandardised units.

| Dyad Type | Group | ROI Pair | Task | Slope estimate | Lower HPD | Upper HPD | No overlap |
|---|---|---|---|---|---|---|---|
| Pseudo | Intergen | LIFG~RIFG | Alone | 0 | 0 | 0.001 | No overlap |
| Real | Intergen | LIFG~RIFG | Alone | 0 | -0.003 | 0.003 | |
| Pseudo | Intergen | LIFG~RIFG | Together | 0 | 0 | 0.001 | No overlap |
| Real | Intergen | LIFG~RIFG | Together | 0.001 | -0.002 | 0.004 | |
| Pseudo | Intergen | LTPJ~RTPJ | Alone | 0 | 0 | 0.001 | No overlap |
| Real | Intergen | LTPJ~RTPJ | Alone | 0.003 | 0 | 0.007 | |
| Pseudo | Intergen | LTPJ~RTPJ | Together | 0 | -0.001 | 0 | |
| Real | Intergen | LTPJ~RTPJ | Together | -0.003 | -0.006 | 0 | |
| Pseudo | Intergen | LIFG~LIFG | Alone | 0 | 0 | 0.001 | |
| Real | Intergen | LIFG~LIFG | Alone | -0.002 | -0.006 | 0.002 | |
| Pseudo | Intergen | LIFG~LIFG | Together | 0 | 0 | 0 | |
| Real | Intergen | LIFG~LIFG | Together | -0.001 | -0.004 | 0.003 | |
| Pseudo | Intergen | LTPJ~LTPJ | Alone | 0.001 | 0 | 0.001 | No overlap |
| Real | Intergen | LTPJ~LTPJ | Alone | -0.002 | -0.006 | 0.002 | |
| Pseudo | Intergen | LTPJ~LTPJ | Together | -0.001 | -0.001 | 0 | No overlap |
| Real | Intergen | LTPJ~LTPJ | Together | 0.004 | 0 | 0.007 | |
| Pseudo | Intergen | LIFG~LTPJ | Alone | 0 | 0 | 0 | |
| Real | Intergen | LIFG~LTPJ | Alone | 0 | -0.003 | 0.003 | |
| Pseudo | Intergen | LIFG~LTPJ | Together | 0 | -0.001 | 0 | |
| Real | Intergen | LIFG~LTPJ | Together | -0.002 | -0.004 | 0.001 | |
| Pseudo | Intergen | LIFG~RTPJ | Alone | 0 | 0 | 0.001 | |
| Real | Intergen | LIFG~RTPJ | Alone | 0.004 | 0 | 0.007 | No overlap |
| Pseudo | Intergen | LIFG~RTPJ | Together | 0 | 0 | 0 | |
| Real | Intergen | LIFG~RTPJ | Together | -0.002 | -0.005 | 0.001 | |
| Pseudo | Intergen | RIFG~RIFG | Alone | 0.001 | 0 | 0.001 | No overlap |
| Real | Intergen | RIFG~RIFG | Alone | 0.002 | -0.002 | 0.006 | |
| Pseudo | Intergen | RIFG~RIFG | Together | 0.001 | 0.001 | 0.002 | No overlap |
| Real | Intergen | RIFG~RIFG | Together | 0.002 | -0.001 | 0.006 | |
| Pseudo | Intergen | RTPJ~RTPJ | Alone | 0 | 0 | 0.001 | |
| Real | Intergen | RTPJ~RTPJ | Alone | 0.006 | -0.002 | 0.013 | |
| Pseudo | Intergen | RTPJ~RTPJ | Together | 0 | -0.001 | 0 | |
| Real | Intergen | RTPJ~RTPJ | Together | -0.005 | -0.013 | 0.002 | |
| Pseudo | Intergen | LTPJ~RIFG | Alone | 0.001 | 0 | 0.001 | No overlap |
| Real | Intergen | LTPJ~RIFG | Alone | 0.002 | -0.001 | 0.005 | |
| Pseudo | Intergen | LTPJ~RIFG | Together | 0 | 0 | 0.001 | No overlap |
| Real | Intergen | LTPJ~RIFG | Together | -0.001 | -0.003 | 0.002 | |
| Pseudo | Intergen | RIFG~RTPJ | Alone | 0.001 | 0 | 0.001 | No overlap |
| Real | Intergen | RIFG~RTPJ | Alone | 0.003 | 0 | 0.007 | |
| Pseudo | Intergen | RIFG~RTPJ | Together | 0.001 | 0.001 | 0.002 | No overlap |
| Real | Intergen | RIFG~RTPJ | Together | -0.003 | -0.007 | 0 | |
| Pseudo | Samegen | LIFG~RIFG | Alone | 0 | 0 | 0 | |
| Real | Samegen | LIFG~RIFG | Alone | -0.002 | -0.005 | 0.001 | |
| Pseudo | Samegen | LIFG~RIFG | Together | 0 | 0 | 0 | |

| Dyad Type | Group | ROI Pair | Task | Slope Estimate | Lower HPD | Upper HPD | No overlap |
|---|---|---|---|---|---|---|---|
| Real | Samegen | LIFG~RIFG | Together | 0.001 | -0.001 | 0.004 | |
| Pseudo | Samegen | LTPJ~RTPJ | Alone | 0 | 0 | 0.001 | No overlap |
| Real | Samegen | LTPJ~RTPJ | Alone | 0 | -0.004 | 0.003 | |
| Pseudo | Samegen | LTPJ~RTPJ | Together | 0.001 | 0 | 0.001 | No overlap |
| Real | Samegen | LTPJ~RTPJ | Together | 0.001 | -0.002 | 0.005 | |
| Pseudo | Samegen | LIFG~LIFG | Alone | 0 | 0 | 0.001 | |
| Real | Samegen | LIFG~LIFG | Alone | -0.003 | -0.007 | 0.001 | |
| Pseudo | Samegen | LIFG~LIFG | Together | 0 | 0 | 0.001 | No overlap |
| Real | Samegen | LIFG~LIFG | Together | 0.002 | -0.002 | 0.006 | |
| Pseudo | Samegen | LTPJ~LTPJ | Alone | 0 | 0 | 0.001 | No overlap |
| Real | Samegen | LTPJ~LTPJ | Alone | -0.001 | -0.004 | 0.003 | |
| Pseudo | Samegen | LTPJ~LTPJ | Together | 0 | -0.001 | 0 | |
| Real | Samegen | LTPJ~LTPJ | Together | -0.001 | -0.004 | 0.003 | |
| Pseudo | Samegen | LIFG~LTPJ | Alone | 0 | 0 | 0 | |
| Real | Samegen | LIFG~LTPJ | Alone | 0.001 | -0.002 | 0.004 | |
| Pseudo | Samegen | LIFG~LTPJ | Together | 0 | 0 | 0 | |
| Real | Samegen | LIFG~LTPJ | Together | 0.003 | 0.001 | 0.006 | No overlap |
| Pseudo | Samegen | LIFG~RTPJ | Alone | 0.001 | 0 | 0.001 | No overlap |
| Real | Samegen | LIFG~RTPJ | Alone | -0.002 | -0.006 | 0.001 | |
| Pseudo | Samegen | LIFG~RTPJ | Together | 0.001 | 0.001 | 0.001 | No overlap |
| Real | Samegen | LIFG~RTPJ | Together | 0.002 | -0.001 | 0.005 | |
| Pseudo | Samegen | RIFG~RIFG | Alone | -0.001 | -0.001 | 0 | No overlap |
| Real | Samegen | RIFG~RIFG | Alone | 0 | -0.004 | 0.004 | |
| Pseudo | Samegen | RIFG~RIFG | Together | 0 | 0 | 0 | |
| Real | Samegen | RIFG~RIFG | Together | -0.002 | -0.005 | 0.002 | |
| Pseudo | Samegen | RTPJ~RTPJ | Alone | 0 | 0 | 0.001 | |
| Real | Samegen | RTPJ~RTPJ | Alone | 0.003 | -0.004 | 0.009 | |
| Pseudo | Samegen | RTPJ~RTPJ | Together | 0.002 | 0.002 | 0.003 | No overlap |
| Real | Samegen | RTPJ~RTPJ | Together | 0.009 | 0.002 | 0.016 | No overlap |
| Pseudo | Samegen | LTPJ~RIFG | Alone | 0 | 0 | 0 | |
| Real | Samegen | LTPJ~RIFG | Alone | 0.001 | -0.002 | 0.004 | |
| Pseudo | Samegen | LTPJ~RIFG | Together | 0 | 0 | 0 | |
| Real | Samegen | LTPJ~RIFG | Together | -0.002 | -0.005 | 0 | |
| Pseudo | Samegen | RIFG~RTPJ | Alone | 0 | 0 | 0 | |
| Real | Samegen | RIFG~RTPJ | Alone | -0.002 | -0.006 | 0.001 | |
| Pseudo | Samegen | RIFG~RTPJ | Together | 0.001 | 0.001 | 0.001 | No overlap |
| Real | Samegen | RIFG~RTPJ | Together | 0.002 | -0.001 | 0.006 | |

**S5 Appendix – Table D.** HbO. Contrasts between real and pseudo dyads' slopes of change in INS across sessions per ROI pair and task in unstandardised units.

| Contrast | Group | ROI Pair | Task | Estimate | Lower HPD | Upper HPD | No overlap |
|---|---|---|---|---|---|---|---|
| Real - Pseudo | Intergen | LIFG~RIFG | Alone | 0 | -0.002 | 0.003 | |
| Real - Pseudo | Intergen | LIFG~RIFG | Together | -0.001 | -0.003 | 0.002 | |
| Real - Pseudo | Intergen | LTPJ~RTPJ | Alone | -0.003 | -0.006 | 0.001 | |
| Real - Pseudo | Intergen | LTPJ~RTPJ | Together | 0.003 | -0.001 | 0.006 | |
| Real - Pseudo | Intergen | LIFG~LIFG | Alone | 0.002 | -0.002 | 0.006 | |
| Real - Pseudo | Intergen | LIFG~LIFG | Together | 0.001 | -0.003 | 0.004 | |
| Real - Pseudo | Intergen | LTPJ~LTPJ | Alone | 0.003 | -0.001 | 0.007 | |
| Real - Pseudo | Intergen | LTPJ~LTPJ | Together | -0.004 | -0.008 | -0.001 | No overlap |
| Real - Pseudo | Intergen | LIFG~LTPJ | Alone | 0 | -0.003 | 0.003 | |
| Real - Pseudo | Intergen | LIFG~LTPJ | Together | 0.002 | -0.001 | 0.004 | |
| Real - Pseudo | Intergen | LIFG~RTPJ | Alone | -0.004 | -0.007 | 0 | No overlap |
| Real - Pseudo | Intergen | LIFG~RTPJ | Together | 0.002 | -0.001 | 0.005 | |
| Real - Pseudo | Intergen | RIFG~RIFG | Alone | -0.001 | -0.005 | 0.003 | |
| Real - Pseudo | Intergen | RIFG~RIFG | Together | -0.001 | -0.005 | 0.003 | |
| Real - Pseudo | Intergen | RTPJ~RTPJ | Alone | -0.005 | -0.012 | 0.002 | |
| Real - Pseudo | Intergen | RTPJ~RTPJ | Together | 0.005 | -0.002 | 0.012 | |
| Real - Pseudo | Intergen | LTPJ~RIFG | Alone | -0.001 | -0.004 | 0.002 | |
| Real - Pseudo | Intergen | LTPJ~RIFG | Together | 0.001 | -0.001 | 0.004 | |
| Real - Pseudo | Intergen | RIFG~RTPJ | Alone | -0.002 | -0.006 | 0.001 | |
| Real - Pseudo | Intergen | RIFG~RTPJ | Together | 0.005 | 0.001 | 0.008 | No overlap |
| Real - Pseudo | Samegen | LIFG~RIFG | Alone | 0.002 | -0.001 | 0.005 | |
| Real - Pseudo | Samegen | LIFG~RIFG | Together | -0.001 | -0.003 | 0.002 | |
| Real - Pseudo | Samegen | LTPJ~RTPJ | Alone | 0.001 | -0.003 | 0.004 | |
| Real - Pseudo | Samegen | LTPJ~RTPJ | Together | -0.001 | -0.004 | 0.003 | |
| Real - Pseudo | Samegen | LIFG~LIFG | Alone | 0.003 | -0.001 | 0.007 | |
| Real - Pseudo | Samegen | LIFG~LIFG | Together | -0.002 | -0.005 | 0.002 | |
| Real - Pseudo | Samegen | LTPJ~LTPJ | Alone | 0.001 | -0.003 | 0.005 | |
| Real - Pseudo | Samegen | LTPJ~LTPJ | Together | 0 | -0.003 | 0.004 | |
| Real - Pseudo | Samegen | LIFG~LTPJ | Alone | -0.001 | -0.004 | 0.002 | |
| Real - Pseudo | Samegen | LIFG~LTPJ | Together | -0.003 | -0.006 | 0 | No overlap |
| Real - Pseudo | Samegen | LIFG~RTPJ | Alone | 0.003 | 0 | 0.006 | |
| Real - Pseudo | Samegen | LIFG~RTPJ | Together | -0.001 | -0.004 | 0.002 | |
| Real - Pseudo | Samegen | RIFG~RIFG | Alone | -0.001 | -0.005 | 0.003 | |
| Real - Pseudo | Samegen | RIFG~RIFG | Together | 0.002 | -0.002 | 0.006 | |
| Real - Pseudo | Samegen | RTPJ~RTPJ | Alone | -0.002 | -0.009 | 0.005 | |
| Real - Pseudo | Samegen | RTPJ~RTPJ | Together | -0.007 | -0.014 | 0 | No overlap |
| Real - Pseudo | Samegen | LTPJ~RIFG | Alone | -0.001 | -0.004 | 0.002 | |
| Real - Pseudo | Samegen | LTPJ~RIFG | Together | 0.002 | 0 | 0.005 | |
| Real - Pseudo | Samegen | RIFG~RTPJ | Alone | 0.002 | -0.001 | 0.006 | |
| Real - Pseudo | Samegen | RIFG~RTPJ | Together | -0.002 | -0.005 | 0.002 | |

**S5 Appendix – Table E.** HbO. Composition of non-homologous ROI pairs, to assess whether an ROI belonging to an older or younger member of a dyad influenced INS levels for all sessions (top) and across sessions (bottom). In contrast column, 'o' indicated that ROI belongs to older adult and 'y' indicates that ROI belongs to younger adult.

| Contrast | Task | Estimate | Lower HPD | Upper HPD | No overlap |
|---|---|---|---|---|---|
| *All sessions* | | | | | |
| o_l_ifg_y_l_tpj - o_l_tpj_y_l_ifg | Together | 0.007 | 0 | 0.014 | trend |
| o_l_ifg_y_r_ifg - o_r_ifg_y_l_ifg | Together | -0.001 | -0.009 | 0.006 | |
| o_l_tpj_y_r_ifg - o_r_ifg_y_l_tpj | Together | 0 | -0.007 | 0.008 | |
| *Across sessions* | | | | | |
| o_l_ifg_y_r_tpj - o_r_tpj_y_l_ifg | Alone | 0 | -0.011 | 0.012 | |
| o_r_ifg_y_r_tpj - o_r_tpj_y_r_ifg | Together | -0.002 | -0.014 | 0.009 | |

**S5 Appendix – Table F.** HbO. INS levels for all sessions: contrasts between tasks, groups and the task*group interaction per ROI pair in unstandardised units.

| Contrast | ROI Pair | Estimate | Lower HPD | Upper HPD | No overlap |
|---|---|---|---|---|---|
| Tog-Alo | LIFG~RIFG | 0.01 | 0.001 | 0.019 | No overlap |
| Int-Sam | LIFG~RIFG | -0.001 | -0.012 | 0.01 | |
| Tog-Alo Int | LIFG~RIFG | 0.006 | -0.001 | 0.012 | |
| Tog-Alo Sam | LIFG~RIFG | 0.004 | -0.002 | 0.011 | |
| Int-Sam Alo | LIFG~RIFG | -0.001 | -0.009 | 0.007 | |
| Int-Sam Tog | LIFG~RIFG | 0 | -0.006 | 0.006 | |
| Tog-Alo | LIFG~LIFG | 0.007 | -0.005 | 0.018 | |
| Int-Sam | LIFG~LIFG | 0.002 | -0.011 | 0.014 | |
| Tog-Alo Int | LIFG~LIFG | 0.005 | -0.003 | 0.014 | |
| Tog-Alo Sam | LIFG~LIFG | 0.001 | -0.007 | 0.01 | |
| Int-Sam Alo | LIFG~LIFG | -0.001 | -0.01 | 0.009 | |
| Int-Sam Tog | LIFG~LIFG | 0.003 | -0.004 | 0.01 | |
| Tog-Alo | LTPJ~LTPJ | 0.013 | 0.001 | 0.025 | No overlap |
| Int-Sam | LTPJ~LTPJ | 0.004 | -0.012 | 0.02 | |
| Tog-Alo Int | LTPJ~LTPJ | 0.009 | 0 | 0.017 | No overlap |
| Tog-Alo Sam | LTPJ~LTPJ | 0.004 | -0.004 | 0.013 | |
| Int-Sam Alo | LTPJ~LTPJ | 0 | -0.011 | 0.011 | |
| Int-Sam Tog | LTPJ~LTPJ | 0.004 | -0.005 | 0.013 | |
| Tog-Alo | LIFG~LTPJ | 0.005 | -0.004 | 0.013 | |
| Int-Sam | LIFG~LTPJ | 0.006 | -0.005 | 0.017 | |
| Tog-Alo Int | LIFG~LTPJ | 0.003 | -0.003 | 0.009 | |
| Tog-Alo Sam | LIFG~LTPJ | 0.001 | -0.005 | 0.008 | |
| Int-Sam Alo | LIFG~LTPJ | 0.002 | -0.005 | 0.01 | |
| Int-Sam Tog | LIFG~LTPJ | 0.004 | -0.002 | 0.01 | |
| Tog-Alo | RIFG~RIFG | 0.004 | -0.009 | 0.018 | |
| Int-Sam | RIFG~RIFG | 0.007 | -0.008 | 0.021 | |
| Tog-Alo Int | RIFG~RIFG | -0.002 | -0.011 | 0.008 | |
| Tog-Alo Sam | RIFG~RIFG | 0.006 | -0.003 | 0.015 | |
| Int-Sam Alo | RIFG~RIFG | 0.007 | -0.004 | 0.017 | |
| Int-Sam Tog | RIFG~RIFG | 0 | -0.009 | 0.008 | |
| Tog-Alo | RIFG~LTPJ | 0.01 | 0.001 | 0.018 | No overlap |
| Int-Sam | RIFG~LTPJ | -0.001 | -0.011 | 0.008 | |
| Tog-Alo Int | RIFG~LTPJ | 0.005 | -0.001 | 0.011 | |
| Tog-Alo Sam | RIFG~LTPJ | 0.005 | -0.001 | 0.011 | |
| Int-Sam Alo | RIFG~LTPJ | -0.001 | -0.008 | 0.006 | |
| Int-Sam Tog | RIFG~LTPJ | -0.001 | -0.006 | 0.005 | |
| Tog-Alo | RIFG~RTPJ | 0.013 | 0.002 | 0.024 | No overlap |
| Int-Sam | RIFG~RTPJ | -0.006 | -0.02 | 0.007 | |
| Tog-Alo Int | RIFG~RTPJ | 0.006 | -0.003 | 0.014 | |
| Tog-Alo Sam | RIFG~RTPJ | 0.008 | 0 | 0.015 | No overlap |
| Int-Sam Alo | RIFG~RTPJ | -0.002 | -0.012 | 0.007 | |
| Int-Sam Tog | RIFG~RTPJ | -0.004 | -0.012 | 0.004 | |

**S5 Appendix – Table G.** HbO. Contrasts between 1st and 2nd instance of drawing together for all sessions combined: contrasts between drawing instance and drawing instance* group interaction per ROI pair in unstandardised units. Tog1 = 1st instance of drawing together, Tog2 = 2nd instance of drawing together. Int–Same 1 = group contrast for 1st instance of drawing together, Int–Same 2 = group contrast for 2nd instance of drawing together.

| Contrast | ROI Pair | Estimate | Lower HPD | Upper HPD | No overlap |
|---|---|---|---|---|---|
| Tog1-Tog2 | LIFG~RIFG | 0.006 | -0.005 | 0.017 | |
| Int1-2 | LIFG~RIFG | 0.000 | -0.008 | 0.008 | |
| Same1-2 | LIFG~RIFG | 0.006 | -0.002 | 0.013 | |
| Int-Same 1 | LIFG~RIFG | -0.003 | -0.012 | 0.006 | |
| Int-Same 2 | LIFG~RIFG | 0.003 | -0.006 | 0.012 | |
| Tog1-Tog2 | LIFG~LIFG | -0.001 | -0.015 | 0.014 | |
| Int1-2 | LIFG~LIFG | 0.001 | -0.009 | 0.011 | |
| Same1-2 | LIFG~LIFG | -0.002 | -0.012 | 0.008 | |
| Int-Same 1 | LIFG~LIFG | 0.005 | -0.006 | 0.015 | |
| Int-Same 2 | LIFG~LIFG | 0.001 | -0.009 | 0.012 | |
| Tog1-Tog2 | LTPJ~LTPJ | 0.004 | -0.012 | 0.019 | |
| Int1-2 | LTPJ~LTPJ | 0.001 | -0.010 | 0.012 | |
| Same1-2 | LTPJ~LTPJ | 0.003 | -0.007 | 0.014 | |
| Int-Same 1 | LTPJ~LTPJ | 0.003 | -0.009 | 0.015 | |
| Int-Same 2 | LTPJ~LTPJ | 0.005 | -0.007 | 0.017 | |
| Tog1-Tog2 | LIFG~LTPJ | -0.003 | -0.013 | 0.007 | |
| Int1-2 | LIFG~LTPJ | 0.000 | -0.007 | 0.007 | |
| Same1-2 | LIFG~LTPJ | -0.003 | -0.010 | 0.004 | |
| Int-Same 1 | LIFG~LTPJ | 0.007 | -0.002 | 0.015 | |
| Int-Same 2 | LIFG~LTPJ | 0.004 | -0.005 | 0.013 | |
| Tog1-Tog2 | RIFG~RIFG | 0.002 | -0.014 | 0.018 | |

**Supplementary Materials:** Moffat, Dumas and Cross (2026). *Social interactions between people of same and different generations shape longitudinal changes in interpersonal neural synchrony, loneliness, and social connection.*

**S5 Appendix – Table H.** HbO. Change in INS across sessions: contrasts between tasks, groups and the task*group interaction per ROI pair in unstandardised units.

| Contrast | ROI Pair | Estimate | Lower HPD | Upper HPD | No overlap |
|---|---|---|---|---|---|
| Tog-Alo | LTPJ~LTPJ | 0.006 | -0.002 | 0.014 | |
| Int-Sam | LTPJ~LTPJ | 0.003 | -0.005 | 0.01 | |
| Tog-Alo Int | LTPJ~LTPJ | 0.006 | 0 | 0.011 | No overlap |
| Tog-Alo Sam | LTPJ~LTPJ | 0 | -0.006 | 0.005 | |
| Int-Sam Alo | LTPJ~LTPJ | -0.002 | -0.007 | 0.004 | |
| Int-Sam Tog | LTPJ~LTPJ | 0.004 | -0.001 | 0.01 | |
| Tog-Alo | LIFG~LTPJ | 0 | -0.005 | 0.005 | |
| Int-Sam | LIFG~LTPJ | -0.006 | -0.012 | -0.001 | No overlap |
| Tog-Alo Int | LIFG~LTPJ | -0.002 | -0.006 | 0.002 | |
| Tog-Alo Sam | LIFG~LTPJ | 0.002 | -0.002 | 0.006 | |
| Int-Sam Alo | LIFG~LTPJ | -0.001 | -0.005 | 0.003 | |
| Int-Sam Tog | LIFG~LTPJ | -0.005 | -0.009 | -0.001 | No overlap |
| Tog-Alo | LIFG~RTPJ | -0.002 | -0.008 | 0.005 | |
| Int-Sam | LIFG~RTPJ | 0.002 | -0.005 | 0.009 | |
| Tog-Alo Int | LIFG~RTPJ | -0.006 | -0.011 | -0.001 | No overlap |
| Tog-Alo Sam | LIFG~RTPJ | 0.004 | 0 | 0.009 | |
| Int-Sam Alo | LIFG~RTPJ | 0.006 | 0.001 | 0.011 | No overlap |
| Int-Sam Tog | LIFG~RTPJ | -0.004 | -0.009 | 0 | |
| Tog-Alo | RTPJ~RTPJ | -0.004 | -0.019 | 0.009 | |
| Int-Sam | RTPJ~RTPJ | -0.011 | -0.026 | 0.003 | |
| Tog-Alo Int | RTPJ~RTPJ | -0.011 | -0.021 | 0 | No overlap |
| Tog-Alo Sam | RTPJ~RTPJ | 0.006 | -0.003 | 0.016 | |
| Int-Sam Alo | RTPJ~RTPJ | 0.003 | -0.007 | 0.013 | |
| Int-Sam Tog | RTPJ~RTPJ | -0.014 | -0.024 | -0.004 | No overlap |
| Tog-Alo | RIFG~RTPJ | -0.002 | -0.009 | 0.005 | |
| Int-Sam | RIFG~RTPJ | -0.001 | -0.008 | 0.007 | |
| Tog-Alo Int | RIFG~RTPJ | -0.007 | -0.012 | -0.002 | No overlap |
| Tog-Alo Sam | RIFG~RTPJ | 0.004 | 0 | 0.009 | |
| Int-Sam Alo | RIFG~RTPJ | 0.005 | 0 | 0.011 | No overlap |
| Int-Sam Tog | RIFG~RTPJ | -0.006 | -0.011 | -0.001 | No overlap |

**Supplementary Materials:** Moffat, Dumas and Cross (2026). *Social interactions between people of same and different generations shape longitudinal changes in interpersonal neural synchrony, loneliness, and social connection.*

**S5 Appendix – Table I.** HbO. Relationships between session and INS in unstandardised units. Effects are presented in the following order: Main, group, task, and simple effects.

| Measure | Group | Task | ROI Pairs | Estimate | Lower HPD | Upper HPD | No overlap |
|---|---|---|---|---|---|---|---|
| *Main effect* | | | | | | | |
| session | | | LIFG~RIFG | -0.0004 | -0.0024 | 0.0016 | |
| session | | | LIFG~LIFG | -0.0002 | -0.0025 | 0.0019 | |
| session | | | LTPJ~LTPJ | 0.0004 | -0.0011 | 0.0020 | |
| session | | | LIFG~LTPJ | 0.0002 | -0.0017 | 0.0021 | |
| session | | | LIFG~RTPJ | 0.0008 | -0.0015 | 0.0030 | |
| session | | | RIFG~RIFG | 0.0046 | 0.0002 | 0.0091 | |
| session | | | RTPJ~RTPJ | -0.0003 | -0.0018 | 0.0012 | No overlap |
| session | | | RIFG~LTPJ | -0.0012 | -0.0032 | 0.0008 | |
| session | | | RIFG~RTPJ | 0.0003 | -0.0010 | 0.0016 | |
| *Group effect* | | | | | | | |
| session | Intergen | | LIFG~RIFG | 0.0002 | -0.0020 | 0.0024 | |
| session | Samegen | | LIFG~RIFG | -0.0007 | -0.0030 | 0.0017 | |
| session | Intergen | | LIFG~LIFG | -0.0005 | -0.0033 | 0.0023 | |
| session | Samegen | | LIFG~LIFG | -0.0003 | -0.0034 | 0.0026 | |
| session | Intergen | | LTPJ~LTPJ | 0.0009 | -0.0022 | 0.0039 | |
| session | Samegen | | LTPJ~LTPJ | -0.0014 | -0.0046 | 0.0019 | |
| session | Intergen | | LIFG~LTPJ | -0.0012 | -0.0033 | 0.0009 | |
| session | Samegen | | LIFG~LTPJ | 0.0021 | -0.0001 | 0.0044 | |
| session | Intergen | | LIFG~RTPJ | 0.0010 | -0.0016 | 0.0036 | |
| session | Samegen | | LIFG~RTPJ | -0.0006 | -0.0033 | 0.0020 | |
| session | Intergen | | RIFG~RIFG | 0.0012 | -0.0020 | 0.0044 | |
| session | Samegen | | RIFG~RIFG | 0.0004 | -0.0028 | 0.0036 | |
| session | Intergen | | RTPJ~RTPJ | 0.0015 | -0.0047 | 0.0078 | |
| session | Samegen | | RTPJ~RTPJ | 0.0077 | 0.0013 | 0.0138 | No overlap |
| session | Intergen | | RIFG~LTPJ | 0.0003 | -0.0018 | 0.0024 | |
| session | Samegen | | RIFG~LTPJ | -0.0009 | -0.0030 | 0.0013 | |
| session | Intergen | | RIFG~RTPJ | -0.0003 | -0.0030 | 0.0026 | |
| session | Samegen | | RIFG~RTPJ | -0.0020 | -0.0049 | 0.0008 | |
| *Task effect* | | | | | | | |
| session | | Alone | LIFG~RIFG | -0.0017 | -0.0040 | 0.0006 | |
| session | | Together | LIFG~RIFG | 0.0012 | -0.0010 | 0.0032 | |
| session | | Alone | LIFG~LIFG | -0.0008 | -0.0038 | 0.0023 | |
| session | | Together | LIFG~LIFG | 0.0000 | -0.0028 | 0.0027 | |
| session | | Alone | LTPJ~LTPJ | -0.0014 | -0.0046 | 0.0019 | |
| session | | Together | LTPJ~LTPJ | 0.0009 | -0.0020 | 0.0039 | |
| session | | Alone | LIFG~LTPJ | 0.0009 | -0.0015 | 0.0031 | |
| session | | Together | LIFG~LTPJ | 0.0000 | -0.0020 | 0.0021 | |
| session | | Alone | LIFG~RTPJ | -0.0001 | -0.0028 | 0.0027 | |
| session | | Together | LIFG~RTPJ | 0.0005 | -0.0019 | 0.0030 | |
| session | | Alone | RIFG~RIFG | 0.0012 | -0.0020 | 0.0046 | |
| session | | Together | RIFG~RIFG | 0.0004 | -0.0027 | 0.0034 | |
| session | | Alone | RTPJ~RTPJ | 0.0040 | -0.0020 | 0.0105 | |

| Measure | Group | Task | ROI Pairs | Estimate | Lower HPD | Upper HPD | No overlap |
|---|---|---|---|---|---|---|---|
| session | | Together | RTPJ~RTPJ | 0.0051 | -0.0008 | 0.0111 | |
| session | | Alone | RIFG~LTPJ | 0.0013 | -0.0010 | 0.0035 | |
| session | | Together | RIFG~LTPJ | -0.0019 | -0.0039 | 0.0002 | |
| session | | Alone | RIFG~RTPJ | -0.0016 | -0.0045 | 0.0013 | |
| session | | Together | RIFG~RTPJ | -0.0007 | -0.0034 | 0.0019 | |
| *Simple effect* | | | | | | | |
| session | Intergen | Alone | LIFG~RIFG | -0.0007 | -0.0040 | 0.0026 | |
| session | Intergen | Together | LIFG~RIFG | 0.0011 | -0.0018 | 0.0040 | |
| session | Samegen | Alone | LIFG~RIFG | -0.0026 | -0.0060 | 0.0008 | |
| session | Samegen | Together | LIFG~RIFG | 0.0012 | -0.0019 | 0.0042 | |
| session | Intergen | Alone | LIFG~LIFG | -0.0003 | -0.0046 | 0.0037 | |
| session | Intergen | Together | LIFG~LIFG | -0.0007 | -0.0043 | 0.0032 | |
| session | Samegen | Alone | LIFG~LIFG | -0.0013 | -0.0058 | 0.0032 | |
| session | Samegen | Together | LIFG~LIFG | 0.0006 | -0.0033 | 0.0046 | |
| session | Intergen | Alone | LTPJ~LTPJ | -0.0016 | -0.0060 | 0.0028 | |
| session | Intergen | Together | LTPJ~LTPJ | 0.0034 | -0.0006 | 0.0075 | |
| session | Samegen | Alone | LTPJ~LTPJ | -0.0012 | -0.0058 | 0.0037 | |
| session | Samegen | Together | LTPJ~LTPJ | -0.0015 | -0.0056 | 0.0029 | |
| session | Intergen | Alone | LIFG~LTPJ | 0.0005 | -0.0025 | 0.0036 | |
| session | Intergen | Together | LIFG~LTPJ | -0.0029 | -0.0058 | -0.0001 | No overlap |
| session | Samegen | Alone | LIFG~LTPJ | 0.0012 | -0.0021 | 0.0045 | |
| session | Samegen | Together | LIFG~LTPJ | 0.0029 | 0.0000 | 0.0059 | |
| session | Intergen | Alone | LIFG~RTPJ | 0.0034 | -0.0003 | 0.0072 | |
| session | Intergen | Together | LIFG~RTPJ | -0.0015 | -0.0049 | 0.0020 | |
| session | Samegen | Alone | LIFG~RTPJ | -0.0037 | -0.0078 | 0.0002 | |
| session | Samegen | Together | LIFG~RTPJ | 0.0024 | -0.0010 | 0.0060 | |
| session | Intergen | Alone | RIFG~RIFG | 0.0004 | -0.0044 | 0.0051 | |
| session | Intergen | Together | RIFG~RIFG | 0.0021 | -0.0023 | 0.0062 | |
| session | Samegen | Alone | RIFG~RIFG | 0.0021 | -0.0025 | 0.0070 | |
| session | Samegen | Together | RIFG~RIFG | -0.0014 | -0.0056 | 0.0029 | |
| session | Intergen | Alone | RTPJ~RTPJ | 0.0061 | -0.0024 | 0.0148 | |
| session | Intergen | Together | RTPJ~RTPJ | -0.0032 | -0.0119 | 0.0053 | |
| session | Samegen | Alone | RTPJ~RTPJ | 0.0021 | -0.0074 | 0.0111 | |
| session | Samegen | Together | RTPJ~RTPJ | 0.0134 | 0.0052 | 0.0217 | No overlap |
| session | Intergen | Alone | RIFG~LTPJ | 0.0020 | -0.0012 | 0.0051 | |
| session | Intergen | Together | RIFG~LTPJ | -0.0015 | -0.0043 | 0.0013 | |
| session | Samegen | Alone | RIFG~LTPJ | 0.0005 | -0.0028 | 0.0037 | |
| session | Samegen | Together | RIFG~LTPJ | -0.0022 | -0.0052 | 0.0007 | |
| session | Intergen | Alone | RIFG~RTPJ | 0.0025 | -0.0013 | 0.0067 | |
| session | Intergen | Together | RIFG~RTPJ | -0.0031 | -0.0069 | 0.0007 | |
| session | Samegen | Alone | RIFG~RTPJ | -0.0058 | -0.0100 | -0.0016 | No overlap |
| session | Samegen | Together | RIFG~RTPJ | 0.0016 | -0.0020 | 0.0054 | |

**S5 Appendix – Table J.** HbO. Contrasts pertaining to session ~ INS relationship in unstandardised units. Contrasts are presented in the following order: Group, task, and interaction contrasts.

| Measure | Group | Task | ROI Pairs | Estimate | Lower HPD | Upper HPD | No overlap |
|---|---|---|---|---|---|---|---|
| *Group contrast* | | | | | | | |
| session | Intergen-Samegen | | LIFG~RIFG | 0.0009 | -0.0023 | 0.0041 | |
| session | Intergen-Samegen | | LIFG~LIFG | -0.0002 | -0.0043 | 0.0039 | |
| session | Intergen-Samegen | | LTPJ~LTPJ | 0.0022 | -0.0023 | 0.0067 | |
| session | Intergen-Samegen | | LIFG~LTPJ | -0.0033 | -0.0064 | -0.0002 | No overlap |
| session | Intergen-Samegen | | LIFG~RTPJ | 0.0016 | -0.0021 | 0.0053 | |
| session | Intergen-Samegen | | RIFG~RIFG | 0.0008 | -0.0037 | 0.0054 | |
| session | Intergen-Samegen | | RTPJ~RTPJ | -0.0062 | -0.0150 | 0.0027 | |
| session | Intergen-Samegen | | RIFG~LTPJ | 0.0011 | -0.0020 | 0.0041 | |
| session | Intergen-Samegen | | RIFG~RTPJ | 0.0018 | -0.0021 | 0.0059 | |
| *Task contrast* | | | | | | | |
| session | | Alone-Together | LIFG~RIFG | -0.0028 | -0.0060 | 0.0002 | |
| session | | Alone-Together | LIFG~LIFG | -0.0007 | -0.0049 | 0.0032 | |
| session | | Alone-Together | LTPJ~LTPJ | -0.0023 | -0.0066 | 0.0019 | |
| session | | Alone-Together | LIFG~LTPJ | 0.0008 | -0.0023 | 0.0038 | |
| session | | Alone-Together | LIFG~RTPJ | -0.0006 | -0.0044 | 0.0030 | |
| session | | Alone-Together | RIFG~RIFG | 0.0009 | -0.0036 | 0.0054 | |
| session | | Alone-Together | RTPJ~RTPJ | -0.0010 | -0.0099 | 0.0072 | |
| session | | Alone-Together | RIFG~LTPJ | 0.0032 | 0.0002 | 0.0062 | No overlap |
| session | | Alone-Together | RIFG~RTPJ | -0.0009 | -0.0048 | 0.0029 | |
| *Interactions* | | | | | | | |
| session | Intergen-Samegen | Alone | LIFG~RIFG | 0.0019 | -0.0029 | 0.0065 | |
| session | Intergen-Samegen | Together | LIFG~RIFG | 0 | -0.0044 | 0.0041 | |
| session | Intergen-Samegen | Alone | LIFG~LIFG | 0.001 | -0.0052 | 0.0069 | |
| session | Intergen-Samegen | Together | LIFG~LIFG | -0.0012 | -0.0065 | 0.0043 | |
| session | Intergen-Samegen | Alone | LTPJ~LTPJ | -5.00E-04 | -0.0068 | 0.0063 | |
| session | Intergen-Samegen | Together | LTPJ~LTPJ | 0.0049 | -0.001 | 0.0107 | |
| session | Intergen-Samegen | Alone | LIFG~LTPJ | -7.00E-04 | -0.0051 | 0.0039 | |
| session | Intergen-Samegen | Together | LIFG~LTPJ | -0.0058 | -0.0099 | -0.0017 | No overlap |
| session | Intergen-Samegen | Alone | LIFG~RTPJ | 0.0072 | 0.0018 | 0.0127 | No overlap |
| session | Intergen-Samegen | Together | LIFG~RTPJ | -0.004 | -0.0091 | 7.00E-04 | |
| session | Intergen-Samegen | Alone | RIFG~RIFG | -0.0017 | -0.0082 | 0.0052 | |
| session | Intergen-Samegen | Together | RIFG~RIFG | 0.0034 | -0.0027 | 0.0094 | |
| session | Intergen-Samegen | Alone | RTPJ~RTPJ | 0.0042 | -0.0087 | 0.0167 | |
| session | Intergen-Samegen | Together | RTPJ~RTPJ | -0.0165 | -0.0285 | -0.0048 | No overlap |
| session | Intergen-Samegen | Alone | RIFG~LTPJ | 0.0015 | -0.0031 | 0.006 | |
| session | Intergen-Samegen | Together | RIFG~LTPJ | 7.00E-04 | -0.0033 | 0.0049 | |
| session | Intergen-Samegen | Alone | RIFG~RTPJ | 0.0083 | 0.0024 | 0.0141 | No overlap |
| session | Intergen-Samegen | Together | RIFG~RTPJ | -0.0047 | -0.0101 | 6.00E-04 | |
| session | Intergen | Alone-Together | LIFG~RIFG | -0.0019 | -0.0062 | 0.0025 | |
| session | Samegen | Alone-Together | LIFG~RIFG | -0.0038 | -0.0082 | 0.0007 | |
| session | Intergen | Alone-Together | LIFG~LIFG | 0.0004 | -0.0054 | 0.0058 | |
| session | Samegen | Alone-Together | LIFG~LIFG | -0.0018 | -0.0078 | 0.0041 | |

| Measure | Group | Task | ROI Pairs | Estimate | Lower HPD | Upper HPD | No overlap |
|---|---|---|---|---|---|---|---|
| session | Intergen | Alone-Together | LTPJ~LTPJ | -0.0050 | -0.0111 | 0.0007 | |
| session | Samegen | Alone-Together | LTPJ~LTPJ | 0.0003 | -0.0059 | 0.0065 | |
| session | Intergen | Alone-Together | LIFG~LTPJ | 0.0034 | -0.0008 | 0.0074 | |
| session | Samegen | Alone-Together | LIFG~LTPJ | -0.0017 | -0.0061 | 0.0027 | |
| session | Intergen | Alone-Together | LIFG~RTPJ | 0.0049 | -0.0002 | 0.0099 | |
| session | Samegen | Alone-Together | LIFG~RTPJ | -0.0062 | -0.0116 | -0.0011 | No overlap |
| session | Intergen | Alone-Together | RIFG~RIFG | -0.0017 | -0.0080 | 0.0046 | |
| session | Samegen | Alone-Together | RIFG~RIFG | 0.0035 | -0.0029 | 0.0098 | |
| session | Intergen | Alone-Together | RTPJ~RTPJ | 0.0093 | -0.0028 | 0.0210 | |
| session | Samegen | Alone-Together | RTPJ~RTPJ | -0.0113 | -0.0232 | 0.0011 | |
| session | Intergen | Alone-Together | RIFG~LTPJ | 0.0035 | -0.0006 | 0.0078 | |
| session | Samegen | Alone-Together | RIFG~LTPJ | 0.0028 | -0.0016 | 0.0071 | |
| session | Intergen | Alone-Together | RIFG~RTPJ | 0.0056 | 0.0003 | 0.0111 | No overlap |
| session | Samegen | Alone-Together | RIFG~RTPJ | -0.0074 | -0.0131 | -0.0020 | No overlap |

**S5 Appendix – Table K.** HbO. Relationships between loneliness (sum) and INS in unstandardised units. Effects are presented in the following order: Main, group, task, and simple effects.

| Measure | Group | Task | ROI Pairs | Estimate | Lower HPD | Upper HPD | No overlap |
|---|---|---|---|---|---|---|---|
| *Main effect* | | | | | | | |
| sum loneliness | | | LIFG~RIFG | 0.0003 | -0.001 | 0.0016 | |
| sum loneliness | | | LIFG~LIFG | 0.0002 | -0.0015 | 0.0018 | |
| sum loneliness | | | LTPJ~LTPJ | 0.0004 | -0.0016 | 0.0024 | |
| sum loneliness | | | LIFG~LTPJ | 0.0004 | -0.0009 | 0.0017 | |
| sum loneliness | | | LIFG~RTPJ | -0.0012 | -0.0027 | 0.0003 | |
| sum loneliness | | | RIFG~RIFG | -0.0003 | -0.0021 | 0.0015 | |
| sum loneliness | | | RTPJ~RTPJ | -0.0009 | -0.0041 | 0.0021 | |
| sum loneliness | | | RIFG~LTPJ | -0.0001 | -0.0013 | 0.0011 | |
| sum loneliness | | | RIFG~RTPJ | -0.0006 | -0.0023 | 0.0011 | |
| *Group effect* | | | | | | | |
| sum loneliness | Intergen | | LIFG~RIFG | -0.0005 | -0.0023 | 0.0013 | |
| sum loneliness | Samegen | | LIFG~RIFG | 0.001 | -0.0008 | 0.0029 | |
| sum loneliness | Intergen | | LIFG~LIFG | 0.0001 | -0.0022 | 0.0023 | |
| sum loneliness | Samegen | | LIFG~LIFG | 0.0004 | -0.0019 | 0.0027 | |
| sum loneliness | Intergen | | LTPJ~LTPJ | 0.0007 | -0.002 | 0.0034 | |
| sum loneliness | Samegen | | LTPJ~LTPJ | 0.0001 | -0.0027 | 0.003 | |
| sum loneliness | Intergen | | LIFG~LTPJ | 0.0002 | -0.0016 | 0.002 | |
| sum loneliness | Samegen | | LIFG~LTPJ | 0.0005 | -0.0014 | 0.0024 | |
| sum loneliness | Intergen | | LIFG~RTPJ | -0.0012 | -0.0032 | 0.0009 | |
| sum loneliness | Samegen | | LIFG~RTPJ | -0.0011 | -0.0033 | 0.001 | |
| sum loneliness | Intergen | | RIFG~RIFG | -0.0003 | -0.0028 | 0.0022 | |
| sum loneliness | Samegen | | RIFG~RIFG | -0.0004 | -0.0029 | 0.0022 | |
| sum loneliness | Intergen | | RTPJ~RTPJ | -0.0004 | -0.0045 | 0.0037 | |
| sum loneliness | Samegen | | RTPJ~RTPJ | -0.0014 | -0.0061 | 0.0033 | |
| sum loneliness | Intergen | | RIFG~LTPJ | 0 | -0.0016 | 0.0017 | |
| sum loneliness | Samegen | | RIFG~LTPJ | -0.0003 | -0.0021 | 0.0014 | |
| sum loneliness | Intergen | | RIFG~RTPJ | -0.0017 | -0.004 | 0.0007 | |
| sum loneliness | Samegen | | RIFG~RTPJ | 0.0005 | -0.0018 | 0.0029 | |
| *Task effect* | | | | | | | |
| sum loneliness | | Alone | LIFG~RIFG | -0.0001 | -0.002 | 0.0018 | |
| sum loneliness | | Together | LIFG~RIFG | 0.0006 | -0.0009 | 0.0022 | |
| sum loneliness | | Alone | LIFG~LIFG | 0.0004 | -0.002 | 0.0029 | |
| sum loneliness | | Together | LIFG~LIFG | 0 | -0.0019 | 0.002 | |
| sum loneliness | | Alone | LTPJ~LTPJ | 0.0016 | -0.0013 | 0.0043 | |
| sum loneliness | | Together | LTPJ~LTPJ | -0.0008 | -0.003 | 0.0015 | |
| sum loneliness | | Alone | LIFG~LTPJ | 0.0007 | -0.0012 | 0.0026 | |
| sum loneliness | | Together | LIFG~LTPJ | 0 | -0.0015 | 0.0016 | |
| sum loneliness | | Alone | LIFG~RTPJ | -0.0017 | -0.0039 | 0.0005 | |
| sum loneliness | | Together | LIFG~RTPJ | -0.0006 | -0.0024 | 0.0011 | |
| sum loneliness | | Alone | RIFG~RIFG | -0.0017 | -0.0044 | 0.001 | |
| sum loneliness | | Together | RIFG~RIFG | 0.0011 | -0.0011 | 0.0032 | |
| sum loneliness | | Alone | RTPJ~RTPJ | -0.0013 | -0.0059 | 0.0031 | |

| Measure | Group | Task | ROI Pairs | Estimate | Lower HPD | Upper HPD | No overlap |
|---|---|---|---|---|---|---|---|
| sum loneliness | | Together | RTPJ~RTPJ | -0.0005 | -0.0043 | 0.0034 | |
| sum loneliness | | Alone | RIFG~LTPJ | -0.0003 | -0.0021 | 0.0015 | |
| sum loneliness | | Together | RIFG~LTPJ | 0.0001 | -0.0014 | 0.0015 | |
| sum loneliness | | Alone | RIFG~RTPJ | -0.0016 | -0.004 | 0.0008 | |
| sum loneliness | | Together | RIFG~RTPJ | 0.0005 | -0.0015 | 0.0024 | |
| *Simple effect* | | | | | | | |
| sum loneliness | Intergen | Alone | LIFG~RIFG | -0.0014 | -0.0040 | 0.0013 | |
| sum loneliness | Intergen | Together | LIFG~RIFG | 0.0004 | -0.0018 | 0.0024 | |
| sum loneliness | Samegen | Alone | LIFG~RIFG | 0.0012 | -0.0016 | 0.0038 | |
| sum loneliness | Samegen | Together | LIFG~RIFG | 0.0009 | -0.0012 | 0.0031 | |
| sum loneliness | Intergen | Alone | LIFG~LIFG | 0.0017 | -0.0017 | 0.0051 | |
| sum loneliness | Intergen | Together | LIFG~LIFG | -0.0016 | -0.0043 | 0.0010 | |
| sum loneliness | Samegen | Alone | LIFG~LIFG | -0.0009 | -0.0044 | 0.0027 | |
| sum loneliness | Samegen | Together | LIFG~LIFG | 0.0016 | -0.0012 | 0.0044 | |
| sum loneliness | Intergen | Alone | LTPJ~LTPJ | 0.0020 | -0.0018 | 0.0058 | |
| sum loneliness | Intergen | Together | LTPJ~LTPJ | -0.0005 | -0.0036 | 0.0026 | |
| sum loneliness | Samegen | Alone | LTPJ~LTPJ | 0.0012 | -0.0028 | 0.0052 | |
| sum loneliness | Samegen | Together | LTPJ~LTPJ | -0.0010 | -0.0043 | 0.0023 | |
| sum loneliness | Intergen | Alone | LIFG~LTPJ | 0.0001 | -0.0025 | 0.0027 | |
| sum loneliness | Intergen | Together | LIFG~LTPJ | 0.0004 | -0.0016 | 0.0026 | |
| sum loneliness | Samegen | Alone | LIFG~LTPJ | 0.0014 | -0.0014 | 0.0041 | |
| sum loneliness | Samegen | Together | LIFG~LTPJ | -0.0003 | -0.0026 | 0.0019 | |
| sum loneliness | Intergen | Alone | LIFG~RTPJ | -0.0024 | -0.0054 | 0.0006 | |
| sum loneliness | Intergen | Together | LIFG~RTPJ | 0.0000 | -0.0024 | 0.0025 | |
| sum loneliness | Samegen | Alone | LIFG~RTPJ | -0.0010 | -0.0042 | 0.0021 | |
| sum loneliness | Samegen | Together | LIFG~RTPJ | -0.0013 | -0.0038 | 0.0013 | |
| sum loneliness | Intergen | Alone | RIFG~RIFG | -0.0009 | -0.0047 | 0.0028 | |
| sum loneliness | Intergen | Together | RIFG~RIFG | 0.0003 | -0.0027 | 0.0033 | |
| sum loneliness | Samegen | Alone | RIFG~RIFG | -0.0026 | -0.0064 | 0.0012 | |
| sum loneliness | Samegen | Together | RIFG~RIFG | 0.0019 | -0.0012 | 0.0050 | |
| sum loneliness | Intergen | Alone | RTPJ~RTPJ | -0.0005 | -0.0065 | 0.0053 | |
| sum loneliness | Intergen | Together | RTPJ~RTPJ | -0.0002 | -0.0054 | 0.0048 | |
| sum loneliness | Samegen | Alone | RTPJ~RTPJ | -0.0021 | -0.0089 | 0.0048 | |
| sum loneliness | Samegen | Together | RTPJ~RTPJ | -0.0007 | -0.0066 | 0.0051 | |
| sum loneliness | Intergen | Alone | RIFG~LTPJ | -0.0009 | -0.0035 | 0.0016 | |
| sum loneliness | Intergen | Together | RIFG~LTPJ | 0.0009 | -0.0011 | 0.0029 | |
| sum loneliness | Samegen | Alone | RIFG~LTPJ | 0.0002 | -0.0023 | 0.0029 | |
| sum loneliness | Samegen | Together | RIFG~LTPJ | -0.0008 | -0.0029 | 0.0012 | |
| sum loneliness | Intergen | Alone | RIFG~RTPJ | -0.0036 | -0.0069 | -0.0003 | No overlap |
| sum loneliness | Intergen | Together | RIFG~RTPJ | 0.0002 | -0.0026 | 0.0028 | |
| sum loneliness | Samegen | Alone | RIFG~RTPJ | 0.0003 | -0.0032 | 0.0037 | |
| sum loneliness | Samegen | Together | RIFG~RTPJ | 0.0007 | -0.0019 | 0.0036 | |
| sum loneliness | Intergen | Alone | LIFG~RIFG | -0.0014 | -0.0040 | 0.0013 | |

**S5 Appendix – Table L.** HbO. Contrasts pertaining to loneliness (sum) ~ INS relationship in unstandardised units. Contrasts are presented in the following order: Group, task, and interaction contrasts.

| Measure | Group | Task | ROI Pairs | Estimate | Lower HPD | Upper HPD | No overlap |
|---|---|---|---|---|---|---|---|
| *Group contrast* | | | | | | | |
| sum loneliness | Intergen-Samegen | | LIFG~RIFG | -0.0015 | -0.0041 | 0.0010 | |
| sum loneliness | Intergen-Samegen | | LIFG~LIFG | -0.0003 | -0.0036 | 0.0029 | |
| sum loneliness | Intergen-Samegen | | LTPJ~LTPJ | 0.0007 | -0.0032 | 0.0047 | |
| sum loneliness | Intergen-Samegen | | LIFG~LTPJ | -0.0003 | -0.0030 | 0.0023 | |
| sum loneliness | Intergen-Samegen | | LIFG~RTPJ | -0.0001 | -0.0029 | 0.0029 | |
| sum loneliness | Intergen-Samegen | | RIFG~RIFG | 0.0001 | -0.0035 | 0.0036 | |
| sum loneliness | Intergen-Samegen | | RTPJ~RTPJ | 0.0010 | -0.0051 | 0.0073 | |
| sum loneliness | Intergen-Samegen | | RIFG~LTPJ | 0.0003 | -0.0021 | 0.0027 | |
| sum loneliness | Intergen-Samegen | | RIFG~RTPJ | -0.0022 | -0.0055 | 0.0010 | |
| *Task contrast* | | | | | | | |
| sum loneliness | | Alone-Together | LIFG~RIFG | -0.0007 | -0.0030 | 0.0016 | |
| sum loneliness | | Alone-Together | LIFG~LIFG | 0.0004 | -0.0026 | 0.0034 | |
| sum loneliness | | Alone-Together | LTPJ~LTPJ | 0.0023 | -0.0009 | 0.0055 | |
| sum loneliness | | Alone-Together | LIFG~LTPJ | 0.0007 | -0.0015 | 0.0029 | |
| sum loneliness | | Alone-Together | LIFG~RTPJ | -0.0011 | -0.0037 | 0.0015 | |
| sum loneliness | | Alone-Together | RIFG~RIFG | -0.0028 | -0.0062 | 0.0004 | |
| sum loneliness | | Alone-Together | RTPJ~RTPJ | -0.0009 | -0.0065 | 0.0050 | |
| sum loneliness | | Alone-Together | RIFG~LTPJ | -0.0004 | -0.0026 | 0.0019 | |
| sum loneliness | | Alone-Together | RIFG~RTPJ | -0.0021 | -0.0049 | 0.0007 | |
| *Interactions* | | | | | | | |
| sum loneliness | Intergen-Samegen | Alone | LIFG~RIFG | -0.0025 | -0.0062 | 0.0014 | |
| sum loneliness | Intergen-Samegen | Together | LIFG~RIFG | -0.0005 | -0.0035 | 0.0026 | |
| sum loneliness | Intergen-Samegen | Alone | LIFG~LIFG | 0.0026 | -0.0023 | 0.0075 | |
| sum loneliness | Intergen-Samegen | Together | LIFG~LIFG | -0.0032 | -0.0070 | 0.0007 | |
| sum loneliness | Intergen-Samegen | Alone | LTPJ~LTPJ | 0.0008 | -0.0047 | 0.0063 | |
| sum loneliness | Intergen-Samegen | Together | LTPJ~LTPJ | 0.0005 | -0.0040 | 0.0051 | |
| sum loneliness | Intergen-Samegen | Alone | LIFG~LTPJ | -0.0013 | -0.0050 | 0.0025 | |
| sum loneliness | Intergen-Samegen | Together | LIFG~LTPJ | 0.0007 | -0.0024 | 0.0037 | |
| sum loneliness | Intergen-Samegen | Alone | LIFG~RTPJ | -0.0014 | -0.0057 | 0.0029 | |
| sum loneliness | Intergen-Samegen | Together | LIFG~RTPJ | 0.0013 | -0.0021 | 0.0048 | |
| sum loneliness | Intergen-Samegen | Alone | RIFG~RIFG | 0.0017 | -0.0036 | 0.0071 | |
| sum loneliness | Intergen-Samegen | Together | RIFG~RIFG | -0.0016 | -0.0060 | 0.0025 | |
| sum loneliness | Intergen-Samegen | Alone | RTPJ~RTPJ | 0.0016 | -0.0074 | 0.0107 | |
| sum loneliness | Intergen-Samegen | Together | RTPJ~RTPJ | 0.0004 | -0.0073 | 0.0083 | |
| sum loneliness | Intergen-Samegen | Alone | RIFG~LTPJ | -0.0012 | -0.0048 | 0.0024 | |
| sum loneliness | Intergen-Samegen | Together | RIFG~LTPJ | 0.0017 | -0.0011 | 0.0046 | |
| sum loneliness | Intergen-Samegen | Alone | RIFG~RTPJ | -0.0039 | -0.0086 | 0.0008 | |
| sum loneliness | Intergen-Samegen | Together | RIFG~RTPJ | -0.0005 | -0.0043 | 0.0034 | |
| sum loneliness | Intergen | Alone-Together | LIFG~RIFG | -0.0017 | -0.0049 | 0.0015 | |
| sum loneliness | Samegen | Alone-Together | LIFG~RIFG | 0.0002 | -0.0031 | 0.0035 | |
| sum loneliness | Intergen | Alone-Together | LIFG~LIFG | 0.0033 | -0.0007 | 0.0076 | |
| sum loneliness | Samegen | Alone-Together | LIFG~LIFG | -0.0025 | -0.0068 | 0.0019 | |

| Measure | Group | Task | ROI Pairs | Estimate | Lower HPD | Upper HPD | No overlap |
|---|---|---|---|---|---|---|---|
| sum loneliness | Intergen | Alone-Together | LTPJ~LTPJ | 0.0025 | -0.0019 | 0.0068 | |
| sum loneliness | Samegen | Alone-Together | LTPJ~LTPJ | 0.0022 | -0.0024 | 0.0068 | |
| sum loneliness | Intergen | Alone-Together | LIFG~LTPJ | -0.0003 | -0.0034 | 0.0028 | |
| sum loneliness | Samegen | Alone-Together | LIFG~LTPJ | 0.0017 | -0.0015 | 0.0050 | |
| sum loneliness | Intergen | Alone-Together | LIFG~RTPJ | -0.0024 | -0.0060 | 0.0013 | |
| sum loneliness | Samegen | Alone-Together | LIFG~RTPJ | 0.0003 | -0.0035 | 0.0041 | |
| sum loneliness | Intergen | Alone-Together | RIFG~RIFG | -0.0011 | -0.0058 | 0.0033 | |
| sum loneliness | Samegen | Alone-Together | RIFG~RIFG | -0.0045 | -0.0090 | 0.0002 | |
| sum loneliness | Intergen | Alone-Together | RTPJ~RTPJ | -0.0003 | -0.0078 | 0.0072 | |
| sum loneliness | Samegen | Alone-Together | RTPJ~RTPJ | -0.0014 | -0.0103 | 0.0072 | |
| sum loneliness | Intergen | Alone-Together | RIFG~LTPJ | -0.0018 | -0.0049 | 0.0013 | |
| sum loneliness | Samegen | Alone-Together | RIFG~LTPJ | 0.0011 | -0.0021 | 0.0042 | |
| sum loneliness | Intergen | Alone-Together | RIFG~RTPJ | -0.0038 | -0.0077 | 0.0001 | |
| sum loneliness | Samegen | Alone-Together | RIFG~RTPJ | -0.0004 | -0.0045 | 0.0036 | |

**Supplementary Materials:** Moffat, Dumas and Cross (2026). *Social interactions between people of same and different generations shape longitudinal changes in interpersonal neural synchrony, loneliness, and social connection.*

**S5 Appendix – Table M.** HbO. Relationships between loneliness (dif) and INS in unstandardised units. Effects are presented in the following order: Main, group, task, and simple effects.

| Measure | Group | Task | ROI Pairs | Estimate | Lower HPD | Upper HPD | No overlap |
|---|---|---|---|---|---|---|---|
| *Main effect* | | | | | | | |
| dif loneliness | | | LIFG~RIFG | -0.0001 | -0.0025 | 0.0022 | |
| dif loneliness | | | LIFG~LIFG | -0.0006 | -0.0035 | 0.0023 | |
| dif loneliness | | | LTPJ~LTPJ | -0.0016 | -0.005 | 0.0018 | |
| dif loneliness | | | LIFG~LTPJ | -0.0014 | -0.0037 | 0.0009 | |
| dif loneliness | | | LIFG~RTPJ | 0 | -0.0027 | 0.0027 | |
| dif loneliness | | | RIFG~RIFG | -0.0014 | -0.0047 | 0.002 | |
| dif loneliness | | | RTPJ~RTPJ | -0.0001 | -0.0057 | 0.0056 | |
| dif loneliness | | | RIFG~LTPJ | -0.0003 | -0.0025 | 0.0019 | |
| dif loneliness | | | RIFG~RTPJ | -0.0004 | -0.0034 | 0.0025 | |
| *Group effect* | | | | | | | |
| dif loneliness | Intergen | | LIFG~RIFG | 0.0001 | -0.0036 | 0.0038 | |
| dif loneliness | Samegen | | LIFG~RIFG | -0.0003 | -0.0031 | 0.0023 | |
| dif loneliness | Intergen | | LIFG~LIFG | -0.0008 | -0.0055 | 0.0037 | |
| dif loneliness | Samegen | | LIFG~LIFG | -0.0004 | -0.0037 | 0.0032 | |
| dif loneliness | Intergen | | LTPJ~LTPJ | -0.0047 | -0.0099 | 0.0006 | |
| dif loneliness | Samegen | | LTPJ~LTPJ | 0.0016 | -0.0027 | 0.0058 | |
| dif loneliness | Intergen | | LIFG~LTPJ | -0.003 | -0.0066 | 0.0006 | |
| dif loneliness | Samegen | | LIFG~LTPJ | 0.0001 | -0.0026 | 0.0029 | |
| dif loneliness | Intergen | | LIFG~RTPJ | 0.001 | -0.0033 | 0.0053 | |
| dif loneliness | Samegen | | LIFG~RTPJ | -0.0011 | -0.0043 | 0.002 | |
| dif loneliness | Intergen | | RIFG~RIFG | -0.0022 | -0.0077 | 0.0033 | |
| dif loneliness | Samegen | | RIFG~RIFG | -0.0005 | -0.0044 | 0.0032 | |
| dif loneliness | Intergen | | RTPJ~RTPJ | -0.003 | -0.0124 | 0.0066 | |
| dif loneliness | Samegen | | RTPJ~RTPJ | 0.0028 | -0.0037 | 0.0093 | |
| dif loneliness | Intergen | | RIFG~LTPJ | -0.001 | -0.0046 | 0.0025 | |
| dif loneliness | Samegen | | RIFG~LTPJ | 0.0005 | -0.002 | 0.003 | |
| dif loneliness | Intergen | | RIFG~RTPJ | -0.0019 | -0.0069 | 0.003 | |
| dif loneliness | Samegen | | RIFG~RTPJ | 0.0011 | -0.0023 | 0.0047 | |
| *Task effect* | | | | | | | |
| dif loneliness | | Alone | LIFG~RIFG | 0.0012 | -0.0023 | 0.0046 | |
| dif loneliness | | Together | LIFG~RIFG | -0.0015 | -0.0043 | 0.0013 | |
| dif loneliness | | Alone | LIFG~LIFG | -0.0014 | -0.0058 | 0.0028 | |
| dif loneliness | | Together | LIFG~LIFG | 0.0002 | -0.0033 | 0.0038 | |
| dif loneliness | | Alone | LTPJ~LTPJ | -0.0038 | -0.0087 | 0.001 | |
| dif loneliness | | Together | LTPJ~LTPJ | 0.0006 | -0.0035 | 0.0045 | |
| dif loneliness | | Alone | LIFG~LTPJ | -0.0013 | -0.0046 | 0.0021 | |
| dif loneliness | | Together | LIFG~LTPJ | -0.0016 | -0.0044 | 0.0011 | |
| dif loneliness | | Alone | LIFG~RTPJ | 0.0017 | -0.0024 | 0.0057 | |
| dif loneliness | | Together | LIFG~RTPJ | -0.0017 | -0.0049 | 0.0016 | |
| dif loneliness | | Alone | RIFG~RIFG | 0.0026 | -0.0023 | 0.0078 | |
| dif loneliness | | Together | RIFG~RIFG | -0.0054 | -0.0095 | -0.0014 | No overlap |
| dif loneliness | | Alone | RTPJ~RTPJ | -0.0028 | -0.0114 | 0.0057 | |

| Measure | Group | Task | ROI Pairs | Estimate | Lower HPD | Upper HPD | No overlap |
|---|---|---|---|---|---|---|---|
| dif loneliness | | Together | RTPJ~RTPJ | 0.0025 | -0.0049 | 0.0096 | |
| dif loneliness | | Alone | RIFG~LTPJ | 0.0003 | -0.0029 | 0.0037 | |
| dif loneliness | | Together | RIFG~LTPJ | -0.0009 | -0.0035 | 0.0018 | |
| dif loneliness | | Alone | RIFG~RTPJ | 0.002 | -0.0023 | 0.0063 | |
| dif loneliness | | Together | RIFG~RTPJ | -0.0027 | -0.0063 | 0.0008 | |
| *Simple effect* | | | | | | | |
| dif loneliness | Intergen | Alone | LIFG~RIFG | 0.0016 | -0.0039 | 0.0071 | |
| dif loneliness | Intergen | Together | LIFG~RIFG | -0.0015 | -0.0062 | 0.0030 | |
| dif loneliness | Samegen | Alone | LIFG~RIFG | 0.0008 | -0.0031 | 0.0049 | |
| dif loneliness | Samegen | Together | LIFG~RIFG | -0.0015 | -0.0047 | 0.0017 | |
| dif loneliness | Intergen | Alone | LIFG~LIFG | -0.0038 | -0.0106 | 0.0031 | |
| dif loneliness | Intergen | Together | LIFG~LIFG | 0.0021 | -0.0037 | 0.0077 | |
| dif loneliness | Samegen | Alone | LIFG~LIFG | 0.0009 | -0.0043 | 0.0062 | |
| dif loneliness | Samegen | Together | LIFG~LIFG | -0.0016 | -0.0057 | 0.0026 | |
| dif loneliness | Intergen | Alone | LTPJ~LTPJ | -0.0093 | -0.0169 | -0.0016 | No overlap |
| dif loneliness | Intergen | Together | LTPJ~LTPJ | -0.0001 | -0.0066 | 0.0060 | |
| dif loneliness | Samegen | Alone | LTPJ~LTPJ | 0.0017 | -0.0042 | 0.0076 | |
| dif loneliness | Samegen | Together | LTPJ~LTPJ | 0.0015 | -0.0034 | 0.0064 | |
| dif loneliness | Intergen | Alone | LIFG~LTPJ | -0.0017 | -0.0069 | 0.0037 | |
| dif loneliness | Intergen | Together | LIFG~LTPJ | -0.0043 | -0.0088 | -0.0001 | No overlap |
| dif loneliness | Samegen | Alone | LIFG~LTPJ | -0.0009 | -0.0050 | 0.0032 | |
| dif loneliness | Samegen | Together | LIFG~LTPJ | 0.0011 | -0.0022 | 0.0044 | |
| dif loneliness | Intergen | Alone | LIFG~RTPJ | 0.0037 | -0.0028 | 0.0101 | |
| dif loneliness | Intergen | Together | LIFG~RTPJ | -0.0015 | -0.0069 | 0.0038 | |
| dif loneliness | Samegen | Alone | LIFG~RTPJ | -0.0003 | -0.0050 | 0.0044 | |
| dif loneliness | Samegen | Together | LIFG~RTPJ | -0.0018 | -0.0056 | 0.0018 | |
| dif loneliness | Intergen | Alone | RIFG~RIFG | 0.0027 | -0.0053 | 0.0113 | |
| dif loneliness | Intergen | Together | RIFG~RIFG | -0.0071 | -0.0137 | -0.0006 | No overlap |
| dif loneliness | Samegen | Alone | RIFG~RIFG | 0.0026 | -0.0029 | 0.0083 | |
| dif loneliness | Samegen | Together | RIFG~RIFG | -0.0037 | -0.0083 | 0.0007 | |
| dif loneliness | Intergen | Alone | RTPJ~RTPJ | -0.0058 | -0.0199 | 0.0088 | |
| dif loneliness | Intergen | Together | RTPJ~RTPJ | -0.0003 | -0.0126 | 0.0118 | |
| dif loneliness | Samegen | Alone | RTPJ~RTPJ | 0.0003 | -0.0094 | 0.0103 | |
| dif loneliness | Samegen | Together | RTPJ~RTPJ | 0.0053 | -0.0024 | 0.0130 | |
| dif loneliness | Intergen | Alone | RIFG~LTPJ | -0.0004 | -0.0057 | 0.0050 | |
| dif loneliness | Intergen | Together | RIFG~LTPJ | -0.0017 | -0.0062 | 0.0025 | |
| dif loneliness | Samegen | Alone | RIFG~LTPJ | 0.0011 | -0.0027 | 0.0049 | |
| dif loneliness | Samegen | Together | RIFG~LTPJ | -0.0001 | -0.0031 | 0.0030 | |
| dif loneliness | Intergen | Alone | RIFG~RTPJ | -0.0005 | -0.0079 | 0.0064 | |
| dif loneliness | Intergen | Together | RIFG~RTPJ | -0.0033 | -0.0093 | 0.0026 | |
| dif loneliness | Samegen | Alone | RIFG~RTPJ | 0.0045 | -0.0004 | 0.0096 | |
| dif loneliness | Samegen | Together | RIFG~RTPJ | -0.0022 | -0.0063 | 0.0019 | |
| dif loneliness | Intergen | Alone | LIFG~RIFG | 0.0016 | -0.0039 | 0.0071 | |

**S5 Appendix – Table N.** HbO. Contrasts pertaining to loneliness (dif) ~ INS relationship in unstandardised units. Contrasts are presented in the following order: Group, task, and interaction contrasts.

| Measure | Group | Task | ROI Pairs | Estimate | Lower HPD | Upper HPD | No overlap |
|---|---|---|---|---|---|---|---|
| *Group contrast* | | | | | | | |
| dif loneliness | Intergen-Samegen | | LIFG~RIFG | 0.0004 | -0.0042 | 0.0048 | |
| dif loneliness | Intergen-Samegen | | LIFG~LIFG | -0.0005 | -0.0062 | 0.0053 | |
| dif loneliness | Intergen-Samegen | | LTPJ~LTPJ | -0.0063 | -0.0129 | 0.0005 | |
| dif loneliness | Intergen-Samegen | | LIFG~LTPJ | -0.0031 | -0.0076 | 0.0014 | |
| dif loneliness | Intergen-Samegen | | LIFG~RTPJ | 0.0021 | -0.0032 | 0.0072 | |
| dif loneliness | Intergen-Samegen | | RIFG~RIFG | -0.0016 | -0.0082 | 0.0051 | |
| dif loneliness | Intergen-Samegen | | RTPJ~RTPJ | -0.0058 | -0.0177 | 0.0057 | |
| dif loneliness | Intergen-Samegen | | RIFG~LTPJ | -0.0015 | -0.0058 | 0.0028 | |
| dif loneliness | Intergen-Samegen | | RIFG~RTPJ | -0.0031 | -0.0092 | 0.0030 | |
| *Task contrast* | | | | | | | |
| dif loneliness | | Alone-Together | LIFG~RIFG | 0.0027 | -0.0015 | 0.0069 | |
| dif loneliness | | Alone-Together | LIFG~LIFG | -0.0017 | -0.0071 | 0.0037 | |
| dif loneliness | | Alone-Together | LTPJ~LTPJ | -0.0045 | -0.0101 | 0.0014 | |
| dif loneliness | | Alone-Together | LIFG~LTPJ | 0.0003 | -0.0039 | 0.0043 | |
| dif loneliness | | Alone-Together | LIFG~RTPJ | 0.0034 | -0.0017 | 0.0083 | |
| dif loneliness | | Alone-Together | RIFG~RIFG | 0.0081 | 0.0019 | 0.0143 | No overlap |
| dif loneliness | | Alone-Together | RTPJ~RTPJ | -0.0053 | -0.0164 | 0.0056 | |
| dif loneliness | | Alone-Together | RIFG~LTPJ | 0.0012 | -0.0029 | 0.0052 | |
| dif loneliness | | Alone-Together | RIFG~RTPJ | 0.0047 | -0.0006 | 0.0100 | |
| *Interactions* | | | | | | | |
| dif loneliness | Intergen-Samegen | Alone | LIFG~RIFG | 0.0008 | -0.0058 | 0.0078 | |
| dif loneliness | Intergen-Samegen | Together | LIFG~RIFG | 0.0000 | -0.0056 | 0.0055 | |
| dif loneliness | Intergen-Samegen | Alone | LIFG~LIFG | -0.0047 | -0.0133 | 0.0043 | |
| dif loneliness | Intergen-Samegen | Together | LIFG~LIFG | 0.0037 | -0.0037 | 0.0104 | |
| dif loneliness | Intergen-Samegen | Alone | LTPJ~LTPJ | -0.0111 | -0.0205 | -0.0011 | No overlap |
| dif loneliness | Intergen-Samegen | Together | LTPJ~LTPJ | -0.0016 | -0.0094 | 0.0066 | |
| dif loneliness | Intergen-Samegen | Alone | LIFG~LTPJ | -0.0008 | -0.0073 | 0.0059 | |
| dif loneliness | Intergen-Samegen | Together | LIFG~LTPJ | -0.0053 | -0.0106 | 0.0002 | |
| dif loneliness | Intergen-Samegen | Alone | LIFG~RTPJ | 0.0040 | -0.0040 | 0.0119 | |
| dif loneliness | Intergen-Samegen | Together | LIFG~RTPJ | 0.0003 | -0.0060 | 0.0069 | |
| dif loneliness | Intergen-Samegen | Alone | RIFG~RIFG | 0.0001 | -0.0099 | 0.0099 | |
| dif loneliness | Intergen-Samegen | Together | RIFG~RIFG | -0.0034 | -0.0112 | 0.0046 | |
| dif loneliness | Intergen-Samegen | Alone | RTPJ~RTPJ | -0.0060 | -0.0238 | 0.0112 | |
| dif loneliness | Intergen-Samegen | Together | RTPJ~RTPJ | -0.0056 | -0.0199 | 0.0092 | |
| dif loneliness | Intergen-Samegen | Alone | RIFG~LTPJ | -0.0015 | -0.0080 | 0.0050 | |
| dif loneliness | Intergen-Samegen | Together | RIFG~LTPJ | -0.0016 | -0.0068 | 0.0038 | |
| dif loneliness | Intergen-Samegen | Alone | RIFG~RTPJ | -0.0050 | -0.0137 | 0.0038 | |
| dif loneliness | Intergen-Samegen | Together | RIFG~RTPJ | -0.0011 | -0.0084 | 0.0062 | |
| dif loneliness | Intergen | Alone-Together | LIFG~RIFG | 0.0031 | -0.0039 | 0.0101 | |
| dif loneliness | Samegen | Alone-Together | LIFG~RIFG | 0.0023 | -0.0026 | 0.0072 | |
| dif loneliness | Intergen | Alone-Together | LIFG~LIFG | -0.0058 | -0.0142 | 0.0033 | |
| dif loneliness | Samegen | Alone-Together | LIFG~LIFG | 0.0024 | -0.0041 | 0.0089 | |

| Measure | Group | Task | ROI Pairs | Estimate | Lower HPD | Upper HPD | No overlap |
|---|---|---|---|---|---|---|---|
| dif loneliness | Intergen | Alone-Together | LTPJ~LTPJ | -0.0092 | -0.0186 | 0.0000 | |
| dif loneliness | Samegen | Alone-Together | LTPJ~LTPJ | 0.0002 | -0.0066 | 0.0070 | |
| dif loneliness | Intergen | Alone-Together | LIFG~LTPJ | 0.0026 | -0.0040 | 0.0091 | |
| dif loneliness | Samegen | Alone-Together | LIFG~LTPJ | -0.0019 | -0.0068 | 0.0029 | |
| dif loneliness | Intergen | Alone-Together | LIFG~RTPJ | 0.0052 | -0.0028 | 0.0133 | |
| dif loneliness | Samegen | Alone-Together | LIFG~RTPJ | 0.0015 | -0.0042 | 0.0072 | |
| dif loneliness | Intergen | Alone-Together | RIFG~RIFG | 0.0098 | -0.0006 | 0.0198 | |
| dif loneliness | Samegen | Alone-Together | RIFG~RIFG | 0.0063 | -0.0003 | 0.0133 | |
| dif loneliness | Intergen | Alone-Together | RTPJ~RTPJ | -0.0055 | -0.0242 | 0.0132 | |
| dif loneliness | Samegen | Alone-Together | RTPJ~RTPJ | -0.0051 | -0.0170 | 0.0067 | |
| dif loneliness | Intergen | Alone-Together | RIFG~LTPJ | 0.0013 | -0.0053 | 0.0080 | |
| dif loneliness | Samegen | Alone-Together | RIFG~LTPJ | 0.0012 | -0.0035 | 0.0059 | |
| dif loneliness | Intergen | Alone-Together | RIFG~RTPJ | 0.0027 | -0.0060 | 0.0112 | |
| dif loneliness | Samegen | Alone-Together | RIFG~RTPJ | 0.0066 | 0.0007 | 0.0126 | No overlap |

**S5 Appendix – Table O.** HbO. Relationships between social closeness (sum) and INS in unstandardised units. Effects are presented in the following order: Main, group, task, and simple effects.

| Measure | Group | Task | ROI Pairs | Estimate | Lower HPD | Upper HPD | No overlap |
|---|---|---|---|---|---|---|---|
| *Main effect* | | | | | | | |
| sum social closeness | | | LIFG~RIFG | 0.0007 | -0.0006 | 0.0021 | |
| sum social closeness | | | LIFG~LIFG | -0.0007 | -0.0025 | 0.0011 | |
| sum social closeness | | | LTPJ~LTPJ | 0.0005 | -0.0015 | 0.0025 | |
| sum social closeness | | | LIFG~LTPJ | 0.0003 | -0.0011 | 0.0016 | |
| sum social closeness | | | LIFG~RTPJ | 0 | -0.0016 | 0.0017 | |
| sum social closeness | | | RIFG~RIFG | 0 | -0.002 | 0.002 | |
| sum social closeness | | | RTPJ~RTPJ | -0.0012 | -0.0047 | 0.0023 | |
| sum social closeness | | | RIFG~LTPJ | 0.0006 | -0.0007 | 0.0019 | |
| sum social closeness | | | RIFG~RTPJ | 0.0021 | 0.0004 | 0.0039 | No overlap |
| *Group effect* | | | | | | | |
| sum social closeness | Intergen | | LIFG~RIFG | 0.0006 | -0.001 | 0.0022 | |
| sum social closeness | Samegen | | LIFG~RIFG | 0.0009 | -0.0014 | 0.0033 | |
| sum social closeness | Intergen | | LIFG~LIFG | -0.0016 | -0.0036 | 0.0003 | |
| sum social closeness | Samegen | | LIFG~LIFG | 0.0002 | -0.0027 | 0.0032 | |
| sum social closeness | Intergen | | LTPJ~LTPJ | -0.0005 | -0.0029 | 0.0018 | |
| sum social closeness | Samegen | | LTPJ~LTPJ | 0.0015 | -0.0018 | 0.0049 | |
| sum social closeness | Intergen | | LIFG~LTPJ | 0.0004 | -0.0012 | 0.0018 | |
| sum social closeness | Samegen | | LIFG~LTPJ | 0.0002 | -0.0021 | 0.0025 | |
| sum social closeness | Intergen | | LIFG~RTPJ | -0.0006 | -0.0025 | 0.0012 | |
| sum social closeness | Samegen | | LIFG~RTPJ | 0.0006 | -0.0021 | 0.0032 | |
| sum social closeness | Intergen | | RIFG~RIFG | 0.0022 | -0.0001 | 0.0044 | |
| sum social closeness | Samegen | | RIFG~RIFG | -0.0021 | -0.0054 | 0.0012 | |
| sum social closeness | Intergen | | RTPJ~RTPJ | 0.0001 | -0.0039 | 0.0043 | |
| sum social closeness | Samegen | | RTPJ~RTPJ | -0.0025 | -0.0081 | 0.0033 | |
| sum social closeness | Intergen | | RIFG~LTPJ | 0.0006 | -0.0008 | 0.0021 | |
| sum social closeness | Samegen | | RIFG~LTPJ | 0.0005 | -0.0017 | 0.0027 | |
| sum social closeness | Intergen | | RIFG~RTPJ | -0.0001 | -0.0022 | 0.0019 | |
| sum social closeness | Samegen | | RIFG~RTPJ | 0.0044 | 0.0016 | 0.0073 | No overlap |
| *Task effect* | | | | | | | |
| sum social closeness | | Alone | LIFG~RIFG | 0.0011 | -0.001 | 0.0033 | |
| sum social closeness | | Together | LIFG~RIFG | 0.0004 | -0.0013 | 0.0021 | |
| sum social closeness | | Alone | LIFG~LIFG | -0.0029 | -0.0057 | -0.0003 | No overlap |
| sum social closeness | | Together | LIFG~LIFG | 0.0015 | -0.0007 | 0.0036 | |
| sum social closeness | | Alone | LTPJ~LTPJ | 0.0005 | -0.0025 | 0.0035 | |
| sum social closeness | | Together | LTPJ~LTPJ | 0.0005 | -0.0019 | 0.0029 | |
| sum social closeness | | Alone | LIFG~LTPJ | -0.0001 | -0.0022 | 0.0019 | |
| sum social closeness | | Together | LIFG~LTPJ | 0.0007 | -0.001 | 0.0023 | |
| sum social closeness | | Alone | LIFG~RTPJ | 0.0009 | -0.0016 | 0.0032 | |
| sum social closeness | | Together | LIFG~RTPJ | -0.0009 | -0.0029 | 0.0011 | |
| sum social closeness | | Alone | RIFG~RIFG | -0.0011 | -0.0041 | 0.0019 | |
| sum social closeness | | Together | RIFG~RIFG | 0.0012 | -0.0012 | 0.0037 | |
| sum social closeness | | Alone | RTPJ~RTPJ | 0.001 | -0.0042 | 0.0063 | |

| Measure | Group | Task | ROI Pairs | Estimate | Lower HPD | Upper HPD | No overlap |
|---|---|---|---|---|---|---|---|
| sum social closeness | | Together | RTPJ~RTPJ | -0.0033 | -0.0078 | 0.001 | |
| sum social closeness | | Alone | RIFG~LTPJ | 0.0006 | -0.0014 | 0.0027 | |
| sum social closeness | | Together | RIFG~LTPJ | 0.0005 | -0.001 | 0.0022 | |
| sum social closeness | | Alone | RIFG~RTPJ | 0.0033 | 0.0006 | 0.0059 | No overlap |
| sum social closeness | | Together | RIFG~RTPJ | 0.001 | -0.0011 | 0.0031 | |
| *Simple effect* | | | | | | | |
| sum social closeness | Intergen | Alone | LIFG~RIFG | 0.0007 | -0.0017 | 0.0030 | |
| sum social closeness | Intergen | Together | LIFG~RIFG | 0.0004 | -0.0014 | 0.0023 | |
| sum social closeness | Samegen | Alone | LIFG~RIFG | 0.0015 | -0.0020 | 0.0051 | |
| sum social closeness | Samegen | Together | LIFG~RIFG | 0.0003 | -0.0025 | 0.0031 | |
| sum social closeness | Intergen | Alone | LIFG~LIFG | -0.0029 | -0.0058 | 0.0002 | |
| sum social closeness | Intergen | Together | LIFG~LIFG | -0.0003 | -0.0026 | 0.0021 | |
| sum social closeness | Samegen | Alone | LIFG~LIFG | -0.0029 | -0.0074 | 0.0018 | |
| sum social closeness | Samegen | Together | LIFG~LIFG | 0.0033 | -0.0002 | 0.0069 | |
| sum social closeness | Intergen | Alone | LTPJ~LTPJ | -0.0008 | -0.0041 | 0.0027 | |
| sum social closeness | Intergen | Together | LTPJ~LTPJ | -0.0002 | -0.0029 | 0.0025 | |
| sum social closeness | Samegen | Alone | LTPJ~LTPJ | 0.0018 | -0.0031 | 0.0068 | |
| sum social closeness | Samegen | Together | LTPJ~LTPJ | 0.0012 | -0.0029 | 0.0051 | |
| sum social closeness | Intergen | Alone | LIFG~LTPJ | -0.0005 | -0.0027 | 0.0018 | |
| sum social closeness | Intergen | Together | LIFG~LTPJ | 0.0012 | -0.0006 | 0.0030 | |
| sum social closeness | Samegen | Alone | LIFG~LTPJ | 0.0002 | -0.0031 | 0.0038 | |
| sum social closeness | Samegen | Together | LIFG~LTPJ | 0.0001 | -0.0026 | 0.0029 | |
| sum social closeness | Intergen | Alone | LIFG~RTPJ | -0.0006 | -0.0033 | 0.0021 | |
| sum social closeness | Intergen | Together | LIFG~RTPJ | -0.0006 | -0.0028 | 0.0017 | |
| sum social closeness | Samegen | Alone | LIFG~RTPJ | 0.0024 | -0.0017 | 0.0064 | |
| sum social closeness | Samegen | Together | LIFG~RTPJ | -0.0012 | -0.0045 | 0.0020 | |
| sum social closeness | Intergen | Alone | RIFG~RIFG | 0.0015 | -0.0020 | 0.0047 | |
| sum social closeness | Intergen | Together | RIFG~RIFG | 0.0029 | 0.0001 | 0.0054 | No overlap |
| sum social closeness | Samegen | Alone | RIFG~RIFG | -0.0037 | -0.0087 | 0.0013 | |
| sum social closeness | Samegen | Together | RIFG~RIFG | -0.0004 | -0.0044 | 0.0036 | |
| sum social closeness | Intergen | Alone | RTPJ~RTPJ | 0.0011 | -0.0049 | 0.0067 | |
| sum social closeness | Intergen | Together | RTPJ~RTPJ | -0.0009 | -0.0064 | 0.0046 | |
| sum social closeness | Samegen | Alone | RTPJ~RTPJ | 0.0008 | -0.0078 | 0.0097 | |
| sum social closeness | Samegen | Together | RTPJ~RTPJ | -0.0058 | -0.0126 | 0.0013 | |
| sum social closeness | Intergen | Alone | RIFG~LTPJ | -0.0001 | -0.0024 | 0.0021 | |
| sum social closeness | Intergen | Together | RIFG~LTPJ | 0.0013 | -0.0004 | 0.0031 | |
| sum social closeness | Samegen | Alone | RIFG~LTPJ | 0.0014 | -0.0020 | 0.0047 | |
| sum social closeness | Samegen | Together | RIFG~LTPJ | -0.0003 | -0.0029 | 0.0025 | |
| sum social closeness | Intergen | Alone | RIFG~RTPJ | -0.0008 | -0.0038 | 0.0021 | |
| sum social closeness | Intergen | Together | RIFG~RTPJ | 0.0006 | -0.0019 | 0.0030 | |
| sum social closeness | Samegen | Alone | RIFG~RTPJ | 0.0073 | 0.0031 | 0.0117 | No overlap |
| sum social closeness | Samegen | Together | RIFG~RTPJ | 0.0014 | -0.0019 | 0.0049 | |
| sum social closeness | Intergen | Alone | LIFG~RIFG | 0.0007 | -0.0017 | 0.0030 | |

**S5 Appendix – Table P.** HbO. Contrasts pertaining to social closeness (sum) ~ INS relationship in unstandardised units. Contrasts are presented in the following order: Group, task, and interaction contrasts.

| Measure | Group | Task | ROI Pairs | Estimate | Lower HPD | Upper HPD | No overlap |
|---|---|---|---|---|---|---|---|
| *Group contrast* | | | | | | | |
| sum social closeness | Int-Samegen | | LIFG~RIFG | -0.0004 | -0.0032 | 0.0024 | |
| sum social closeness | Int-Samegen | | LIFG~LIFG | -0.0018 | -0.0054 | 0.0018 | |
| sum social closeness | Int-Samegen | | LTPJ~LTPJ | -0.0020 | -0.0060 | 0.0021 | |
| sum social closeness | Int-Samegen | | LIFG~LTPJ | 0.0002 | -0.0025 | 0.0030 | |
| sum social closeness | Int-Samegen | | LIFG~RTPJ | -0.0012 | -0.0044 | 0.0022 | |
| sum social closeness | Int-Samegen | | RIFG~RIFG | 0.0042 | 0.0003 | 0.0083 | No overlap |
| sum social closeness | Int-Samegen | | RTPJ~RTPJ | 0.0026 | -0.0045 | 0.0095 | |
| sum social closeness | Int-Samegen | | RIFG~LTPJ | 0.0001 | -0.0026 | 0.0027 | |
| sum social closeness | Int-Samegen | | RIFG~RTPJ | -0.0045 | -0.0079 | -0.0009 | No overlap |
| *Task contrast* | | | | | | | |
| sum social closeness | | Alone-Together | LIFG~RIFG | 0.0007 | -0.0019 | 0.0034 | |
| sum social closeness | | Alone-Together | LIFG~LIFG | -0.0044 | -0.0079 | -0.0010 | No overlap |
| sum social closeness | | Alone-Together | LTPJ~LTPJ | 0.0000 | -0.0036 | 0.0035 | |
| sum social closeness | | Alone-Together | LIFG~LTPJ | -0.0008 | -0.0034 | 0.0017 | |
| sum social closeness | | Alone-Together | LIFG~RTPJ | 0.0018 | -0.0012 | 0.0048 | |
| sum social closeness | | Alone-Together | RIFG~RIFG | -0.0024 | -0.0062 | 0.0014 | |
| sum social closeness | | Alone-Together | RTPJ~RTPJ | 0.0043 | -0.0026 | 0.0109 | |
| sum social closeness | | Alone-Together | RIFG~LTPJ | 0.0001 | -0.0024 | 0.0027 | |
| sum social closeness | | Alone-Together | RIFG~RTPJ | 0.0022 | -0.0010 | 0.0054 | |
| *Interactions* | | | | | | | |
| sum social closeness | Int-Samegen | Alone | LIFG~RIFG | -0.0008 | -0.0050 | 0.0037 | |
| sum social closeness | Int-Samegen | Together | LIFG~RIFG | 0.0001 | -0.0032 | 0.0034 | |
| sum social closeness | Int-Samegen | Alone | LIFG~LIFG | 0.0000 | -0.0056 | 0.0054 | |
| sum social closeness | Int-Samegen | Together | LIFG~LIFG | -0.0037 | -0.0080 | 0.0006 | |
| sum social closeness | Int-Samegen | Alone | LTPJ~LTPJ | -0.0026 | -0.0085 | 0.0035 | |
| sum social closeness | Int-Samegen | Together | LTPJ~LTPJ | -0.0015 | -0.0063 | 0.0034 | |
| sum social closeness | Int-Samegen | Alone | LIFG~LTPJ | -0.0007 | -0.0047 | 0.0036 | |
| sum social closeness | Int-Samegen | Together | LIFG~LTPJ | 0.0011 | -0.0023 | 0.0044 | |
| sum social closeness | Int-Samegen | Alone | LIFG~RTPJ | -0.0030 | -0.0080 | 0.0018 | |
| sum social closeness | Int-Samegen | Together | LIFG~RTPJ | 0.0006 | -0.0034 | 0.0044 | |
| sum social closeness | Int-Samegen | Alone | RIFG~RIFG | 0.0052 | -0.0008 | 0.0112 | |
| sum social closeness | Int-Samegen | Together | RIFG~RIFG | 0.0033 | -0.0014 | 0.0082 | |
| sum social closeness | Int-Samegen | Alone | RTPJ~RTPJ | 0.0003 | -0.0105 | 0.0105 | |
| sum social closeness | Int-Samegen | Together | RTPJ~RTPJ | 0.0050 | -0.0041 | 0.0136 | |
| sum social closeness | Int-Samegen | Alone | RIFG~LTPJ | -0.0014 | -0.0056 | 0.0025 | |
| sum social closeness | Int-Samegen | Together | RIFG~LTPJ | 0.0016 | -0.0015 | 0.0049 | |
| sum social closeness | Int-Samegen | Alone | RIFG~RTPJ | -0.0081 | -0.0132 | -0.0026 | No overlap |
| sum social closeness | Int-Samegen | Together | RIFG~RTPJ | -0.0008 | -0.0050 | 0.0034 | |
| sum social closeness | Intergen | Alone-Together | LIFG~RIFG | 0.0003 | -0.0025 | 0.0033 | |
| sum social closeness | Samegen | Alone-Together | LIFG~RIFG | 0.0012 | -0.0033 | 0.0055 | |
| sum social closeness | Intergen | Alone-Together | LIFG~LIFG | -0.0026 | -0.0062 | 0.0012 | |
| sum social closeness | Samegen | Alone-Together | LIFG~LIFG | -0.0063 | -0.0120 | -0.0006 | No overlap |

| Measure | Group | Task | ROI Pairs | Estimate | Lower HPD | Upper HPD | No overlap |
|---|---|---|---|---|---|---|---|
| sum social closeness | Intergen | Alone-Together | LTPJ~LTPJ | -0.0005 | -0.0044 | 0.0034 | |
| sum social closeness | Samegen | Alone-Together | LTPJ~LTPJ | 0.0006 | -0.0054 | 0.0066 | |
| sum social closeness | Intergen | Alone-Together | LIFG~LTPJ | -0.0017 | -0.0044 | 0.0011 | |
| sum social closeness | Samegen | Alone-Together | LIFG~LTPJ | 0.0001 | -0.0042 | 0.0043 | |
| sum social closeness | Intergen | Alone-Together | LIFG~RTPJ | 0.0000 | -0.0033 | 0.0035 | |
| sum social closeness | Samegen | Alone-Together | LIFG~RTPJ | 0.0036 | -0.0014 | 0.0086 | |
| sum social closeness | Intergen | Alone-Together | RIFG~RIFG | -0.0014 | -0.0056 | 0.0026 | |
| sum social closeness | Samegen | Alone-Together | RIFG~RIFG | -0.0033 | -0.0095 | 0.0029 | |
| sum social closeness | Intergen | Alone-Together | RTPJ~RTPJ | 0.0020 | -0.0057 | 0.0101 | |
| sum social closeness | Samegen | Alone-Together | RTPJ~RTPJ | 0.0066 | -0.0043 | 0.0175 | |
| sum social closeness | Intergen | Alone-Together | RIFG~LTPJ | -0.0014 | -0.0042 | 0.0014 | |
| sum social closeness | Samegen | Alone-Together | RIFG~LTPJ | 0.0016 | -0.0028 | 0.0057 | |
| sum social closeness | Intergen | Alone-Together | RIFG~RTPJ | -0.0014 | -0.0050 | 0.0023 | |
| sum social closeness | Samegen | Alone-Together | RIFG~RTPJ | 0.0059 | 0.0007 | 0.0113 | No overlap |

**S5 Appendix – Table Q.** HbO. Relationships between social closeness (dif) and INS in unstandardised units. Effects are presented in the following order: Main, group, task, and simple effects.

| Measure | Group | Task | ROI Pairs | Estimate | Lower HPD | Upper HPD | No overlap |
|---|---|---|---|---|---|---|---|
| *Main effect* | | | | | | | |
| dif social closeness | | | LIFG~RIFG | -0.0004 | -0.0027 | 0.0018 | |
| dif social closeness | | | LIFG~LIFG | -0.0003 | -0.0031 | 0.0025 | |
| dif social closeness | | | LTPJ~LTPJ | -0.001 | -0.0042 | 0.0021 | |
| dif social closeness | | | LIFG~LTPJ | -0.0005 | -0.0026 | 0.0017 | |
| dif social closeness | | | LIFG~RTPJ | -0.0014 | -0.0039 | 0.0012 | |
| dif social closeness | | | RIFG~RIFG | 0.0018 | -0.0015 | 0.005 | |
| dif social closeness | | | RTPJ~RTPJ | 0.0018 | -0.0035 | 0.007 | |
| dif social closeness | | | RIFG~LTPJ | 0.0011 | -0.001 | 0.0032 | |
| dif social closeness | | | RIFG~RTPJ | -0.002 | -0.0048 | 0.0009 | |
| *Group effect* | | | | | | | |
| dif social closeness | Intergen | | LIFG~RIFG | -0.0002 | -0.0032 | 0.0027 | |
| dif social closeness | Samegen | | LIFG~RIFG | -0.0005 | -0.004 | 0.0028 | |
| dif social closeness | Intergen | | LIFG~LIFG | -0.0011 | -0.0048 | 0.0025 | |
| dif social closeness | Samegen | | LIFG~LIFG | 0.0005 | -0.0037 | 0.0047 | |
| dif social closeness | Intergen | | LTPJ~LTPJ | -0.0005 | -0.0047 | 0.0038 | |
| dif social closeness | Samegen | | LTPJ~LTPJ | -0.0015 | -0.0063 | 0.0033 | |
| dif social closeness | Intergen | | LIFG~LTPJ | -0.0004 | -0.0032 | 0.0026 | |
| dif social closeness | Samegen | | LIFG~LTPJ | -0.0005 | -0.0038 | 0.0027 | |
| dif social closeness | Intergen | | LIFG~RTPJ | -0.0027 | -0.0061 | 0.0008 | |
| dif social closeness | Samegen | | LIFG~RTPJ | 0 | -0.0039 | 0.0035 | |
| dif social closeness | Intergen | | RIFG~RIFG | 0.0004 | -0.0039 | 0.0046 | |
| dif social closeness | Samegen | | RIFG~RIFG | 0.0031 | -0.0016 | 0.008 | |
| dif social closeness | Intergen | | RTPJ~RTPJ | 0.0005 | -0.0073 | 0.0086 | |
| dif social closeness | Samegen | | RTPJ~RTPJ | 0.0032 | -0.0038 | 0.0103 | |
| dif social closeness | Intergen | | RIFG~LTPJ | 0.0013 | -0.0014 | 0.004 | |
| dif social closeness | Samegen | | RIFG~LTPJ | 0.0009 | -0.0023 | 0.0041 | |
| dif social closeness | Intergen | | RIFG~RTPJ | -0.0016 | -0.0055 | 0.0023 | |
| dif social closeness | Samegen | | RIFG~RTPJ | -0.0024 | -0.0066 | 0.0018 | |
| *Task effect* | | | | | | | |
| dif social closeness | | Alone | LIFG~RIFG | -0.0007 | -0.0042 | 0.0027 | |
| dif social closeness | | Together | LIFG~RIFG | -0.0001 | -0.0027 | 0.0027 | |
| dif social closeness | | Alone | LIFG~LIFG | 0.0013 | -0.0028 | 0.0057 | |
| dif social closeness | | Together | LIFG~LIFG | -0.002 | -0.0054 | 0.0014 | |
| dif social closeness | | Alone | LTPJ~LTPJ | -0.0002 | -0.0049 | 0.0046 | |
| dif social closeness | | Together | LTPJ~LTPJ | -0.0019 | -0.0056 | 0.002 | |
| dif social closeness | | Alone | LIFG~LTPJ | -0.0006 | -0.0038 | 0.0027 | |
| dif social closeness | | Together | LIFG~LTPJ | -0.0003 | -0.0028 | 0.0024 | |
| dif social closeness | | Alone | LIFG~RTPJ | -0.0023 | -0.0062 | 0.0016 | |
| dif social closeness | | Together | LIFG~RTPJ | -0.0004 | -0.0035 | 0.0026 | |
| dif social closeness | | Alone | RIFG~RIFG | 0.0024 | -0.0026 | 0.0072 | |
| dif social closeness | | Together | RIFG~RIFG | 0.0012 | -0.0027 | 0.0051 | |
| dif social closeness | | Alone | RTPJ~RTPJ | -0.0005 | -0.0085 | 0.0073 | |

| Measure | Group | Task | ROI Pairs | Estimate | Lower HPD | Upper HPD | No overlap |
|---|---|---|---|---|---|---|---|
| dif social closeness | | Together | RTPJ~RTPJ | 0.0041 | -0.0022 | 0.0109 | |
| dif social closeness | | Alone | RIFG~LTPJ | 0.0014 | -0.0019 | 0.0048 | |
| dif social closeness | | Together | RIFG~LTPJ | 0.0008 | -0.0017 | 0.0034 | |
| dif social closeness | | Alone | RIFG~RTPJ | -0.0037 | -0.008 | 0.0006 | |
| dif social closeness | | Together | RIFG~RTPJ | -0.0004 | -0.0038 | 0.003 | |
| *Simple effect* | | | | | | | |
| dif social closeness | Intergen | Alone | LIFG~RIFG | -0.0019 | -0.0063 | 0.0027 | |
| dif social closeness | Intergen | Together | LIFG~RIFG | 0.0014 | -0.0020 | 0.0049 | |
| dif social closeness | Samegen | Alone | LIFG~RIFG | 0.0005 | -0.0047 | 0.0056 | |
| dif social closeness | Samegen | Together | LIFG~RIFG | -0.0017 | -0.0058 | 0.0024 | |
| dif social closeness | Intergen | Alone | LIFG~LIFG | -0.0019 | -0.0077 | 0.0037 | |
| dif social closeness | Intergen | Together | LIFG~LIFG | -0.0004 | -0.0047 | 0.0040 | |
| dif social closeness | Samegen | Alone | LIFG~LIFG | 0.0045 | -0.0018 | 0.0109 | |
| dif social closeness | Samegen | Together | LIFG~LIFG | -0.0035 | -0.0088 | 0.0014 | |
| dif social closeness | Intergen | Alone | LTPJ~LTPJ | 0.0002 | -0.0059 | 0.0065 | |
| dif social closeness | Intergen | Together | LTPJ~LTPJ | -0.0013 | -0.0062 | 0.0036 | |
| dif social closeness | Samegen | Alone | LTPJ~LTPJ | -0.0006 | -0.0077 | 0.0065 | |
| dif social closeness | Samegen | Together | LTPJ~LTPJ | -0.0025 | -0.0081 | 0.0033 | |
| dif social closeness | Intergen | Alone | LIFG~LTPJ | 0.0002 | -0.0042 | 0.0045 | |
| dif social closeness | Intergen | Together | LIFG~LTPJ | -0.0008 | -0.0042 | 0.0026 | |
| dif social closeness | Samegen | Alone | LIFG~LTPJ | -0.0014 | -0.0062 | 0.0035 | |
| dif social closeness | Samegen | Together | LIFG~LTPJ | 0.0003 | -0.0036 | 0.0042 | |
| dif social closeness | Intergen | Alone | LIFG~RTPJ | -0.0031 | -0.0083 | 0.0022 | |
| dif social closeness | Intergen | Together | LIFG~RTPJ | -0.0024 | -0.0065 | 0.0018 | |
| dif social closeness | Samegen | Alone | LIFG~RTPJ | -0.0016 | -0.0071 | 0.0042 | |
| dif social closeness | Samegen | Together | LIFG~RTPJ | 0.0015 | -0.0028 | 0.0060 | |
| dif social closeness | Intergen | Alone | RIFG~RIFG | -0.0037 | -0.0102 | 0.0027 | |
| dif social closeness | Intergen | Together | RIFG~RIFG | 0.0044 | -0.0005 | 0.0095 | |
| dif social closeness | Samegen | Alone | RIFG~RIFG | 0.0084 | 0.0012 | 0.0158 | No overlap |
| dif social closeness | Samegen | Together | RIFG~RIFG | -0.0021 | -0.0080 | 0.0039 | |
| dif social closeness | Intergen | Alone | RTPJ~RTPJ | -0.0015 | -0.0138 | 0.0099 | |
| dif social closeness | Intergen | Together | RTPJ~RTPJ | 0.0023 | -0.0081 | 0.0122 | |
| dif social closeness | Samegen | Alone | RTPJ~RTPJ | 0.0005 | -0.0102 | 0.0108 | |
| dif social closeness | Samegen | Together | RTPJ~RTPJ | 0.0059 | -0.0023 | 0.0143 | |
| dif social closeness | Intergen | Alone | RIFG~LTPJ | 0.0000 | -0.0042 | 0.0043 | |
| dif social closeness | Intergen | Together | RIFG~LTPJ | 0.0026 | -0.0005 | 0.0060 | |
| dif social closeness | Samegen | Alone | RIFG~LTPJ | 0.0027 | -0.0021 | 0.0078 | |
| dif social closeness | Samegen | Together | RIFG~LTPJ | -0.0010 | -0.0048 | 0.0029 | |
| dif social closeness | Intergen | Alone | RIFG~RTPJ | -0.0029 | -0.0087 | 0.0029 | |
| dif social closeness | Intergen | Together | RIFG~RTPJ | -0.0004 | -0.0049 | 0.0043 | |
| dif social closeness | Samegen | Alone | RIFG~RTPJ | -0.0044 | -0.0107 | 0.0020 | |
| dif social closeness | Samegen | Together | RIFG~RTPJ | -0.0004 | -0.0053 | 0.0046 | |
| dif social closeness | Intergen | Alone | LIFG~RIFG | -0.0019 | -0.0063 | 0.0027 | |

**S5 Appendix – Table R.** HbO. Contrasts pertaining to social closeness (dif) ~ INS relationship in unstandardised units. Contrasts are presented in the following order: Group, task, and interaction contrasts.

| Measure | Group | Task | ROI Pairs | Estimate | Lower HPD | Upper HPD | No overlap |
|---|---|---|---|---|---|---|---|
| *Group contrast* | | | | | | | |
| dif social closeness | Int-Samegen | | LIFG~RIFG | 0.0003 | -0.0041 | 0.0048 | |
| dif social closeness | Int-Samegen | | LIFG~LIFG | -0.0016 | -0.0072 | 0.0040 | |
| dif social closeness | Int-Samegen | | LTPJ~LTPJ | 0.0010 | -0.0054 | 0.0073 | |
| dif social closeness | Int-Samegen | | LIFG~LTPJ | 0.0002 | -0.0041 | 0.0046 | |
| dif social closeness | Int-Samegen | | LIFG~RTPJ | -0.0027 | -0.0077 | 0.0024 | |
| dif social closeness | Int-Samegen | | RIFG~RIFG | -0.0028 | -0.0091 | 0.0037 | |
| dif social closeness | Int-Samegen | | RTPJ~RTPJ | -0.0028 | -0.0136 | 0.0079 | |
| dif social closeness | Int-Samegen | | RIFG~LTPJ | 0.0004 | -0.0037 | 0.0045 | |
| dif social closeness | Int-Samegen | | RIFG~RTPJ | 0.0007 | -0.0050 | 0.0064 | |
| *Task contrast* | | | | | | | |
| dif social closeness | | Alone-Together | LIFG~RIFG | -0.0005 | -0.0047 | 0.0037 | |
| dif social closeness | | Alone-Together | LIFG~LIFG | 0.0033 | -0.0021 | 0.0085 | |
| dif social closeness | | Alone-Together | LTPJ~LTPJ | 0.0017 | -0.0040 | 0.0073 | |
| dif social closeness | | Alone-Together | LIFG~LTPJ | -0.0003 | -0.0043 | 0.0037 | |
| dif social closeness | | Alone-Together | LIFG~RTPJ | -0.0019 | -0.0067 | 0.0028 | |
| dif social closeness | | Alone-Together | RIFG~RIFG | 0.0012 | -0.0049 | 0.0073 | |
| dif social closeness | | Alone-Together | RTPJ~RTPJ | -0.0046 | -0.0145 | 0.0054 | |
| dif social closeness | | Alone-Together | RIFG~LTPJ | 0.0005 | -0.0035 | 0.0047 | |
| dif social closeness | | Alone-Together | RIFG~RTPJ | -0.0033 | -0.0084 | 0.0020 | |
| *Interactions* | | | | | | | |
| dif social closeness | Int-Samegen | Alone | LIFG~RIFG | -0.0024 | -0.0093 | 0.0044 | |
| dif social closeness | Int-Samegen | Together | LIFG~RIFG | 0.0031 | -0.0022 | 0.0086 | |
| dif social closeness | Int-Samegen | Alone | LIFG~LIFG | -0.0064 | -0.0150 | 0.0023 | |
| dif social closeness | Int-Samegen | Together | LIFG~LIFG | 0.0032 | -0.0034 | 0.0101 | |
| dif social closeness | Int-Samegen | Alone | LTPJ~LTPJ | 0.0008 | -0.0084 | 0.0103 | |
| dif social closeness | Int-Samegen | Together | LTPJ~LTPJ | 0.0012 | -0.0063 | 0.0085 | |
| dif social closeness | Int-Samegen | Alone | LIFG~LTPJ | 0.0016 | -0.0049 | 0.0081 | |
| dif social closeness | Int-Samegen | Together | LIFG~LTPJ | -0.0011 | -0.0064 | 0.0041 | |
| dif social closeness | Int-Samegen | Alone | LIFG~RTPJ | -0.0014 | -0.0091 | 0.0062 | |
| dif social closeness | Int-Samegen | Together | LIFG~RTPJ | -0.0039 | -0.0098 | 0.0021 | |
| dif social closeness | Int-Samegen | Alone | RIFG~RIFG | -0.0121 | -0.0221 | -0.0026 | No overlap |
| dif social closeness | Int-Samegen | Together | RIFG~RIFG | 0.0066 | -0.0011 | 0.0144 | |
| dif social closeness | Int-Samegen | Alone | RTPJ~RTPJ | -0.0020 | -0.0180 | 0.0138 | |
| dif social closeness | Int-Samegen | Together | RTPJ~RTPJ | -0.0036 | -0.0167 | 0.0095 | |
| dif social closeness | Int-Samegen | Alone | RIFG~LTPJ | -0.0028 | -0.0094 | 0.0034 | |
| dif social closeness | Int-Samegen | Together | RIFG~LTPJ | 0.0036 | -0.0015 | 0.0086 | |
| dif social closeness | Int-Samegen | Alone | RIFG~RTPJ | 0.0015 | -0.0073 | 0.0097 | |
| dif social closeness | Int-Samegen | Together | RIFG~RTPJ | 0.0000 | -0.0068 | 0.0068 | |
| dif social closeness | Intergen | Alone-Together | LIFG~RIFG | -0.0033 | -0.0086 | 0.0023 | |
| dif social closeness | Samegen | Alone-Together | LIFG~RIFG | 0.0022 | -0.0040 | 0.0087 | |
| dif social closeness | Intergen | Alone-Together | LIFG~LIFG | -0.0015 | -0.0084 | 0.0057 | |
| dif social closeness | Samegen | Alone-Together | LIFG~LIFG | 0.0081 | 0.0000 | 0.0159 | |

| Measure | Group | Task | ROI Pairs | Estimate | Lower HPD | Upper HPD | No overlap |
|---|---|---|---|---|---|---|---|
| dif social closeness | Intergen | Alone-Together | LTPJ~LTPJ | 0.0015 | -0.0060 | 0.0086 | |
| dif social closeness | Samegen | Alone-Together | LTPJ~LTPJ | 0.0019 | -0.0065 | 0.0107 | |
| dif social closeness | Intergen | Alone-Together | LIFG~LTPJ | 0.0010 | -0.0042 | 0.0062 | |
| dif social closeness | Samegen | Alone-Together | LIFG~LTPJ | -0.0017 | -0.0077 | 0.0044 | |
| dif social closeness | Intergen | Alone-Together | LIFG~RTPJ | -0.0007 | -0.0072 | 0.0058 | |
| dif social closeness | Samegen | Alone-Together | LIFG~RTPJ | -0.0031 | -0.0101 | 0.0036 | |
| dif social closeness | Intergen | Alone-Together | RIFG~RIFG | -0.0081 | -0.0160 | -0.0002 | No overlap |
| dif social closeness | Samegen | Alone-Together | RIFG~RIFG | 0.0105 | 0.0013 | 0.0198 | No overlap |
| dif social closeness | Intergen | Alone-Together | RTPJ~RTPJ | -0.0038 | -0.0192 | 0.0111 | |
| dif social closeness | Samegen | Alone-Together | RTPJ~RTPJ | -0.0054 | -0.0182 | 0.0070 | |
| dif social closeness | Intergen | Alone-Together | RIFG~LTPJ | -0.0026 | -0.0079 | 0.0026 | |
| dif social closeness | Samegen | Alone-Together | RIFG~LTPJ | 0.0037 | -0.0025 | 0.0098 | |
| dif social closeness | Intergen | Alone-Together | RIFG~RTPJ | -0.0025 | -0.0097 | 0.0042 | |
| dif social closeness | Samegen | Alone-Together | RIFG~RTPJ | -0.0041 | -0.0117 | 0.0035 | |

**Supplementary Materials:** Moffat, Dumas and Cross (2026). *Social interactions between people of same and different generations shape longitudinal changes in interpersonal neural synchrony, loneliness, and social connection.*

**S5 Appendix – Table S.** HbO. Relationships between allophilia (sum) and INS in unstandardised units. Effects are presented in the following order: Main, group, task, and simple effects.

| Measure | Group | Task | ROI Pairs | Estimate | Lower HPD | Upper HPD | No overlap |
|---|---|---|---|---|---|---|---|
| *Main effect* | | | | | | | |
| sum allophilia | | | LIFG~RIFG | -0.0002 | -0.0003 | 0 | No overlap |
| sum allophilia | | | LIFG~LIFG | 0.0001 | -0.0001 | 0.0003 | |
| sum allophilia | | | LTPJ~LTPJ | -0.0004 | -0.0006 | -0.0001 | No overlap |
| sum allophilia | | | LIFG~LTPJ | -0.0001 | -0.0003 | 0.0001 | |
| sum allophilia | | | LIFG~RTPJ | 0 | -0.0002 | 0.0002 | |
| sum allophilia | | | RIFG~RIFG | 0 | -0.0002 | 0.0002 | |
| sum allophilia | | | RTPJ~RTPJ | 0 | -0.0004 | 0.0004 | |
| sum allophilia | | | RIFG~LTPJ | -0.0001 | -0.0002 | 0.0001 | |
| sum allophilia | | | RIFG~RTPJ | 0 | -0.0003 | 0.0002 | |
| *Group effect* | | | | | | | |
| sum allophilia | Intergen | | LIFG~RIFG | -0.0001 | -0.0003 | 0.0002 | |
| sum allophilia | Samegen | | LIFG~RIFG | -0.0003 | -0.0005 | -0.0001 | No overlap |
| sum allophilia | Intergen | | LIFG~LIFG | 0.0001 | -0.0002 | 0.0004 | |
| sum allophilia | Samegen | | LIFG~LIFG | 0.0001 | -0.0002 | 0.0004 | |
| sum allophilia | Intergen | | LTPJ~LTPJ | -0.0004 | -0.0008 | -0.0001 | No overlap |
| sum allophilia | Samegen | | LTPJ~LTPJ | -0.0003 | -0.0007 | 0 | |
| sum allophilia | Intergen | | LIFG~LTPJ | -0.0001 | -0.0004 | 0.0001 | |
| sum allophilia | Samegen | | LIFG~LTPJ | -0.0001 | -0.0003 | 0.0002 | |
| sum allophilia | Intergen | | LIFG~RTPJ | 0.0001 | -0.0002 | 0.0003 | |
| sum allophilia | Samegen | | LIFG~RTPJ | -0.0001 | -0.0004 | 0.0002 | |
| sum allophilia | Intergen | | RIFG~RIFG | 0 | -0.0003 | 0.0004 | |
| sum allophilia | Samegen | | RIFG~RIFG | -0.0001 | -0.0004 | 0.0003 | |
| sum allophilia | Intergen | | RTPJ~RTPJ | 0.0001 | -0.0005 | 0.0007 | |
| sum allophilia | Samegen | | RTPJ~RTPJ | -0.0001 | -0.0006 | 0.0004 | |
| sum allophilia | Intergen | | RIFG~LTPJ | -0.0001 | -0.0003 | 0.0001 | |
| sum allophilia | Samegen | | RIFG~LTPJ | -0.0001 | -0.0003 | 0.0001 | |
| sum allophilia | Intergen | | RIFG~RTPJ | 0.0002 | -0.0001 | 0.0005 | |
| sum allophilia | Samegen | | RIFG~RTPJ | -0.0003 | -0.0006 | 0 | |
| *Task effect* | | | | | | | |
| sum allophilia | | Alone | LIFG~RIFG | -0.0002 | -0.0004 | 0.0001 | |
| sum allophilia | | Together | LIFG~RIFG | -0.0002 | -0.0004 | 0 | |
| sum allophilia | | Alone | LIFG~LIFG | 0.0002 | -0.0001 | 0.0005 | |
| sum allophilia | | Together | LIFG~LIFG | 0.0001 | -0.0002 | 0.0003 | |
| sum allophilia | | Alone | LTPJ~LTPJ | -0.0004 | -0.0007 | -0.0001 | No overlap |
| sum allophilia | | Together | LTPJ~LTPJ | -0.0003 | -0.0006 | 0 | No overlap |
| sum allophilia | | Alone | LIFG~LTPJ | -0.0001 | -0.0003 | 0.0002 | |
| sum allophilia | | Together | LIFG~LTPJ | -0.0001 | -0.0003 | 0.0001 | |
| sum allophilia | | Alone | LIFG~RTPJ | -0.0001 | -0.0004 | 0.0002 | |
| sum allophilia | | Together | LIFG~RTPJ | 0.0001 | -0.0002 | 0.0003 | |
| sum allophilia | | Alone | RIFG~RIFG | -0.0002 | -0.0005 | 0.0002 | |
| sum allophilia | | Together | RIFG~RIFG | 0.0001 | -0.0001 | 0.0004 | |
| sum allophilia | | Alone | RTPJ~RTPJ | -0.0001 | -0.0007 | 0.0005 | |

| Measure | Group | Task | ROI Pairs | Estimate | Lower HPD | Upper HPD | No overlap |
|---|---|---|---|---|---|---|---|
| sum allophilia | | Together | RTPJ~RTPJ | 0.0001 | -0.0004 | 0.0006 | |
| sum allophilia | | Alone | RIFG~LTPJ | -0.0001 | -0.0003 | 0.0001 | |
| sum allophilia | | Together | RIFG~LTPJ | -0.0001 | -0.0003 | 0.0001 | |
| sum allophilia | | Alone | RIFG~RTPJ | -0.0001 | -0.0005 | 0.0002 | |
| sum allophilia | | Together | RIFG~RTPJ | 0.0001 | -0.0002 | 0.0003 | |
| *Simple effect* | | | | | | | |
| sum allophilia | Intergen | Alone | LIFG~RIFG | 0.0000 | -0.0003 | 0.0003 | |
| sum allophilia | Intergen | Together | LIFG~RIFG | -0.0001 | -0.0004 | 0.0002 | |
| sum allophilia | Samegen | Alone | LIFG~RIFG | -0.0003 | -0.0007 | 0.0000 | |
| sum allophilia | Samegen | Together | LIFG~RIFG | -0.0003 | -0.0005 | 0.0000 | |
| sum allophilia | Intergen | Alone | LIFG~LIFG | 0.0001 | -0.0004 | 0.0005 | |
| sum allophilia | Intergen | Together | LIFG~LIFG | 0.0001 | -0.0002 | 0.0005 | |
| sum allophilia | Samegen | Alone | LIFG~LIFG | 0.0003 | -0.0002 | 0.0007 | |
| sum allophilia | Samegen | Together | LIFG~LIFG | 0.0000 | -0.0004 | 0.0003 | |
| sum allophilia | Intergen | Alone | LTPJ~LTPJ | -0.0004 | -0.0009 | 0.0001 | |
| sum allophilia | Intergen | Together | LTPJ~LTPJ | -0.0004 | -0.0008 | 0.0000 | No overlap |
| sum allophilia | Samegen | Alone | LTPJ~LTPJ | -0.0004 | -0.0009 | 0.0001 | |
| sum allophilia | Samegen | Together | LTPJ~LTPJ | -0.0002 | -0.0007 | 0.0002 | |
| sum allophilia | Intergen | Alone | LIFG~LTPJ | 0.0001 | -0.0003 | 0.0004 | |
| sum allophilia | Intergen | Together | LIFG~LTPJ | -0.0003 | -0.0006 | 0.0000 | No overlap |
| sum allophilia | Samegen | Alone | LIFG~LTPJ | -0.0002 | -0.0006 | 0.0001 | |
| sum allophilia | Samegen | Together | LIFG~LTPJ | 0.0001 | -0.0002 | 0.0003 | |
| sum allophilia | Intergen | Alone | LIFG~RTPJ | 0.0000 | -0.0004 | 0.0004 | |
| sum allophilia | Intergen | Together | LIFG~RTPJ | 0.0002 | -0.0002 | 0.0005 | |
| sum allophilia | Samegen | Alone | LIFG~RTPJ | -0.0001 | -0.0006 | 0.0002 | |
| sum allophilia | Samegen | Together | LIFG~RTPJ | 0.0000 | -0.0003 | 0.0003 | |
| sum allophilia | Intergen | Alone | RIFG~RIFG | -0.0001 | -0.0006 | 0.0003 | |
| sum allophilia | Intergen | Together | RIFG~RIFG | 0.0002 | -0.0002 | 0.0006 | |
| sum allophilia | Samegen | Alone | RIFG~RIFG | -0.0002 | -0.0007 | 0.0003 | |
| sum allophilia | Samegen | Together | RIFG~RIFG | 0.0001 | -0.0003 | 0.0005 | |
| sum allophilia | Intergen | Alone | RTPJ~RTPJ | 0.0003 | -0.0006 | 0.0012 | |
| sum allophilia | Intergen | Together | RTPJ~RTPJ | -0.0001 | -0.0009 | 0.0007 | |
| sum allophilia | Samegen | Alone | RTPJ~RTPJ | -0.0005 | -0.0013 | 0.0002 | |
| sum allophilia | Samegen | Together | RTPJ~RTPJ | 0.0004 | -0.0003 | 0.0010 | |
| sum allophilia | Intergen | Alone | RIFG~LTPJ | -0.0001 | -0.0004 | 0.0002 | |
| sum allophilia | Intergen | Together | RIFG~LTPJ | -0.0001 | -0.0003 | 0.0002 | |
| sum allophilia | Samegen | Alone | RIFG~LTPJ | -0.0001 | -0.0004 | 0.0003 | |
| sum allophilia | Samegen | Together | RIFG~LTPJ | -0.0001 | -0.0003 | 0.0002 | |
| sum allophilia | Intergen | Alone | RIFG~RTPJ | 0.0001 | -0.0003 | 0.0005 | |
| sum allophilia | Intergen | Together | RIFG~RTPJ | 0.0003 | -0.0001 | 0.0006 | |
| sum allophilia | Samegen | Alone | RIFG~RTPJ | -0.0004 | -0.0008 | 0.0001 | |
| sum allophilia | Samegen | Together | RIFG~RTPJ | -0.0002 | -0.0005 | 0.0002 | |
| sum allophilia | Intergen | Alone | LIFG~RIFG | 0.0000 | -0.0003 | 0.0003 | |

**Supplementary Materials:** Moffat, Dumas and Cross (2026). *Social interactions between people of same and different generations shape longitudinal changes in interpersonal neural synchrony, loneliness, and social connection.*

**S5 Appendix – Table T.** HbO. Contrasts pertaining to allophilia (sum) ~ INS relationship in unstandardised units. Contrasts are presented in the following order: Group, task, and interaction contrasts.

| Measure | Group | Task | ROI Pairs | Estimate | Lower HPD | Upper HPD | No overlap |
|---|---|---|---|---|---|---|---|
| *Group contrast* | | | | | | | |
| sum allophilia | Int-Samegen | | LIFG~RIFG | 0.0002 | -0.0001 | 0.0006 | |
| sum allophilia | Int-Samegen | | LIFG~LIFG | 0.0000 | -0.0005 | 0.0004 | |
| sum allophilia | Int-Samegen | | LTPJ~LTPJ | -0.0001 | -0.0006 | 0.0004 | |
| sum allophilia | Int-Samegen | | LIFG~LTPJ | 0.0000 | -0.0004 | 0.0003 | |
| sum allophilia | Int-Samegen | | LIFG~RTPJ | 0.0002 | -0.0002 | 0.0005 | |
| sum allophilia | Int-Samegen | | RIFG~RIFG | 0.0001 | -0.0004 | 0.0006 | |
| sum allophilia | Int-Samegen | | RTPJ~RTPJ | 0.0002 | -0.0006 | 0.0010 | |
| sum allophilia | Int-Samegen | | RIFG~LTPJ | 0.0000 | -0.0003 | 0.0003 | |
| sum allophilia | Int-Samegen | | RIFG~RTPJ | 0.0005 | 0.0000 | 0.0009 | No overlap |
| *Task contrast* | | | | | | | |
| sum allophilia | | Alone-Together | LIFG~RIFG | 0.0000 | -0.0003 | 0.0003 | |
| sum allophilia | | Alone-Together | LIFG~LIFG | 0.0001 | -0.0003 | 0.0005 | |
| sum allophilia | | Alone-Together | LTPJ~LTPJ | -0.0001 | -0.0005 | 0.0003 | |
| sum allophilia | | Alone-Together | LIFG~LTPJ | 0.0000 | -0.0002 | 0.0003 | |
| sum allophilia | | Alone-Together | LIFG~RTPJ | -0.0001 | -0.0005 | 0.0002 | |
| sum allophilia | | Alone-Together | RIFG~RIFG | -0.0003 | -0.0007 | 0.0001 | |
| sum allophilia | | Alone-Together | RTPJ~RTPJ | -0.0003 | -0.0010 | 0.0005 | |
| sum allophilia | | Alone-Together | RIFG~LTPJ | 0.0000 | -0.0003 | 0.0002 | |
| sum allophilia | | Alone-Together | RIFG~RTPJ | -0.0002 | -0.0006 | 0.0002 | |
| *Interactions* | | | | | | | |
| sum allophilia | Int-Samegen | Alone | LIFG~RIFG | 0.0003 | -0.0001 | 0.0008 | |
| sum allophilia | Int-Samegen | Together | LIFG~RIFG | 0.0001 | -0.0002 | 0.0005 | |
| sum allophilia | Int-Samegen | Alone | LIFG~LIFG | -0.0002 | -0.0008 | 0.0004 | |
| sum allophilia | Int-Samegen | Together | LIFG~LIFG | 0.0001 | -0.0004 | 0.0006 | |
| sum allophilia | Int-Samegen | Alone | LTPJ~LTPJ | 0.0000 | -0.0007 | 0.0007 | |
| sum allophilia | Int-Samegen | Together | LTPJ~LTPJ | -0.0002 | -0.0007 | 0.0004 | |
| sum allophilia | Int-Samegen | Alone | LIFG~LTPJ | 0.0003 | -0.0002 | 0.0008 | |
| sum allophilia | Int-Samegen | Together | LIFG~LTPJ | -0.0004 | -0.0008 | 0.0000 | |
| sum allophilia | Int-Samegen | Alone | LIFG~RTPJ | 0.0001 | -0.0004 | 0.0007 | |
| sum allophilia | Int-Samegen | Together | LIFG~RTPJ | 0.0002 | -0.0003 | 0.0006 | |
| sum allophilia | Int-Samegen | Alone | RIFG~RIFG | 0.0001 | -0.0006 | 0.0007 | |
| sum allophilia | Int-Samegen | Together | RIFG~RIFG | 0.0001 | -0.0004 | 0.0007 | |
| sum allophilia | Int-Samegen | Alone | RTPJ~RTPJ | 0.0009 | -0.0003 | 0.0020 | |
| sum allophilia | Int-Samegen | Together | RTPJ~RTPJ | -0.0004 | -0.0014 | 0.0006 | |
| sum allophilia | Int-Samegen | Alone | RIFG~LTPJ | -0.0001 | -0.0005 | 0.0004 | |
| sum allophilia | Int-Samegen | Together | RIFG~LTPJ | 0.0000 | -0.0003 | 0.0004 | |
| sum allophilia | Int-Samegen | Alone | RIFG~RTPJ | 0.0005 | -0.0002 | 0.0011 | |
| sum allophilia | Int-Samegen | Together | RIFG~RTPJ | 0.0004 | -0.0001 | 0.0009 | |
| sum allophilia | Intergen | Alone-Together | LIFG~RIFG | 0.0001 | -0.0003 | 0.0005 | |
| sum allophilia | Samegen | Alone-Together | LIFG~RIFG | -0.0001 | -0.0005 | 0.0003 | |
| sum allophilia | Intergen | Alone-Together | LIFG~LIFG | -0.0001 | -0.0006 | 0.0005 | |
| sum allophilia | Samegen | Alone-Together | LIFG~LIFG | 0.0003 | -0.0003 | 0.0008 | |

| Measure | Group | Task | ROI Pairs | Estimate | Lower HPD | Upper HPD | No overlap |
|---|---|---|---|---|---|---|---|
| sum allophilia | Intergen | Alone-Together | LTPJ~LTPJ | 0.0000 | -0.0005 | 0.0005 | |
| sum allophilia | Samegen | Alone-Together | LTPJ~LTPJ | -0.0001 | -0.0007 | 0.0004 | |
| sum allophilia | Intergen | Alone-Together | LIFG~LTPJ | 0.0004 | 0.0000 | 0.0008 | No overlap |
| sum allophilia | Samegen | Alone-Together | LIFG~LTPJ | -0.0003 | -0.0007 | 0.0001 | |
| sum allophilia | Intergen | Alone-Together | LIFG~RTPJ | -0.0002 | -0.0006 | 0.0003 | |
| sum allophilia | Samegen | Alone-Together | LIFG~RTPJ | -0.0001 | -0.0006 | 0.0004 | |
| sum allophilia | Intergen | Alone-Together | RIFG~RIFG | -0.0003 | -0.0009 | 0.0002 | |
| sum allophilia | Samegen | Alone-Together | RIFG~RIFG | -0.0003 | -0.0009 | 0.0004 | |
| sum allophilia | Intergen | Alone-Together | RTPJ~RTPJ | 0.0004 | -0.0008 | 0.0015 | |
| sum allophilia | Samegen | Alone-Together | RTPJ~RTPJ | -0.0009 | -0.0018 | 0.0000 | |
| sum allophilia | Intergen | Alone-Together | RIFG~LTPJ | -0.0001 | -0.0005 | 0.0003 | |
| sum allophilia | Samegen | Alone-Together | RIFG~LTPJ | 0.0000 | -0.0004 | 0.0004 | |
| sum allophilia | Intergen | Alone-Together | RIFG~RTPJ | -0.0002 | -0.0007 | 0.0003 | |
| sum allophilia | Samegen | Alone-Together | RIFG~RTPJ | -0.0002 | -0.0007 | 0.0003 | |

**S5 Appendix – Table U.** HbO. Relationships between allophilia (dif) and INS in unstandardised units. Effects are presented in the following order: Main, group, task, and simple effects.

| Measure | Group | Task | ROI Pairs | Estimate | Lower HPD | Upper HPD | No overlap |
|---|---|---|---|---|---|---|---|
| *Main effect* | | | | | | | |
| dif allophilia | | | LIFG~RIFG | 0.0001 | -0.0002 | 0.0003 | |
| dif allophilia | | | LIFG~LIFG | 0 | -0.0002 | 0.0003 | |
| dif allophilia | | | LTPJ~LTPJ | 0.0002 | -0.0002 | 0.0005 | |
| dif allophilia | | | LIFG~LTPJ | -0.0001 | -0.0004 | 0.0001 | |
| dif allophilia | | | LIFG~RTPJ | -0.0001 | -0.0004 | 0.0001 | |
| dif allophilia | | | RIFG~RIFG | 0.0001 | -0.0002 | 0.0004 | |
| dif allophilia | | | RTPJ~RTPJ | 0.0001 | -0.0003 | 0.0006 | |
| dif allophilia | | | RIFG~LTPJ | -0.0001 | -0.0003 | 0.0001 | |
| dif allophilia | | | RIFG~RTPJ | 0 | -0.0003 | 0.0003 | |
| *Group effect* | | | | | | | |
| dif allophilia | Intergen | | LIFG~RIFG | 0.0002 | -0.0001 | 0.0005 | |
| dif allophilia | Samegen | | LIFG~RIFG | -0.0001 | -0.0004 | 0.0003 | |
| dif allophilia | Intergen | | LIFG~LIFG | 0.0001 | -0.0003 | 0.0004 | |
| dif allophilia | Samegen | | LIFG~LIFG | 0 | -0.0004 | 0.0004 | |
| dif allophilia | Intergen | | LTPJ~LTPJ | 0.0002 | -0.0002 | 0.0006 | |
| dif allophilia | Samegen | | LTPJ~LTPJ | 0.0001 | -0.0003 | 0.0007 | |
| dif allophilia | Intergen | | LIFG~LTPJ | -0.0002 | -0.0005 | 0.0001 | |
| dif allophilia | Samegen | | LIFG~LTPJ | -0.0001 | -0.0004 | 0.0002 | |
| dif allophilia | Intergen | | LIFG~RTPJ | 0 | -0.0003 | 0.0003 | |
| dif allophilia | Samegen | | LIFG~RTPJ | -0.0002 | -0.0006 | 0.0001 | |
| dif allophilia | Intergen | | RIFG~RIFG | 0.0002 | -0.0003 | 0.0006 | |
| dif allophilia | Samegen | | RIFG~RIFG | 0.0001 | -0.0004 | 0.0005 | |
| dif allophilia | Intergen | | RTPJ~RTPJ | 0.0003 | -0.0003 | 0.0009 | |
| dif allophilia | Samegen | | RTPJ~RTPJ | -0.0001 | -0.0007 | 0.0006 | |
| dif allophilia | Intergen | | RIFG~LTPJ | -0.0003 | -0.0005 | 0 | |
| dif allophilia | Samegen | | RIFG~LTPJ | 0 | -0.0003 | 0.0003 | |
| dif allophilia | Intergen | | RIFG~RTPJ | 0.0001 | -0.0003 | 0.0004 | |
| dif allophilia | Samegen | | RIFG~RTPJ | -0.0001 | -0.0005 | 0.0003 | |
| *Task effect* | | | | | | | |
| dif allophilia | | Alone | LIFG~RIFG | 0 | -0.0003 | 0.0003 | |
| dif allophilia | | Together | LIFG~RIFG | 0.0001 | -0.0001 | 0.0004 | |
| dif allophilia | | Alone | LIFG~LIFG | 0.0002 | -0.0002 | 0.0006 | |
| dif allophilia | | Together | LIFG~LIFG | -0.0001 | -0.0004 | 0.0003 | |
| dif allophilia | | Alone | LTPJ~LTPJ | 0.0001 | -0.0003 | 0.0006 | |
| dif allophilia | | Together | LTPJ~LTPJ | 0.0002 | -0.0002 | 0.0006 | |
| dif allophilia | | Alone | LIFG~LTPJ | -0.0001 | -0.0004 | 0.0002 | |
| dif allophilia | | Together | LIFG~LTPJ | -0.0001 | -0.0004 | 0.0001 | |
| dif allophilia | | Alone | LIFG~RTPJ | -0.0002 | -0.0006 | 0.0001 | |
| dif allophilia | | Together | LIFG~RTPJ | 0 | -0.0003 | 0.0003 | |
| dif allophilia | | Alone | RIFG~RIFG | -0.0003 | -0.0007 | 0.0002 | |
| dif allophilia | | Together | RIFG~RIFG | 0.0005 | 0.0002 | 0.0009 | No overlap |
| dif allophilia | | Alone | RTPJ~RTPJ | 0.0004 | -0.0003 | 0.0011 | |

| Measure | Group | Task | ROI Pairs | Estimate | Lower HPD | Upper HPD | No overlap |
|---|---|---|---|---|---|---|---|
| dif allophilia | | Together | RTPJ~RTPJ | -0.0001 | -0.0007 | 0.0005 | |
| dif allophilia | | Alone | RIFG~LTPJ | -0.0002 | -0.0005 | 0.0001 | |
| dif allophilia | | Together | RIFG~LTPJ | -0.0001 | -0.0004 | 0.0001 | |
| dif allophilia | | Alone | RIFG~RTPJ | -0.0001 | -0.0004 | 0.0003 | |
| dif allophilia | | Together | RIFG~RTPJ | 0 | -0.0003 | 0.0003 | |
| *Simple effect* | | | | | | | |
| dif allophilia | Intergen | Alone | LIFG~RIFG | 0.0002 | -0.0002 | 0.0006 | |
| dif allophilia | Intergen | Together | LIFG~RIFG | 0.0001 | -0.0002 | 0.0005 | |
| dif allophilia | Samegen | Alone | LIFG~RIFG | -0.0002 | -0.0007 | 0.0002 | |
| dif allophilia | Samegen | Together | LIFG~RIFG | 0.0001 | -0.0002 | 0.0005 | |
| dif allophilia | Intergen | Alone | LIFG~LIFG | 0.0002 | -0.0004 | 0.0007 | |
| dif allophilia | Intergen | Together | LIFG~LIFG | -0.0001 | -0.0006 | 0.0004 | |
| dif allophilia | Samegen | Alone | LIFG~LIFG | 0.0002 | -0.0004 | 0.0008 | |
| dif allophilia | Samegen | Together | LIFG~LIFG | -0.0001 | -0.0006 | 0.0004 | |
| dif allophilia | Intergen | Alone | LTPJ~LTPJ | 0.0002 | -0.0004 | 0.0008 | |
| dif allophilia | Intergen | Together | LTPJ~LTPJ | 0.0002 | -0.0004 | 0.0007 | |
| dif allophilia | Samegen | Alone | LTPJ~LTPJ | 0.0000 | -0.0007 | 0.0007 | |
| dif allophilia | Samegen | Together | LTPJ~LTPJ | 0.0003 | -0.0003 | 0.0009 | |
| dif allophilia | Intergen | Alone | LIFG~LTPJ | -0.0002 | -0.0006 | 0.0002 | |
| dif allophilia | Intergen | Together | LIFG~LTPJ | -0.0002 | -0.0005 | 0.0002 | |
| dif allophilia | Samegen | Alone | LIFG~LTPJ | -0.0001 | -0.0005 | 0.0004 | |
| dif allophilia | Samegen | Together | LIFG~LTPJ | -0.0001 | -0.0005 | 0.0002 | |
| dif allophilia | Intergen | Alone | LIFG~RTPJ | 0.0000 | -0.0005 | 0.0004 | |
| dif allophilia | Intergen | Together | LIFG~RTPJ | 0.0001 | -0.0003 | 0.0005 | |
| dif allophilia | Samegen | Alone | LIFG~RTPJ | -0.0004 | -0.0009 | 0.0001 | |
| dif allophilia | Samegen | Together | LIFG~RTPJ | 0.0000 | -0.0005 | 0.0004 | |
| dif allophilia | Intergen | Alone | RIFG~RIFG | -0.0002 | -0.0008 | 0.0004 | |
| dif allophilia | Intergen | Together | RIFG~RIFG | 0.0005 | 0.0000 | 0.0010 | |
| dif allophilia | Samegen | Alone | RIFG~RIFG | -0.0003 | -0.0010 | 0.0003 | |
| dif allophilia | Samegen | Together | RIFG~RIFG | 0.0005 | 0.0000 | 0.0011 | |
| dif allophilia | Intergen | Alone | RTPJ~RTPJ | 0.0009 | -0.0001 | 0.0018 | |
| dif allophilia | Intergen | Together | RTPJ~RTPJ | -0.0003 | -0.0011 | 0.0006 | |
| dif allophilia | Samegen | Alone | RTPJ~RTPJ | -0.0001 | -0.0011 | 0.0009 | |
| dif allophilia | Samegen | Together | RTPJ~RTPJ | 0.0000 | -0.0008 | 0.0008 | |
| dif allophilia | Intergen | Alone | RIFG~LTPJ | -0.0004 | -0.0008 | 0.0000 | |
| dif allophilia | Intergen | Together | RIFG~LTPJ | -0.0001 | -0.0005 | 0.0002 | |
| dif allophilia | Samegen | Alone | RIFG~LTPJ | 0.0000 | -0.0004 | 0.0005 | |
| dif allophilia | Samegen | Together | RIFG~LTPJ | -0.0001 | -0.0004 | 0.0003 | |
| dif allophilia | Intergen | Alone | RIFG~RTPJ | 0.0000 | -0.0005 | 0.0005 | |
| dif allophilia | Intergen | Together | RIFG~RTPJ | 0.0002 | -0.0003 | 0.0006 | |
| dif allophilia | Samegen | Alone | RIFG~RTPJ | -0.0001 | -0.0007 | 0.0005 | |
| dif allophilia | Samegen | Together | RIFG~RTPJ | -0.0001 | -0.0006 | 0.0003 | |
| dif allophilia | Intergen | Alone | LIFG~RIFG | 0.0002 | -0.0002 | 0.0006 | |

**S5 Appendix – Table V.** HbO. Contrasts pertaining to allophilia (dif) ~ INS relationship in unstandardised units. Contrasts are presented in the following order: Group, task, and interaction contrasts.

| Measure | Group | Task | ROI Pairs | Estimate | Lower HPD | Upper HPD | No overlap |
|---|---|---|---|---|---|---|---|
| *Group contrast* | | | | | | | |
| dif allophilia | Int-Samegen | | LIFG~RIFG | 0.0002 | -0.0002 | 0.0007 | |
| dif allophilia | Int-Samegen | | LIFG~LIFG | 0.0000 | -0.0005 | 0.0006 | |
| dif allophilia | Int-Samegen | | LTPJ~LTPJ | 0.0000 | -0.0006 | 0.0007 | |
| dif allophilia | Int-Samegen | | LIFG~LTPJ | -0.0001 | -0.0005 | 0.0004 | |
| dif allophilia | Int-Samegen | | LIFG~RTPJ | 0.0002 | -0.0003 | 0.0007 | |
| dif allophilia | Int-Samegen | | RIFG~RIFG | 0.0001 | -0.0006 | 0.0007 | |
| dif allophilia | Int-Samegen | | RTPJ~RTPJ | 0.0004 | -0.0006 | 0.0013 | |
| dif allophilia | Int-Samegen | | RIFG~LTPJ | -0.0002 | -0.0006 | 0.0002 | |
| dif allophilia | Int-Samegen | | RIFG~RTPJ | 0.0002 | -0.0003 | 0.0008 | |
| *Task contrast* | | | | | | | |
| dif allophilia | | Alone-Together | LIFG~RIFG | -0.0002 | -0.0006 | 0.0002 | |
| dif allophilia | | Alone-Together | LIFG~LIFG | 0.0003 | -0.0002 | 0.0008 | |
| dif allophilia | | Alone-Together | LTPJ~LTPJ | -0.0001 | -0.0007 | 0.0004 | |
| dif allophilia | | Alone-Together | LIFG~LTPJ | 0.0000 | -0.0004 | 0.0004 | |
| dif allophilia | | Alone-Together | LIFG~RTPJ | -0.0002 | -0.0007 | 0.0002 | |
| dif allophilia | | Alone-Together | RIFG~RIFG | -0.0008 | -0.0013 | -0.0002 | No overlap |
| dif allophilia | | Alone-Together | RTPJ~RTPJ | 0.0005 | -0.0003 | 0.0014 | |
| dif allophilia | | Alone-Together | RIFG~LTPJ | -0.0001 | -0.0004 | 0.0003 | |
| dif allophilia | | Alone-Together | RIFG~RTPJ | -0.0001 | -0.0005 | 0.0004 | |
| *Interactions* | | | | | | | |
| dif allophilia | Int-Samegen | Alone | LIFG~RIFG | 0.0004 | -0.0002 | 0.0011 | |
| dif allophilia | Int-Samegen | Together | LIFG~RIFG | 0.0000 | -0.0005 | 0.0005 | |
| dif allophilia | Int-Samegen | Alone | LIFG~LIFG | 0.0000 | -0.0008 | 0.0008 | |
| dif allophilia | Int-Samegen | Together | LIFG~LIFG | 0.0000 | -0.0006 | 0.0007 | |
| dif allophilia | Int-Samegen | Alone | LTPJ~LTPJ | 0.0002 | -0.0007 | 0.0011 | |
| dif allophilia | Int-Samegen | Together | LTPJ~LTPJ | -0.0001 | -0.0009 | 0.0007 | |
| dif allophilia | Int-Samegen | Alone | LIFG~LTPJ | -0.0001 | -0.0007 | 0.0005 | |
| dif allophilia | Int-Samegen | Together | LIFG~LTPJ | 0.0000 | -0.0005 | 0.0005 | |
| dif allophilia | Int-Samegen | Alone | LIFG~RTPJ | 0.0004 | -0.0004 | 0.0011 | |
| dif allophilia | Int-Samegen | Together | LIFG~RTPJ | 0.0001 | -0.0005 | 0.0007 | |
| dif allophilia | Int-Samegen | Alone | RIFG~RIFG | 0.0002 | -0.0007 | 0.0011 | |
| dif allophilia | Int-Samegen | Together | RIFG~RIFG | 0.0000 | -0.0008 | 0.0007 | |
| dif allophilia | Int-Samegen | Alone | RTPJ~RTPJ | 0.0009 | -0.0004 | 0.0023 | |
| dif allophilia | Int-Samegen | Together | RTPJ~RTPJ | -0.0002 | -0.0014 | 0.0010 | |
| dif allophilia | Int-Samegen | Alone | RIFG~LTPJ | -0.0004 | -0.0010 | 0.0002 | |
| dif allophilia | Int-Samegen | Together | RIFG~LTPJ | -0.0001 | -0.0006 | 0.0004 | |
| dif allophilia | Int-Samegen | Alone | RIFG~RTPJ | 0.0001 | -0.0007 | 0.0009 | |
| dif allophilia | Int-Samegen | Together | RIFG~RTPJ | 0.0003 | -0.0003 | 0.0010 | |
| dif allophilia | Intergen | Alone-Together | LIFG~RIFG | 0.0001 | -0.0005 | 0.0006 | |
| dif allophilia | Samegen | Alone-Together | LIFG~RIFG | -0.0004 | -0.0009 | 0.0002 | |
| dif allophilia | Intergen | Alone-Together | LIFG~LIFG | 0.0003 | -0.0004 | 0.0009 | |
| dif allophilia | Samegen | Alone-Together | LIFG~LIFG | 0.0003 | -0.0005 | 0.0010 | |

| Measure | Group | Task | ROI Pairs | Estimate | Lower HPD | Upper HPD | No overlap |
|---|---|---|---|---|---|---|---|
| sum allophilia | Intergen | Alone-Together | LTPJ~LTPJ | 0.0001 | -0.0007 | 0.0008 | |
| sum allophilia | Samegen | Alone-Together | LTPJ~LTPJ | -0.0003 | -0.0011 | 0.0005 | |
| sum allophilia | Intergen | Alone-Together | LIFG~LTPJ | 0.0000 | -0.0005 | 0.0005 | |
| sum allophilia | Samegen | Alone-Together | LIFG~LTPJ | 0.0000 | -0.0005 | 0.0006 | |
| sum allophilia | Intergen | Alone-Together | LIFG~RTPJ | -0.0001 | -0.0007 | 0.0005 | |
| sum allophilia | Samegen | Alone-Together | LIFG~RTPJ | -0.0003 | -0.0010 | 0.0003 | |
| sum allophilia | Intergen | Alone-Together | RIFG~RIFG | -0.0007 | -0.0015 | 0.0001 | |
| sum allophilia | Samegen | Alone-Together | RIFG~RIFG | -0.0009 | -0.0017 | -0.0001 | No overlap |
| sum allophilia | Intergen | Alone-Together | RTPJ~RTPJ | 0.0011 | -0.0002 | 0.0023 | |
| sum allophilia | Samegen | Alone-Together | RTPJ~RTPJ | 0.0000 | -0.0013 | 0.0012 | |
| sum allophilia | Intergen | Alone-Together | RIFG~LTPJ | -0.0002 | -0.0007 | 0.0003 | |
| sum allophilia | Samegen | Alone-Together | RIFG~LTPJ | 0.0001 | -0.0004 | 0.0007 | |
| sum allophilia | Intergen | Alone-Together | RIFG~RTPJ | -0.0002 | -0.0008 | 0.0004 | |
| sum allophilia | Samegen | Alone-Together | RIFG~RTPJ | 0.0000 | -0.0007 | 0.0007 | |

**Supplementary Materials:** Moffat, Dumas and Cross (2026). *Social interactions between people of same and different generations shape longitudinal changes in interpersonal neural synchrony, loneliness, and social connection.*

**S6 Appendix – Table A.** HbR. Estimates of INS levels for all sessions combined for real and pseudo dyads per ROI pair and task in unstandardised units.

| Dyad Type | Group | ROI Pair | Task | Estimate | Lower HPD | Upper HPD |
|---|---|---|---|---|---|---|
| Pseudo | Intergen | LIFG~RIFG | Alone | 0.238 | 0.238 | 0.239 |
| Real | Intergen | LIFG~RIFG | Alone | 0.233 | 0.228 | 0.238 |
| Pseudo | Intergen | LIFG~RIFG | Together | 0.239 | 0.238 | 0.239 |
| Real | Intergen | LIFG~RIFG | Together | 0.244 | 0.241 | 0.248 |
| Pseudo | Intergen | LTPJ~RTPJ | Alone | 0.236 | 0.236 | 0.237 |
| Real | Intergen | LTPJ~RTPJ | Alone | 0.238 | 0.231 | 0.243 |
| Pseudo | Intergen | LTPJ~RTPJ | Together | 0.237 | 0.236 | 0.237 |
| Real | Intergen | LTPJ~RTPJ | Together | 0.238 | 0.233 | 0.242 |
| Pseudo | Intergen | LIFG~LIFG | Alone | 0.24 | 0.239 | 0.241 |
| Real | Intergen | LIFG~LIFG | Alone | 0.237 | 0.231 | 0.244 |
| Pseudo | Intergen | LIFG~LIFG | Together | 0.24 | 0.24 | 0.241 |
| Real | Intergen | LIFG~LIFG | Together | 0.241 | 0.236 | 0.246 |
| Pseudo | Intergen | LTPJ~LTPJ | Alone | 0.237 | 0.237 | 0.238 |
| Real | Intergen | LTPJ~LTPJ | Alone | 0.241 | 0.234 | 0.247 |
| Pseudo | Intergen | LTPJ~LTPJ | Together | 0.237 | 0.237 | 0.238 |
| Real | Intergen | LTPJ~LTPJ | Together | 0.244 | 0.239 | 0.249 |
| Pseudo | Intergen | LIFG~LTPJ | Alone | 0.238 | 0.237 | 0.238 |
| Real | Intergen | LIFG~LTPJ | Alone | 0.24 | 0.236 | 0.245 |
| Pseudo | Intergen | LIFG~LTPJ | Together | 0.239 | 0.238 | 0.239 |
| Real | Intergen | LIFG~LTPJ | Together | 0.244 | 0.24 | 0.248 |
| Pseudo | Intergen | LIFG~RTPJ | Alone | 0.237 | 0.236 | 0.237 |
| Real | Intergen | LIFG~RTPJ | Alone | 0.235 | 0.229 | 0.241 |
| Pseudo | Intergen | LIFG~RTPJ | Together | 0.238 | 0.237 | 0.238 |
| Real | Intergen | LIFG~RTPJ | Together | 0.24 | 0.236 | 0.245 |
| Pseudo | Intergen | RIFG~RIFG | Alone | 0.237 | 0.236 | 0.238 |
| Real | Intergen | RIFG~RIFG | Alone | 0.241 | 0.233 | 0.247 |
| Pseudo | Intergen | RIFG~RIFG | Together | 0.237 | 0.237 | 0.238 |
| Real | Intergen | RIFG~RIFG | Together | 0.238 | 0.233 | 0.243 |
| Pseudo | Intergen | RTPJ~RTPJ | Alone | 0.235 | 0.233 | 0.236 |
| Real | Intergen | RTPJ~RTPJ | Alone | 0.233 | 0.221 | 0.245 |
| Pseudo | Intergen | RTPJ~RTPJ | Together | 0.236 | 0.235 | 0.237 |
| Real | Intergen | RTPJ~RTPJ | Together | 0.235 | 0.225 | 0.244 |
| Pseudo | Intergen | LTPJ~RIFG | Alone | 0.237 | 0.236 | 0.237 |
| Real | Intergen | LTPJ~RIFG | Alone | 0.236 | 0.231 | 0.24 |
| Pseudo | Intergen | LTPJ~RIFG | Together | 0.237 | 0.237 | 0.237 |
| Real | Intergen | LTPJ~RIFG | Together | 0.241 | 0.237 | 0.244 |
| Pseudo | Intergen | RIFG~RTPJ | Alone | 0.235 | 0.235 | 0.236 |
| Real | Intergen | RIFG~RTPJ | Alone | 0.23 | 0.224 | 0.236 |
| Pseudo | Intergen | RIFG~RTPJ | Together | 0.236 | 0.236 | 0.237 |
| Real | Intergen | RIFG~RTPJ | Together | 0.239 | 0.234 | 0.244 |
| Pseudo | Samegen | LIFG~RIFG | Alone | 0.24 | 0.24 | 0.24 |
| Real | Samegen | LIFG~RIFG | Alone | 0.237 | 0.232 | 0.241 |
| Pseudo | Samegen | LIFG~RIFG | Together | 0.24 | 0.24 | 0.241 |

| Dyad Type | Group | ROI Pair | Task | Estimate | Lower HPD | Upper HPD |
|---|---|---|---|---|---|---|
| Real | Samegen | LIFG~RIFG | Together | 0.24 | 0.236 | 0.244 |
| Pseudo | Samegen | LTPJ~RTPJ | Alone | 0.242 | 0.241 | 0.242 |
| Real | Samegen | LTPJ~RTPJ | Alone | 0.244 | 0.239 | 0.25 |
| Pseudo | Samegen | LTPJ~RTPJ | Together | 0.241 | 0.241 | 0.242 |
| Real | Samegen | LTPJ~RTPJ | Together | 0.244 | 0.24 | 0.249 |
| Pseudo | Samegen | LIFG~LIFG | Alone | 0.241 | 0.24 | 0.241 |
| Real | Samegen | LIFG~LIFG | Alone | 0.245 | 0.238 | 0.252 |
| Pseudo | Samegen | LIFG~LIFG | Together | 0.241 | 0.241 | 0.242 |
| Real | Samegen | LIFG~LIFG | Together | 0.244 | 0.239 | 0.249 |
| Pseudo | Samegen | LTPJ~LTPJ | Alone | 0.241 | 0.24 | 0.242 |
| Real | Samegen | LTPJ~LTPJ | Alone | 0.243 | 0.237 | 0.25 |
| Pseudo | Samegen | LTPJ~LTPJ | Together | 0.242 | 0.242 | 0.243 |
| Real | Samegen | LTPJ~LTPJ | Together | 0.245 | 0.24 | 0.25 |
| Pseudo | Samegen | LIFG~LTPJ | Alone | 0.24 | 0.24 | 0.241 |
| Real | Samegen | LIFG~LTPJ | Alone | 0.241 | 0.236 | 0.246 |
| Pseudo | Samegen | LIFG~LTPJ | Together | 0.241 | 0.24 | 0.241 |
| Real | Samegen | LIFG~LTPJ | Together | 0.241 | 0.238 | 0.245 |
| Pseudo | Samegen | LIFG~RTPJ | Alone | 0.24 | 0.24 | 0.241 |
| Real | Samegen | LIFG~RTPJ | Alone | 0.243 | 0.238 | 0.249 |
| Pseudo | Samegen | LIFG~RTPJ | Together | 0.24 | 0.24 | 0.24 |
| Real | Samegen | LIFG~RTPJ | Together | 0.239 | 0.235 | 0.244 |
| Pseudo | Samegen | RIFG~RIFG | Alone | 0.239 | 0.238 | 0.239 |
| Real | Samegen | RIFG~RIFG | Alone | 0.237 | 0.23 | 0.244 |
| Pseudo | Samegen | RIFG~RIFG | Together | 0.24 | 0.239 | 0.24 |
| Real | Samegen | RIFG~RIFG | Together | 0.246 | 0.24 | 0.251 |
| Pseudo | Samegen | RTPJ~RTPJ | Alone | 0.242 | 0.241 | 0.243 |
| Real | Samegen | RTPJ~RTPJ | Alone | 0.248 | 0.238 | 0.259 |
| Pseudo | Samegen | RTPJ~RTPJ | Together | 0.24 | 0.24 | 0.241 |
| Real | Samegen | RTPJ~RTPJ | Together | 0.244 | 0.236 | 0.252 |
| Pseudo | Samegen | LTPJ~RIFG | Alone | 0.239 | 0.238 | 0.239 |
| Real | Samegen | LTPJ~RIFG | Alone | 0.24 | 0.235 | 0.244 |
| Pseudo | Samegen | LTPJ~RIFG | Together | 0.24 | 0.24 | 0.24 |
| Real | Samegen | LTPJ~RIFG | Together | 0.244 | 0.24 | 0.247 |
| Pseudo | Samegen | RIFG~RTPJ | Alone | 0.239 | 0.239 | 0.24 |
| Real | Samegen | RIFG~RTPJ | Alone | 0.239 | 0.233 | 0.245 |
| Pseudo | Samegen | RIFG~RTPJ | Together | 0.239 | 0.239 | 0.24 |
| Real | Samegen | RIFG~RTPJ | Together | 0.242 | 0.237 | 0.246 |

**S6 Appendix – Table B.** HbR. Contrasts between real and pseudo dyads' INS levels for all sessions combined per ROI pair and task in unstandardised units.

| Contrast | Group | ROI Pair | Task | Estimate | Lower HPD | Upper HPD | No overlap |
|---|---|---|---|---|---|---|---|
| Real - Pseudo | Intergen | LIFG~RIFG | Alone | 0.005 | 0 | 0.009 | |
| Real - Pseudo | Intergen | LIFG~RIFG | Together | -0.006 | -0.009 | -0.002 | No overlap |
| Real - Pseudo | Intergen | LTPJ~RTPJ | Alone | -0.001 | -0.007 | 0.005 | |
| Real - Pseudo | Intergen | LTPJ~RTPJ | Together | -0.001 | -0.006 | 0.004 | |
| Real - Pseudo | Intergen | LIFG~LIFG | Alone | 0.003 | -0.004 | 0.01 | |
| Real - Pseudo | Intergen | LIFG~LIFG | Together | -0.001 | -0.006 | 0.005 | |
| Real - Pseudo | Intergen | LTPJ~LTPJ | Alone | -0.003 | -0.01 | 0.003 | |
| Real - Pseudo | Intergen | LTPJ~LTPJ | Together | -0.007 | -0.012 | -0.002 | No overlap |
| Real - Pseudo | Intergen | LIFG~LTPJ | Alone | -0.003 | -0.007 | 0.002 | |
| Real - Pseudo | Intergen | LIFG~LTPJ | Together | -0.005 | -0.009 | -0.002 | No overlap |
| Real - Pseudo | Intergen | LIFG~RTPJ | Alone | 0.001 | -0.004 | 0.007 | |
| Real - Pseudo | Intergen | LIFG~RTPJ | Together | -0.002 | -0.007 | 0.002 | |
| Real - Pseudo | Intergen | RIFG~RIFG | Alone | -0.004 | -0.011 | 0.003 | |
| Real - Pseudo | Intergen | RIFG~RIFG | Together | -0.001 | -0.006 | 0.004 | |
| Real - Pseudo | Intergen | RTPJ~RTPJ | Alone | 0.002 | -0.011 | 0.014 | |
| Real - Pseudo | Intergen | RTPJ~RTPJ | Together | 0.001 | -0.009 | 0.011 | |
| Real - Pseudo | Intergen | LTPJ~RIFG | Alone | 0.001 | -0.004 | 0.006 | |
| Real - Pseudo | Intergen | LTPJ~RIFG | Together | -0.004 | -0.007 | 0 | |
| Real - Pseudo | Intergen | RIFG~RTPJ | Alone | 0.005 | -0.001 | 0.011 | |
| Real - Pseudo | Intergen | RIFG~RTPJ | Together | -0.003 | -0.007 | 0.002 | |
| Real - Pseudo | Samegen | LIFG~RIFG | Alone | 0.003 | -0.002 | 0.008 | |
| Real - Pseudo | Samegen | LIFG~RIFG | Together | 0 | -0.003 | 0.004 | |
| Real - Pseudo | Samegen | LTPJ~RTPJ | Alone | -0.003 | -0.008 | 0.003 | |
| Real - Pseudo | Samegen | LTPJ~RTPJ | Together | -0.003 | -0.008 | 0.001 | |
| Real - Pseudo | Samegen | LIFG~LIFG | Alone | -0.004 | -0.011 | 0.003 | |
| Real - Pseudo | Samegen | LIFG~LIFG | Together | -0.003 | -0.008 | 0.003 | |
| Real - Pseudo | Samegen | LTPJ~LTPJ | Alone | -0.002 | -0.009 | 0.004 | |
| Real - Pseudo | Samegen | LTPJ~LTPJ | Together | -0.003 | -0.008 | 0.002 | |
| Real - Pseudo | Samegen | LIFG~LTPJ | Alone | -0.001 | -0.006 | 0.004 | |
| Real - Pseudo | Samegen | LIFG~LTPJ | Together | 0 | -0.004 | 0.003 | |
| Real - Pseudo | Samegen | LIFG~RTPJ | Alone | -0.003 | -0.008 | 0.003 | |
| Real - Pseudo | Samegen | LIFG~RTPJ | Together | 0.001 | -0.004 | 0.005 | |
| Real - Pseudo | Samegen | RIFG~RIFG | Alone | 0.002 | -0.005 | 0.009 | |
| Real - Pseudo | Samegen | RIFG~RIFG | Together | -0.006 | -0.011 | 0 | No overlap |
| Real - Pseudo | Samegen | RTPJ~RTPJ | Alone | -0.006 | -0.017 | 0.004 | |
| Real - Pseudo | Samegen | RTPJ~RTPJ | Together | -0.003 | -0.012 | 0.005 | |
| Real - Pseudo | Samegen | LTPJ~RIFG | Alone | -0.001 | -0.006 | 0.004 | |
| Real - Pseudo | Samegen | LTPJ~RIFG | Together | -0.004 | -0.008 | 0 | No overlap |
| Real - Pseudo | Samegen | RIFG~RTPJ | Alone | 0 | -0.005 | 0.006 | |
| Real - Pseudo | Samegen | RIFG~RTPJ | Together | -0.002 | -0.007 | 0.002 | |

**S6 Appendix – Table C.** HbR. Estimates of slope of change in INS across sessions for real and pseudo dyads per ROI pair and task in unstandardised units.

| Dyad Type | Group | ROI Pair | Task | Slope Estimate | Lower HPD | Upper HPD | No overlap |
|---|---|---|---|---|---|---|---|
| Pseudo | Intergen | LIFG~RIFG | Alone | 0.001 | 0 | 0.001 | No overlap |
| Real | Intergen | LIFG~RIFG | Alone | -0.001 | -0.003 | 0.002 | |
| Pseudo | Intergen | LIFG~RIFG | Together | 0.001 | 0 | 0.001 | No overlap |
| Real | Intergen | LIFG~RIFG | Together | 0 | -0.003 | 0.002 | |
| Pseudo | Intergen | LTPJ~RTPJ | Alone | 0 | 0 | 0.001 | No overlap |
| Real | Intergen | LTPJ~RTPJ | Alone | 0.002 | -0.002 | 0.005 | |
| Pseudo | Intergen | LTPJ~RTPJ | Together | 0 | -0.001 | 0 | No overlap |
| Real | Intergen | LTPJ~RTPJ | Together | -0.002 | -0.005 | 0.001 | |
| Pseudo | Intergen | LIFG~LIFG | Alone | 0 | 0 | 0.001 | |
| Real | Intergen | LIFG~LIFG | Alone | 0 | -0.004 | 0.004 | |
| Pseudo | Intergen | LIFG~LIFG | Together | 0 | 0 | 0.001 | No overlap |
| Real | Intergen | LIFG~LIFG | Together | -0.004 | -0.007 | 0 | No overlap |
| Pseudo | Intergen | LTPJ~LTPJ | Alone | 0.001 | 0 | 0.001 | No overlap |
| Real | Intergen | LTPJ~LTPJ | Alone | -0.001 | -0.005 | 0.003 | |
| Pseudo | Intergen | LTPJ~LTPJ | Together | 0 | -0.001 | 0 | |
| Real | Intergen | LTPJ~LTPJ | Together | 0.001 | -0.003 | 0.004 | |
| Pseudo | Intergen | LIFG~LTPJ | Alone | 0 | 0 | 0.001 | |
| Real | Intergen | LIFG~LTPJ | Alone | 0 | -0.003 | 0.003 | |
| Pseudo | Intergen | LIFG~LTPJ | Together | 0 | 0 | 0 | |
| Real | Intergen | LIFG~LTPJ | Together | 0 | -0.002 | 0.003 | |
| Pseudo | Intergen | LIFG~RTPJ | Alone | 0 | 0 | 0.001 | |
| Real | Intergen | LIFG~RTPJ | Alone | -0.002 | -0.006 | 0.001 | |
| Pseudo | Intergen | LIFG~RTPJ | Together | 0 | -0.001 | 0 | |
| Real | Intergen | LIFG~RTPJ | Together | 0.002 | -0.001 | 0.005 | |
| Pseudo | Intergen | RIFG~RIFG | Alone | 0.001 | 0 | 0.001 | No overlap |
| Real | Intergen | RIFG~RIFG | Alone | 0 | -0.004 | 0.005 | |
| Pseudo | Intergen | RIFG~RIFG | Together | 0.001 | 0.001 | 0.001 | No overlap |
| Real | Intergen | RIFG~RIFG | Together | 0 | -0.004 | 0.004 | |
| Pseudo | Intergen | RTPJ~RTPJ | Alone | 0.001 | 0 | 0.002 | |
| Real | Intergen | RTPJ~RTPJ | Alone | 0.01 | 0.003 | 0.018 | No overlap |
| Pseudo | Intergen | RTPJ~RTPJ | Together | 0 | -0.001 | 0.001 | |
| Real | Intergen | RTPJ~RTPJ | Together | -0.003 | -0.011 | 0.004 | |
| Pseudo | Intergen | LTPJ~RIFG | Alone | 0.001 | 0.001 | 0.001 | No overlap |
| Real | Intergen | LTPJ~RIFG | Alone | 0.003 | 0 | 0.006 | No overlap |
| Pseudo | Intergen | LTPJ~RIFG | Together | 0 | 0 | 0.001 | |
| Real | Intergen | LTPJ~RIFG | Together | 0 | -0.003 | 0.003 | |
| Pseudo | Intergen | RIFG~RTPJ | Alone | 0.001 | 0 | 0.001 | No overlap |
| Real | Intergen | RIFG~RTPJ | Alone | 0.004 | 0.001 | 0.008 | No overlap |
| Pseudo | Intergen | RIFG~RTPJ | Together | 0.001 | 0 | 0.001 | No overlap |
| Real | Intergen | RIFG~RTPJ | Together | -0.001 | -0.005 | 0.002 | |
| Pseudo | Samegen | LIFG~RIFG | Alone | 0 | 0 | 0 | |
| Real | Samegen | LIFG~RIFG | Alone | -0.002 | -0.005 | 0 | |
| Pseudo | Samegen | LIFG~RIFG | Together | 0 | 0 | 0.001 | No overlap |

| Dyad Type | Group | ROI Pair | Task | Slope Estimate | Lower HPD | Upper HPD | No overlap |
|---|---|---|---|---|---|---|---|
| Real | Samegen | LIFG~RIFG | Together | 0 | -0.002 | 0.003 | |
| Pseudo | Samegen | LTPJ~RTPJ | Alone | 0 | 0 | 0.001 | No overlap |
| Real | Samegen | LTPJ~RTPJ | Alone | 0.001 | -0.003 | 0.004 | |
| Pseudo | Samegen | LTPJ~RTPJ | Together | 0.001 | 0 | 0.001 | No overlap |
| Real | Samegen | LTPJ~RTPJ | Together | 0.003 | 0 | 0.006 | |
| Pseudo | Samegen | LIFG~LIFG | Alone | 0.001 | 0 | 0.001 | No overlap |
| Real | Samegen | LIFG~LIFG | Alone | 0.001 | -0.003 | 0.005 | |
| Pseudo | Samegen | LIFG~LIFG | Together | 0 | 0 | 0.001 | No overlap |
| Real | Samegen | LIFG~LIFG | Together | -0.002 | -0.005 | 0.002 | |
| Pseudo | Samegen | LTPJ~LTPJ | Alone | 0.001 | 0 | 0.001 | No overlap |
| Real | Samegen | LTPJ~LTPJ | Alone | 0 | -0.004 | 0.003 | |
| Pseudo | Samegen | LTPJ~LTPJ | Together | 0 | 0 | 0 | |
| Real | Samegen | LTPJ~LTPJ | Together | 0.001 | -0.002 | 0.005 | |
| Pseudo | Samegen | LIFG~LTPJ | Alone | 0 | 0 | 0.001 | No overlap |
| Real | Samegen | LIFG~LTPJ | Alone | 0.001 | -0.002 | 0.003 | |
| Pseudo | Samegen | LIFG~LTPJ | Together | 0 | 0 | 0 | |
| Real | Samegen | LIFG~LTPJ | Together | 0.001 | -0.002 | 0.004 | |
| Pseudo | Samegen | LIFG~RTPJ | Alone | 0 | 0 | 0 | |
| Real | Samegen | LIFG~RTPJ | Alone | 0 | -0.004 | 0.003 | |
| Pseudo | Samegen | LIFG~RTPJ | Together | 0.001 | 0 | 0.001 | No overlap |
| Real | Samegen | LIFG~RTPJ | Together | 0 | -0.003 | 0.003 | |
| Pseudo | Samegen | RIFG~RIFG | Alone | -0.001 | -0.001 | 0 | No overlap |
| Real | Samegen | RIFG~RIFG | Alone | 0.003 | -0.001 | 0.007 | |
| Pseudo | Samegen | RIFG~RIFG | Together | 0 | 0 | 0.001 | No overlap |
| Real | Samegen | RIFG~RIFG | Together | -0.001 | -0.005 | 0.002 | |
| Pseudo | Samegen | RTPJ~RTPJ | Alone | 0 | 0 | 0.001 | |
| Real | Samegen | RTPJ~RTPJ | Alone | -0.003 | -0.01 | 0.004 | |
| Pseudo | Samegen | RTPJ~RTPJ | Together | 0.002 | 0.001 | 0.002 | No overlap |
| Real | Samegen | RTPJ~RTPJ | Together | 0.004 | -0.002 | 0.011 | |
| Pseudo | Samegen | LTPJ~RIFG | Alone | 0 | 0 | 0 | |
| Real | Samegen | LTPJ~RIFG | Alone | 0.001 | -0.002 | 0.004 | |
| Pseudo | Samegen | LTPJ~RIFG | Together | 0 | 0 | 0.001 | No overlap |
| Real | Samegen | LTPJ~RIFG | Together | 0.003 | 0 | 0.005 | No overlap |
| Pseudo | Samegen | RIFG~RTPJ | Alone | 0 | -0.001 | 0 | No overlap |
| Real | Samegen | RIFG~RTPJ | Alone | 0 | -0.004 | 0.003 | |
| Pseudo | Samegen | RIFG~RTPJ | Together | 0.001 | 0.001 | 0.001 | No overlap |
| Real | Samegen | RIFG~RTPJ | Together | 0 | -0.003 | 0.003 | |

**S6 Appendix – Table D.** HbR. Contrasts between real and pseudo dyads' slopes of change in INS across sessions per ROI pair and task in unstandardised units.

| Contrast | Group | ROI Pair | Task | Estimate | Lower HPD | Upper HPD | No overlap |
|---|---|---|---|---|---|---|---|
| Real - Pseudo | Intergen | LIFG~RIFG | Alone | 0.001 | -0.002 | 0.004 | |
| Real - Pseudo | Intergen | LIFG~RIFG | Together | 0.001 | -0.002 | 0.003 | |
| Real - Pseudo | Intergen | LTPJ~RTPJ | Alone | -0.001 | -0.005 | 0.002 | |
| Real - Pseudo | Intergen | LTPJ~RTPJ | Together | 0.001 | -0.002 | 0.005 | |
| Real - Pseudo | Intergen | LIFG~LIFG | Alone | 0 | -0.003 | 0.004 | |
| Real - Pseudo | Intergen | LIFG~LIFG | Together | 0.004 | 0.001 | 0.008 | No overlap |
| Real - Pseudo | Intergen | LTPJ~LTPJ | Alone | 0.002 | -0.002 | 0.006 | |
| Real - Pseudo | Intergen | LTPJ~LTPJ | Together | -0.001 | -0.005 | 0.003 | |
| Real - Pseudo | Intergen | LIFG~LTPJ | Alone | 0 | -0.002 | 0.003 | |
| Real - Pseudo | Intergen | LIFG~LTPJ | Together | 0 | -0.003 | 0.002 | |
| Real - Pseudo | Intergen | LIFG~RTPJ | Alone | 0.003 | -0.001 | 0.006 | |
| Real - Pseudo | Intergen | LIFG~RTPJ | Together | -0.002 | -0.005 | 0.001 | |
| Real - Pseudo | Intergen | RIFG~RIFG | Alone | 0 | -0.004 | 0.005 | |
| Real - Pseudo | Intergen | RIFG~RIFG | Together | 0.001 | -0.003 | 0.005 | |
| Real - Pseudo | Intergen | RTPJ~RTPJ | Alone | -0.01 | -0.017 | -0.002 | No overlap |
| Real - Pseudo | Intergen | RTPJ~RTPJ | Together | 0.003 | -0.004 | 0.01 | |
| Real - Pseudo | Intergen | LTPJ~RIFG | Alone | -0.002 | -0.005 | 0.001 | |
| Real - Pseudo | Intergen | LTPJ~RIFG | Together | 0 | -0.002 | 0.003 | |
| Real - Pseudo | Intergen | RIFG~RTPJ | Alone | -0.004 | -0.007 | 0 | No overlap |
| Real - Pseudo | Intergen | RIFG~RTPJ | Together | 0.002 | -0.002 | 0.005 | |
| Real - Pseudo | Samegen | LIFG~RIFG | Alone | 0.002 | 0 | 0.005 | |
| Real - Pseudo | Samegen | LIFG~RIFG | Together | 0 | -0.003 | 0.003 | |
| Real - Pseudo | Samegen | LTPJ~RTPJ | Alone | 0 | -0.004 | 0.003 | |
| Real - Pseudo | Samegen | LTPJ~RTPJ | Together | -0.002 | -0.005 | 0.001 | |
| Real - Pseudo | Samegen | LIFG~LIFG | Alone | 0 | -0.004 | 0.004 | |
| Real - Pseudo | Samegen | LIFG~LIFG | Together | 0.002 | -0.002 | 0.006 | |
| Real - Pseudo | Samegen | LTPJ~LTPJ | Alone | 0.001 | -0.003 | 0.005 | |
| Real - Pseudo | Samegen | LTPJ~LTPJ | Together | -0.001 | -0.005 | 0.003 | |
| Real - Pseudo | Samegen | LIFG~LTPJ | Alone | 0 | -0.003 | 0.003 | |
| Real - Pseudo | Samegen | LIFG~LTPJ | Together | -0.001 | -0.004 | 0.002 | |
| Real - Pseudo | Samegen | LIFG~RTPJ | Alone | 0.001 | -0.003 | 0.004 | |
| Real - Pseudo | Samegen | LIFG~RTPJ | Together | 0.001 | -0.002 | 0.004 | |
| Real - Pseudo | Samegen | RIFG~RIFG | Alone | -0.003 | -0.007 | 0 | |
| Real - Pseudo | Samegen | RIFG~RIFG | Together | 0.002 | -0.002 | 0.005 | |
| Real - Pseudo | Samegen | RTPJ~RTPJ | Alone | 0.003 | -0.004 | 0.01 | |
| Real - Pseudo | Samegen | RTPJ~RTPJ | Together | -0.003 | -0.009 | 0.003 | |
| Real - Pseudo | Samegen | LTPJ~RIFG | Alone | -0.001 | -0.003 | 0.002 | |
| Real - Pseudo | Samegen | LTPJ~RIFG | Together | -0.002 | -0.005 | 0 | |
| Real - Pseudo | Samegen | RIFG~RTPJ | Alone | 0 | -0.004 | 0.003 | |
| Real - Pseudo | Samegen | RIFG~RTPJ | Together | 0.001 | -0.002 | 0.004 | |

**S6 Appendix – Table E.** HbR. Composition of non-homologous ROI pairs, to assess whether an ROI belonging to an older or younger member of a dyad influenced INS levels for all sessions (top) and across sessions (bottom). In contrast column, 'o' indicated that ROI belongs to older adult and 'y' indicates that ROI belongs to younger adult.

| Contrast | Task | Estimate | Lower HPD | Upper HPD | No overlap |
|---|---|---|---|---|---|
| *All sessions* | | | | | |
| o_l_ifg_y_l_tpj - o_l_tpj_y_l_ifg | Together | 0.006 | -0.001 | 0.014 | trend |
| o_l_ifg_y_r_ifg - o_r_ifg_y_l_ifg | Together | 0 | -0.007 | 0.007 | |
| o_l_tpj_y_r_ifg - o_r_ifg_y_l_tpj | Together | -0.008 | -0.015 | 0 | No overlap |
| *Across sessions* | | | | | |
| o_r_ifg_y_r_tpj - o_r_tpj_y_r_ifg | Alone | -0.004 | -0.017 | 0.008 | |

**S6 Appendix – Table F.** HbR. INS levels for all sessions: contrasts between tasks, groups and the task*group interaction per ROI pair in unstandardised units.

| Contrast | ROI Pair | Estimate | Lower HPD | Upper HPD | No overlap |
|---|---|---|---|---|---|
| Tog-Alo | LIFG~RIFG | 0.014 | 0.006 | 0.023 | No overlap |
| Int-Sam | LIFG~RIFG | 0 | -0.01 | 0.01 | |
| Tog-Alo Int | LIFG~RIFG | 0.011 | 0.005 | 0.017 | No overlap |
| Tog-Alo Sam | LIFG~RIFG | 0.003 | -0.003 | 0.009 | |
| Int-Sam Alo | LIFG~RIFG | -0.004 | -0.011 | 0.003 | |
| Int-Sam Tog | LIFG~RIFG | 0.004 | -0.002 | 0.01 | |
| Tog-Alo | LTPJ~LTPJ | 0.006 | -0.006 | 0.017 | |
| Int-Sam | LTPJ~LTPJ | -0.004 | -0.017 | 0.009 | |
| Tog-Alo Int | LTPJ~LTPJ | 0.004 | -0.004 | 0.013 | |
| Tog-Alo Sam | LTPJ~LTPJ | 0.002 | -0.006 | 0.01 | |
| Int-Sam Alo | LTPJ~LTPJ | -0.003 | -0.012 | 0.007 | |
| Int-Sam Tog | LTPJ~LTPJ | -0.001 | -0.009 | 0.007 | |
| Tog-Alo | LIFG~LTPJ | 0.004 | -0.005 | 0.012 | |
| Int-Sam | LIFG~LTPJ | 0.002 | -0.008 | 0.013 | |
| Tog-Alo Int | LIFG~LTPJ | 0.004 | -0.002 | 0.01 | |
| Tog-Alo Sam | LIFG~LTPJ | 0 | -0.006 | 0.006 | |
| Int-Sam Alo | LIFG~LTPJ | -0.001 | -0.008 | 0.007 | |
| Int-Sam Tog | LIFG~LTPJ | 0.003 | -0.003 | 0.009 | |
| Tog-Alo | RIFG~RIFG | 0.006 | -0.006 | 0.019 | |
| Int-Sam | RIFG~RIFG | -0.004 | -0.017 | 0.01 | |
| Tog-Alo Int | RIFG~RIFG | -0.002 | -0.011 | 0.007 | |
| Tog-Alo Sam | RIFG~RIFG | 0.009 | 0 | 0.018 | |
| Int-Sam Alo | RIFG~RIFG | 0.004 | -0.007 | 0.014 | |
| Int-Sam Tog | RIFG~RIFG | -0.007 | -0.016 | 0.001 | |
| Tog-Alo | RIFG~LTPJ | 0.009 | 0 | 0.018 | No overlap |
| Int-Sam | RIFG~LTPJ | -0.007 | -0.017 | 0.002 | |
| Tog-Alo Int | RIFG~LTPJ | 0.005 | -0.001 | 0.011 | |
| Tog-Alo Sam | RIFG~LTPJ | 0.004 | -0.002 | 0.01 | |
| Int-Sam Alo | RIFG~LTPJ | -0.004 | -0.011 | 0.003 | |
| Int-Sam Tog | RIFG~LTPJ | -0.003 | -0.009 | 0.002 | |

**S6 Appendix – Table G.** HbR. Change in INS across sessions: contrasts between tasks, groups and the task*group interaction per ROI pair in unstandardised units.

| Contrast | ROI Pair | Estimate | Lower HPD | Upper HPD | No overlap |
|---|---|---|---|---|---|
| Tog-Alo | LIFG~LIFG | -0.007 | -0.014 | 0.001 | |
| Int-Sam | LIFG~LIFG | -0.004 | -0.011 | 0.004 | |
| Tog-Alo Int | LIFG~LIFG | -0.004 | -0.009 | 0.002 | |
| Tog-Alo Sam | LIFG~LIFG | -0.003 | -0.008 | 0.002 | |
| Int-Sam Alo | LIFG~LIFG | -0.001 | -0.007 | 0.004 | |
| Int-Sam Tog | LIFG~LIFG | -0.002 | -0.007 | 0.003 | |
| Tog-Alo | RTPJ~RTPJ | -0.006 | -0.02 | 0.007 | |
| Int-Sam | RTPJ~RTPJ | 0.005 | -0.008 | 0.019 | |
| Tog-Alo Int | RTPJ~RTPJ | -0.014 | -0.023 | -0.004 | No overlap |
| Tog-Alo Sam | RTPJ~RTPJ | 0.007 | -0.002 | 0.016 | |
| Int-Sam Alo | RTPJ~RTPJ | 0.013 | 0.004 | 0.023 | No overlap |
| Int-Sam Tog | RTPJ~RTPJ | -0.008 | -0.017 | 0.002 | |
| Tog-Alo | RIFG~RTPJ | -0.006 | -0.012 | 0.001 | |
| Int-Sam | RIFG~RTPJ | 0.003 | -0.004 | 0.01 | |
| Tog-Alo Int | RIFG~RTPJ | -0.006 | -0.01 | -0.001 | No overlap |
| Tog-Alo Sam | RIFG~RTPJ | 0 | -0.005 | 0.005 | |
| Int-Sam Alo | RIFG~RTPJ | 0.004 | -0.001 | 0.009 | |
| Int-Sam Tog | RIFG~RTPJ | -0.001 | -0.006 | 0.003 | |

**S6 Appendix – Table H.** HbR. Relationships between session and INS in unstandardised units. Effects are presented in the following order: Main, group, task, and simple effects.

| Measure | Group | Task | ROI Pairs | Estimate | Lower HPD | Upper HPD | No overlap |
|---|---|---|---|---|---|---|---|
| *Main effect* | | | | | | | |
| session | | | LIFG~RIFG | -0.0009 | -0.0023 | 0.0006 | |
| session | | | LIFG~LIFG | -0.0009 | -0.0030 | 0.0012 | |
| session | | | LTPJ~LTPJ | -0.0004 | -0.0026 | 0.0016 | |
| session | | | LIFG~LTPJ | 0.0002 | -0.0013 | 0.0017 | |
| session | | | RIFG~RIFG | 0.0010 | -0.0012 | 0.0032 | |
| session | | | RTPJ~RTPJ | 0.0003 | -0.0039 | 0.0043 | |
| session | | | RIFG~LTPJ | 0.0012 | -0.0003 | 0.0027 | |
| session | | | RIFG~RTPJ | 0.0010 | -0.0009 | 0.0029 | |
| *Group effect* | | | | | | | |
| session | Intergen | | LIFG~RIFG | -0.0005 | -0.0026 | 0.0015 | |
| session | Samegen | | LIFG~RIFG | -0.0012 | -0.0033 | 0.0010 | |
| session | Intergen | | LIFG~LIFG | -0.0024 | -0.0053 | 0.0005 | |
| session | Samegen | | LIFG~LIFG | 0.0006 | -0.0025 | 0.0036 | |
| session | Intergen | | LTPJ~LTPJ | 0.0002 | -0.0026 | 0.0031 | |
| session | Samegen | | LTPJ~LTPJ | -0.0011 | -0.0042 | 0.0019 | |
| session | Intergen | | LIFG~LTPJ | -0.0004 | -0.0025 | 0.0017 | |
| session | Samegen | | LIFG~LTPJ | 0.0008 | -0.0014 | 0.0030 | |
| session | Intergen | | RIFG~RIFG | 0.0002 | -0.0030 | 0.0033 | |
| session | Samegen | | RIFG~RIFG | 0.0018 | -0.0014 | 0.0050 | |
| session | Intergen | | RTPJ~RTPJ | 0.0017 | -0.0041 | 0.0074 | |
| session | Samegen | | RTPJ~RTPJ | -0.0011 | -0.0069 | 0.0047 | |
| session | Intergen | | RIFG~LTPJ | 0.0014 | -0.0007 | 0.0035 | |
| session | Samegen | | RIFG~LTPJ | 0.0010 | -0.0012 | 0.0031 | |
| session | Intergen | | RIFG~RTPJ | 0.0011 | -0.0015 | 0.0038 | |
| session | Samegen | | RIFG~RTPJ | 0.0008 | -0.0019 | 0.0035 | |
| *Task effect* | | | | | | | |
| session | | Alone | LIFG~RIFG | -0.0013 | -0.0035 | 0.0009 | |
| session | | Together | LIFG~RIFG | -0.0004 | -0.0023 | 0.0016 | |
| session | | Alone | LIFG~LIFG | 0.0010 | -0.0022 | 0.0041 | |
| session | | Together | LIFG~LIFG | -0.0028 | -0.0057 | -0.0001 | No overlap |
| session | | Alone | LTPJ~LTPJ | -0.0010 | -0.0041 | 0.0021 | |
| session | | Together | LTPJ~LTPJ | 0.0001 | -0.0027 | 0.0028 | |
| session | | Alone | LIFG~LTPJ | 0.0001 | -0.0021 | 0.0023 | |
| session | | Together | LIFG~LTPJ | 0.0003 | -0.0017 | 0.0024 | |
| session | | Alone | RIFG~RIFG | 0.0026 | -0.0007 | 0.0059 | |
| session | | Together | RIFG~RIFG | -0.0006 | -0.0036 | 0.0023 | |
| session | | Alone | RTPJ~RTPJ | 0.0005 | -0.0055 | 0.0064 | |
| session | | Together | RTPJ~RTPJ | 0.0001 | -0.0053 | 0.0058 | |
| session | | Alone | RIFG~LTPJ | 0.0018 | -0.0004 | 0.0041 | |
| session | | Together | RIFG~LTPJ | 0.0006 | -0.0014 | 0.0027 | |
| session | | Alone | RIFG~RTPJ | 0.0018 | -0.0011 | 0.0045 | |
| session | | Together | RIFG~RTPJ | 0.0002 | -0.0023 | 0.0028 | |

| Measure | Group | Task | ROI Pairs | Estimate | Lower HPD | Upper HPD | No overlap |
|---|---|---|---|---|---|---|---|
| *Simple effect* | | | | | | | |
| session | Intergen | Alone | LIFG~RIFG | -0.0002 | -0.0033 | 0.0028 | |
| session | Intergen | Together | LIFG~RIFG | -0.0008 | -0.0036 | 0.0019 | |
| session | Samegen | Alone | LIFG~RIFG | -0.0024 | -0.0054 | 0.0008 | |
| session | Samegen | Together | LIFG~RIFG | 0.0000 | -0.0029 | 0.0028 | |
| session | Intergen | Alone | LIFG~LIFG | -0.0007 | -0.0049 | 0.0036 | |
| session | Intergen | Together | LIFG~LIFG | -0.0041 | -0.0079 | -0.0001 | No overlap |
| session | Samegen | Alone | LIFG~LIFG | 0.0027 | -0.0019 | 0.0072 | |
| session | Samegen | Together | LIFG~LIFG | -0.0015 | -0.0057 | 0.0025 | |
| session | Intergen | Alone | LTPJ~LTPJ | 0.0005 | -0.0037 | 0.0048 | |
| session | Intergen | Together | LTPJ~LTPJ | 0.0000 | -0.0038 | 0.0038 | |
| session | Samegen | Alone | LTPJ~LTPJ | -0.0024 | -0.0069 | 0.0021 | |
| session | Samegen | Together | LTPJ~LTPJ | 0.0002 | -0.0037 | 0.0044 | |
| session | Intergen | Alone | LIFG~LTPJ | -0.0009 | -0.0040 | 0.0021 | |
| session | Intergen | Together | LIFG~LTPJ | 0.0002 | -0.0026 | 0.0030 | |
| session | Samegen | Alone | LIFG~LTPJ | 0.0011 | -0.0021 | 0.0044 | |
| session | Samegen | Together | LIFG~LTPJ | 0.0004 | -0.0025 | 0.0033 | |
| session | Intergen | Alone | RIFG~RIFG | 0.0007 | -0.0039 | 0.0053 | |
| session | Intergen | Together | RIFG~RIFG | -0.0003 | -0.0043 | 0.0039 | |
| session | Samegen | Alone | RIFG~RIFG | 0.0045 | -0.0001 | 0.0093 | |
| session | Samegen | Together | RIFG~RIFG | -0.0009 | -0.0051 | 0.0033 | |
| session | Intergen | Alone | RTPJ~RTPJ | 0.0086 | 0.0005 | 0.0166 | No overlap |
| session | Intergen | Together | RTPJ~RTPJ | -0.0052 | -0.0134 | 0.0029 | |
| session | Samegen | Alone | RTPJ~RTPJ | -0.0076 | -0.0160 | 0.0012 | |
| session | Samegen | Together | RTPJ~RTPJ | 0.0054 | -0.0024 | 0.0127 | |
| session | Intergen | Alone | RIFG~LTPJ | 0.0033 | 0.0002 | 0.0064 | No overlap |
| session | Intergen | Together | RIFG~LTPJ | -0.0005 | -0.0033 | 0.0023 | |
| session | Samegen | Alone | RIFG~LTPJ | 0.0002 | -0.0031 | 0.0034 | |
| session | Samegen | Together | RIFG~LTPJ | 0.0017 | -0.0012 | 0.0046 | |
| session | Intergen | Alone | RIFG~RTPJ | 0.0034 | -0.0006 | 0.0072 | |
| session | Intergen | Together | RIFG~RTPJ | -0.0011 | -0.0047 | 0.0025 | |
| session | Samegen | Alone | RIFG~RTPJ | 0.0001 | -0.0040 | 0.0041 | |
| session | Samegen | Together | RIFG~RTPJ | 0.0015 | -0.0021 | 0.0051 | |

**S6 Appendix – Table I.** HbR. Contrasts pertaining to session ~ INS relationship in unstandardised units. Contrasts are presented in the following order: Group, task, and interaction contrasts.

| Measure | Group | Task | ROI Pairs | Estimate | Lower HPD | Upper HPD | No overlap |
|---|---|---|---|---|---|---|---|
| *Group contrast* | | | | | | | |
| session | Intergen-Samegen | | LIFG~RIFG | 0.0007 | -0.0023 | 0.0036 | |
| session | Intergen-Samegen | | LIFG~LIFG | -0.0030 | -0.0072 | 0.0013 | |
| session | Intergen-Samegen | | LTPJ~LTPJ | 0.0013 | -0.0029 | 0.0054 | |
| session | Intergen-Samegen | | LIFG~LTPJ | -0.0011 | -0.0041 | 0.0020 | |
| session | Intergen-Samegen | | RIFG~RIFG | -0.0016 | -0.0062 | 0.0028 | |
| session | Intergen-Samegen | | RTPJ~RTPJ | 0.0028 | -0.0054 | 0.0111 | |
| session | Intergen-Samegen | | RIFG~LTPJ | 0.0004 | -0.0026 | 0.0035 | |
| session | Intergen-Samegen | | RIFG~RTPJ | 0.0003 | -0.0034 | 0.0042 | |
| *Task contrast* | | | | | | | |
| session | | Alone-Together | LIFG~RIFG | -0.0009 | -0.0039 | 0.0020 | |
| session | | Alone-Together | LIFG~LIFG | 0.0038 | -0.0004 | 0.0079 | |
| session | | Alone-Together | LTPJ~LTPJ | -0.0011 | -0.0051 | 0.0032 | |
| session | | Alone-Together | LIFG~LTPJ | -0.0002 | -0.0032 | 0.0028 | |
| session | | Alone-Together | RIFG~RIFG | 0.0033 | -0.0012 | 0.0076 | |
| session | | Alone-Together | RTPJ~RTPJ | 0.0004 | -0.0078 | 0.0083 | |
| session | | Alone-Together | RIFG~LTPJ | 0.0011 | -0.0019 | 0.0040 | |
| session | | Alone-Together | RIFG~RTPJ | 0.0016 | -0.0021 | 0.0054 | |
| *Interactions* | | | | | | | |
| session | Intergen-Samegen | Alone | LIFG~RIFG | 0.0021 | -0.0023 | 0.0064 | |
| session | Intergen-Samegen | Together | LIFG~RIFG | -0.0008 | -0.0047 | 0.0032 | |
| session | Intergen-Samegen | Alone | LIFG~LIFG | -0.0034 | -0.0095 | 0.0030 | |
| session | Intergen-Samegen | Together | LIFG~LIFG | -0.0026 | -0.0081 | 0.0031 | |
| session | Intergen-Samegen | Alone | LTPJ~LTPJ | 0.0029 | -0.0036 | 0.0089 | |
| session | Intergen-Samegen | Together | LTPJ~LTPJ | -0.0003 | -0.0057 | 0.0055 | |
| session | Intergen-Samegen | Alone | LIFG~LTPJ | -0.0020 | -0.0065 | 0.0024 | |
| session | Intergen-Samegen | Together | LIFG~LTPJ | -0.0002 | -0.0041 | 0.0040 | |
| session | Intergen-Samegen | Alone | RIFG~RIFG | -0.0038 | -0.0102 | 0.0030 | |
| session | Intergen-Samegen | Together | RIFG~RIFG | 0.0006 | -0.0052 | 0.0066 | |
| session | Intergen-Samegen | Alone | RTPJ~RTPJ | 0.0161 | 0.0044 | 0.0279 | No overlap |
| session | Intergen-Samegen | Together | RTPJ~RTPJ | -0.0104 | -0.0217 | 0.0006 | |
| session | Intergen-Samegen | Alone | RIFG~LTPJ | 0.0031 | -0.0013 | 0.0076 | |
| session | Intergen-Samegen | Together | RIFG~LTPJ | -0.0022 | -0.0062 | 0.0018 | |
| session | Intergen-Samegen | Alone | RIFG~RTPJ | 0.0032 | -0.0024 | 0.0090 | |
| session | Intergen-Samegen | Together | RIFG~RTPJ | -0.0026 | -0.0078 | 0.0023 | |
| session | Intergen | Alone-Together | LIFG~RIFG | 0.0006 | -0.0034 | 0.0048 | |
| session | Samegen | Alone-Together | LIFG~RIFG | -0.0023 | -0.0065 | 0.0018 | |
| session | Intergen | Alone-Together | LIFG~LIFG | 0.0034 | -0.0024 | 0.0091 | |
| session | Samegen | Alone-Together | LIFG~LIFG | 0.0042 | -0.0019 | 0.0101 | |
| session | Intergen | Alone-Together | LTPJ~LTPJ | 0.0005 | -0.0053 | 0.0062 | |
| session | Samegen | Alone-Together | LTPJ~LTPJ | -0.0026 | -0.0085 | 0.0036 | |
| session | Intergen | Alone-Together | LIFG~LTPJ | -0.0011 | -0.0052 | 0.0030 | |
| session | Samegen | Alone-Together | LIFG~LTPJ | 0.0007 | -0.0037 | 0.0049 | |

| Measure | Group | Task | ROI Pairs | Estimate | Lower HPD | Upper HPD | No overlap |
|---|---|---|---|---|---|---|---|
| session | Intergen | Alone-Together | RIFG~RIFG | 0.0010 | -0.0051 | 0.0072 | |
| session | Samegen | Alone-Together | RIFG~RIFG | 0.0054 | -0.0008 | 0.0116 | |
| session | Intergen | Alone-Together | RTPJ~RTPJ | 0.0137 | 0.0022 | 0.0248 | No overlap |
| session | Samegen | Alone-Together | RTPJ~RTPJ | -0.0129 | -0.0247 | -0.0018 | No overlap |
| session | Intergen | Alone-Together | RIFG~LTPJ | 0.0039 | -0.0003 | 0.0081 | |
| session | Samegen | Alone-Together | RIFG~LTPJ | -0.0015 | -0.0059 | 0.0028 | |
| session | Intergen | Alone-Together | RIFG~RTPJ | 0.0045 | -0.0008 | 0.0097 | |
| session | Samegen | Alone-Together | RIFG~RTPJ | -0.0014 | -0.0066 | 0.0041 | |

**Supplementary Materials:** Moffat, Dumas and Cross (2026). *Social interactions between people of same and different generations shape longitudinal changes in interpersonal neural synchrony, loneliness, and social connection.*

**S6 Appendix – Table J.** HbR. Relationships between loneliness (sum) and INS in unstandardised units. Effects are presented in the following order: Main, group, task, and simple effects.

| Measure | Group | Task | ROI Pairs | Estimate | Lower HPD | Upper HPD | No overlap |
|---|---|---|---|---|---|---|---|
| *Main effect* | | | | | | | |
| sum loneliness | | | LIFG~RIFG | -0.0017 | -0.0029 | -0.0004 | No overlap |
| sum loneliness | | | LIFG~LIFG | -0.0007 | -0.0023 | 0.0009 | |
| sum loneliness | | | LTPJ~LTPJ | -0.0004 | -0.0020 | 0.0011 | |
| sum loneliness | | | LIFG~LTPJ | -0.0010 | -0.0023 | 0.0002 | |
| sum loneliness | | | RIFG~RIFG | -0.0010 | -0.0027 | 0.0008 | |
| sum loneliness | | | RTPJ~RTPJ | 0.0003 | -0.0025 | 0.0032 | |
| sum loneliness | | | RIFG~LTPJ | -0.0004 | -0.0016 | 0.0007 | |
| sum loneliness | | | RIFG~RTPJ | -0.0008 | -0.0022 | 0.0007 | |
| *Group effect* | | | | | | | |
| sum loneliness | Intergen | | LIFG~RIFG | -0.0014 | -0.0032 | 0.0003 | |
| sum loneliness | Samegen | | LIFG~RIFG | -0.0020 | -0.0038 | -0.0003 | No overlap |
| sum loneliness | Intergen | | LIFG~LIFG | -0.0011 | -0.0033 | 0.0011 | |
| sum loneliness | Samegen | | LIFG~LIFG | -0.0003 | -0.0027 | 0.0020 | |
| sum loneliness | Intergen | | LTPJ~LTPJ | -0.0010 | -0.0032 | 0.0012 | |
| sum loneliness | Samegen | | LTPJ~LTPJ | 0.0002 | -0.0021 | 0.0026 | |
| sum loneliness | Intergen | | LIFG~LTPJ | -0.0014 | -0.0031 | 0.0004 | |
| sum loneliness | Samegen | | LIFG~LTPJ | -0.0007 | -0.0026 | 0.0012 | |
| sum loneliness | Intergen | | RIFG~RIFG | -0.0009 | -0.0033 | 0.0015 | |
| sum loneliness | Samegen | | RIFG~RIFG | -0.0011 | -0.0035 | 0.0014 | |
| sum loneliness | Intergen | | RTPJ~RTPJ | -0.0003 | -0.0040 | 0.0034 | |
| sum loneliness | Samegen | | RTPJ~RTPJ | 0.0009 | -0.0035 | 0.0051 | |
| sum loneliness | Intergen | | RIFG~LTPJ | 0.0007 | -0.0009 | 0.0023 | |
| sum loneliness | Samegen | | RIFG~LTPJ | -0.0015 | -0.0031 | 0.0002 | |
| sum loneliness | Intergen | | RIFG~RTPJ | 0.0002 | -0.0018 | 0.0023 | |
| sum loneliness | Samegen | | RIFG~RTPJ | -0.0017 | -0.0038 | 0.0004 | |
| *Task effect* | | | | | | | |
| sum loneliness | | Alone | LIFG~RIFG | -0.0020 | -0.0038 | -0.0001 | No overlap |
| sum loneliness | | Together | LIFG~RIFG | -0.0014 | -0.0028 | 0.0000 | |
| sum loneliness | | Alone | LIFG~LIFG | -0.0011 | -0.0036 | 0.0014 | |
| sum loneliness | | Together | LIFG~LIFG | -0.0003 | -0.0022 | 0.0017 | |
| sum loneliness | | Alone | LTPJ~LTPJ | 0.0002 | -0.0023 | 0.0027 | |
| sum loneliness | | Together | LTPJ~LTPJ | -0.0011 | -0.0031 | 0.0008 | |
| sum loneliness | | Alone | LIFG~LTPJ | -0.0010 | -0.0029 | 0.0008 | |
| sum loneliness | | Together | LIFG~LTPJ | -0.0010 | -0.0025 | 0.0004 | |
| sum loneliness | | Alone | RIFG~RIFG | -0.0008 | -0.0035 | 0.0018 | |
| sum loneliness | | Together | RIFG~RIFG | -0.0011 | -0.0032 | 0.0010 | |
| sum loneliness | | Alone | RTPJ~RTPJ | 0.0010 | -0.0032 | 0.0053 | |
| sum loneliness | | Together | RTPJ~RTPJ | -0.0003 | -0.0040 | 0.0032 | |
| sum loneliness | | Alone | RIFG~LTPJ | 0.0001 | -0.0017 | 0.0019 | |
| sum loneliness | | Together | RIFG~LTPJ | -0.0009 | -0.0023 | 0.0005 | |
| sum loneliness | | Alone | RIFG~RTPJ | -0.0007 | -0.0029 | 0.0015 | |
| sum loneliness | | Together | RIFG~RTPJ | -0.0008 | -0.0026 | 0.0009 | |

| Measure | Group | Task | ROI Pairs | Estimate | Lower HPD | Upper HPD | No overlap |
|---|---|---|---|---|---|---|---|
| *Main effect* | | | | | | | |
| sum loneliness | Intergen | Alone | LIFG~RIFG | -0.0008 | -0.0034 | 0.0017 | |
| sum loneliness | Intergen | Together | LIFG~RIFG | -0.0020 | -0.0040 | 0.0000 | |
| sum loneliness | Samegen | Alone | LIFG~RIFG | -0.0031 | -0.0057 | -0.0006 | No overlap |
| sum loneliness | Samegen | Together | LIFG~RIFG | -0.0008 | -0.0029 | 0.0013 | |
| sum loneliness | Intergen | Alone | LIFG~LIFG | -0.0002 | -0.0037 | 0.0031 | |
| sum loneliness | Intergen | Together | LIFG~LIFG | -0.0020 | -0.0047 | 0.0006 | |
| sum loneliness | Samegen | Alone | LIFG~LIFG | -0.0021 | -0.0056 | 0.0016 | |
| sum loneliness | Samegen | Together | LIFG~LIFG | 0.0014 | -0.0014 | 0.0042 | |
| sum loneliness | Intergen | Alone | LTPJ~LTPJ | -0.0011 | -0.0044 | 0.0024 | |
| sum loneliness | Intergen | Together | LTPJ~LTPJ | -0.0010 | -0.0036 | 0.0017 | |
| sum loneliness | Samegen | Alone | LTPJ~LTPJ | 0.0015 | -0.0022 | 0.0050 | |
| sum loneliness | Samegen | Together | LTPJ~LTPJ | -0.0012 | -0.0041 | 0.0016 | |
| sum loneliness | Intergen | Alone | LIFG~LTPJ | -0.0007 | -0.0033 | 0.0019 | |
| sum loneliness | Intergen | Together | LIFG~LTPJ | -0.0020 | -0.0041 | 0.0000 | |
| sum loneliness | Samegen | Alone | LIFG~LTPJ | -0.0014 | -0.0041 | 0.0013 | |
| sum loneliness | Samegen | Together | LIFG~LTPJ | 0.0000 | -0.0021 | 0.0022 | |
| sum loneliness | Intergen | Alone | RIFG~RIFG | -0.0008 | -0.0044 | 0.0030 | |
| sum loneliness | Intergen | Together | RIFG~RIFG | -0.0010 | -0.0040 | 0.0018 | |
| sum loneliness | Samegen | Alone | RIFG~RIFG | -0.0009 | -0.0047 | 0.0028 | |
| sum loneliness | Samegen | Together | RIFG~RIFG | -0.0012 | -0.0041 | 0.0017 | |
| sum loneliness | Intergen | Alone | RTPJ~RTPJ | -0.0008 | -0.0064 | 0.0047 | |
| sum loneliness | Intergen | Together | RTPJ~RTPJ | 0.0003 | -0.0046 | 0.0049 | |
| sum loneliness | Samegen | Alone | RTPJ~RTPJ | 0.0027 | -0.0038 | 0.0090 | |
| sum loneliness | Samegen | Together | RTPJ~RTPJ | -0.0010 | -0.0065 | 0.0044 | |
| sum loneliness | Intergen | Alone | RIFG~LTPJ | 0.0014 | -0.0011 | 0.0039 | |
| sum loneliness | Intergen | Together | RIFG~LTPJ | 0.0000 | -0.0020 | 0.0019 | |
| sum loneliness | Samegen | Alone | RIFG~LTPJ | -0.0011 | -0.0037 | 0.0015 | |
| sum loneliness | Samegen | Together | RIFG~LTPJ | -0.0018 | -0.0039 | 0.0001 | |
| sum loneliness | Intergen | Alone | RIFG~RTPJ | -0.0003 | -0.0033 | 0.0029 | |
| sum loneliness | Intergen | Together | RIFG~RTPJ | 0.0007 | -0.0018 | 0.0031 | |
| sum loneliness | Samegen | Alone | RIFG~RTPJ | -0.0011 | -0.0043 | 0.0021 | |
| sum loneliness | Samegen | Together | RIFG~RTPJ | -0.0023 | -0.0048 | 0.0002 | |

**Supplementary Materials:** Moffat, Dumas and Cross (2026). *Social interactions between people of same and different generations shape longitudinal changes in interpersonal neural synchrony, loneliness, and social connection.*

**S6 Appendix – Table K.** HbR. Contrasts pertaining to loneliness (sum) ~ INS relationship in unstandardised units. Contrasts are presented in the following order: Group, task, and interaction contrasts.

| Measure | Group | Task | ROI Pairs | Estimate | Lower HPD | Upper HPD | No overlap |
|---|---|---|---|---|---|---|---|
| *Group contrast* | | | | | | | |
| sum loneliness | Intergen-Samegen | | LIFG~RIFG | 0.0006 | -0.0018 | 0.0031 | |
| sum loneliness | Intergen-Samegen | | LIFG~LIFG | -0.0008 | -0.0039 | 0.0025 | |
| sum loneliness | Intergen-Samegen | | LTPJ~LTPJ | -0.0012 | -0.0044 | 0.0020 | |
| sum loneliness | Intergen-Samegen | | LIFG~LTPJ | -0.0007 | -0.0033 | 0.0019 | |
| sum loneliness | Intergen-Samegen | | RIFG~RIFG | 0.0001 | -0.0033 | 0.0037 | |
| sum loneliness | Intergen-Samegen | | RTPJ~RTPJ | -0.0011 | -0.0068 | 0.0046 | |
| sum loneliness | Intergen-Samegen | | RIFG~LTPJ | 0.0022 | -0.0001 | 0.0045 | |
| sum loneliness | Intergen-Samegen | | RIFG~RTPJ | 0.0019 | -0.0010 | 0.0049 | |
| *Task contrast* | | | | | | | |
| sum loneliness | | Alone-Together | LIFG~RIFG | -0.0006 | -0.0027 | 0.0016 | |
| sum loneliness | | Alone-Together | LIFG~LIFG | -0.0008 | -0.0039 | 0.0023 | |
| sum loneliness | | Alone-Together | LTPJ~LTPJ | 0.0013 | -0.0018 | 0.0044 | |
| sum loneliness | | Alone-Together | LIFG~LTPJ | 0.0000 | -0.0023 | 0.0021 | |
| sum loneliness | | Alone-Together | RIFG~RIFG | 0.0003 | -0.0029 | 0.0036 | |
| sum loneliness | | Alone-Together | RTPJ~RTPJ | 0.0013 | -0.0041 | 0.0068 | |
| sum loneliness | | Alone-Together | RIFG~LTPJ | 0.0010 | -0.0012 | 0.0033 | |
| sum loneliness | | Alone-Together | RIFG~RTPJ | 0.0001 | -0.0026 | 0.0029 | |
| *Interactions* | | | | | | | |
| sum loneliness | Intergen-Samegen | Alone | LIFG~RIFG | 0.0023 | -0.0012 | 0.0060 | |
| sum loneliness | Intergen-Samegen | Together | LIFG~RIFG | -0.0012 | -0.0039 | 0.0018 | |
| sum loneliness | Intergen-Samegen | Alone | LIFG~LIFG | 0.0019 | -0.0031 | 0.0068 | |
| sum loneliness | Intergen-Samegen | Together | LIFG~LIFG | -0.0034 | -0.0073 | 0.0004 | |
| sum loneliness | Intergen-Samegen | Alone | LTPJ~LTPJ | -0.0026 | -0.0075 | 0.0023 | |
| sum loneliness | Intergen-Samegen | Together | LTPJ~LTPJ | 0.0002 | -0.0037 | 0.0041 | |
| sum loneliness | Intergen-Samegen | Alone | LIFG~LTPJ | 0.0007 | -0.0031 | 0.0044 | |
| sum loneliness | Intergen-Samegen | Together | LIFG~LTPJ | -0.0020 | -0.0051 | 0.0010 | |
| sum loneliness | Intergen-Samegen | Alone | RIFG~RIFG | 0.0001 | -0.0050 | 0.0054 | |
| sum loneliness | Intergen-Samegen | Together | RIFG~RIFG | 0.0002 | -0.0038 | 0.0044 | |
| sum loneliness | Intergen-Samegen | Alone | RTPJ~RTPJ | -0.0035 | -0.0121 | 0.0049 | |
| sum loneliness | Intergen-Samegen | Together | RTPJ~RTPJ | 0.0012 | -0.0059 | 0.0084 | |
| sum loneliness | Intergen-Samegen | Alone | RIFG~LTPJ | 0.0025 | -0.0010 | 0.0062 | |
| sum loneliness | Intergen-Samegen | Together | RIFG~LTPJ | 0.0018 | -0.0010 | 0.0046 | |
| sum loneliness | Intergen-Samegen | Alone | RIFG~RTPJ | 0.0008 | -0.0035 | 0.0053 | |
| sum loneliness | Intergen-Samegen | Together | RIFG~RTPJ | 0.0029 | -0.0007 | 0.0063 | |
| sum loneliness | Intergen | Alone-Together | LIFG~RIFG | 0.0012 | -0.0018 | 0.0042 | |
| sum loneliness | Samegen | Alone-Together | LIFG~RIFG | -0.0023 | -0.0054 | 0.0007 | |
| sum loneliness | Intergen | Alone-Together | LIFG~LIFG | 0.0018 | -0.0024 | 0.0062 | |
| sum loneliness | Samegen | Alone-Together | LIFG~LIFG | -0.0035 | -0.0079 | 0.0011 | |
| sum loneliness | Intergen | Alone-Together | LTPJ~LTPJ | -0.0001 | -0.0044 | 0.0042 | |
| sum loneliness | Samegen | Alone-Together | LTPJ~LTPJ | 0.0027 | -0.0017 | 0.0073 | |
| sum loneliness | Intergen | Alone-Together | LIFG~LTPJ | 0.0013 | -0.0018 | 0.0043 | |
| sum loneliness | Samegen | Alone-Together | LIFG~LTPJ | -0.0014 | -0.0045 | 0.0018 | |

| Measure | Group | Task | ROI Pairs | Estimate | Lower HPD | Upper HPD | No overlap |
|---|---|---|---|---|---|---|---|
| sum loneliness | Intergen | Alone-Together | RIFG~RIFG | 0.0002 | -0.0042 | 0.0048 | |
| sum loneliness | Samegen | Alone-Together | RIFG~RIFG | 0.0004 | -0.0042 | 0.0049 | |
| sum loneliness | Intergen | Alone-Together | RTPJ~RTPJ | -0.0010 | -0.0081 | 0.0060 | |
| sum loneliness | Samegen | Alone-Together | RTPJ~RTPJ | 0.0037 | -0.0045 | 0.0120 | |
| sum loneliness | Intergen | Alone-Together | RIFG~LTPJ | 0.0014 | -0.0018 | 0.0045 | |
| sum loneliness | Samegen | Alone-Together | RIFG~LTPJ | 0.0007 | -0.0025 | 0.0039 | |
| sum loneliness | Intergen | Alone-Together | RIFG~RTPJ | -0.0010 | -0.0047 | 0.0029 | |
| sum loneliness | Samegen | Alone-Together | RIFG~RTPJ | 0.0011 | -0.0027 | 0.0050 | |

**S6 Appendix – Table L.** HbR. Relationships between loneliness (dif) and INS in unstandardised units. Effects are presented in the following order: Main, group, task, and simple effects.

| Measure | Group | Task | ROI Pairs | Estimate | Lower HPD | Upper HPD | No overlap |
|---|---|---|---|---|---|---|---|
| *Main effect* | | | | | | | |
| dif loneliness | | | LIFG~RIFG | 0.0010 | -0.0013 | 0.0032 | |
| dif loneliness | | | LIFG~LIFG | 0.0002 | -0.0026 | 0.0031 | |
| dif loneliness | | | LTPJ~LTPJ | -0.0035 | -0.0064 | -0.0007 | No overlap |
| dif loneliness | | | LIFG~LTPJ | 0.0010 | -0.0013 | 0.0032 | |
| dif loneliness | | | RIFG~RIFG | 0.0010 | -0.0022 | 0.0043 | |
| dif loneliness | | | RTPJ~RTPJ | -0.0018 | -0.0072 | 0.0035 | |
| dif loneliness | | | RIFG~LTPJ | -0.0002 | -0.0024 | 0.0019 | |
| dif loneliness | | | RIFG~RTPJ | 0.0011 | -0.0015 | 0.0039 | |
| *Group effect* | | | | | | | |
| dif loneliness | Intergen | | LIFG~RIFG | 0.0007 | -0.0029 | 0.0041 | |
| dif loneliness | Samegen | | LIFG~RIFG | 0.0013 | -0.0012 | 0.0040 | |
| dif loneliness | Intergen | | LIFG~LIFG | 0.0013 | -0.0033 | 0.0059 | |
| dif loneliness | Samegen | | LIFG~LIFG | -0.0009 | -0.0043 | 0.0026 | |
| dif loneliness | Intergen | | LTPJ~LTPJ | -0.0051 | -0.0098 | -0.0005 | No overlap |
| dif loneliness | Samegen | | LTPJ~LTPJ | -0.0019 | -0.0055 | 0.0014 | |
| dif loneliness | Intergen | | LIFG~LTPJ | 0.0009 | -0.0026 | 0.0044 | |
| dif loneliness | Samegen | | LIFG~LTPJ | 0.0010 | -0.0017 | 0.0039 | |
| dif loneliness | Intergen | | RIFG~RIFG | 0.0004 | -0.0047 | 0.0059 | |
| dif loneliness | Samegen | | RIFG~RIFG | 0.0016 | -0.0019 | 0.0054 | |
| dif loneliness | Intergen | | RTPJ~RTPJ | -0.0027 | -0.0117 | 0.0061 | |
| dif loneliness | Samegen | | RTPJ~RTPJ | -0.0009 | -0.0067 | 0.0051 | |
| dif loneliness | Intergen | | RIFG~LTPJ | -0.0015 | -0.0050 | 0.0020 | |
| dif loneliness | Samegen | | RIFG~LTPJ | 0.0011 | -0.0013 | 0.0037 | |
| dif loneliness | Intergen | | RIFG~RTPJ | -0.0005 | -0.0049 | 0.0039 | |
| dif loneliness | Samegen | | RIFG~RTPJ | 0.0027 | -0.0004 | 0.0058 | |
| *Task effect* | | | | | | | |
| dif loneliness | | Alone | LIFG~RIFG | 0.0016 | -0.0016 | 0.0049 | |
| dif loneliness | | Together | LIFG~RIFG | 0.0004 | -0.0022 | 0.0031 | |
| dif loneliness | | Alone | LIFG~LIFG | 0.0007 | -0.0039 | 0.0049 | |
| dif loneliness | | Together | LIFG~LIFG | -0.0002 | -0.0037 | 0.0034 | |
| dif loneliness | | Alone | LTPJ~LTPJ | -0.0029 | -0.0074 | 0.0015 | |
| dif loneliness | | Together | LTPJ~LTPJ | -0.0042 | -0.0077 | -0.0006 | No overlap |
| dif loneliness | | Alone | LIFG~LTPJ | 0.0019 | -0.0014 | 0.0052 | |
| dif loneliness | | Together | LIFG~LTPJ | 0.0000 | -0.0028 | 0.0026 | |
| dif loneliness | | Alone | RIFG~RIFG | 0.0037 | -0.0011 | 0.0087 | |
| dif loneliness | | Together | RIFG~RIFG | -0.0017 | -0.0056 | 0.0022 | |
| dif loneliness | | Alone | RTPJ~RTPJ | -0.0041 | -0.0124 | 0.0038 | |
| dif loneliness | | Together | RTPJ~RTPJ | 0.0006 | -0.0062 | 0.0072 | |
| dif loneliness | | Alone | RIFG~LTPJ | 0.0007 | -0.0025 | 0.0040 | |
| dif loneliness | | Together | RIFG~LTPJ | -0.0011 | -0.0038 | 0.0014 | |
| dif loneliness | | Alone | RIFG~RTPJ | 0.0037 | -0.0004 | 0.0079 | |
| dif loneliness | | Together | RIFG~RTPJ | -0.0015 | -0.0048 | 0.0018 | |

| Measure | Group | Task | ROI Pairs | Estimate | Lower HPD | Upper HPD | No overlap |
|---|---|---|---|---|---|---|---|
| *Main effect* | | | | | | | |
| dif loneliness | Intergen | Alone | LIFG~RIFG | 0.0006 | -0.0045 | 0.0060 | |
| dif loneliness | Intergen | Together | LIFG~RIFG | 0.0007 | -0.0036 | 0.0049 | |
| dif loneliness | Samegen | Alone | LIFG~RIFG | 0.0025 | -0.0013 | 0.0063 | |
| dif loneliness | Samegen | Together | LIFG~RIFG | 0.0001 | -0.0029 | 0.0032 | |
| dif loneliness | Intergen | Alone | LIFG~LIFG | 0.0012 | -0.0058 | 0.0082 | |
| dif loneliness | Intergen | Together | LIFG~LIFG | 0.0014 | -0.0044 | 0.0072 | |
| dif loneliness | Samegen | Alone | LIFG~LIFG | 0.0001 | -0.0053 | 0.0055 | |
| dif loneliness | Samegen | Together | LIFG~LIFG | -0.0018 | -0.0059 | 0.0024 | |
| dif loneliness | Intergen | Alone | LTPJ~LTPJ | -0.0048 | -0.0117 | 0.0025 | |
| dif loneliness | Intergen | Together | LTPJ~LTPJ | -0.0054 | -0.0113 | 0.0002 | |
| dif loneliness | Samegen | Alone | LTPJ~LTPJ | -0.0009 | -0.0062 | 0.0045 | |
| dif loneliness | Samegen | Together | LTPJ~LTPJ | -0.0029 | -0.0070 | 0.0015 | |
| dif loneliness | Intergen | Alone | LIFG~LTPJ | 0.0009 | -0.0044 | 0.0060 | |
| dif loneliness | Intergen | Together | LIFG~LTPJ | 0.0009 | -0.0035 | 0.0051 | |
| dif loneliness | Samegen | Alone | LIFG~LTPJ | 0.0030 | -0.0011 | 0.0069 | |
| dif loneliness | Samegen | Together | LIFG~LTPJ | -0.0009 | -0.0041 | 0.0023 | |
| dif loneliness | Intergen | Alone | RIFG~RIFG | 0.0049 | -0.0031 | 0.0129 | |
| dif loneliness | Intergen | Together | RIFG~RIFG | -0.0041 | -0.0106 | 0.0023 | |
| dif loneliness | Samegen | Alone | RIFG~RIFG | 0.0024 | -0.0031 | 0.0079 | |
| dif loneliness | Samegen | Together | RIFG~RIFG | 0.0008 | -0.0036 | 0.0051 | |
| dif loneliness | Intergen | Alone | RTPJ~RTPJ | -0.0050 | -0.0179 | 0.0087 | |
| dif loneliness | Intergen | Together | RTPJ~RTPJ | -0.0005 | -0.0121 | 0.0108 | |
| dif loneliness | Samegen | Alone | RTPJ~RTPJ | -0.0033 | -0.0123 | 0.0061 | |
| dif loneliness | Samegen | Together | RTPJ~RTPJ | 0.0016 | -0.0054 | 0.0086 | |
| dif loneliness | Intergen | Alone | RIFG~LTPJ | -0.0009 | -0.0063 | 0.0045 | |
| dif loneliness | Intergen | Together | RIFG~LTPJ | -0.0022 | -0.0065 | 0.0020 | |
| dif loneliness | Samegen | Alone | RIFG~LTPJ | 0.0023 | -0.0015 | 0.0061 | |
| dif loneliness | Samegen | Together | RIFG~LTPJ | -0.0001 | -0.0031 | 0.0029 | |
| dif loneliness | Intergen | Alone | RIFG~RTPJ | 0.0030 | -0.0039 | 0.0097 | |
| dif loneliness | Intergen | Together | RIFG~RTPJ | -0.0039 | -0.0094 | 0.0016 | |
| dif loneliness | Samegen | Alone | RIFG~RTPJ | 0.0044 | -0.0003 | 0.0091 | |
| dif loneliness | Samegen | Together | RIFG~RTPJ | 0.0009 | -0.0028 | 0.0046 | |

**S6 Appendix – Table M.** HbR. Contrasts pertaining to loneliness (dif) ~ INS relationship in unstandardised units. Contrasts are presented in the following order: Group, task, and interaction contrasts.

| Measure | Group | Task | ROI Pairs | Estimate | Lower HPD | Upper HPD | No overlap |
|---|---|---|---|---|---|---|---|
| *Group contrast* | | | | | | | |
| dif loneliness | Intergen-Samegen | | LIFG~RIFG | -0.0007 | -0.0050 | 0.0037 | |
| dif loneliness | Intergen-Samegen | | LIFG~LIFG | 0.0021 | -0.0035 | 0.0080 | |
| dif loneliness | Intergen-Samegen | | LTPJ~LTPJ | -0.0031 | -0.0088 | 0.0027 | |
| dif loneliness | Intergen-Samegen | | LIFG~LTPJ | -0.0002 | -0.0047 | 0.0042 | |
| dif loneliness | Intergen-Samegen | | RIFG~RIFG | -0.0012 | -0.0076 | 0.0052 | |
| dif loneliness | Intergen-Samegen | | RTPJ~RTPJ | -0.0018 | -0.0126 | 0.0088 | |
| dif loneliness | Intergen-Samegen | | RIFG~LTPJ | -0.0026 | -0.0069 | 0.0016 | |
| dif loneliness | Intergen-Samegen | | RIFG~RTPJ | -0.0031 | -0.0084 | 0.0024 | |
| *Task contrast* | | | | | | | |
| dif loneliness | | Alone-Together | LIFG~RIFG | 0.0012 | -0.0028 | 0.0050 | |
| dif loneliness | | Alone-Together | LIFG~LIFG | 0.0009 | -0.0048 | 0.0064 | |
| dif loneliness | | Alone-Together | LTPJ~LTPJ | 0.0013 | -0.0042 | 0.0071 | |
| dif loneliness | | Alone-Together | LIFG~LTPJ | 0.0019 | -0.0020 | 0.0060 | |
| dif loneliness | | Alone-Together | RIFG~RIFG | 0.0053 | -0.0008 | 0.0115 | |
| dif loneliness | | Alone-Together | RTPJ~RTPJ | -0.0047 | -0.0152 | 0.0055 | |
| dif loneliness | | Alone-Together | RIFG~LTPJ | 0.0019 | -0.0023 | 0.0060 | |
| dif loneliness | | Alone-Together | RIFG~RTPJ | 0.0052 | 0.0001 | 0.0104 | No overlap |
| *Interactions* | | | | | | | |
| dif loneliness | Intergen-Samegen | Alone | LIFG~RIFG | -0.0019 | -0.0087 | 0.0042 | |
| dif loneliness | Intergen-Samegen | Together | LIFG~RIFG | 0.0006 | -0.0047 | 0.0057 | |
| dif loneliness | Intergen-Samegen | Alone | LIFG~LIFG | 0.0011 | -0.0076 | 0.0100 | |
| dif loneliness | Intergen-Samegen | Together | LIFG~LIFG | 0.0032 | -0.0039 | 0.0102 | |
| dif loneliness | Intergen-Samegen | Alone | LTPJ~LTPJ | -0.0038 | -0.0124 | 0.0052 | |
| dif loneliness | Intergen-Samegen | Together | LTPJ~LTPJ | -0.0024 | -0.0096 | 0.0046 | |
| dif loneliness | Intergen-Samegen | Alone | LIFG~LTPJ | -0.0021 | -0.0084 | 0.0047 | |
| dif loneliness | Intergen-Samegen | Together | LIFG~LTPJ | 0.0018 | -0.0038 | 0.0070 | |
| dif loneliness | Intergen-Samegen | Alone | RIFG~RIFG | 0.0025 | -0.0074 | 0.0121 | |
| dif loneliness | Intergen-Samegen | Together | RIFG~RIFG | -0.0048 | -0.0127 | 0.0029 | |
| dif loneliness | Intergen-Samegen | Alone | RTPJ~RTPJ | -0.0016 | -0.0180 | 0.0144 | |
| dif loneliness | Intergen-Samegen | Together | RTPJ~RTPJ | -0.0020 | -0.0155 | 0.0112 | |
| dif loneliness | Intergen-Samegen | Alone | RIFG~LTPJ | -0.0032 | -0.0099 | 0.0033 | |
| dif loneliness | Intergen-Samegen | Together | RIFG~LTPJ | -0.0020 | -0.0073 | 0.0031 | |
| dif loneliness | Intergen-Samegen | Alone | RIFG~RTPJ | -0.0014 | -0.0096 | 0.0068 | |
| dif loneliness | Intergen-Samegen | Together | RIFG~RTPJ | -0.0049 | -0.0115 | 0.0018 | |
| dif loneliness | Intergen | Alone-Together | LIFG~RIFG | -0.0001 | -0.0066 | 0.0063 | |
| dif loneliness | Samegen | Alone-Together | LIFG~RIFG | 0.0024 | -0.0022 | 0.0068 | |
| dif loneliness | Intergen | Alone-Together | LIFG~LIFG | -0.0001 | -0.0093 | 0.0087 | |
| dif loneliness | Samegen | Alone-Together | LIFG~LIFG | 0.0019 | -0.0047 | 0.0085 | |
| dif loneliness | Intergen | Alone-Together | LTPJ~LTPJ | 0.0006 | -0.0087 | 0.0093 | |
| dif loneliness | Samegen | Alone-Together | LTPJ~LTPJ | 0.0020 | -0.0046 | 0.0088 | |
| dif loneliness | Intergen | Alone-Together | LIFG~LTPJ | 0.0000 | -0.0064 | 0.0066 | |
| dif loneliness | Samegen | Alone-Together | LIFG~LTPJ | 0.0038 | -0.0008 | 0.0086 | |

| Measure | Group | Task | ROI Pairs | Estimate | Lower HPD | Upper HPD | No overlap |
|---|---|---|---|---|---|---|---|
| dif loneliness | Intergen | Alone-Together | RIFG~RIFG | 0.0090 | -0.0011 | 0.0191 | |
| dif loneliness | Samegen | Alone-Together | RIFG~RIFG | 0.0017 | -0.0050 | 0.0086 | |
| dif loneliness | Intergen | Alone-Together | RTPJ~RTPJ | -0.0044 | -0.0219 | 0.0128 | |
| dif loneliness | Samegen | Alone-Together | RTPJ~RTPJ | -0.0049 | -0.0162 | 0.0061 | |
| dif loneliness | Intergen | Alone-Together | RIFG~LTPJ | 0.0013 | -0.0056 | 0.0080 | |
| dif loneliness | Samegen | Alone-Together | RIFG~LTPJ | 0.0024 | -0.0023 | 0.0072 | |
| dif loneliness | Intergen | Alone-Together | RIFG~RTPJ | 0.0069 | -0.0014 | 0.0156 | |
| dif loneliness | Samegen | Alone-Together | RIFG~RTPJ | 0.0034 | -0.0022 | 0.0093 | |

**S6 Appendix – Table N.** HbR. Relationships between social closeness (sum) and INS in unstandardised units. Effects are presented in the following order: Main, group, task, and simple effects.

| Measure | Group | Task | ROI Pairs | Estimate | Lower HPD | Upper HPD | No overlap |
|---|---|---|---|---|---|---|---|
| *Main effect* | | | | | | | |
| sum social closeness | | | LIFG~RIFG | 0.0000 | -0.0013 | 0.0013 | |
| sum social closeness | | | LIFG~LIFG | -0.0007 | -0.0025 | 0.0011 | |
| sum social closeness | | | LTPJ~LTPJ | 0.0008 | -0.0009 | 0.0026 | |
| sum social closeness | | | LIFG~LTPJ | 0.0002 | -0.0012 | 0.0016 | |
| sum social closeness | | | RIFG~RIFG | -0.0009 | -0.0028 | 0.0011 | |
| sum social closeness | | | RTPJ~RTPJ | 0.0025 | -0.0007 | 0.0059 | |
| sum social closeness | | | RIFG~LTPJ | 0.0007 | -0.0006 | 0.0020 | |
| sum social closeness | | | RIFG~RTPJ | -0.0004 | -0.0020 | 0.0012 | |
| *Group effect* | | | | | | | |
| sum social closeness | Intergen | | LIFG~RIFG | 0.0000 | -0.0014 | 0.0016 | |
| sum social closeness | Samegen | | LIFG~RIFG | 0.0000 | -0.0022 | 0.0022 | |
| sum social closeness | Intergen | | LIFG~LIFG | 0.0002 | -0.0018 | 0.0021 | |
| sum social closeness | Samegen | | LIFG~LIFG | -0.0016 | -0.0046 | 0.0014 | |
| sum social closeness | Intergen | | LTPJ~LTPJ | -0.0007 | -0.0026 | 0.0013 | |
| sum social closeness | Samegen | | LTPJ~LTPJ | 0.0023 | -0.0006 | 0.0054 | |
| sum social closeness | Intergen | | LIFG~LTPJ | 0.0003 | -0.0012 | 0.0018 | |
| sum social closeness | Samegen | | LIFG~LTPJ | 0.0002 | -0.0020 | 0.0025 | |
| sum social closeness | Intergen | | RIFG~RIFG | 0.0004 | -0.0017 | 0.0026 | |
| sum social closeness | Samegen | | RIFG~RIFG | -0.0022 | -0.0054 | 0.0011 | |
| sum social closeness | Intergen | | RTPJ~RTPJ | 0.0024 | -0.0015 | 0.0062 | |
| sum social closeness | Samegen | | RTPJ~RTPJ | 0.0027 | -0.0027 | 0.0079 | |
| sum social closeness | Intergen | | RIFG~LTPJ | 0.0004 | -0.0011 | 0.0017 | |
| sum social closeness | Samegen | | RIFG~LTPJ | 0.0010 | -0.0011 | 0.0032 | |
| sum social closeness | Intergen | | RIFG~RTPJ | 0.0013 | -0.0005 | 0.0032 | |
| sum social closeness | Samegen | | RIFG~RTPJ | -0.0020 | -0.0047 | 0.0006 | |
| *Task effect* | | | | | | | |
| sum social closeness | | Alone | LIFG~RIFG | -0.0006 | -0.0026 | 0.0014 | |
| sum social closeness | | Together | LIFG~RIFG | 0.0006 | -0.0010 | 0.0022 | |
| sum social closeness | | Alone | LIFG~LIFG | -0.0016 | -0.0043 | 0.0013 | |
| sum social closeness | | Together | LIFG~LIFG | 0.0002 | -0.0020 | 0.0023 | |
| sum social closeness | | Alone | LTPJ~LTPJ | 0.0007 | -0.0021 | 0.0035 | |
| sum social closeness | | Together | LTPJ~LTPJ | 0.0010 | -0.0011 | 0.0032 | |
| sum social closeness | | Alone | LIFG~LTPJ | 0.0001 | -0.0020 | 0.0020 | |
| sum social closeness | | Together | LIFG~LTPJ | 0.0004 | -0.0012 | 0.0022 | |
| sum social closeness | | Alone | RIFG~RIFG | -0.0017 | -0.0046 | 0.0014 | |
| sum social closeness | | Together | RIFG~RIFG | -0.0002 | -0.0025 | 0.0022 | |
| sum social closeness | | Alone | RTPJ~RTPJ | 0.0037 | -0.0015 | 0.0084 | |
| sum social closeness | | Together | RTPJ~RTPJ | 0.0014 | -0.0028 | 0.0055 | |
| sum social closeness | | Alone | RIFG~LTPJ | 0.0004 | -0.0015 | 0.0025 | |
| sum social closeness | | Together | RIFG~LTPJ | 0.0009 | -0.0006 | 0.0026 | |
| sum social closeness | | Alone | RIFG~RTPJ | 0.0010 | -0.0014 | 0.0035 | |
| sum social closeness | | Together | RIFG~RTPJ | -0.0017 | -0.0037 | 0.0002 | |

| Measure | Group | Task | ROI Pairs | Estimate | Lower HPD | Upper HPD | No overlap |
|---|---|---|---|---|---|---|---|
| *Main effect* | | | | | | | |
| sum social closeness | Intergen | Alone | LIFG~RIFG | -0.0006 | -0.0028 | 0.0017 | |
| sum social closeness | Intergen | Together | LIFG~RIFG | 0.0007 | -0.0012 | 0.0025 | |
| sum social closeness | Samegen | Alone | LIFG~RIFG | -0.0006 | -0.0040 | 0.0026 | |
| sum social closeness | Samegen | Together | LIFG~RIFG | 0.0005 | -0.0022 | 0.0031 | |
| sum social closeness | Intergen | Alone | LIFG~LIFG | 0.0006 | -0.0025 | 0.0036 | |
| sum social closeness | Intergen | Together | LIFG~LIFG | -0.0001 | -0.0025 | 0.0023 | |
| sum social closeness | Samegen | Alone | LIFG~LIFG | -0.0037 | -0.0082 | 0.0012 | |
| sum social closeness | Samegen | Together | LIFG~LIFG | 0.0004 | -0.0031 | 0.0041 | |
| sum social closeness | Intergen | Alone | LTPJ~LTPJ | -0.0024 | -0.0055 | 0.0006 | |
| sum social closeness | Intergen | Together | LTPJ~LTPJ | 0.0011 | -0.0013 | 0.0035 | |
| sum social closeness | Samegen | Alone | LTPJ~LTPJ | 0.0037 | -0.0009 | 0.0084 | |
| sum social closeness | Samegen | Together | LTPJ~LTPJ | 0.0010 | -0.0027 | 0.0046 | |
| sum social closeness | Intergen | Alone | LIFG~LTPJ | 0.0010 | -0.0013 | 0.0033 | |
| sum social closeness | Intergen | Together | LIFG~LTPJ | -0.0004 | -0.0022 | 0.0014 | |
| sum social closeness | Samegen | Alone | LIFG~LTPJ | -0.0008 | -0.0042 | 0.0026 | |
| sum social closeness | Samegen | Together | LIFG~LTPJ | 0.0013 | -0.0015 | 0.0040 | |
| sum social closeness | Intergen | Alone | RIFG~RIFG | -0.0003 | -0.0038 | 0.0029 | |
| sum social closeness | Intergen | Together | RIFG~RIFG | 0.0011 | -0.0015 | 0.0037 | |
| sum social closeness | Samegen | Alone | RIFG~RIFG | -0.0030 | -0.0079 | 0.0021 | |
| sum social closeness | Samegen | Together | RIFG~RIFG | -0.0014 | -0.0053 | 0.0025 | |
| sum social closeness | Intergen | Alone | RTPJ~RTPJ | 0.0016 | -0.0039 | 0.0071 | |
| sum social closeness | Intergen | Together | RTPJ~RTPJ | 0.0032 | -0.0019 | 0.0084 | |
| sum social closeness | Samegen | Alone | RTPJ~RTPJ | 0.0058 | -0.0026 | 0.0139 | |
| sum social closeness | Samegen | Together | RTPJ~RTPJ | -0.0004 | -0.0070 | 0.0061 | |
| sum social closeness | Intergen | Alone | RIFG~LTPJ | -0.0002 | -0.0024 | 0.0020 | |
| sum social closeness | Intergen | Together | RIFG~LTPJ | 0.0009 | -0.0009 | 0.0026 | |
| sum social closeness | Samegen | Alone | RIFG~LTPJ | 0.0011 | -0.0023 | 0.0044 | |
| sum social closeness | Samegen | Together | RIFG~LTPJ | 0.0009 | -0.0018 | 0.0036 | |
| sum social closeness | Intergen | Alone | RIFG~RTPJ | 0.0021 | -0.0007 | 0.0049 | |
| sum social closeness | Intergen | Together | RIFG~RTPJ | 0.0005 | -0.0018 | 0.0028 | |
| sum social closeness | Samegen | Alone | RIFG~RTPJ | -0.0001 | -0.0043 | 0.0039 | |
| sum social closeness | Samegen | Together | RIFG~RTPJ | -0.0039 | -0.0072 | -0.0008 | No overlap |

**S6 Appendix – Table O.** HbR. Contrasts pertaining to social closeness (sum) ~ INS relationship in unstandardised units. Contrasts are presented in the following order: Group, task, and interaction contrasts.

| Measure | Group | Task | ROI Pairs | Estimate | Lower HPD | Upper HPD | No overlap |
|---|---|---|---|---|---|---|---|
| *Group contrast* | | | | | | | |
| sum social closeness | Int-Samegen | | LIFG~RIFG | 0.0001 | -0.0025 | 0.0028 | |
| sum social closeness | Int-Samegen | | LIFG~LIFG | 0.0018 | -0.0017 | 0.0054 | |
| sum social closeness | Int-Samegen | | LTPJ~LTPJ | -0.0030 | -0.0067 | 0.0004 | |
| sum social closeness | Int-Samegen | | LIFG~LTPJ | 0.0001 | -0.0027 | 0.0029 | |
| sum social closeness | Int-Samegen | | RIFG~RIFG | 0.0025 | -0.0015 | 0.0064 | |
| sum social closeness | Int-Samegen | | RTPJ~RTPJ | -0.0003 | -0.0069 | 0.0061 | |
| sum social closeness | Int-Samegen | | RIFG~LTPJ | -0.0006 | -0.0032 | 0.0020 | |
| sum social closeness | Int-Samegen | | RIFG~RTPJ | 0.0033 | 0.0001 | 0.0065 | No overlap |
| *Task contrast* | | | | | | | |
| sum social closeness | | Alone-Together | LIFG~RIFG | -0.0012 | -0.0036 | 0.0013 | |
| sum social closeness | | Alone-Together | LIFG~LIFG | -0.0017 | -0.0052 | 0.0018 | |
| sum social closeness | | Alone-Together | LTPJ~LTPJ | -0.0004 | -0.0039 | 0.0030 | |
| sum social closeness | | Alone-Together | LIFG~LTPJ | -0.0004 | -0.0029 | 0.0020 | |
| sum social closeness | | Alone-Together | RIFG~RIFG | -0.0015 | -0.0052 | 0.0022 | |
| sum social closeness | | Alone-Together | RTPJ~RTPJ | 0.0023 | -0.0040 | 0.0088 | |
| sum social closeness | | Alone-Together | RIFG~LTPJ | -0.0005 | -0.0030 | 0.0020 | |
| sum social closeness | | Alone-Together | RIFG~RTPJ | 0.0027 | -0.0004 | 0.0058 | |
| *Interactions* | | | | | | | |
| sum social closeness | Int-Samegen | Alone | LIFG~RIFG | 0.0000 | -0.0039 | 0.0041 | |
| sum social closeness | Int-Samegen | Together | LIFG~RIFG | 0.0002 | -0.0031 | 0.0033 | |
| sum social closeness | Int-Samegen | Alone | LIFG~LIFG | 0.0042 | -0.0013 | 0.0099 | |
| sum social closeness | Int-Samegen | Together | LIFG~LIFG | -0.0005 | -0.0048 | 0.0039 | |
| sum social closeness | Int-Samegen | Alone | LTPJ~LTPJ | -0.0061 | -0.0117 | -0.0006 | No overlap |
| sum social closeness | Int-Samegen | Together | LTPJ~LTPJ | 0.0001 | -0.0042 | 0.0045 | |
| sum social closeness | Int-Samegen | Alone | LIFG~LTPJ | 0.0018 | -0.0022 | 0.0060 | |
| sum social closeness | Int-Samegen | Together | LIFG~LTPJ | -0.0017 | -0.0049 | 0.0016 | |
| sum social closeness | Int-Samegen | Alone | RIFG~RIFG | 0.0027 | -0.0036 | 0.0083 | |
| sum social closeness | Int-Samegen | Together | RIFG~RIFG | 0.0024 | -0.0023 | 0.0071 | |
| sum social closeness | Int-Samegen | Alone | RTPJ~RTPJ | -0.0042 | -0.0141 | 0.0057 | |
| sum social closeness | Int-Samegen | Together | RTPJ~RTPJ | 0.0036 | -0.0049 | 0.0119 | |
| sum social closeness | Int-Samegen | Alone | RIFG~LTPJ | -0.0013 | -0.0052 | 0.0029 | |
| sum social closeness | Int-Samegen | Together | RIFG~LTPJ | 0.0000 | -0.0032 | 0.0032 | |
| sum social closeness | Int-Samegen | Alone | RIFG~RTPJ | 0.0022 | -0.0026 | 0.0074 | |
| sum social closeness | Int-Samegen | Together | RIFG~RTPJ | 0.0044 | 0.0004 | 0.0084 | No overlap |
| sum social closeness | Intergen | Alone-Together | LIFG~RIFG | -0.0013 | -0.0039 | 0.0015 | |
| sum social closeness | Samegen | Alone-Together | LIFG~RIFG | -0.0011 | -0.0050 | 0.0031 | |
| sum social closeness | Intergen | Alone-Together | LIFG~LIFG | 0.0007 | -0.0032 | 0.0045 | |
| sum social closeness | Samegen | Alone-Together | LIFG~LIFG | -0.0041 | -0.0097 | 0.0019 | |
| sum social closeness | Intergen | Alone-Together | LTPJ~LTPJ | -0.0035 | -0.0074 | 0.0002 | |
| sum social closeness | Samegen | Alone-Together | LTPJ~LTPJ | 0.0028 | -0.0030 | 0.0087 | |
| sum social closeness | Intergen | Alone-Together | LIFG~LTPJ | 0.0014 | -0.0013 | 0.0041 | |
| sum social closeness | Samegen | Alone-Together | LIFG~LTPJ | -0.0021 | -0.0063 | 0.0020 | |

| Measure | Group | Task | ROI Pairs | Estimate | Lower HPD | Upper HPD | No overlap |
|---|---|---|---|---|---|---|---|
| sum social closeness | Intergen | Alone-Together | RIFG~RIFG | -0.0014 | -0.0054 | 0.0028 | |
| sum social closeness | Samegen | Alone-Together | RIFG~RIFG | -0.0016 | -0.0080 | 0.0045 | |
| sum social closeness | Intergen | Alone-Together | RTPJ~RTPJ | -0.0016 | -0.0091 | 0.0058 | |
| sum social closeness | Samegen | Alone-Together | RTPJ~RTPJ | 0.0062 | -0.0041 | 0.0166 | |
| sum social closeness | Intergen | Alone-Together | RIFG~LTPJ | -0.0011 | -0.0039 | 0.0017 | |
| sum social closeness | Samegen | Alone-Together | RIFG~LTPJ | 0.0002 | -0.0040 | 0.0044 | |
| sum social closeness | Intergen | Alone-Together | RIFG~RTPJ | 0.0016 | -0.0019 | 0.0052 | |
| sum social closeness | Samegen | Alone-Together | RIFG~RTPJ | 0.0038 | -0.0015 | 0.0087 | |

**S6 Appendix – Table P.** HbR. Relationships between social closeness (dif) and INS in unstandardised units. Effects are presented in the following order: Main, group, task, and simple effects.

| Measure | Group | Task | ROI Pairs | Estimate | Lower HPD | Upper HPD | No overlap |
|---|---|---|---|---|---|---|---|
| *Main effect* | | | | | | | |
| dif social closeness | | | LIFG~RIFG | 0.0019 | -0.0002 | 0.0040 | |
| dif social closeness | | | LIFG~LIFG | 0.0016 | -0.0011 | 0.0045 | |
| dif social closeness | | | LTPJ~LTPJ | 0.0009 | -0.0019 | 0.0037 | |
| dif social closeness | | | LIFG~LTPJ | 0.0012 | -0.0010 | 0.0033 | |
| dif social closeness | | | RIFG~RIFG | 0.0027 | -0.0005 | 0.0058 | |
| dif social closeness | | | RTPJ~RTPJ | -0.0026 | -0.0075 | 0.0023 | |
| dif social closeness | | | RIFG~LTPJ | 0.0003 | -0.0019 | 0.0022 | |
| dif social closeness | | | RIFG~RTPJ | 0.0009 | -0.0017 | 0.0036 | |
| *Group effect* | | | | | | | |
| dif social closeness | Intergen | | LIFG~RIFG | 0.0013 | -0.0015 | 0.0040 | |
| dif social closeness | Samegen | | LIFG~RIFG | 0.0024 | -0.0008 | 0.0056 | |
| dif social closeness | Intergen | | LIFG~LIFG | 0.0001 | -0.0036 | 0.0037 | |
| dif social closeness | Samegen | | LIFG~LIFG | 0.0032 | -0.0009 | 0.0075 | |
| dif social closeness | Intergen | | LTPJ~LTPJ | 0.0004 | -0.0033 | 0.0040 | |
| dif social closeness | Samegen | | LTPJ~LTPJ | 0.0014 | -0.0029 | 0.0056 | |
| dif social closeness | Intergen | | LIFG~LTPJ | 0.0017 | -0.0011 | 0.0045 | |
| dif social closeness | Samegen | | LIFG~LTPJ | 0.0006 | -0.0025 | 0.0040 | |
| dif social closeness | Intergen | | RIFG~RIFG | 0.0047 | 0.0006 | 0.0087 | No overlap |
| dif social closeness | Samegen | | RIFG~RIFG | 0.0007 | -0.0042 | 0.0054 | |
| dif social closeness | Intergen | | RTPJ~RTPJ | -0.0063 | -0.0137 | 0.0009 | |
| dif social closeness | Samegen | | RTPJ~RTPJ | 0.0010 | -0.0054 | 0.0074 | |
| dif social closeness | Intergen | | RIFG~LTPJ | 0.0023 | -0.0003 | 0.0051 | |
| dif social closeness | Samegen | | RIFG~LTPJ | -0.0019 | -0.0049 | 0.0014 | |
| dif social closeness | Intergen | | RIFG~RTPJ | -0.0005 | -0.0040 | 0.0031 | |
| dif social closeness | Samegen | | RIFG~RTPJ | 0.0023 | -0.0016 | 0.0061 | |
| *Task effect* | | | | | | | |
| dif social closeness | | Alone | LIFG~RIFG | 0.0042 | 0.0011 | 0.0074 | No overlap |
| dif social closeness | | Together | LIFG~RIFG | -0.0004 | -0.0030 | 0.0020 | |
| dif social closeness | | Alone | LIFG~LIFG | 0.0039 | -0.0005 | 0.0082 | |
| dif social closeness | | Together | LIFG~LIFG | -0.0006 | -0.0039 | 0.0028 | |
| dif social closeness | | Alone | LTPJ~LTPJ | 0.0006 | -0.0038 | 0.0049 | |
| dif social closeness | | Together | LTPJ~LTPJ | 0.0012 | -0.0022 | 0.0046 | |
| dif social closeness | | Alone | LIFG~LTPJ | 0.0015 | -0.0018 | 0.0046 | |
| dif social closeness | | Together | LIFG~LTPJ | 0.0009 | -0.0017 | 0.0034 | |
| dif social closeness | | Alone | RIFG~RIFG | 0.0038 | -0.0012 | 0.0084 | |
| dif social closeness | | Together | RIFG~RIFG | 0.0016 | -0.0022 | 0.0054 | |
| dif social closeness | | Alone | RTPJ~RTPJ | -0.0046 | -0.0123 | 0.0025 | |
| dif social closeness | | Together | RTPJ~RTPJ | -0.0007 | -0.0067 | 0.0053 | |
| dif social closeness | | Alone | RIFG~LTPJ | 0.0007 | -0.0025 | 0.0040 | |
| dif social closeness | | Together | RIFG~LTPJ | -0.0002 | -0.0027 | 0.0023 | |
| dif social closeness | | Alone | RIFG~RTPJ | 0.0014 | -0.0027 | 0.0054 | |
| dif social closeness | | Together | RIFG~RTPJ | 0.0004 | -0.0027 | 0.0036 | |

| Measure | Group | Task | ROI Pairs | Estimate | Lower HPD | Upper HPD | No overlap |
|---|---|---|---|---|---|---|---|
| *Simple effect* | | | | | | | |
| dif social closeness | Intergen | Alone | LIFG~RIFG | 0.0032 | -0.0011 | 0.0073 | |
| dif social closeness | Intergen | Together | LIFG~RIFG | -0.0006 | -0.0038 | 0.0027 | |
| dif social closeness | Samegen | Alone | LIFG~RIFG | 0.0052 | 0.0004 | 0.0099 | No overlap |
| dif social closeness | Samegen | Together | LIFG~RIFG | -0.0002 | -0.0041 | 0.0035 | |
| dif social closeness | Intergen | Alone | LIFG~LIFG | 0.0017 | -0.0040 | 0.0075 | |
| dif social closeness | Intergen | Together | LIFG~LIFG | -0.0015 | -0.0057 | 0.0030 | |
| dif social closeness | Samegen | Alone | LIFG~LIFG | 0.0061 | -0.0004 | 0.0126 | |
| dif social closeness | Samegen | Together | LIFG~LIFG | 0.0002 | -0.0050 | 0.0054 | |
| dif social closeness | Intergen | Alone | LTPJ~LTPJ | 0.0028 | -0.0029 | 0.0084 | |
| dif social closeness | Intergen | Together | LTPJ~LTPJ | -0.0021 | -0.0065 | 0.0023 | |
| dif social closeness | Samegen | Alone | LTPJ~LTPJ | -0.0016 | -0.0083 | 0.0049 | |
| dif social closeness | Samegen | Together | LTPJ~LTPJ | 0.0044 | -0.0008 | 0.0096 | |
| dif social closeness | Intergen | Alone | LIFG~LTPJ | 0.0030 | -0.0013 | 0.0072 | |
| dif social closeness | Intergen | Together | LIFG~LTPJ | 0.0005 | -0.0028 | 0.0039 | |
| dif social closeness | Samegen | Alone | LIFG~LTPJ | 0.0000 | -0.0047 | 0.0049 | |
| dif social closeness | Samegen | Together | LIFG~LTPJ | 0.0013 | -0.0026 | 0.0051 | |
| dif social closeness | Intergen | Alone | RIFG~RIFG | 0.0047 | -0.0016 | 0.0110 | |
| dif social closeness | Intergen | Together | RIFG~RIFG | 0.0047 | 0.0000 | 0.0096 | |
| dif social closeness | Samegen | Alone | RIFG~RIFG | 0.0029 | -0.0046 | 0.0100 | |
| dif social closeness | Samegen | Together | RIFG~RIFG | -0.0014 | -0.0071 | 0.0048 | |
| dif social closeness | Intergen | Alone | RTPJ~RTPJ | -0.0042 | -0.0153 | 0.0066 | |
| dif social closeness | Intergen | Together | RTPJ~RTPJ | -0.0084 | -0.0176 | 0.0010 | |
| dif social closeness | Samegen | Alone | RTPJ~RTPJ | -0.0050 | -0.0148 | 0.0048 | |
| dif social closeness | Samegen | Together | RTPJ~RTPJ | 0.0071 | -0.0006 | 0.0146 | |
| dif social closeness | Intergen | Alone | RIFG~LTPJ | 0.0018 | -0.0024 | 0.0059 | |
| dif social closeness | Intergen | Together | RIFG~LTPJ | 0.0029 | -0.0003 | 0.0062 | |
| dif social closeness | Samegen | Alone | RIFG~LTPJ | -0.0004 | -0.0053 | 0.0045 | |
| dif social closeness | Samegen | Together | RIFG~LTPJ | -0.0033 | -0.0073 | 0.0005 | |
| dif social closeness | Intergen | Alone | RIFG~RTPJ | -0.0010 | -0.0066 | 0.0044 | |
| dif social closeness | Intergen | Together | RIFG~RTPJ | 0.0001 | -0.0044 | 0.0043 | |
| dif social closeness | Samegen | Alone | RIFG~RTPJ | 0.0038 | -0.0021 | 0.0096 | |
| dif social closeness | Samegen | Together | RIFG~RTPJ | 0.0008 | -0.0037 | 0.0055 | |

**Supplementary Materials:** Moffat, Dumas and Cross (2026). *Social interactions between people of same and different generations shape longitudinal changes in interpersonal neural synchrony, loneliness, and social connection.*

**S6 Appendix – Table Q.** HbR. Contrasts pertaining to social closeness (dif) ~ INS relationship in unstandardised units. Contrasts are presented in the following order: Group, task, and interaction contrasts.

| Measure | Group | Task | ROI Pairs | Estimate | Lower HPD | Upper HPD | No overlap |
|---|---|---|---|---|---|---|---|
| *Group contrast* | | | | | | | |
| dif social closeness | Int-Samegen | | LIFG~RIFG | -0.0012 | -0.0053 | 0.0030 | |
| dif social closeness | Int-Samegen | | LIFG~LIFG | -0.0031 | -0.0087 | 0.0024 | |
| dif social closeness | Int-Samegen | | LTPJ~LTPJ | -0.0011 | -0.0067 | 0.0044 | |
| dif social closeness | Int-Samegen | | LIFG~LTPJ | 0.0011 | -0.0031 | 0.0054 | |
| dif social closeness | Int-Samegen | | RIFG~RIFG | 0.0039 | -0.0022 | 0.0104 | |
| dif social closeness | Int-Samegen | | RTPJ~RTPJ | -0.0073 | -0.0169 | 0.0024 | |
| dif social closeness | Int-Samegen | | RIFG~LTPJ | 0.0042 | -0.0001 | 0.0083 | |
| dif social closeness | Int-Samegen | | RIFG~RTPJ | -0.0027 | -0.0080 | 0.0024 | |
| *Task contrast* | | | | | | | |
| dif social closeness | | Alone-Together | LIFG~RIFG | 0.0045 | 0.0006 | 0.0084 | No overlap |
| dif social closeness | | Alone-Together | LIFG~LIFG | 0.0045 | -0.0009 | 0.0101 | |
| dif social closeness | | Alone-Together | LTPJ~LTPJ | -0.0005 | -0.0060 | 0.0050 | |
| dif social closeness | | Alone-Together | LIFG~LTPJ | 0.0006 | -0.0033 | 0.0045 | |
| dif social closeness | | Alone-Together | RIFG~RIFG | 0.0022 | -0.0038 | 0.0082 | |
| dif social closeness | | Alone-Together | RTPJ~RTPJ | -0.0040 | -0.0132 | 0.0055 | |
| dif social closeness | | Alone-Together | RIFG~LTPJ | 0.0009 | -0.0031 | 0.0050 | |
| dif social closeness | | Alone-Together | RIFG~RTPJ | 0.0009 | -0.0040 | 0.0061 | |
| *Interactions* | | | | | | | |
| dif social closeness | Int-Samegen | Alone | LIFG~RIFG | -0.0020 | -0.0084 | 0.0043 | |
| dif social closeness | Int-Samegen | Together | LIFG~RIFG | -0.0003 | -0.0053 | 0.0047 | |
| dif social closeness | Int-Samegen | Alone | LIFG~LIFG | -0.0044 | -0.0132 | 0.0042 | |
| dif social closeness | Int-Samegen | Together | LIFG~LIFG | -0.0017 | -0.0084 | 0.0051 | |
| dif social closeness | Int-Samegen | Alone | LTPJ~LTPJ | 0.0044 | -0.0045 | 0.0130 | |
| dif social closeness | Int-Samegen | Together | LTPJ~LTPJ | -0.0065 | -0.0131 | 0.0004 | |
| dif social closeness | Int-Samegen | Alone | LIFG~LTPJ | 0.0029 | -0.0035 | 0.0094 | |
| dif social closeness | Int-Samegen | Together | LIFG~LTPJ | -0.0008 | -0.0058 | 0.0045 | |
| dif social closeness | Int-Samegen | Alone | RIFG~RIFG | 0.0018 | -0.0080 | 0.0116 | |
| dif social closeness | Int-Samegen | Together | RIFG~RIFG | 0.0061 | -0.0016 | 0.0137 | |
| dif social closeness | Int-Samegen | Alone | RTPJ~RTPJ | 0.0007 | -0.0143 | 0.0151 | |
| dif social closeness | Int-Samegen | Together | RTPJ~RTPJ | -0.0155 | -0.0274 | -0.0032 | No overlap |
| dif social closeness | Int-Samegen | Alone | RIFG~LTPJ | 0.0021 | -0.0042 | 0.0087 | |
| dif social closeness | Int-Samegen | Together | RIFG~LTPJ | 0.0063 | 0.0011 | 0.0112 | No overlap |
| dif social closeness | Int-Samegen | Alone | RIFG~RTPJ | -0.0049 | -0.0130 | 0.0031 | |
| dif social closeness | Int-Samegen | Together | RIFG~RTPJ | -0.0006 | -0.0068 | 0.0059 | |
| dif social closeness | Intergen | Alone-Together | LIFG~RIFG | 0.0037 | -0.0013 | 0.0088 | |
| dif social closeness | Samegen | Alone-Together | LIFG~RIFG | 0.0054 | -0.0006 | 0.0112 | |
| dif social closeness | Intergen | Alone-Together | LIFG~LIFG | 0.0032 | -0.0041 | 0.0103 | |
| dif social closeness | Samegen | Alone-Together | LIFG~LIFG | 0.0058 | -0.0025 | 0.0139 | |
| dif social closeness | Intergen | Alone-Together | LTPJ~LTPJ | 0.0049 | -0.0020 | 0.0122 | |
| dif social closeness | Samegen | Alone-Together | LTPJ~LTPJ | -0.0059 | -0.0142 | 0.0023 | |
| dif social closeness | Intergen | Alone-Together | LIFG~LTPJ | 0.0024 | -0.0027 | 0.0076 | |
| dif social closeness | Samegen | Alone-Together | LIFG~LTPJ | -0.0012 | -0.0072 | 0.0047 | |

| Measure | Group | Task | ROI Pairs | Estimate | Lower HPD | Upper HPD | No overlap |
|---|---|---|---|---|---|---|---|
| dif social closeness | Intergen | Alone-Together | RIFG~RIFG | 0.0000 | -0.0076 | 0.0080 | |
| dif social closeness | Samegen | Alone-Together | RIFG~RIFG | 0.0043 | -0.0050 | 0.0134 | |
| dif social closeness | Intergen | Alone-Together | RTPJ~RTPJ | 0.0041 | -0.0095 | 0.0190 | |
| dif social closeness | Samegen | Alone-Together | RTPJ~RTPJ | -0.0120 | -0.0241 | 0.0000 | |
| dif social closeness | Intergen | Alone-Together | RIFG~LTPJ | -0.0011 | -0.0064 | 0.0041 | |
| dif social closeness | Samegen | Alone-Together | RIFG~LTPJ | 0.0030 | -0.0030 | 0.0092 | |
| dif social closeness | Intergen | Alone-Together | RIFG~RTPJ | -0.0012 | -0.0078 | 0.0060 | |
| dif social closeness | Samegen | Alone-Together | RIFG~RTPJ | 0.0031 | -0.0043 | 0.0103 | |

**S6 Appendix – Table R.** HbR. Relationships between allophilia (sum) and INS in unstandardised units. Effects are presented in the following order: Main, group, task, and simple effects.

| Measure | Group | Task | ROI Pairs | Estimate | Lower HPD | Upper HPD | No overlap |
|---|---|---|---|---|---|---|---|
| *Main effect* | | | | | | | |
| sum allophilia | | | LIFG~RIFG | 0.0000 | -0.0002 | 0.0001 | |
| sum allophilia | | | LIFG~LIFG | 0.0001 | -0.0002 | 0.0003 | |
| sum allophilia | | | LTPJ~LTPJ | 0.0001 | -0.0001 | 0.0003 | |
| sum allophilia | | | LIFG~LTPJ | 0.0001 | 0.0000 | 0.0003 | |
| sum allophilia | | | RIFG~RIFG | 0.0000 | -0.0002 | 0.0003 | |
| sum allophilia | | | RTPJ~RTPJ | 0.0000 | -0.0004 | 0.0004 | |
| sum allophilia | | | RIFG~LTPJ | -0.0001 | -0.0002 | 0.0001 | |
| sum allophilia | | | RIFG~RTPJ | 0.0000 | -0.0002 | 0.0002 | |
| *Group effect* | | | | | | | |
| sum allophilia | Intergen | | LIFG~RIFG | 0.0001 | -0.0002 | 0.0003 | |
| sum allophilia | Samegen | | LIFG~RIFG | -0.0001 | -0.0004 | 0.0001 | |
| sum allophilia | Intergen | | LIFG~LIFG | 0.0000 | -0.0002 | 0.0003 | |
| sum allophilia | Samegen | | LIFG~LIFG | 0.0001 | -0.0002 | 0.0004 | |
| sum allophilia | Intergen | | LTPJ~LTPJ | 0.0002 | 0.0000 | 0.0005 | |
| sum allophilia | Samegen | | LTPJ~LTPJ | 0.0000 | -0.0003 | 0.0003 | |
| sum allophilia | Intergen | | LIFG~LTPJ | 0.0001 | -0.0001 | 0.0003 | |
| sum allophilia | Samegen | | LIFG~LTPJ | 0.0001 | -0.0001 | 0.0004 | |
| sum allophilia | Intergen | | RIFG~RIFG | 0.0002 | -0.0001 | 0.0005 | |
| sum allophilia | Samegen | | RIFG~RIFG | -0.0001 | -0.0004 | 0.0002 | |
| sum allophilia | Intergen | | RTPJ~RTPJ | 0.0001 | -0.0005 | 0.0007 | |
| sum allophilia | Samegen | | RTPJ~RTPJ | -0.0001 | -0.0006 | 0.0003 | |
| sum allophilia | Intergen | | RIFG~LTPJ | 0.0000 | -0.0002 | 0.0002 | |
| sum allophilia | Samegen | | RIFG~LTPJ | -0.0002 | -0.0004 | 0.0001 | |
| sum allophilia | Intergen | | RIFG~RTPJ | 0.0000 | -0.0003 | 0.0003 | |
| sum allophilia | Samegen | | RIFG~RTPJ | 0.0001 | -0.0002 | 0.0003 | |
| *Task effect* | | | | | | | |
| sum allophilia | | Alone | LIFG~RIFG | -0.0001 | -0.0003 | 0.0001 | |
| sum allophilia | | Together | LIFG~RIFG | 0.0000 | -0.0002 | 0.0002 | |
| sum allophilia | | Alone | LIFG~LIFG | 0.0001 | -0.0002 | 0.0004 | |
| sum allophilia | | Together | LIFG~LIFG | 0.0000 | -0.0002 | 0.0003 | |
| sum allophilia | | Alone | LTPJ~LTPJ | 0.0003 | 0.0000 | 0.0006 | |
| sum allophilia | | Together | LTPJ~LTPJ | -0.0001 | -0.0003 | 0.0002 | |
| sum allophilia | | Alone | LIFG~LTPJ | 0.0001 | -0.0001 | 0.0003 | |
| sum allophilia | | Together | LIFG~LTPJ | 0.0001 | 0.0000 | 0.0003 | |
| sum allophilia | | Alone | RIFG~RIFG | -0.0001 | -0.0004 | 0.0002 | |
| sum allophilia | | Together | RIFG~RIFG | 0.0002 | -0.0001 | 0.0005 | |
| sum allophilia | | Alone | RTPJ~RTPJ | -0.0004 | -0.0010 | 0.0001 | |
| sum allophilia | | Together | RTPJ~RTPJ | 0.0004 | -0.0001 | 0.0009 | |
| sum allophilia | | Alone | RIFG~LTPJ | -0.0001 | -0.0003 | 0.0001 | |
| sum allophilia | | Together | RIFG~LTPJ | -0.0001 | -0.0003 | 0.0001 | |
| sum allophilia | | Alone | RIFG~RTPJ | -0.0001 | -0.0004 | 0.0002 | |
| sum allophilia | | Together | RIFG~RTPJ | 0.0001 | -0.0001 | 0.0004 | |

| Measure | Group | Task | ROI Pairs | Estimate | Lower HPD | Upper HPD | No overlap |
|---|---|---|---|---|---|---|---|
| *Simple effect* | | | | | | | |
| sum allophilia | Intergen | Alone | LIFG~RIFG | 0.0000 | -0.0003 | 0.0004 | |
| sum allophilia | Intergen | Together | LIFG~RIFG | 0.0001 | -0.0002 | 0.0003 | |
| sum allophilia | Samegen | Alone | LIFG~RIFG | -0.0003 | -0.0006 | 0.0001 | |
| sum allophilia | Samegen | Together | LIFG~RIFG | 0.0000 | -0.0003 | 0.0002 | |
| sum allophilia | Intergen | Alone | LIFG~LIFG | -0.0001 | -0.0005 | 0.0003 | |
| sum allophilia | Intergen | Together | LIFG~LIFG | 0.0001 | -0.0002 | 0.0005 | |
| sum allophilia | Samegen | Alone | LIFG~LIFG | 0.0002 | -0.0002 | 0.0007 | |
| sum allophilia | Samegen | Together | LIFG~LIFG | -0.0001 | -0.0004 | 0.0003 | |
| sum allophilia | Intergen | Alone | LTPJ~LTPJ | 0.0006 | 0.0002 | 0.0010 | No overlap |
| sum allophilia | Intergen | Together | LTPJ~LTPJ | -0.0001 | -0.0004 | 0.0002 | |
| sum allophilia | Samegen | Alone | LTPJ~LTPJ | 0.0000 | -0.0005 | 0.0004 | |
| sum allophilia | Samegen | Together | LTPJ~LTPJ | 0.0000 | -0.0003 | 0.0003 | |
| sum allophilia | Intergen | Alone | LIFG~LTPJ | 0.0000 | -0.0003 | 0.0003 | |
| sum allophilia | Intergen | Together | LIFG~LTPJ | 0.0002 | -0.0001 | 0.0004 | |
| sum allophilia | Samegen | Alone | LIFG~LTPJ | 0.0002 | -0.0002 | 0.0005 | |
| sum allophilia | Samegen | Together | LIFG~LTPJ | 0.0001 | -0.0002 | 0.0004 | |
| sum allophilia | Intergen | Alone | RIFG~RIFG | 0.0000 | -0.0004 | 0.0005 | |
| sum allophilia | Intergen | Together | RIFG~RIFG | 0.0004 | 0.0000 | 0.0007 | |
| sum allophilia | Samegen | Alone | RIFG~RIFG | -0.0002 | -0.0007 | 0.0003 | |
| sum allophilia | Samegen | Together | RIFG~RIFG | 0.0000 | -0.0004 | 0.0004 | |
| sum allophilia | Intergen | Alone | RTPJ~RTPJ | -0.0001 | -0.0010 | 0.0007 | |
| sum allophilia | Intergen | Together | RTPJ~RTPJ | 0.0004 | -0.0004 | 0.0011 | |
| sum allophilia | Samegen | Alone | RTPJ~RTPJ | -0.0007 | -0.0014 | 0.0000 | No overlap |
| sum allophilia | Samegen | Together | RTPJ~RTPJ | 0.0005 | -0.0001 | 0.0010 | |
| sum allophilia | Intergen | Alone | RIFG~LTPJ | 0.0000 | -0.0003 | 0.0003 | |
| sum allophilia | Intergen | Together | RIFG~LTPJ | 0.0000 | -0.0002 | 0.0002 | |
| sum allophilia | Samegen | Alone | RIFG~LTPJ | -0.0002 | -0.0005 | 0.0002 | |
| sum allophilia | Samegen | Together | RIFG~LTPJ | -0.0001 | -0.0004 | 0.0001 | |
| sum allophilia | Intergen | Alone | RIFG~RTPJ | -0.0001 | -0.0005 | 0.0004 | |
| sum allophilia | Intergen | Together | RIFG~RTPJ | 0.0000 | -0.0003 | 0.0003 | |
| sum allophilia | Samegen | Alone | RIFG~RTPJ | -0.0001 | -0.0005 | 0.0003 | |
| sum allophilia | Samegen | Together | RIFG~RTPJ | 0.0002 | -0.0001 | 0.0005 | |

**S6 Appendix – Table S.** HbR. Contrasts pertaining to allophilia (sum) ~ INS relationship in unstandardised units. Contrasts are presented in the following order: Group, task, and interaction contrasts.

| Measure | Group | Task | ROI Pairs | Estimate | Lower HPD | Upper HPD | No overlap |
|---|---|---|---|---|---|---|---|
| *Group contrast* | | | | | | | |
| sum allophilia | Int-Samegen | | LIFG~RIFG | 0.0002 | -0.0001 | 0.0005 | |
| sum allophilia | Int-Samegen | | LIFG~LIFG | -0.0001 | -0.0004 | 0.0004 | |
| sum allophilia | Int-Samegen | | LTPJ~LTPJ | 0.0003 | -0.0001 | 0.0007 | |
| sum allophilia | Int-Samegen | | LIFG~LTPJ | 0.0000 | -0.0004 | 0.0003 | |
| sum allophilia | Int-Samegen | | RIFG~RIFG | 0.0003 | -0.0001 | 0.0008 | |
| sum allophilia | Int-Samegen | | RTPJ~RTPJ | 0.0002 | -0.0005 | 0.0010 | |
| sum allophilia | Int-Samegen | | RIFG~LTPJ | 0.0002 | -0.0001 | 0.0005 | |
| sum allophilia | Int-Samegen | | RIFG~RTPJ | -0.0001 | -0.0005 | 0.0003 | |
| *Task contrast* | | | | | | | |
| sum allophilia | | Alone-Together | LIFG~RIFG | -0.0001 | -0.0004 | 0.0001 | |
| sum allophilia | | Alone-Together | LIFG~LIFG | 0.0001 | -0.0004 | 0.0004 | |
| sum allophilia | | Alone-Together | LTPJ~LTPJ | 0.0003 | 0.0000 | 0.0007 | |
| sum allophilia | | Alone-Together | LIFG~LTPJ | 0.0000 | -0.0003 | 0.0002 | |
| sum allophilia | | Alone-Together | RIFG~RIFG | -0.0003 | -0.0007 | 0.0001 | |
| sum allophilia | | Alone-Together | RTPJ~RTPJ | -0.0008 | -0.0015 | -0.0001 | No overlap |
| sum allophilia | | Alone-Together | RIFG~LTPJ | 0.0000 | -0.0003 | 0.0003 | |
| sum allophilia | | Alone-Together | RIFG~RTPJ | -0.0002 | -0.0005 | 0.0002 | |
| *Interactions* | | | | | | | |
| sum allophilia | Int-Samegen | Alone | LIFG~RIFG | 0.0003 | -0.0002 | 0.0008 | |
| sum allophilia | Int-Samegen | Together | LIFG~RIFG | 0.0001 | -0.0003 | 0.0005 | |
| sum allophilia | Int-Samegen | Alone | LIFG~LIFG | -0.0003 | -0.0009 | 0.0003 | |
| sum allophilia | Int-Samegen | Together | LIFG~LIFG | 0.0002 | -0.0003 | 0.0007 | |
| sum allophilia | Int-Samegen | Alone | LTPJ~LTPJ | 0.0006 | 0.0000 | 0.0012 | |
| sum allophilia | Int-Samegen | Together | LTPJ~LTPJ | -0.0001 | -0.0006 | 0.0004 | |
| sum allophilia | Int-Samegen | Alone | LIFG~LTPJ | -0.0002 | -0.0007 | 0.0003 | |
| sum allophilia | Int-Samegen | Together | LIFG~LTPJ | 0.0001 | -0.0003 | 0.0005 | |
| sum allophilia | Int-Samegen | Alone | RIFG~RIFG | 0.0003 | -0.0004 | 0.0010 | |
| sum allophilia | Int-Samegen | Together | RIFG~RIFG | 0.0003 | -0.0002 | 0.0009 | |
| sum allophilia | Int-Samegen | Alone | RTPJ~RTPJ | 0.0006 | -0.0005 | 0.0017 | |
| sum allophilia | Int-Samegen | Together | RTPJ~RTPJ | -0.0001 | -0.0010 | 0.0008 | |
| sum allophilia | Int-Samegen | Alone | RIFG~LTPJ | 0.0002 | -0.0003 | 0.0006 | |
| sum allophilia | Int-Samegen | Together | RIFG~LTPJ | 0.0001 | -0.0002 | 0.0005 | |
| sum allophilia | Int-Samegen | Alone | RIFG~RTPJ | 0.0000 | -0.0006 | 0.0006 | |
| sum allophilia | Int-Samegen | Together | RIFG~RTPJ | -0.0002 | -0.0007 | 0.0003 | |
| sum allophilia | Intergen | Alone-Together | LIFG~RIFG | 0.0000 | -0.0004 | 0.0003 | |
| sum allophilia | Samegen | Alone-Together | LIFG~RIFG | -0.0002 | -0.0006 | 0.0002 | |
| sum allophilia | Intergen | Alone-Together | LIFG~LIFG | -0.0002 | -0.0007 | 0.0003 | |
| sum allophilia | Samegen | Alone-Together | LIFG~LIFG | 0.0003 | -0.0003 | 0.0009 | |
| sum allophilia | Intergen | Alone-Together | LTPJ~LTPJ | 0.0007 | 0.0002 | 0.0012 | No overlap |
| sum allophilia | Samegen | Alone-Together | LTPJ~LTPJ | 0.0000 | -0.0006 | 0.0005 | |
| sum allophilia | Intergen | Alone-Together | LIFG~LTPJ | -0.0002 | -0.0006 | 0.0002 | |
| sum allophilia | Samegen | Alone-Together | LIFG~LTPJ | 0.0001 | -0.0003 | 0.0005 | |

| Measure | Group | Task | ROI Pairs | Estimate | Lower HPD | Upper HPD | No overlap |
|---|---|---|---|---|---|---|---|
| sum allophilia | Intergen | Alone-Together | RIFG~RIFG | -0.0003 | -0.0009 | 0.0002 | |
| sum allophilia | Samegen | Alone-Together | RIFG~RIFG | -0.0003 | -0.0009 | 0.0004 | |
| sum allophilia | Intergen | Alone-Together | RTPJ~RTPJ | -0.0005 | -0.0016 | 0.0006 | |
| sum allophilia | Samegen | Alone-Together | RTPJ~RTPJ | -0.0012 | -0.0020 | -0.0003 | No overlap |
| sum allophilia | Intergen | Alone-Together | RIFG~LTPJ | 0.0000 | -0.0004 | 0.0004 | |
| sum allophilia | Samegen | Alone-Together | RIFG~LTPJ | 0.0000 | -0.0004 | 0.0004 | |
| sum allophilia | Intergen | Alone-Together | RIFG~RTPJ | -0.0001 | -0.0006 | 0.0004 | |
| sum allophilia | Samegen | Alone-Together | RIFG~RTPJ | -0.0003 | -0.0008 | 0.0002 | |

**S6 Appendix – Table T.** HbR. Relationships between allophilia (dif) and INS in unstandardised units. Effects are presented in the following order: Main, group, task, and simple effects.

| Measure | Group | Task | ROI Pairs | Estimate | Lower HPD | Upper HPD | No overlap |
|---|---|---|---|---|---|---|---|
| *Main effect* | | | | | | | |
| dif allophilia | | | LIFG~RIFG | 0.0000 | -0.0002 | 0.0003 | |
| dif allophilia | | | LIFG~LIFG | 0.0001 | -0.0002 | 0.0003 | |
| dif allophilia | | | LTPJ~LTPJ | 0.0001 | -0.0002 | 0.0003 | |
| dif allophilia | | | LIFG~LTPJ | -0.0001 | -0.0003 | 0.0002 | |
| dif allophilia | | | RIFG~RIFG | 0.0001 | -0.0002 | 0.0004 | |
| dif allophilia | | | RTPJ~RTPJ | -0.0002 | -0.0006 | 0.0003 | |
| dif allophilia | | | RIFG~LTPJ | -0.0001 | -0.0003 | 0.0001 | |
| dif allophilia | | | RIFG~RTPJ | 0.0000 | -0.0002 | 0.0003 | |
| *Group effect* | | | | | | | |
| dif allophilia | Intergen | | LIFG~RIFG | 0.0002 | -0.0001 | 0.0005 | |
| dif allophilia | Samegen | | LIFG~RIFG | -0.0001 | -0.0004 | 0.0002 | |
| dif allophilia | Intergen | | LIFG~LIFG | 0.0002 | -0.0002 | 0.0006 | |
| dif allophilia | Samegen | | LIFG~LIFG | -0.0001 | -0.0005 | 0.0003 | |
| dif allophilia | Intergen | | LTPJ~LTPJ | 0.0004 | 0.0000 | 0.0007 | No overlap |
| dif allophilia | Samegen | | LTPJ~LTPJ | -0.0002 | -0.0006 | 0.0001 | |
| dif allophilia | Intergen | | LIFG~LTPJ | 0.0001 | -0.0002 | 0.0003 | |
| dif allophilia | Samegen | | LIFG~LTPJ | -0.0002 | -0.0005 | 0.0001 | |
| dif allophilia | Intergen | | RIFG~RIFG | -0.0002 | -0.0006 | 0.0002 | |
| dif allophilia | Samegen | | RIFG~RIFG | 0.0004 | -0.0001 | 0.0008 | |
| dif allophilia | Intergen | | RTPJ~RTPJ | 0.0002 | -0.0004 | 0.0007 | |
| dif allophilia | Samegen | | RTPJ~RTPJ | -0.0005 | -0.0011 | 0.0001 | |
| dif allophilia | Intergen | | RIFG~LTPJ | -0.0002 | -0.0005 | 0.0001 | |
| dif allophilia | Samegen | | RIFG~LTPJ | 0.0001 | -0.0002 | 0.0004 | |
| dif allophilia | Intergen | | RIFG~RTPJ | -0.0002 | -0.0005 | 0.0002 | |
| dif allophilia | Samegen | | RIFG~RTPJ | 0.0002 | -0.0002 | 0.0006 | |
| *Task effect* | | | | | | | |
| dif allophilia | | Alone | LIFG~RIFG | -0.0002 | -0.0005 | 0.0001 | |
| dif allophilia | | Together | LIFG~RIFG | 0.0003 | 0.0000 | 0.0005 | |
| dif allophilia | | Alone | LIFG~LIFG | 0.0000 | -0.0004 | 0.0004 | |
| dif allophilia | | Together | LIFG~LIFG | 0.0001 | -0.0002 | 0.0005 | |
| dif allophilia | | Alone | LTPJ~LTPJ | -0.0001 | -0.0005 | 0.0003 | |
| dif allophilia | | Together | LTPJ~LTPJ | 0.0002 | -0.0001 | 0.0006 | |
| dif allophilia | | Alone | LIFG~LTPJ | -0.0002 | -0.0005 | 0.0002 | |
| dif allophilia | | Together | LIFG~LTPJ | 0.0000 | -0.0002 | 0.0003 | |
| dif allophilia | | Alone | RIFG~RIFG | 0.0000 | -0.0005 | 0.0004 | |
| dif allophilia | | Together | RIFG~RIFG | 0.0002 | -0.0002 | 0.0006 | |
| dif allophilia | | Alone | RTPJ~RTPJ | -0.0001 | -0.0007 | 0.0005 | |
| dif allophilia | | Together | RTPJ~RTPJ | -0.0002 | -0.0008 | 0.0003 | |
| dif allophilia | | Alone | RIFG~LTPJ | -0.0001 | -0.0004 | 0.0002 | |
| dif allophilia | | Together | RIFG~LTPJ | 0.0000 | -0.0003 | 0.0002 | |
| dif allophilia | | Alone | RIFG~RTPJ | -0.0001 | -0.0005 | 0.0002 | |
| dif allophilia | | Together | RIFG~RTPJ | 0.0002 | -0.0001 | 0.0005 | |

| Measure | Group | Task | ROI Pairs | Estimate | Lower HPD | Upper HPD | No overlap |
|---|---|---|---|---|---|---|---|
| *Simple effect* | | | | | | | |
| dif allophilia | Intergen | Alone | LIFG~RIFG | 0.0000 | -0.0004 | 0.0004 | |
| dif allophilia | Intergen | Together | LIFG~RIFG | 0.0004 | 0.0000 | 0.0007 | |
| dif allophilia | Samegen | Alone | LIFG~RIFG | -0.0004 | -0.0008 | 0.0001 | |
| dif allophilia | Samegen | Together | LIFG~RIFG | 0.0001 | -0.0002 | 0.0005 | |
| dif allophilia | Intergen | Alone | LIFG~LIFG | 0.0001 | -0.0005 | 0.0007 | |
| dif allophilia | Intergen | Together | LIFG~LIFG | 0.0002 | -0.0002 | 0.0007 | |
| dif allophilia | Samegen | Alone | LIFG~LIFG | -0.0001 | -0.0008 | 0.0005 | |
| dif allophilia | Samegen | Together | LIFG~LIFG | 0.0000 | -0.0004 | 0.0005 | |
| dif allophilia | Intergen | Alone | LTPJ~LTPJ | 0.0001 | -0.0005 | 0.0006 | |
| dif allophilia | Intergen | Together | LTPJ~LTPJ | 0.0007 | 0.0002 | 0.0012 | No overlap |
| dif allophilia | Samegen | Alone | LTPJ~LTPJ | -0.0002 | -0.0008 | 0.0004 | |
| dif allophilia | Samegen | Together | LTPJ~LTPJ | -0.0002 | -0.0007 | 0.0002 | |
| dif allophilia | Intergen | Alone | LIFG~LTPJ | 0.0001 | -0.0003 | 0.0005 | |
| dif allophilia | Intergen | Together | LIFG~LTPJ | 0.0000 | -0.0003 | 0.0004 | |
| dif allophilia | Samegen | Alone | LIFG~LTPJ | -0.0004 | -0.0008 | 0.0001 | |
| dif allophilia | Samegen | Together | LIFG~LTPJ | 0.0000 | -0.0004 | 0.0004 | |
| dif allophilia | Intergen | Alone | RIFG~RIFG | -0.0005 | -0.0011 | 0.0001 | |
| dif allophilia | Intergen | Together | RIFG~RIFG | 0.0001 | -0.0004 | 0.0006 | |
| dif allophilia | Samegen | Alone | RIFG~RIFG | 0.0004 | -0.0002 | 0.0011 | |
| dif allophilia | Samegen | Together | RIFG~RIFG | 0.0003 | -0.0002 | 0.0008 | |
| dif allophilia | Intergen | Alone | RTPJ~RTPJ | -0.0001 | -0.0010 | 0.0007 | |
| dif allophilia | Intergen | Together | RTPJ~RTPJ | 0.0005 | -0.0003 | 0.0012 | |
| dif allophilia | Samegen | Alone | RTPJ~RTPJ | -0.0001 | -0.0010 | 0.0009 | |
| dif allophilia | Samegen | Together | RTPJ~RTPJ | -0.0009 | -0.0017 | -0.0001 | No overlap |
| dif allophilia | Intergen | Alone | RIFG~LTPJ | -0.0002 | -0.0006 | 0.0002 | |
| dif allophilia | Intergen | Together | RIFG~LTPJ | -0.0003 | -0.0006 | 0.0001 | |
| dif allophilia | Samegen | Alone | RIFG~LTPJ | 0.0000 | -0.0004 | 0.0004 | |
| dif allophilia | Samegen | Together | RIFG~LTPJ | 0.0002 | -0.0001 | 0.0005 | |
| dif allophilia | Intergen | Alone | RIFG~RTPJ | -0.0003 | -0.0008 | 0.0002 | |
| dif allophilia | Intergen | Together | RIFG~RTPJ | -0.0001 | -0.0005 | 0.0003 | |
| dif allophilia | Samegen | Alone | RIFG~RTPJ | 0.0000 | -0.0006 | 0.0005 | |
| dif allophilia | Samegen | Together | RIFG~RTPJ | 0.0004 | 0.0000 | 0.0009 | No overlap |

**S6 Appendix – Table U.** HbR. Contrasts pertaining to allophilia (dif) ~ INS relationship in unstandardised units. Contrasts are presented in the following order: Group, task, and interaction contrasts.

| Measure | Group | Task | ROI Pairs | Estimate | Lower HPD | Upper HPD | No overlap |
|---|---|---|---|---|---|---|---|
| *Group contrast* | | | | | | | |
| dif allophilia | Int-Samegen | | LIFG~RIFG | 0.0003 | -0.0001 | 0.0007 | |
| dif allophilia | Int-Samegen | | LIFG~LIFG | 0.0002 | -0.0003 | 0.0008 | |
| dif allophilia | Int-Samegen | | LTPJ~LTPJ | 0.0006 | 0.0001 | 0.0012 | No overlap |
| dif allophilia | Int-Samegen | | LIFG~LTPJ | 0.0003 | -0.0002 | 0.0007 | |
| dif allophilia | Int-Samegen | | RIFG~RIFG | -0.0006 | -0.0012 | 0.0000 | |
| dif allophilia | Int-Samegen | | RTPJ~RTPJ | 0.0006 | -0.0002 | 0.0015 | |
| dif allophilia | Int-Samegen | | RIFG~LTPJ | -0.0003 | -0.0007 | 0.0001 | |
| dif allophilia | Int-Samegen | | RIFG~RTPJ | -0.0004 | -0.0009 | 0.0001 | |
| *Task contrast* | | | | | | | |
| dif allophilia | | Alone-Together | LIFG~RIFG | -0.0004 | -0.0008 | -0.0001 | No overlap |
| dif allophilia | | Alone-Together | LIFG~LIFG | -0.0002 | -0.0007 | 0.0004 | |
| dif allophilia | | Alone-Together | LTPJ~LTPJ | -0.0003 | -0.0008 | 0.0002 | |
| dif allophilia | | Alone-Together | LIFG~LTPJ | -0.0002 | -0.0005 | 0.0002 | |
| dif allophilia | | Alone-Together | RIFG~RIFG | -0.0002 | -0.0008 | 0.0003 | |
| dif allophilia | | Alone-Together | RTPJ~RTPJ | 0.0001 | -0.0007 | 0.0010 | |
| dif allophilia | | Alone-Together | RIFG~LTPJ | 0.0000 | -0.0004 | 0.0003 | |
| dif allophilia | | Alone-Together | RIFG~RTPJ | -0.0003 | -0.0008 | 0.0001 | |
| *Interactions* | | | | | | | |
| dif allophilia | Int-Samegen | Alone | LIFG~RIFG | 0.0004 | -0.0002 | 0.0010 | |
| dif allophilia | Int-Samegen | Together | LIFG~RIFG | 0.0002 | -0.0003 | 0.0007 | |
| dif allophilia | Int-Samegen | Alone | LIFG~LIFG | 0.0003 | -0.0006 | 0.0011 | |
| dif allophilia | Int-Samegen | Together | LIFG~LIFG | 0.0002 | -0.0004 | 0.0009 | |
| dif allophilia | Int-Samegen | Alone | LTPJ~LTPJ | 0.0003 | -0.0005 | 0.0011 | |
| dif allophilia | Int-Samegen | Together | LTPJ~LTPJ | 0.0009 | 0.0003 | 0.0016 | No overlap |
| dif allophilia | Int-Samegen | Alone | LIFG~LTPJ | 0.0005 | -0.0002 | 0.0011 | |
| dif allophilia | Int-Samegen | Together | LIFG~LTPJ | 0.0001 | -0.0005 | 0.0006 | |
| dif allophilia | Int-Samegen | Alone | RIFG~RIFG | -0.0010 | -0.0019 | -0.0001 | No overlap |
| dif allophilia | Int-Samegen | Together | RIFG~RIFG | -0.0002 | -0.0009 | 0.0005 | |
| dif allophilia | Int-Samegen | Alone | RTPJ~RTPJ | -0.0001 | -0.0014 | 0.0012 | |
| dif allophilia | Int-Samegen | Together | RTPJ~RTPJ | 0.0013 | 0.0003 | 0.0025 | No overlap |
| dif allophilia | Int-Samegen | Alone | RIFG~LTPJ | -0.0002 | -0.0008 | 0.0004 | |
| dif allophilia | Int-Samegen | Together | RIFG~LTPJ | -0.0005 | -0.0010 | 0.0000 | |
| dif allophilia | Int-Samegen | Alone | RIFG~RTPJ | -0.0003 | -0.0010 | 0.0004 | |
| dif allophilia | Int-Samegen | Together | RIFG~RTPJ | -0.0005 | -0.0011 | 0.0001 | |
| dif allophilia | Intergen | Alone-Together | LIFG~RIFG | -0.0003 | -0.0009 | 0.0002 | |
| dif allophilia | Samegen | Alone-Together | LIFG~RIFG | -0.0005 | -0.0010 | 0.0000 | |
| dif allophilia | Intergen | Alone-Together | LIFG~LIFG | -0.0001 | -0.0008 | 0.0006 | |
| dif allophilia | Samegen | Alone-Together | LIFG~LIFG | -0.0002 | -0.0010 | 0.0006 | |
| dif allophilia | Intergen | Alone-Together | LTPJ~LTPJ | -0.0006 | -0.0013 | 0.0001 | |
| dif allophilia | Samegen | Alone-Together | LTPJ~LTPJ | 0.0000 | -0.0008 | 0.0007 | |
| dif allophilia | Intergen | Alone-Together | LIFG~LTPJ | 0.0000 | -0.0005 | 0.0005 | |
| dif allophilia | Samegen | Alone-Together | LIFG~LTPJ | -0.0004 | -0.0009 | 0.0002 | |

| Measure | Group | Task | ROI Pairs | Estimate | Lower HPD | Upper HPD | No overlap |
|---|---|---|---|---|---|---|---|
| dif allophilia | Intergen | Alone-Together | RIFG~RIFG | -0.0006 | -0.0014 | 0.0001 | |
| dif allophilia | Samegen | Alone-Together | RIFG~RIFG | 0.0002 | -0.0006 | 0.0009 | |
| dif allophilia | Intergen | Alone-Together | RTPJ~RTPJ | -0.0006 | -0.0017 | 0.0006 | |
| dif allophilia | Samegen | Alone-Together | RTPJ~RTPJ | 0.0008 | -0.0003 | 0.0020 | |
| dif allophilia | Intergen | Alone-Together | RIFG~LTPJ | 0.0001 | -0.0004 | 0.0006 | |
| dif allophilia | Samegen | Alone-Together | RIFG~LTPJ | -0.0002 | -0.0007 | 0.0004 | |
| dif allophilia | Intergen | Alone-Together | RIFG~RTPJ | -0.0002 | -0.0008 | 0.0004 | |
| dif allophilia | Samegen | Alone-Together | RIFG~RTPJ | -0.0004 | -0.0011 | 0.0002 | |
